# Three phylogenetic metrics are compatible with natural evolution of the earliest SARS-CoV-2 sequence

**DOI:** 10.64898/2026.07.08.737272

**Authors:** Jean-Noël Lorenzi, François Graner, Thomas Bigot, Etienne Decroly, Virginie Courtier-Orgogozo, Guillaume Achaz

## Abstract

Comparative analyses of coronavirus sequences can unravel important aspects of their complex evolution. Here we develop a method to infer past recombination breakpoints based on minimizing homoplasies count. We test it on three outbreak viruses (SARS-CoV-1, MERS-CoV, and SARS-CoV-2) and various chimeric coronaviruses as positive controls. We identify genomic regions evolving under distinct selective pressures. We also trace possible signs of human-made manipulations using metrics such as synonymous and non-synonymous mutation rates, codon usage and insertion patterns. Our pipeline appears to efficiently detect synthetic sequence optimization or genome re-encoding, but does not identify chimeras of natural viruses. Unlike in positive controls, with our method no signal of human-made manipulation is detected in SARS-CoV-1, MERS-CoV, nor SARS-CoV-2.

---

Understanding the origin of SARS-CoV-2, which was first detected in late 2019 in Wuhan, China (*1, 2*), remains an important question with significant societal and public-health implications. This task is challenging since coronaviruses harbor mosaic genomes shaped by multiple recombination events with closely related viral lineages (*3*). Viruses with the highest genomic similarity to SARS-CoV-2 (96% of identity) were identified in wild horseshoe bats inhabiting Southeast Asia caves (*2, 4*), suggesting a zoonotic origin. However, the progenitor of the pandemic and the pathways by which the virus evolved to infect humans, was transported, and ultimately gave rise to the initial human infections are still unresolved. Several plausible hypotheses have been proposed (*5, 6*), including zoonotic processes independent of research activities and research-related pathways (*7*–*10*). In-depth analyses of samples from the Huanan market did not point to a definitive answer (*11*–*13*). With respect to SARS-CoV-2 sequence evolution, several authors suggested a natural origin (*9, 14, 15*) and others a laboratory manipulation (*16*). For several recent outbreaks, e.g., cholera in Haiti (*17*) and bovine bluetongue in France (*18*), the pathogen sequence was found to be >99.8% identical to other sequences sampled too far in space (Haiti *vs* Nepal) or time (2015 *vs* 2008) for human-independant evolution, thus hinting at human-related pathways, namely human migration and frozen bull sperm, respectively. As far as we know, there is currently no method to test for potential human-related pathways when the sequence of the new pathogen is less similar to available sequences.

For viruses that undergo frequent recombinations, such as coronaviruses (*3*), whole-genome analyses are not suitable for inferring their past evolution because individual genomic regions are likely to harbour distinct evolutionary signals. Accurate reconstruction of viral evolution therefore requires the identification of recombination breakpoints, and the partitioning of the genome into independently evolving segments. Analyzing these regions separately can help reconstruct their distinct evolutionary histories and thus provides a more robust understanding of the evolutionary origins and trajectories of recombinant viruses.

Here we develop a method to delineate recombination breakpoints, by minimizing (through simulated annealing) the number of homoplasies. We analyze SARS-CoV-2 along with two other coronaviruses which caused outbreaks, SARS-CoV-1 in China in 2003 (*19*), and MERS in the Middle-East in 2012 (*20*). We identify genomic segments that evolved under distinct selective pressures.

For each segment, we test for three types of human-made manipulations: long-term cold storage, serial passage and chimeric virus construction containing fragments from other related viruses with or without codon optimization. Each of them, if strong or long enough, might leave prints in patterns of *i*) non-synonymous and synonymous substitutions, *ii*) codon usage and *iii*) insertions-deletions (indels). Cold storage could prevent mutation accumulation and lead to a low rate of substitutions; serial passage and intense selection could affect the relative rate of substitutions and codon usage by accelerating amino-acid replacements; creation of chimeric viruses by combining segments from diverse viruses could affect substitution rates, codon usage and indel rates. While detecting an outlier pattern may hint at human-made manipulations, finding no trace could be due to fine manipulations that are not detectable. To validate our method, we use as positive controls three *in silico* chimeric coronaviruses carrying distinct Spike protein sequences: CoV-2-S^edit^ where the spike gene of a SARS-CoV-2 sequence was replaced with another divergent spike gene, WIV-BANAL-20-236^nat^ where the spike gene of the SARS-CoV-1 sequence was replaced with the spike gene of a close relative, and finally WIV-BANAL-20-236^opt^ where the spike gene was replaced and further optimized for codon usage.

For the three outbreaks (SARS-CoV-1, MERS and SARS-CoV-2), we first search for all complete viral genomes that were more than 88% identical genome-wide to the initial sequence from public databases (Genbank and GISAID). For SARS-CoV-1 and SARS-CoV-2, when a group of sequences is >99% identical to each other, only one representative of the group is kept for the analysis. For MERS, all sequences are included. We thus collect 18 closely related sequences for SARS-CoV-1, 13 for MERS and 16 for SARS-CoV-2 (tables S1-3). The closely related SARS-CoV-2 related viruses were sampled in bats, camels and pangolins; and none in humans. The resulting SARS-CoV-1 and SARS-CoV-2 datasets encompass sequences evenly distributed in terms of percentage of nucleotide identity, whereas most sequences of the MERS dataset display 99% identity to MERS (Fig. S1, data S1). Since sequences of chimeric coronaviruses are not present in GenBank, we test three chimeric viruses that we reconstruct *in silico*: CoV-2-S^edit^, which corresponds to the Wuhan-1 SARS-CoV-2 reference sequence, in which the native Spike sequence is replaced by an edited SARS-CoV-2 Spike sequence (serendipitously found in a bacterial NCBI dataset, see Materials and Methods), WIV-BANAL-20-236^nat^ and BANAL-20-236^opt^, which correspond, respectively, to the native and a modified (with codons *in silico* optimized for human expression) Spike sequence from the Laos BANAL-20-236 virus inserted into the SARS-CoV WIV1 backbone.

For each dataset nucleotide sequences are aligned using our recent Coding / Non-Coding Aligner (CNCA) (*21*), which guarantees that coding regions are aligned by codons (Fig. S2, data S2). Visual inspection (Fig. 1A, Fig. S3) reveals that the closest relative to the human outbreak virus varies depending on genome position, a pattern consistent with the fact that these viruses frequently recombine (*3*). To partition the genome into segments with no detectable recombination (named *segments* hereafter), we devise a new robust and efficient method that can infer the number and positions of past recombination breakpoints from a sequence alignment. In brief, our approach minimizes the number of sites bearing apparent homoplasies. Indeed, past recombination events create homoplasies in phylogenetic trees inferred from full sequences (*22*). We combine a simulated annealing algorithm with Akaike and Bayesian information criteria (AIC, BIC) to control the likelihood improvement (see Materials and Methods, Figs. S4-S7). Our algorithm delimits 16 segments (and thus 15 breakpoints) for SARS-CoV-1, 11 for MERS and 25 for SARS-CoV-2 (Fig. S4). We then compare our pipeline to the commonly used GARD method (*23*), which splits the alignment into segments by maximizing the likelihood product of all local trees, tuning the number of breakpoints and their positions. For SARS-CoV-2, GARD comes up with 59 segments, that is twice more than our pipeline, for a similar number of homoplasies (Fig. S8). Interestingly, breakpoint positions (Fig. S9) are similar for both methods: less than 200-nt difference is found for 10/15 breakpoints for SARS-CoV-1, 2/10 for MERS and 20/24 for SARS-CoV-2.

**Figure 1:**
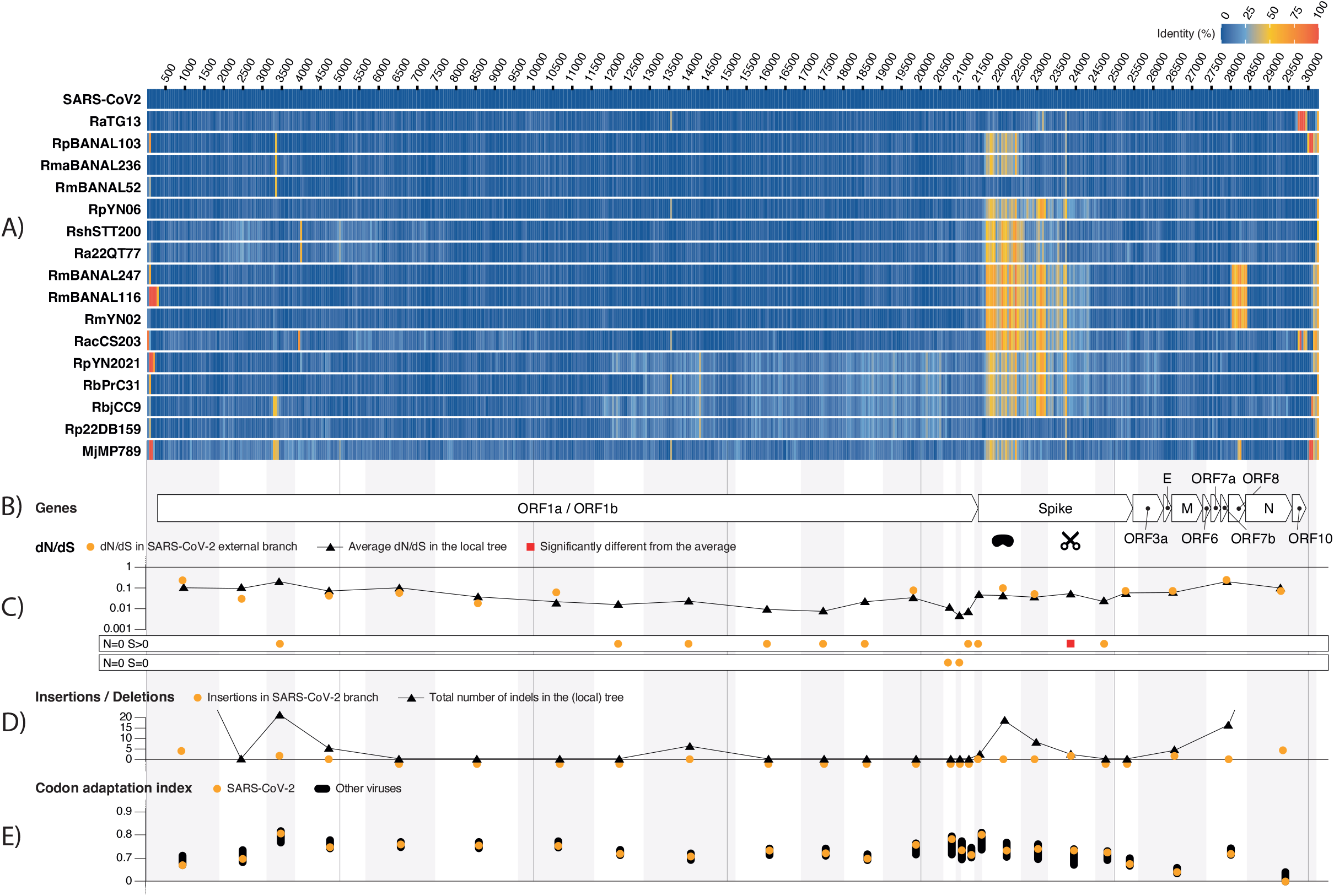
No trace of human-made manipulation is detected in SARS-CoV-2. (**A**) SARS-CoV-2 alignment with its 16 closest relatives. Identity to the SARS-CoV-2 genome is color coded with a sliding window of size 50 nt. (**B**) Localisation of SARS-CoV-2 genes in the alignment. The alternating vertical gray and white stripes indicate the 25 segments with no detectable recombination. (**C**) Statistical tests based on the *dN/dS* ratio assessing whether the external branch leading to SARS-CoV-2 (orange discs and red square) differs from the average ratio in the local phylogeny (black triangles). One significant test (red square) points to an especially conserved region of the spike gene that contains the furin cleavage site (FCS). (**D**) The number of insertions and deletions (indels) in the phylogeny of each segment (black triangles) as well as indels specific to SARS-CoV-2 (orange discs). (**E**) Local codon usage (measured by CAI) of each segment for each virus. The CAI range for all viruses is indicated by a thick black interval whereas the CAI of the SARS-CoV-2 is indicated by orange disks.

For each inferred segment of the three alignments, we compute a local maximum-likelihood phylogenetic tree that represents the local evolutionary history of the segment (Figs. S10, S11). We pay special attention to segments containing the Receptor Binding Domain (RBD) for the three viruses and the 12-nt insertion of the Furin Cleavage Site (FCS) for SARS-CoV-2, as they are key to viral infectivity (*24*). For each segment we generate a *codon-only* segment, concatenating all its codons, discarding non-coding sequences and incomplete codons.

Accounting for local phylogeny, for each *codon-only* segment we compute average Synonymous (*dS*) and Non-synonymous (*dN*) mutation rates for the whole phylogeny using PAML model 0 (*25*). We compare these rates with inferred specific rates (PAML branch model 2) of each external branch leading to a sampled virus. Results show no excess of synonymous or non-synonymous mutations in the external branch leading to SARS-CoV-1, MERS or SARS-CoV-2, contrarily to chimeric viruses which show a high rate of both mutation types (Figs. S12, S13). Furthermore, phylogenetic average d*N*/d*S* ratios are all below 1, with values ranging in [0.02-0.2] for SARS-CoV-1, [0.1-0.5] for MERS and [0.003-0.2] for SARS-CoV-2, a pattern compatible with natural evolution that is not observed for chimeric sequences, where typically inserted spike genes show a dN/dS ratio significantly larger than 1 (Fig. 1C, Figs. S14, S15). This excludes scenarios where one or more segments are subject to recurrent positive selection in SARS-CoV-2. Furthermore, only 17/850 statistical tests (2%) show an external branch statistically different (*p* < 0.001) from the average ratio. No test is significant for any segment of SARS-CoV-1 nor for MERS. For SASR-Cov-2, only the 20th segment (positions 23,252 to 24,481) points to an especially conserved segment of the spike gene (30 synonymous and 0 non-synonymous mutations since the ancestor). This conserved region corresponds to the domain located downstream of the receptor-binding domain of the Spike protein, including the S1/S2 and S2′ cleavage sites, which are essential for furin- and TMPRSS2-mediated proteolytic processing of the Spike protein and for virus–cell fusion (*26*). Using fewer but larger segments for the SARS-CoV-2 dataset (14 segments instead of 25) or specifically merging all small regions of the spike gene into a single segment also result in an absence of statistically supported positive selection, excluding a lack of power caused by too small segments (Fig. S14). Finally, none of the 59 GARD segments of SARS-CoV-2 show a pattern significantly different from other terminal branches either.

For each virus and each *coding-restricted* segment, we compute the Codon Adaptation Index metric, CAI (16), to evaluate whether the codon usage of a segment (the query sequence) differs from the codon usage of the rest of its genome (using all other *coding-restricted* segments of the same virus as reference usage). We observe that the spike gene of CoV-2-S^edit^ and WIV-BANAL-20-236^opt^ chimeric sequences displays obvious patterns of abnormal CAI, which is expected because synthetic replacement of a part of a genome with random codons preserving the amino-acids or with codon optimisation appears as clear outliers (Fig. 1E, Figs. S16, S17). Interestingly, the CAI pattern for the chimeric WIV-BANAL-20-236^opt^ also points (but to a lesser extent) to an abnormal CAI pattern. No segment of SARS-CoV-1, MERS or SARS-Cov-2, displays an abnormal codon usage in any of the viruses. We then further explore for each segment of each virus whether codon usage has changed from its most direct ancestor. To do so, we compute a new *Delta*-CAI metrics, which indicates whether codon usage has become more similar (positive *Delta-*CAI) or less similar (negative *Delta-*CAI) from that of the rest of the genome. This metrics shows that CAI typically does not change much (at most +/-0.05) since the most recent ancestor, with few exceptions (Figs. S18, S19). None of the codon usage for the three query viruses (SARS-CoV-1, MERS or SARS-CoV-2) show a strong shift in CAI. For SARS-CoV-2, the strongest shift is observed for the 17th segment (positions 21,350 to 21,631) for which the CAI evolved toward more homogeneity with the rest of the genome.

Finally, we map insertions and deletion events in the local trees using parsimony. The total number of both insertions and deletions is especially elevated in the first and last segments (Fig. 1D, Figs. S20-S22). Alignments that correspond to these two segments appear to be less reliable than the rest of the alignment (overall poor alignment score and many gaps). The high number of indels at the two edges of genomes in all viruses probably reflects loose evolutionary constraints in these regions. Besides these edges, we notice a relatively higher number of both insertions and deletions in the spike gene of the SARS-CoV-1 and SARS-CoV-2 sets, pointing to relatively frequent insertions inside these regions of the spike genes. In addition, a 6-nt deletion event occurred just at the left edge of the 12-nt FCS insertion in the branch of the phylogeny leading to the subtree of four other viruses (RmBANAL247, RacS203, RmYN02 and RmBANAL116), suggesting that this precise region experienced at least two independent indels, the 6-nt deletion being natural.

In summary, we devise a method to test for evolutionary abnormalities in viral genome sequences. The method first relies on partitioning the aligned genomes into a series of non-recombined segments, each possessing a local phylogenetic tree. To this purpose, we develop an efficient method to detect recombination breakpoints that minimizes the number of apparent homoplasies. Using the local phylogenetic tree of each segment, we evaluate the relative rate of amino-acid replacement, the homogeneity of codon usage in each segment compared to the rest of the genome and the rate of insertion/deletion events.

Based on this approach, *in silico* chimeric viruses with optimized codons reveal significantly different rates of *dN* that pinpoint to human-made manipulations. However, silico chimeric sequences made by mixing pieces of natural viruses, as well as SARS-CoV-1, MERS or SARS-CoV-2, display no clear deviation from natural evolution.

Our method has several limitations. On the methodological side, the shorter the recombination segments, the more difficult it is possible to estimate phylogenetic trees and the various metrics (*dN, dS*, CAI, indel rates) because counts of evolutionary events are very small or vanish. Furthermore, our codon usage test compares usage of whole segments against the whole genome as a reference and this oversimplification may obscure subtle or localized codon optimization. With a suitable robust molecular clock, one could further test whether any branch length is abnormal given the sampling dates. On the application side, our present approach does not allow us to test for certain laboratory-related scenarios, such as recent sampling in the wild, short-term cold storage, cell culture without selection or even chimeras made of natural viruses. In any case, certain scenarios will probably be impossible to test based on sequence analyses. They should rather be analyzed by other approaches such as epidemiological studies or scoring risk criteria (27, 28).

Overall, our study represents a first attempt in designing a sequence-based method to detect traces of human-made manipulation. The COVID-19 pandemic and the increasing number of high-security biological laboratories studying dangerous pathogens worldwide (*29*) have raised public awareness of the risk of lab accidents. Our method can be extended to detect traces of human-made manipulation in any pathogen sequence. In particular, it could help examine future outbreaks that will likely fuel concerns and speculations on whether they have a laboratory origin. While we detect no sign that the original Wuhan-Hu-1 SARS-CoV-2 has been artificially manipulated in a lab, nor strongly selected for gain of functions through serial passages that would result in a strong signal of positive selection, it remains to be explained how this virus happened to spread in Wuhan, while its closest relatives could only be sampled thousands of kilometers away.

## Supporting information

MSA

Supp Mat

## Abbreviations

AIC,: Akaike Information Criterion
BIC,: Bayesian Information Criterion
CAI,: Codon Adaptation Index
FCS,: Furin Cleavage Site
RBD,: Receptor Binding Domain

## Acknowledgements

We thank F. Abdi for implementing the simulated annealing. The authors thank E.P.C. Rocha and R. Kulathinal for constructive comments on the manuscript.

## Authors contributions

Conceptualization: VC, GA, FG; Data curation: JNL; Formal analysis: n/a; Funding acquisition: GA, VC, FG; Investigation: JNL, GA, VC, ED, FG; Methodology: JNL, ED, TB, GA, VC, FG; Project administration: n/a; Resources: n/a; Software: JNL, TB; Supervision: GA, VC, FG; Validation: JNL, GA, VC, FG; Visualization: JNL; Writing – original draft preparation: VC, GA, ED, FG; Writing – review and editing: GA, VC, ED, FG, JNL.

## Data and materials availability

All data are available in the manuscript or the supplementary materials.

