## Supplementary material for "Three phylogenetic metrics are compatible with natural evolution of the earliest SARS-CoV-2 sequence": Supp Mat

**The PDF file includes:**
Materials and Methods
Supplementary Text
Figs. S1 to S22
Tables S1 to S3
References (35 to 57)

**Other Supplementary Materials for this manuscript include the following:**
Data S1 to S#

**Materials and Methods**

Collection of sequences

Closely-related sequences of SARS-CoV-2 (NC_045512.2), SARS-CoV-1 (AY278488.2) and MERS (NC_019843.3) were retrieved from NCBI database by gathering all the sequences annotated with the word "coronavirus" in April 2022 and by running a local BLAST with the three respective sequences as queries. Only sequences with more than 88% identity to the query were kept. For SARS-CoV-1 and SARS-CoV-2, extremely similar nucleotide sequences (more than 99% identity) were downsampled to 1 sequence chosen at random. For MERS, all sequences were kept. Coronavirus sequences published after April 2022 and until December 2023 were also added along the way. In total we used 19 sequences for SARS-CoV-1 (Table S1), 14 sequences for MERS (Table S2) and 17 sequences for SARS-CoV-2 (Table S3).

Artificial chimeric sequences

Three chimeric virus genomic sequences were generated for this study. For CoV-2-$S^{edit}$, the backbone was the Wuhan-1 SARS-CoV-2 reference sequence and the coding sequence of Spike (position 21563-25384) was replaced by another coding sequence of Spike, named here $S^{edit}$. The $S^{edit}$ sequence was serendipitously detected in a whole-genome sequencing assembly data made public on 06-JAN-2023 in NCBI, deposited under PRJNA839565, that reports isolated samples of *Pseudomonas Aeruginosa* taken from patients in 2019 in Henan, China. The $S^{edit}$ sequence was detected within the Eukaryotic expression vector pCDNA3.1 in PRJNA839565. Whereas the encoded Spike protein sequence is 100% identical to SARS-CoV-2 Spike (except that the FCS is missing), the $S^{edit}$ nucleotide sequence is only 71.8% identical to SARS-CoV-2 Spike, suggesting that this sequence may result from codon optimization. For WIV-BANAL-20-236$^{nat}$, the backbone is SARS-CoV WIV1 (Genbank FF367457) and the native Spike sequence of the

Laos coronavirus BANAL-20-236 (*30*) was inserted at BlgII sites as described in (*31*). For WIV-BANAL-20-236$^{opt}$, the backbone was SARS-CoV WIV1 (Genbank FF367457) and a codon-optimized Spike sequence of the Laos coronavirus BANAL-20-236 was inserted at BlgII sites as described in (*31*). Codon-usage optimization for human expression was performed via the IDT technology website (https://eu.idtdna.com/CodonOpt). For each of the three chimeric sequences, we ran the entire analysis simply replacing SARS-CoV-2 (for the first one) or SARS-CoV WIV1 (for the latter two) with the corresponding chimera.

Genome annotations

Annotations were retrieved from NCBI except for RmYN02, RpYN2021 and RshSTT200 which had no annotation and for which we transferred annotations from SARS-CoV-2 (NC_045512.2) with Geneious Prime 2019 after alignment. For the RmYN02 genome (EPI_ISL_412977), the *ORF8* annotation obtained by annotation transfer with Geneious generated a last 3' end codon consisting of only two nucleotides (position 28,088 and 28,089), with no stop codon. We extended the *ORF8* coding sequence up to position 28,102 which generated a stop codon and the resulting protein sequence displayed 100% protein sequence identity with the *ORF8* of BANAL-20-116 (MZ937002.1), supporting this new *ORF8* annotation for RmYN02. For the RpYN2021 genome (OK017806.1), the *Spike* annotation obtained by annotation transfer with Geneious generated a 5' end codon consisting of three nucleotides (position 21,425, 21,424 and 21,425), coding for an L. The *S* coding sequence was extended up to position 21,422, which codes for a M in the same frame.

Nucleotide alignment using CNCA

For each dataset, nucleotide sequences were aligned using our recent Coding / Non-Coding Aligner (CNCA) software, which combines information from both protein sequences and nucleotide sequences (*32*). The three resulting alignments were then visually inspected and validated. No change was made secondarily on the alignments.

New algorithm to infer non-recombinant segments

In brief, the algorithm leverages the fact that recombination creates artefactual signals of homoplasy when recombination is ignored. Homoplasy can be scored by counting the number of polymorphic sites that are not compatible with the phylogenetic tree (*i.e.* that cannot be explained by a single event for each extra allele in the tree). By recursively splitting the alignment into smaller segments, while building local parsimony trees for each corresponding segment in the alignment, the number of sites with homoplasy that are not compatible with the local tree decreases. To overcome the pitfall of over splitting the alignment into many very small segments, we used Akaike and Bayesian information criteria (AIC, BIC).

More precisely, our recursive algorithm runs as follow:

**0. Initiate with *k* = 1 segment**

We initiate our recursive algorithm with a single segment that is the full alignment of $n$ sequences with $k$=1 segment, no breakpoint. We infer the maximum likelihood tree (using a model GTR + F + Gamma) over the entire alignment and record its likelihood.

1. **Find $k$+1 "good" segments**
   a. We introduce a new random breakpoint that splits the alignment into $k$+1 segments. We infer the tree of each segment using a standard parsimony method [*parsimony ratchet* (*33*) in the R package *phangorn*] that is fast and minimizes the number of sites with apparent homoplasy. We then count $H_T$ the total number of sites that harbor homoplasy [using the R package *homoplasyFinder*] while considering the local tree for each segment of the alignment.
   b. The position of all $k$ breakpoints is then optimized by *Simulated Annealing* minimizing $H_T$, while keeping the number of segments identical. At each proposition, the algorithm randomly changes $B$ breakpoints at a time, where $B$ is a random geometric variable with mean 1/0.6 ~ 1.67. Local trees are inferred by parsimony. The acceptance ratio (probability to accept the new configuration) is given by $P_{accept} = \exp(-DH/C)$, with $DH$ the difference in $H_T$ between the current split and the proposed one, and $C$ the fluctuation allowance (or effective temperature) computed as $C = 1000/(i+1)$, for which $i$ is the iteration number (from 0 to $10^4$). The algorithm stops after $10^4$ iterations or when no further change is accepted in 2,500 subsequent propositions.

Steps 1a and 1b are performed 100 independent times but only the best segmentation of $k$+1 segments with the lowest $H_T$ is kept further.

2. **Evaluate Likelihood**
   Once the $k$ breakpoints are placed delimiting $k$+1 segments, we infer one tree for each of the corresponding segments in the alignment by Maximum Likelihood (using a model `GTR + F + Gamma`). We then compute a total likelihood for the segmentation by multiplying the likelihood of each local tree, assuming that they are all independent. Although pseudo-likelyhood is likely imperfect, this choice is in accordance with the hypothesis of independent sites of standard phylogeny using likelihood. We record it and the AIC and BIC are computed assuming that introducing each breakpoint adds $2n$-1 new parameters ($2n$-3 branches of the unrooted tree, 1 for the topology and 1 for the position of the breakpoint). Note that the GTR+F+Gamma model also adds a few parameters, whose number is difficult to estimate, so we chose not to include them in our AIC and BIC calculations.

3. **Go to Step 1** to add a new breakpoint.

The algorithm is run until 50 segments (49 breakpoints) are inferred and the solution with the lowest local minimum AIC or BIC is returned. Visual inspection of the gains in likelihood, AIC and BIC is performed.

The resulting segmentations are 16 segments for the SARS-CoV-1 alignment, 11 segments for the MERS alignment and 25 segments for the SARS-CoV-2 alignment.

To test whether too small segment size can hinder statistical detections of unexpected patterns of substitution or codon usage, we also ran the downstream analysis with a coarser grain using the 14 segments chopping for the SARS-CoV-2 alignment.

GARD segmentation

Segmentation with GARD (*23*) was done using parameters: HYPHYMPI gard --mode Faster --code Universal --type nucleotide --rv GDD --model JTT --rate-classes 4.

Rate of amino acid evolution (d*N*/d*S*)

For each segment of each dataset, we evaluated first a tree topology by maximum likelihood [with RAxML-NG (*34*), substitution model GTR+F+G] with all nucleotides. We then concatenated and phased all codons from the segment and estimated one single d*N*/d*S* ratio between the rate of non-synonymous mutations (d*N*) and the rate of synonymous mutation (d*S*) for all branches of the local segment tree (model 0 in PAML) (*25*). We then estimated, for each external branch of the tree, a second set of parameters with potentially different d*N*/d*S* ratio for the focal branch (branch-model number 2 of PAML). We then ran a chi-square Likelihood Ratio Test to compute a *p*-value for each external branch of the phylogeny, thus testing for the presence of a peculiar d*N*/d*S* ratio in the external branch leading to the different viruses. If any virus displayed an unexpected rate of amino-acid evolution in any part of the alignment, this resulted in a low *p*-value. Again a chi-square Likelihood Ratio Test was applied to test whether positive selection was supported for a given segment.

Codon Usage

To assess codon usage, like in the previous step, we trimmed the alignment to keep only the coding phase of each segment, making sure that codons were in phase. We then computed for each segment a value of Codon Adaptation Index (CAI) (Sharp & Li, 1986), using all codons of the other segments as the reference. Thus here, the CAI measured how similar was the codon usage of the focal segment compared to the rest of the genome. Then, using CODEML (*25*), a putative ancestral coding sequence was reconstructed for each node in the tree. We then computed the CAI of the most proximal ancestor of each virus and reported the difference computed as *Delta*-CAI = CAI_virus - CAI_ancestor, which measured how much the codon usage had changed since the most recent ancestor to the current virus. A positive value corresponds to mutations that tend to homogenize the CAI of the segment with the rest of the genome.

Indels

Indels are defined as regions of the alignments where a constant set of one or more viruses harbor a sequence of gaps, while others have nucleotides. Potentially overlapping indels (where the genomes containing gaps are not the same) are considered as different non-overlapping independent events. In the three alignments, indels were unambiguously delimited from the CNCA multiple sequence alignment and then mapped by parsimony onto the branches of the local segment phylogeny.

**Table S1. List of coronavirus genomes analyzed in this study that are closely related to the initial SARS sequence (AY278488.2, isolate BJ01).** "% id" indicates the percentage of nucleotide identity between the sequence and the initial SARS sequence (AY278488.2) among the aligned nucleotides. "INS" and "DEL" report, respectively, the number of different insertions (stretches of '-' in AY278488.2) and deletions (stretches of '-' in the other viruses) while comparing to SARS-CoV-1. This table contains SARS-CoV-1, and 18 bat viruses: 1 Stoliczka's trident bat ("As" = *Aselliscus stoliczkanus*), 17 from horseshoe bats (*Rhinolophus*): 13 *R. sinicus* ("Rs"), 3 *R. ferrumequinum* ("Rf"), 1 *R. pusillus* ("bRp" or "Rp").

| Accession number | Name | DB | Reference | % id | Length | INS | DEL |
|---|---|---|---|---|---|---|---|
| AY278488.2 | SARS-CoV-1 | NCBI | Qin et al. (2003)(*35*) | 100 | 29 725 | - | - |
| KY417150.1 | Rs4874 | NCBI | Hu et al. (2017)(*36*) | 96.10 | 30 311 | 13 | 5 |
| KY417146.1 | Rs4231 | NCBI | Hu et al. (2017)(*36*) | 95.93 | 29 782 | 11 | 6 |
| KY417151.1 | Rs7327 | NCBI | Hu et al. (2017)(*36*) | 95.81 | 30 307 | 10 | 5 |
| KY417152.1 | Rs9401 | NCBI | Hu et al. (2017)(*36*) | 95.76 | 29 769 | 11 | 6 |
| KC881006.1 | Rs3367 | NCBI | Ge et al. (2013)(*37*) | 95.70 | 29 792 | 12 | 5 |
| KF367457.1 | RsWIV1 | NCBI | Ge et al. (2013)(*37*) | 95.65 | 30 309 | 12 | 6 |
| KY417142.1 | As6526 | NCBI | Hu et al. (2017)(*36*) | 93.98 | 29 725 | 14 | 11 |
| KJ473816.1 | RsYN2013 | NCBI | Wu et al. (2016)(*38*) | 93.97 | 29 142 | 6 | 7 |
| KY417147.1 | Rs4237 | NCBI | Hu et al. (2017)(*36*) | 93.96 | 29 741 | 14 | 11 |
| KY417145.1 | Rf4092 | NCBI | Hu et al. (2017)(*36*) | 93.91 | 29 710 | 8 | 10 |
| KY417148.1 | Rs4247 | NCBI | Hu et al. (2017)(*36*) | 93.83 | 29 743 | 14 | 12 |
| KY417149.1 | Rs4255 | NCBI | Hu et al. (2017)(*36*) | 93.81 | 29 743 | 15 | 10 |
| KY417143.1 | Rs4081 | NCBI | Hu et al. (2017)(*36*) | 93.74 | 29 741 | 14 | 11 |
| FJ588686.1 | Rs672 | NCBI | Yuan et al. (2010)(*39*) | 93.45 | 29 059 | 14 | 12 |
| KP886808.1 | RfYNLF_31C | NCBI | Lau et al. (2015)(*40*) | 93.38 | 29 723 | 7 | 11 |
| KP886809.1 | RfYNLF_34C | NCBI | Lau et al. (2015)(*40*) | 93.37 | 29 723 | 7 | 11 |
| DQ071615.1 | bRp3 | NCBI | Li et al. (2005)(*41*) | 92.60 | 29 736 | 15 | 11 |
| KJ473815.1 | RsGX2013 | NCBI | Wu et al. (2016)(*38*) | 92.20 | 29 161 | 8 | 8 |

**Table S2. List of coronavirus genomes analyzed in this study that are closely related to the initial MERS sequence (NC_019843.3, isolate HCoV-EMC/2012).** "% id" indicates the percentage of nucleotide identity between the sequence and the initial MERS sequence (NC_019843.3) among the aligned nucleotides. "INS" and "DEL" report, respectively, the number of different insertions (stretches of '-' in NC_019843.3) and deletions (stretches of '-' in the other viruses) while comparing to MERS. This table contains MERS and 13 dromedary viruses ("Cd" = *Camelus dromedarius*).

| Accession number | Name | DB | Reference | % id | Length | INS | DEL |
|---|---|---|---|---|---|---|---|
| NC_019843.3 | MERS | NCBI | Zaki et al. (2012)(*42*) | 100 | 30 119 | 0 | 0 |
| MN758607.1 | Cd_KFU-HKU-P | NCBI | Chu et al. (2019) (no publication) | 99.65 | 30 089 | 1 | 0 |
| KT368824.1 | Cd_F13A | NCBI | Sabir et al. (2015)(*43*) | 99.60 | 30 115 | 2 | 0 |
| KJ477102.1 | Cd_NRCE-HKU20 | NCBI | Chu et al. (2014)(*44*) | 99.51 | 29 908 | 2 | 0 |
| KX108943.1 | Cd_D998_15 | NCBI | Lau et al. (2016)(*45*) | 99.46 | 30 088 | 3 | 0 |
| MG923469.1 | Cd_HKU213 | NCBI | Chu et al. (2018)(*46*) | 99.45 | 29 421 | 5 | 0 |
| MG923473.1 | Cd_HKU697 | NCBI | Chu et al. (2018)(*46*) | 99.42 | 29 374 | 8 | 0 |
| OP712624.1 | Cd_NC4713 | NCBI | Kandeil et al. (2022) (no publication) | 99.40 | 30 108 | 2 | 1 |
| OP866280.1 | Cd_CAC4787 | NCBI | Zhou et al. (2023)(*47*) | 99.32 | 30 044 | 4 | 0 |
| OP866291.1 | Cd_CAC11181 | NCBI | Zhou et al. (2023)(*47*) | 99.25 | 29 998 | 4 | 0 |
| MF598718.1 | Cd_415915_W4 | NCBI | Queen et al. (2017)(*48*) | 99.24 | 30 123 | 1 | 1 |
| OP866287.1 | Cd_CAC9670 | NCBI | Zhou et al. (2023) (*47*) | 99.21 | 30 078 | 4 | 1 |
| MF598628.1 | Cd_B36_2015 | NCBI | Queen et al. (2017)(*48*) | 97.01 | 30 123 | 1 | 1 |
| MF598642.1 | Cd_B51_2015 | NCBI | Queen et al. (2017) (*48*) | 88.89 | 30 123 | 1 | 1 |

**Table S3. List of coronavirus genomes analyzed in this study that are closely related to the initial SARS-CoV-2 sequence (NC_045512.2, isolate Wuhan-Hu-1).** "% id" indicates the percentage of nucleotide identity between the sequence and the initial SARS-CoV-2 sequence (NC_045512.2) among the aligned nucleotides. "INS" and "DEL" report, respectively, the number of different insertions (stretches of '-' in NC_045512.2) and deletions (stretches of '-' in the other viruses) while comparing to SARS-CoV-2. This table contains SARS-CoV-2, 1 virus from Sunda pangolin ("Ms" = *Manis javanica*), and 15 from horseshoe bats (*Rhinolophus*): 4 *R. malayanus* ("Rm"), 4 *R. pusillus* ("Rp"), 2 *R. affinis* ("Ra"), 2 *R. blythi* ("Rb"), 1 *R. marshalli* ("Rma"), 1 *R. shameli* ("Rsh"), 1 *R. acuminatus* ("Rac").

| Accession number | Name | DB | Reference | % id | Length | INS | DEL |
|---|---|---|---|---|---|---|---|
| NC_045512.2 | SARS-CoV2 | NCBI | Wu et al. (2020)(*1*) | 100.00 | 29 903 | - | - |
| MZ937000.1 | RmBANAL52 | NCBI | Temmam et al. (2022)(*4*) | 96.96 | 29 838 | 4 | 0 |
| MN996532.2 | RaTG13 | NCBI | Zhou et al. (2020)(*24*)) | 96.15 | 29 855 | 7 | 2 |
| MZ937001.1 | RpBANAL103 | NCBI | Temmam et al. (2022)(*23*) | 96.08 | 29 632 | 8 | 2 |
| MZ937003.2 | RmaBANAL236 | NCBI | Temmam et al. (2022)(*23*) | 96.02 | 29 844 | 8 | 2 |
| MZ081381.1 | RpYN06 | NCBI | Zhou et al. (2021)(*25*) | 94.80 | 29 793 | 12 | 4 |
| EPI_ISL_412977 | RmYN02 | GISAID | Zhou et al. (2020)(*26*) | 93.80 | 29 671 | 19 | 6 |
| MZ937002.1 | RmBANAL116 | NCBI | Temmam et al. (2022)(*23*) | 93.68 | 29 437 | 19 | 6 |
| MZ937004.1 | RmBANAL247 | NCBI | Temmam et al. (2022)(*23*) | 93.67 | 29 645 | 19 | 6 |
| EPI_ISL_852605 | RshSTT200 | GISAID | Delaune et al. (2021)(*27*) | 93.14 | 29 793 | 13 | 2 |
| OR233324.1 | Ra22QT77 | NCBI | Hassanin et al. (2024)(*28*) | 92.77 | 29 751 | 13 | 2 |
| OR233302.1 | Rp22DB159 | NCBI | Hassanin et al. (2024)(*28*) | 92.19 | 29 823 | 3 | 0 |
| MW251308.1 | RacCS203 | NCBI | Wacharapluesadee et al. (2021)(*29*) | 91.68 | 29 832 | 18 | 7 |
| MW703458.1 | RbPrC31 | NCBI | Li et al. (2021)(*54*) | 91.02 | 29 765 | 12 | 5 |
| MT121216.1 | MjMP789 | NCBI | Liu et al. (2020)(*55*) | 90.26 | 29 521 | 10 | 6 |
| OK017806.1 | RpYN2021 | NCBI | Wu et al. (2023)(*56*) | 90.10 | 29 617 | 11 | 3 |
| OK287355.1 | RbJCC9 | NCBI | Xia et al. (2021)<br>(no publication)<br>Li et al. 2024(*57*) | 89.55 | 29 694 | 14 | 3 |

45. S. K. P. Lau, R. Wernery, E. Y. M. Wong, S. Joseph, A. K. L. Tsang, N. A. G. Patteril, S. K. Elizabeth, K.-H. Chan, R. Muhammed, J. Kinne, K.-Y. Yuen, U. Wernery, P. C. Y. Woo, Polyphyletic origin of MERS coronaviruses and isolation of a novel clade A strain from dromedary camels in the United Arab Emirates. *Emerging Microbes & Infections* **5**, 1–9

(2016).

46. D. K. W. Chu, K. P. Y. Hui, R. A. P. M. Perera, E. Miguel, D. Niemeyer, J. Zhao, R. Channappanavar, G. Dudas, J. O. Oladipo, A. Traoré, O. Fassi-Fihri, A. Ali, G. F. Demissié, D. Muth, M. C. W. Chan, J. M. Nicholls, D. K. Meyerholz, S. A. Kuranga, G. Mamo, Z. Zhou, R. T. Y. So, M. G. Hemida, R. J. Webby, F. Roger, A. Rambaut, L. L. M. Poon, S. Perlman, C. Drosten, V. Chevalier, M. Peiris, MERS coronaviruses from camels in Africa exhibit region-dependent genetic diversity. *Proc. Natl. Acad. Sci. U.S.A.* **115**, 3144–3149 (2018).

47. Z. Zhou, A. Ali, E. Walelign, G. F. Demissie, I. El Masry, T. Abayneh, B. Getachew, P. Krishnan, D. Y. M. Ng, E. Gardner, Y. Makonnen, E. Miguel, V. Chevalier, D. K. Chu, R. T. Y. So, S. Von Dobschuetz, G. Mamo, L. L. M. Poon, M. Peiris, Genetic diversity and molecular epidemiology of Middle East Respiratory Syndrome Coronavirus in dromedaries in Ethiopia, 2017–2020. *Emerging Microbes & Infections* **12**, e2164218 (2023).

48. M. F. Yusof, K. Queen, Y. M. Eltahir, C. R. Paden, Z. M. A. H. Al Hammadi, Y. Tao, Y. Li, A. I. Khalafalla, M. Shi, J. Zhang, M. S. A. E. Mohamed, M. H. Abd Elaal Ahmed, I. A. Azeez, O. K. Bensalah, Z. S. Eldahab, F. I. Al Hosani, S. I. Gerber, A. J. Hall, S. Tong, S. S. Al Muhairi, Diversity of Middle East respiratory syndrome coronaviruses in 109 dromedary camels based on full-genome sequencing, Abu Dhabi, United Arab Emirates. *Emerging Microbes & Infections* **6**, 1–10 (2017).

49. H. Zhou, J. Ji, X. Chen, Y. Bi, J. Li, Q. Wang, T. Hu, H. Song, R. Zhao, Y. Chen, M. Cui, Y. Zhang, A. C. Hughes, E. C. Holmes, W. Shi, Identification of novel bat coronaviruses sheds light on the evolutionary origins of SARS-CoV-2 and related viruses. *Cell*, S0092867421007091 (2021).

50. H. Zhou, X. Chen, T. Hu, J. Li, H. Song, Y. Liu, P. Wang, D. Liu, J. Yang, E. C. Holmes, A. C. Hughes, Y. Bi, W. Shi, A Novel Bat Coronavirus Closely Related to SARS-CoV-2 Contains Natural Insertions at the S1/S2 Cleavage Site of the Spike Protein. *Current Biology* **30**, 2196-2203.e3 (2020).

51. D. Delaune, V. Hul, E. A. Karlsson, A. Hassanin, T. P. Ou, A. Baidaliuk, F. Gámbaro, M. Prot, V. T. Tu, S. Chea, A novel SARS-CoV-2 related coronavirus in bats from Cambodia. *Nature communications* **12**, 6563 (2021).

52. A. Hassanin, V. T. Tu, T. Görföl, L. Q. Ngon, P. V. Pham, C. T. Hang, T. A. Tuan, M. Prot, E. Simon‐Lorière, G. Kemenesi, G. E. Tóth, L. Moulin, S. Wurtzer, Phylogeography of horseshoe bat sarbecoviruses in Vietnam and neighbouring countries. Implications for the origins of SARS‐CoV and SARS‐CoV‐2. *Molecular Ecology* **33**, e17486 (2024).

53. S. Wacharapluesadee, C. W. Tan, P. Maneeorn, P. Duengkae, F. Zhu, Y. Joyjinda, T. Kaewpom, W. N. Chia, W. Ampoot, B. L. Lim, K. Worachotsueptrakun, V. C.-W. Chen, N. Sirichan, C. Ruchisrisarod, A. Rodpan, K. Noradechanon, T. Phaichana, N. Jantarat, B. Thongnumchaima, C. Tu, G. Crameri, M. M. Stokes, T. Hemachudha, L.-F. Wang, Evidence for SARS-CoV-2 related coronaviruses circulating in bats and pangolins in Southeast Asia. *Nat Commun* **12**, 972 (2021).

54. L. Li, J. Wang, X. Ma, X. Sun, J. Li, X. Yang, W. Shi, Z. Duan, A novel SARS-CoV-2 related coronavirus with complex recombination isolated from bats in Yunnan province,

China. *Emerging Microbes & Infections* **10**, 1683–1690 (2021).

55. P. Liu, J.-Z. Jiang, X.-F. Wan, Y. Hua, L. Li, J. Zhou, X. Wang, F. Hou, J. Chen, J. Zou, J. Chen, Are pangolins the intermediate host of the 2019 novel coronavirus (SARS-CoV-2)? *PLoS Pathog.* **16**, e1008421 (2020).

56. Z. Wu, Y. Han, Y. Wang, B. Liu, L. Zhao, J. Zhang, H. Su, W. Zhao, L. Liu, S. Bai, A comprehensive survey of bat sarbecoviruses across China in relation to the origins of SARS-CoV and SARS-CoV-2. *National Science Review* **10**, nwac213 (2023).

57. Q. Li, Y. Hou, B. Huang, X. Le, B. Wang, X. Xia, Identification and genetic characterization of five novel bat coronaviruses from Yunnan, China. *BMC Vet Res* **20**, 466 (2024).
