## Supplementary material for "Three phylogenetic metrics are compatible with natural evolution of the earliest SARS-CoV-2 sequence": Supp Figs

### Three phylogenetic quantifiers are compatible with natural evolution for the initial SARS-CoV-2 sequence

#### Supplementary Figures

Figure S1. Percentage of nucleotide identity relative to the reference initial sequence for all the closely-related viruses selected for analysis for each dataset. Percentage of nucleotide identity was computed from pairwise aligned nucleotides relative to the AY278488.2 reference sequence for the SARS-CoV-1 dataset (A), relative to the NC_019843.3 reference sequence for the MERS dataset (B), and relative to the NC_045512.2 reference sequence for the SARS-CoV-2 dataset (C). See tables S1-S3 for numerical values.

(A)

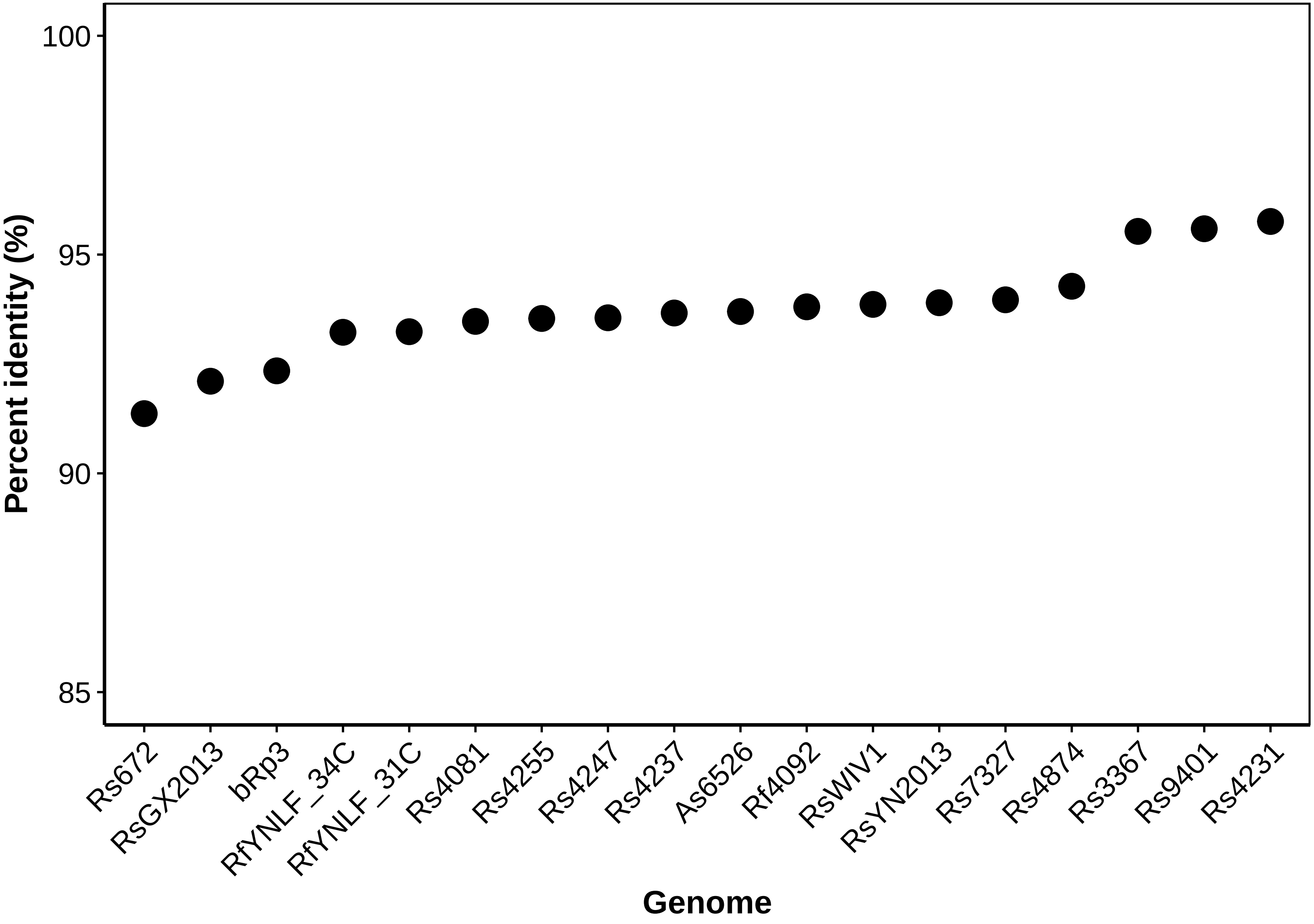
Percent identity (%)
100
95
90
85
Rs672
RsGX2013
bRp3
RfYNLF_34C
RfYNLF_31C
Rs4081
Rs4255
Rs4247
Rs4237
As6526
Rf4092
RsWIV1
RsYN2013
Rs7327
Rs4874
Rs3367
Rs9401
Rs4231
Genome

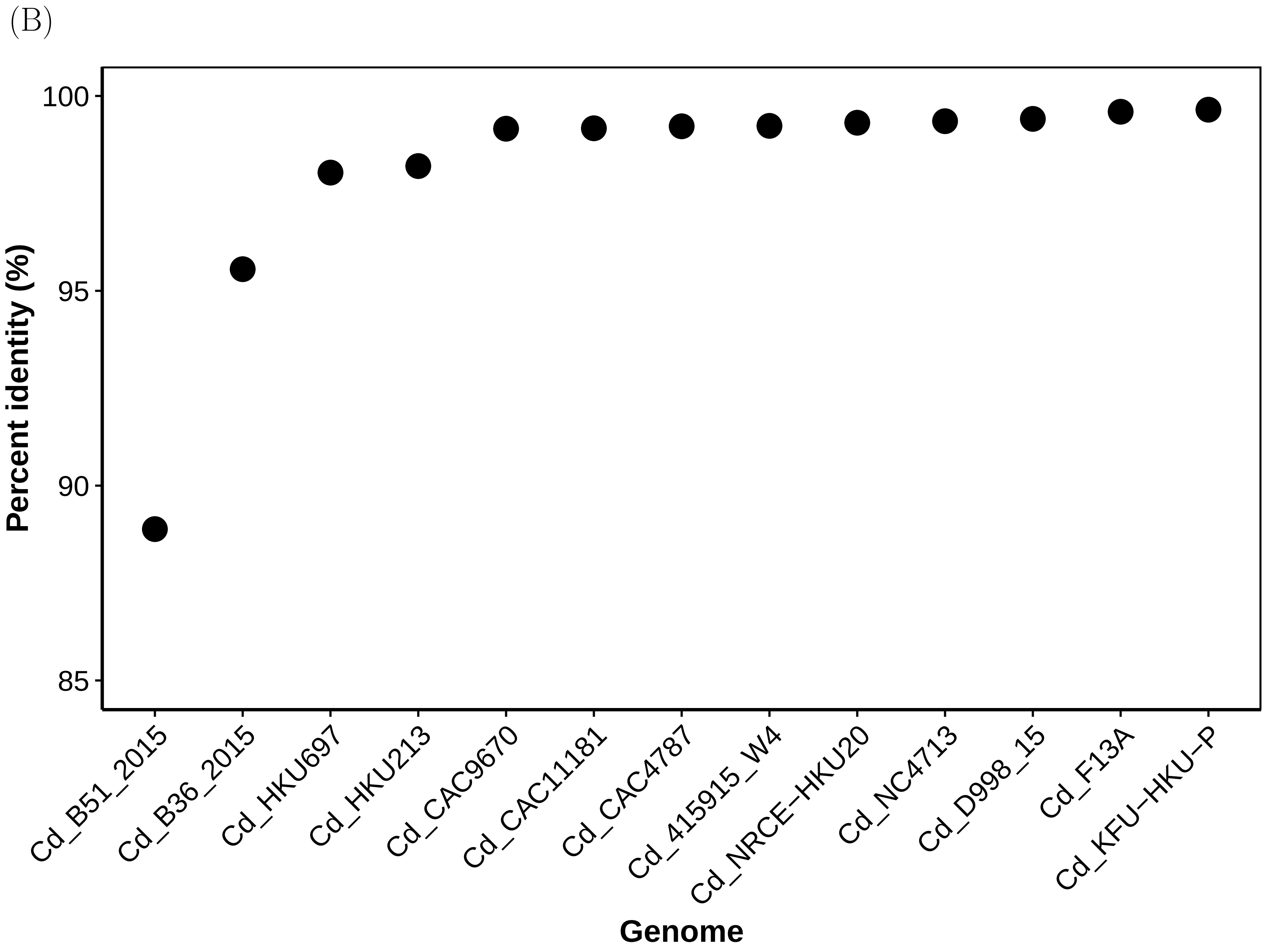

(B)
Percent identity (%)
100
95
90
85
Cd_B51_2015
Cd_B36_2015
Cd_HKU697
Cd_HKU213
Cd_CAC9670
Cd_CAC11181
Cd_CAC4787
Cd_415915_W4
Cd_NRCE−HKU20
Cd_NC4713
Cd_D998_15
Cd_F13A
Cd_KFU−HKU−P
Genome

(C)

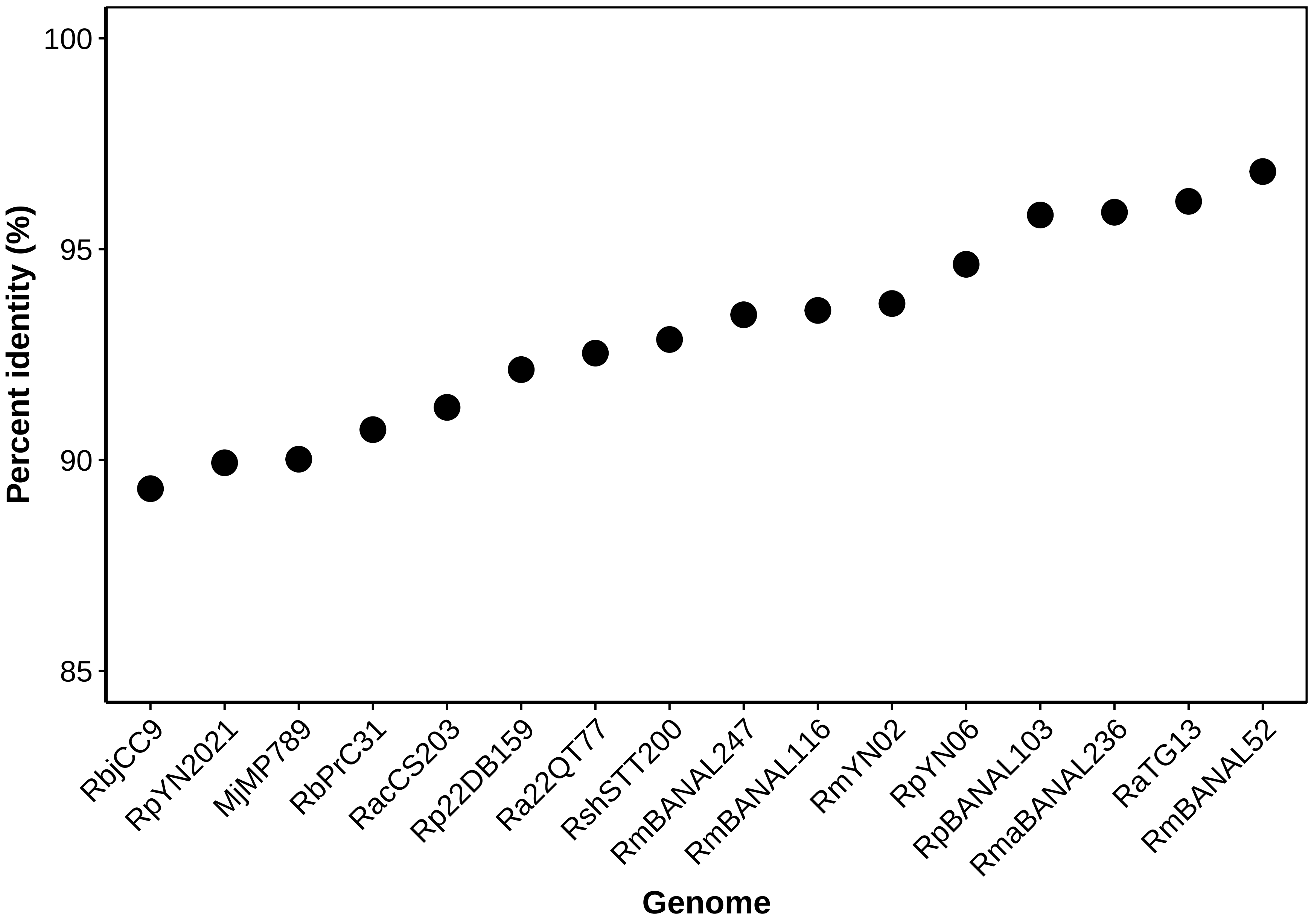

Percent identity (%)
100
95
90
85
RbjCC9
RpYN2021
MjMP789
RbPrC31
RacCS203
Rp22DB159
Ra22QT77
RshSTT200
RmBANAL247
RmBANAL116
RmYN02
RpYN06
RpBANAL103
RmaBANAL236
RaTG13
RmBANAL52
Genome

Figure S2. Circular representation of CNCA multiple sequence alignments of the coronavirus sequences. (A) SARS-CoV-1, (B) MERS and (C) SARS-CoV-2. Numbers outside the disc indicate nucleotide positions. the focal virus is always on the external circle. Colors represent the percentage of nucleotide mismatch of each virus relative to the focal virus and over a window size of how 50 nt.

(A)

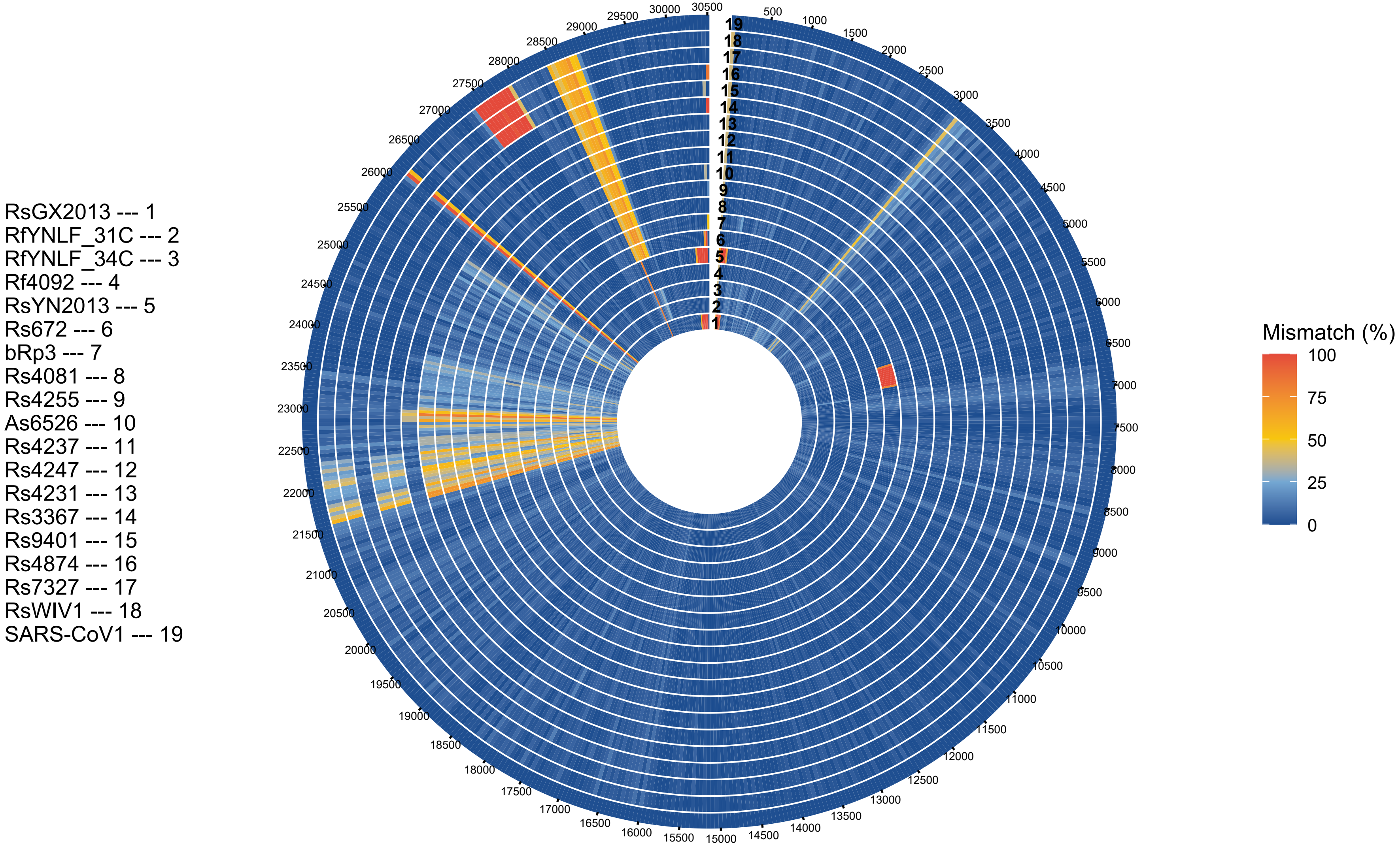
RsGX2013 --- 1
RfYNLF_31C --- 2
RfYNLF_34C --- 3
Rf4092 --- 4
RsYN2013 --- 5
Rs672 --- 6
bRp3 --- 7
Rs4081 --- 8
Rs4255 --- 9
As6526 --- 10
Rs4237 --- 11
Rs4247 --- 12
Rs4231 --- 13
Rs3367 --- 14
Rs9401 --- 15
Rs4874 --- 16
Rs7327 --- 17
RsWIV1 --- 18
SARS-CoV1 --- 19
500
1000
1500
2000
2500
3000
3500
4000
4500
5000
5500
6000
6500
7000
7500
8000
8500
9000
9500
10000
10500
11000
11500
12000
12500
13000
13500
14000
14500
15000
15500
16000
16500
17000
17500
18000
18500
19000
19500
20000
20500
21000
21500
22000
22500
23000
23500
24000
24500
25000
25500
26000
26500
27000
27500
28000
28500
29000
29500
30000
30500
Mismatch (%)
100
75
50
25
0

(B)

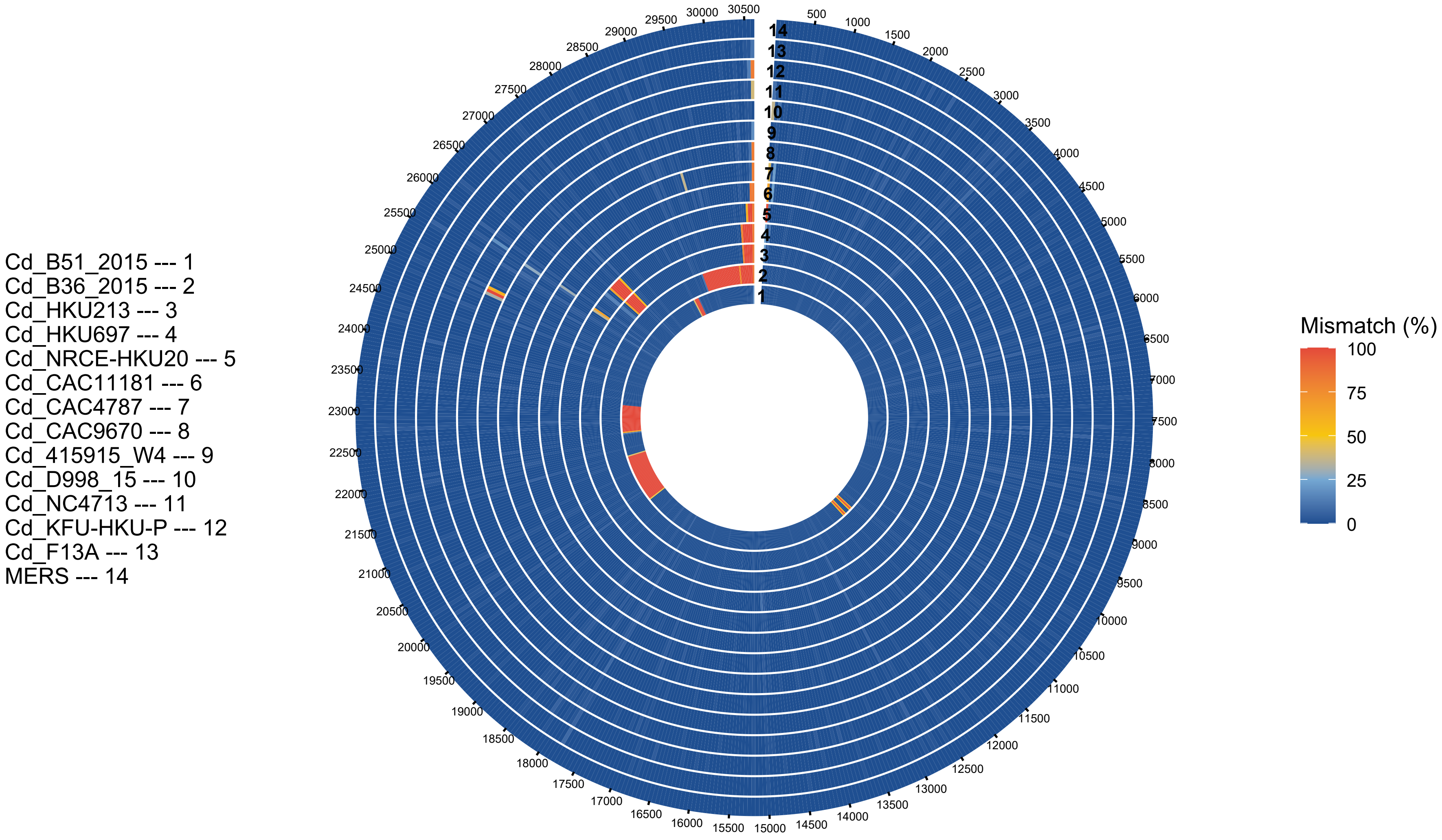
Cd_B51_2015 --- 1
Cd_B36_2015 --- 2
Cd_HKU213 --- 3
Cd_HKU697 --- 4
Cd_NRCE-HKU20 --- 5
Cd_CAC11181 --- 6
Cd_CAC4787 --- 7
Cd_CAC9670 --- 8
Cd_415915_W4 --- 9
Cd_D998_15 --- 10
Cd_NC4713 --- 11
Cd_KFU-HKU-P --- 12
Cd_F13A --- 13
MERS --- 14
Mismatch (%)
100
75
50
25
0

(C)

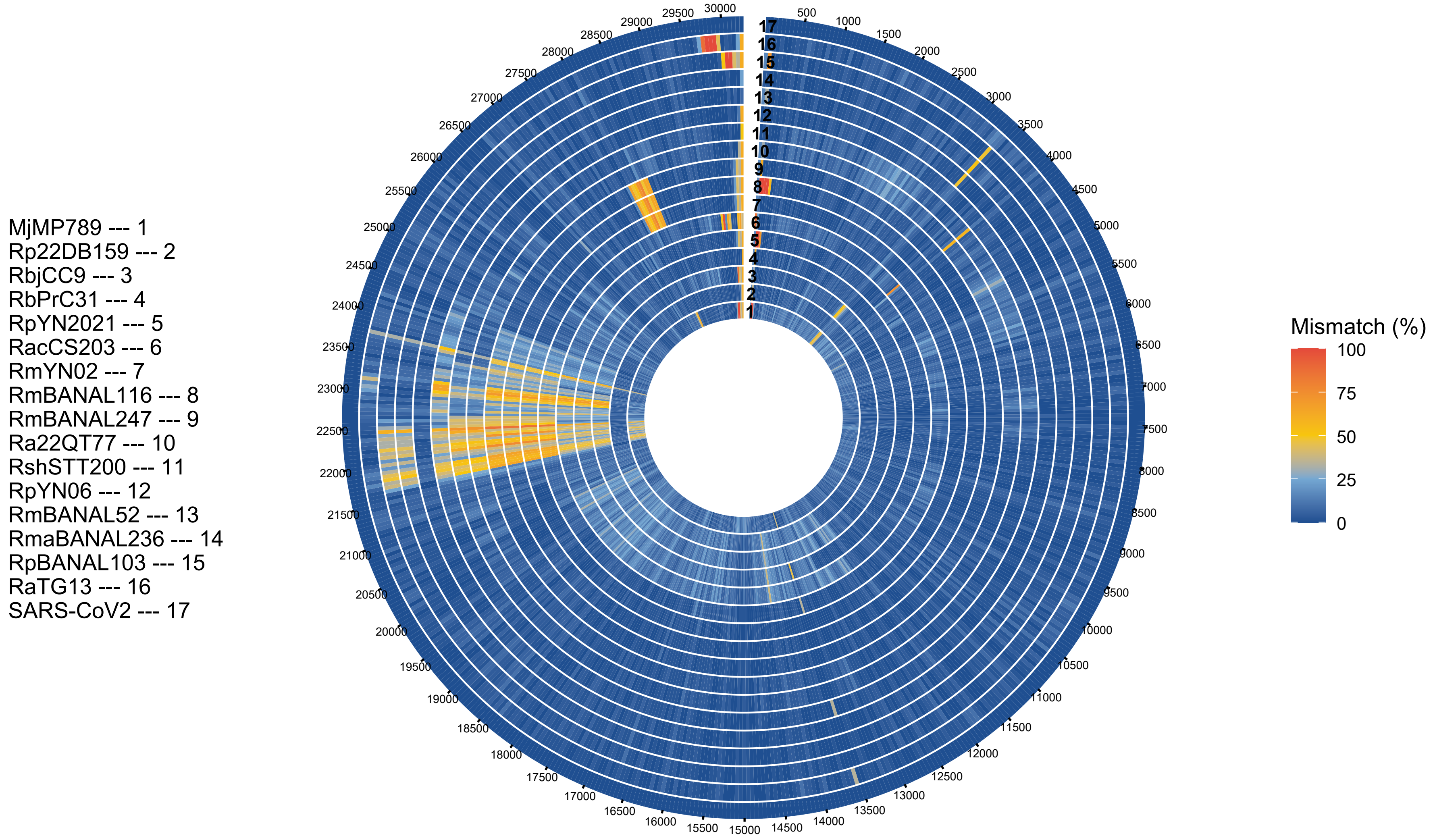
MjMP789 --- 1
Rp22DB159 --- 2
RbjCC9 --- 3
RbPrC31 --- 4
RpYN2021 --- 5
RacCS203 --- 6
RmYN02 --- 7
RmBANAL116 --- 8
RmBANAL247 --- 9
Ra22QT77 --- 10
RshSTT200 --- 11
RpYN06 --- 12
RmBANAL52 --- 13
RmaBANAL236 --- 14
RpBANAL103 --- 15
RaTG13 --- 16
SARS-CoV2 --- 17
Mismatch (%)
100
75
50
25
0

Figure S3. Heatmaps of the genomewide percentages of nucleotide identity for pairs of sequences of the multiple sequence alignment of each dataset. (A) SARS-CoV-1, (B) MERS and (C) SARS-CoV-2. Hierarchical clustering was done with the complete-linkage clustering method. See Data File S1 for numerical values.

(A)

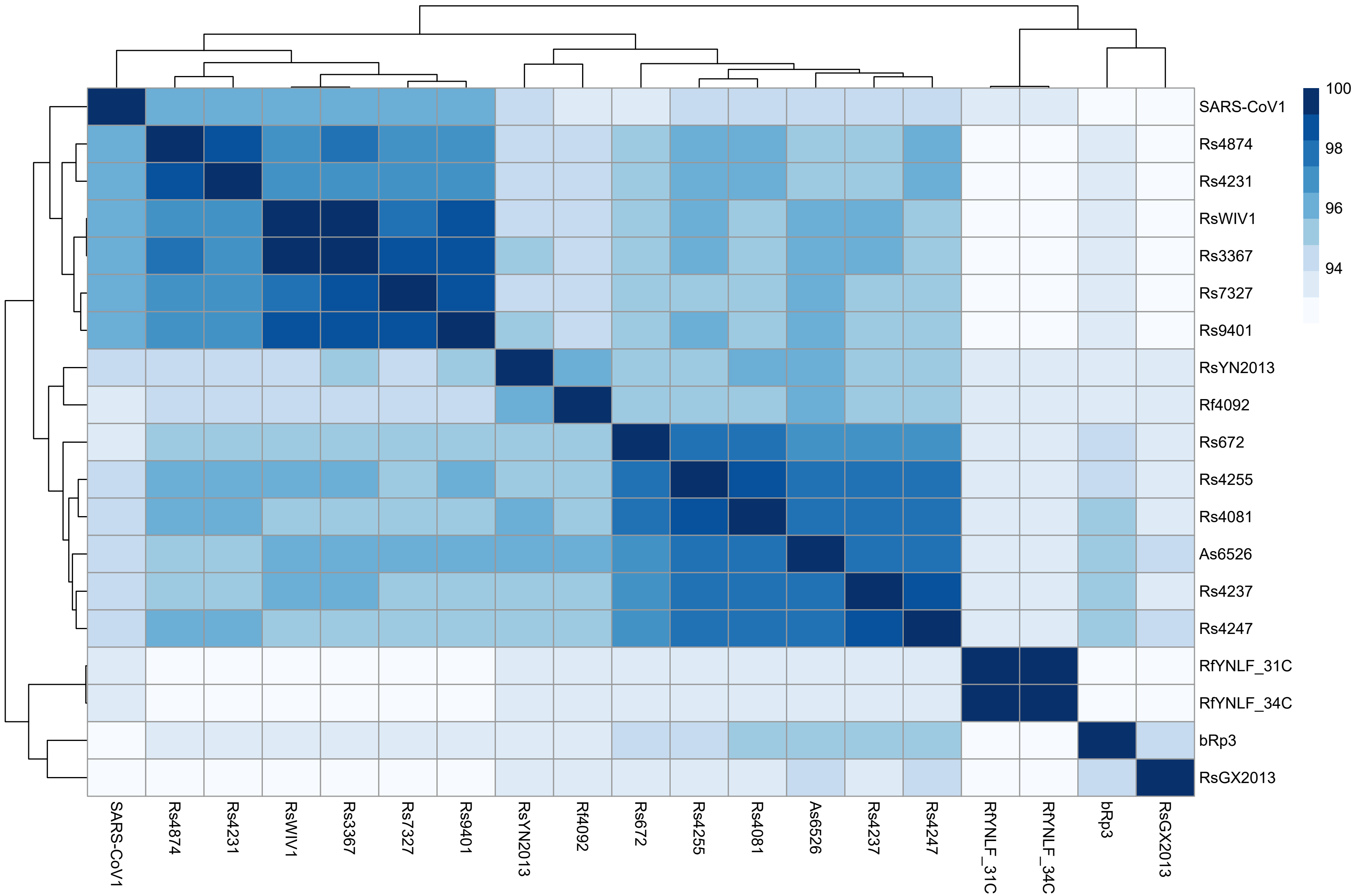

100
98
96
94
SARS-CoV1
Rs4874
Rs4231
RsWIV1
Rs3367
Rs7327
Rs9401
RsYN2013
Rf4092
Rs672
Rs4255
Rs4081
As6526
Rs4237
Rs4247
RfYNLF_31C
RfYNLF_34C
bRp3
RsGX2013

(B)

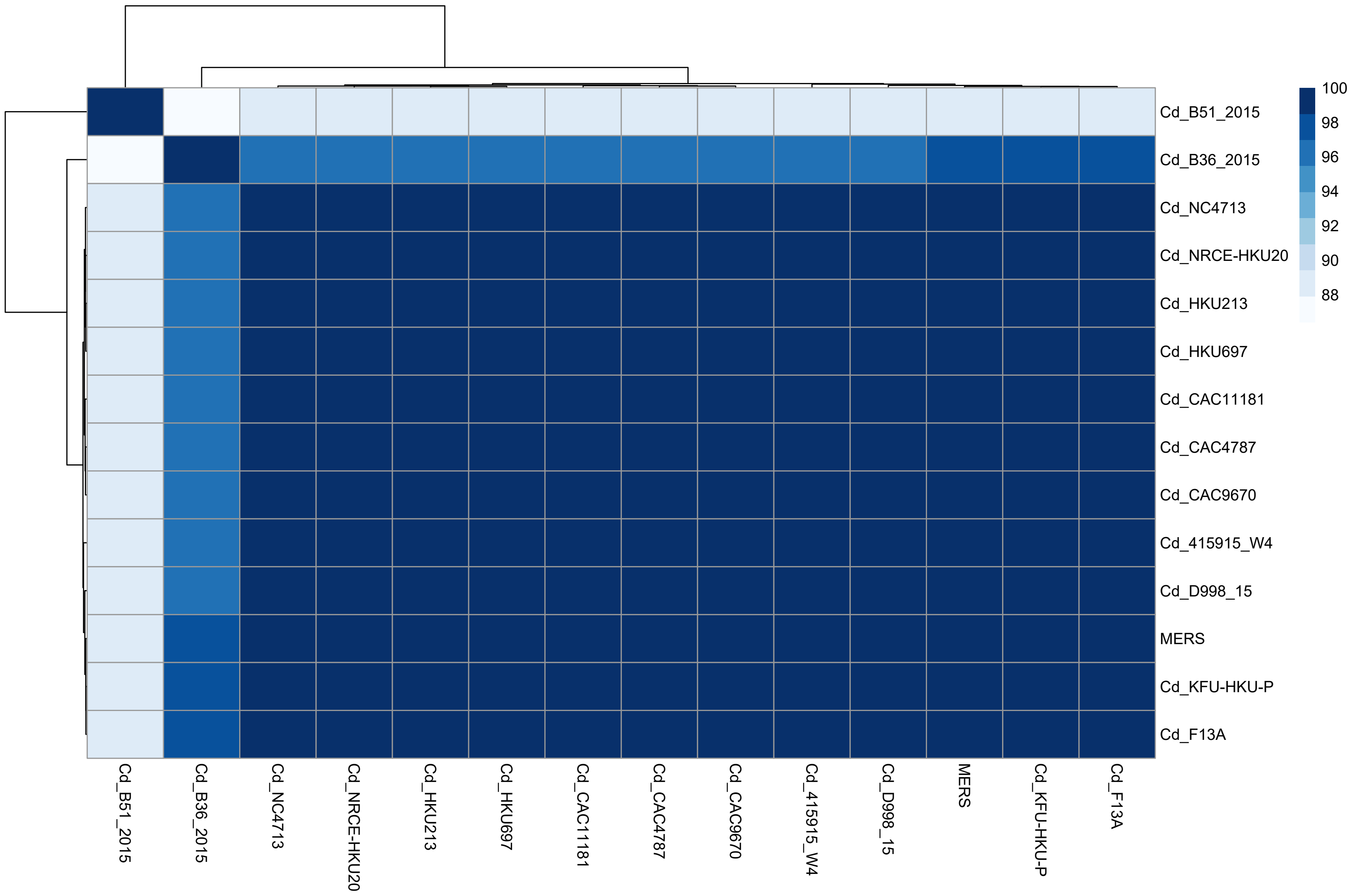

Cd_B51_2015
Cd_B36_2015
Cd_NC4713
Cd_NRCE-HKU20
Cd_HKU213
Cd_HKU697
Cd_CAC11181
Cd_CAC4787
Cd_CAC9670
Cd_415915_W4
Cd_D998_15
MERS
Cd_KFU-HKU-P
Cd_F13A
100
98
96
94
92
90
88

(C)

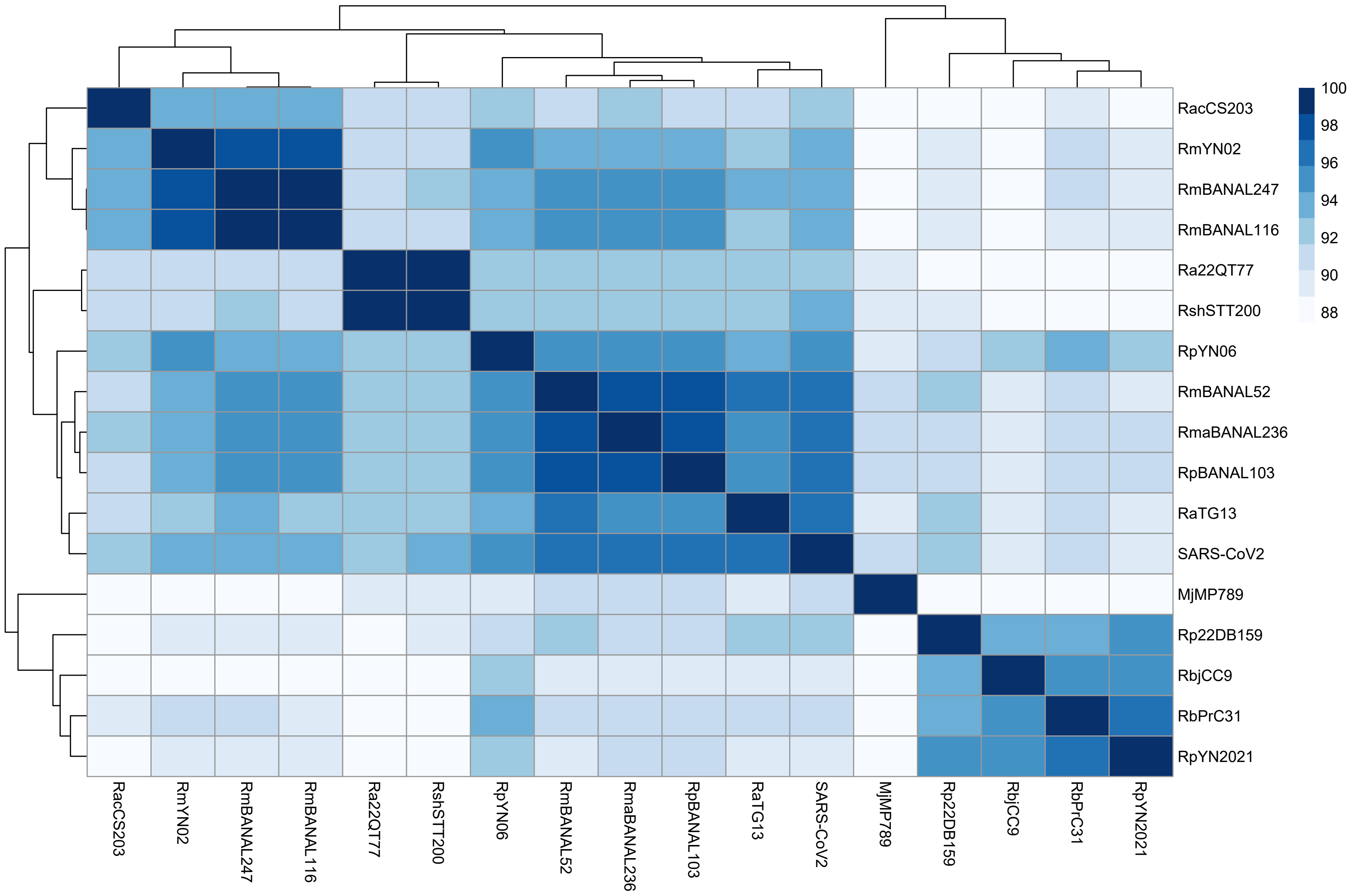
RacCS203
RmYN02
RmBANAL247
RmBANAL116
Ra22QT77
RshSTT200
RpYN06
RmBANAL52
RmaBANAL236
RpBANAL103
RaTG13
SARS-CoV2
MjMP789
Rp22DB159
RbjCC9
RbPrC31
RpYN2021
100
98
96
94
92
90
88

Figure S4. Positions of the inferred breakpoints at each segmentation step from 1 to 30 segments. (A) SARS-CoV-1, (B) MERS and (C) SARS-CoV-2. The $x$-axis indicates the nucleotide positions in the multiple sequence alignment. The $y$-axis indicates the number of segmentation steps. The segmentation chosen for further analysis is colored in red.

(A)

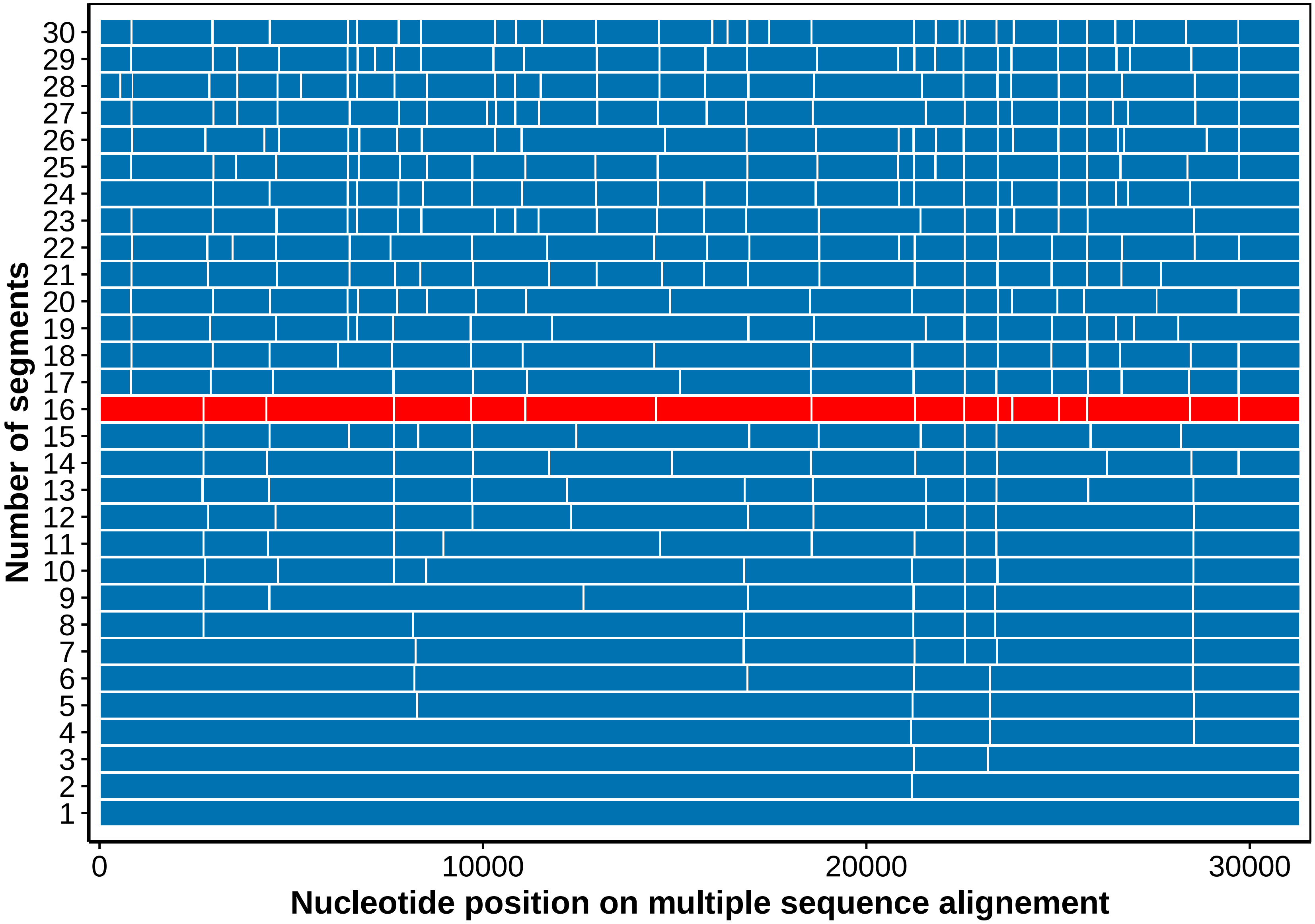

Number of segments
30
29
28
27
26
25
24
23
22
21
20
19
18
17
16
15
14
13
12
11
10
9
8
7
6
5
4
3
2
1
0
10000
20000
30000
Nucleotide position on multiple sequence alignement

(B)

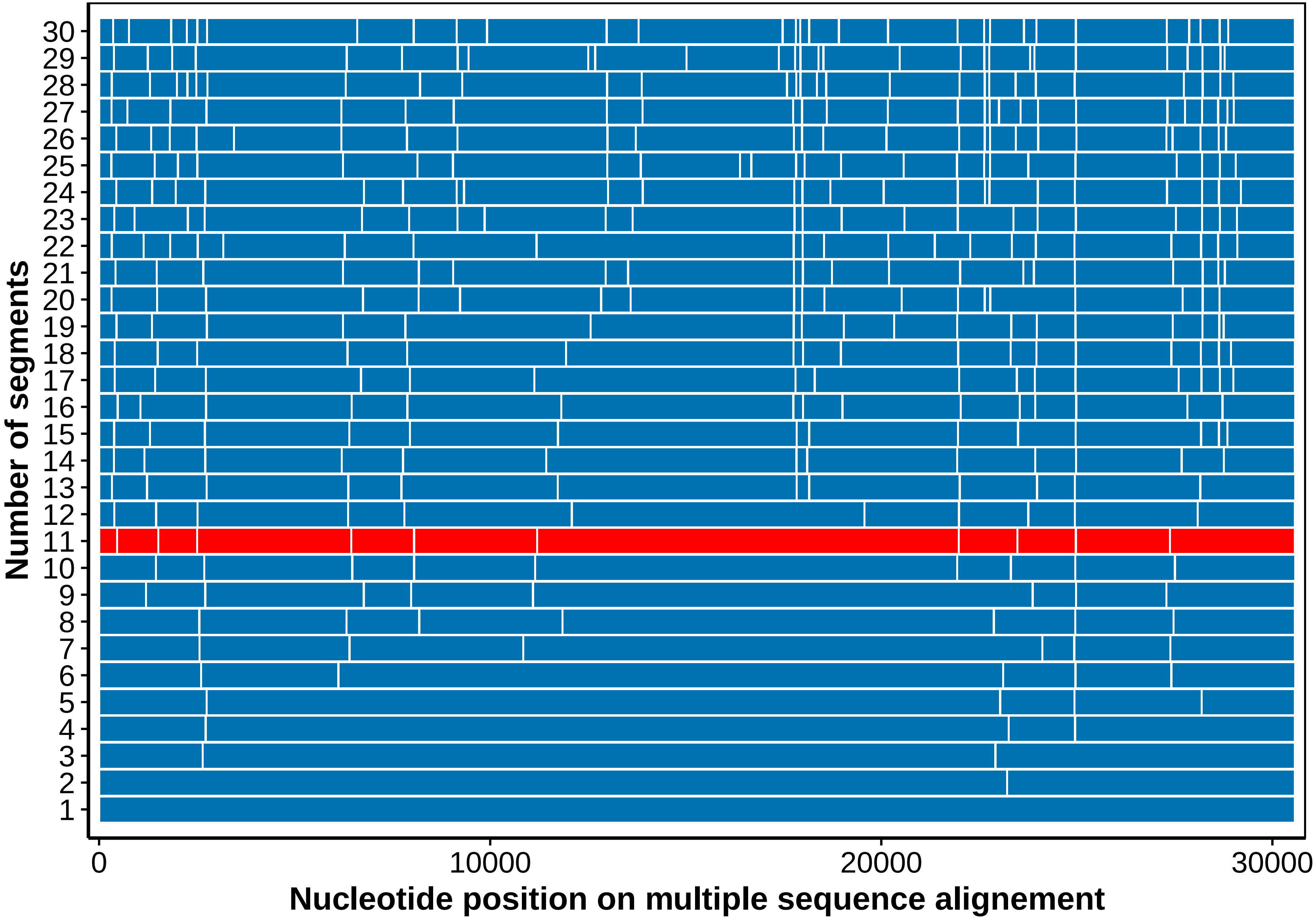

Number of segments
30
29
28
27
26
25
24
23
22
21
20
19
18
17
16
15
14
13
12
11
10
9
8
7
6
5
4
3
2
1
0
10000
20000
30000
Nucleotide position on multiple sequence alignement

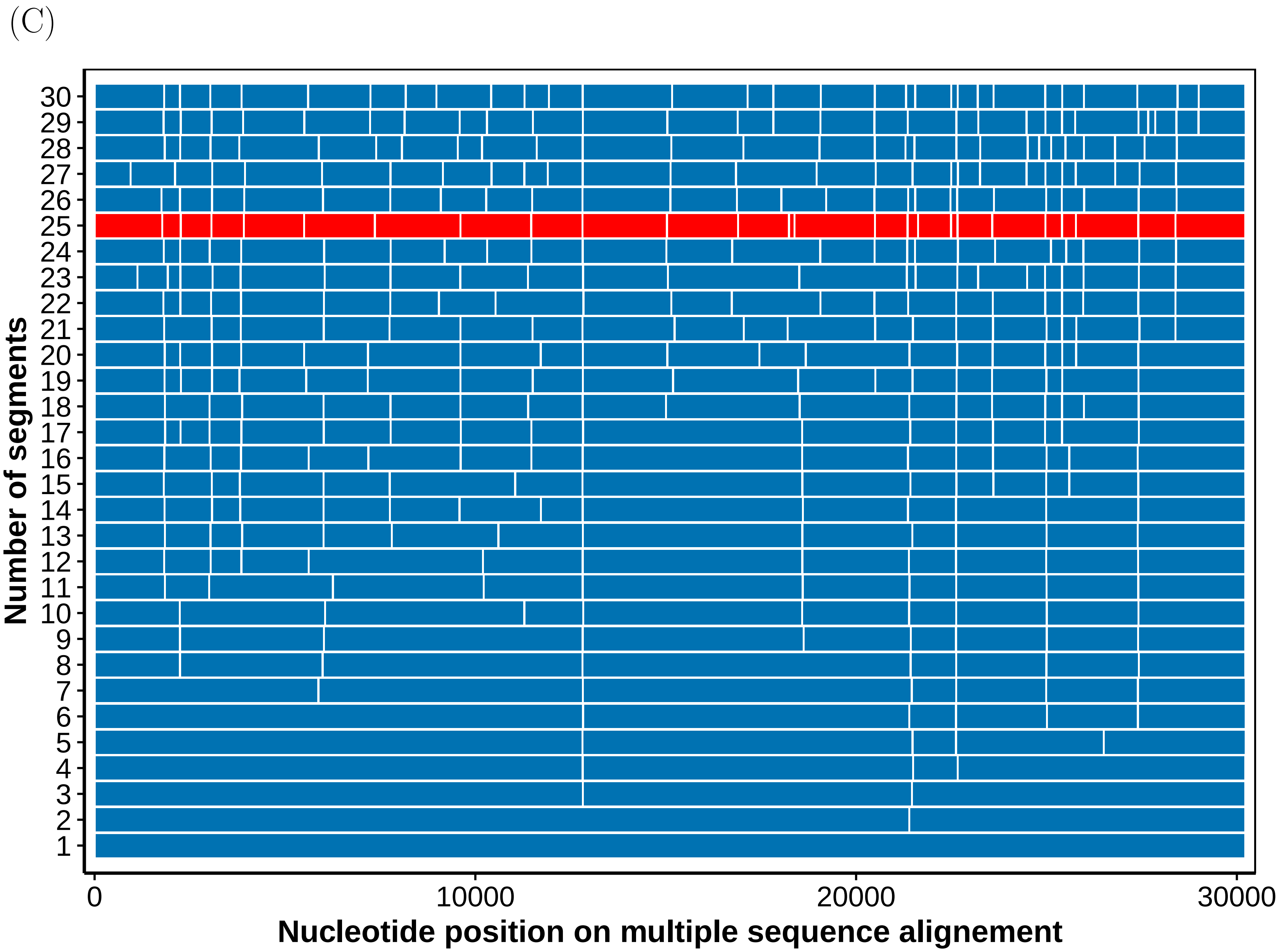

(C)
Number of segments
30
29
28
27
26
25
24
23
22
21
20
19
18
17
16
15
14
13
12
11
10
9
8
7
6
5
4
3
2
1
0
10000
20000
30000
Nucleotide position on multiple sequence alignement

Figure S5. Subsequent improvement of log likelihood as the number of segments increases. (A) SARS-CoV-1, (B) MERS and (C) SARS-CoV-2. The red arrow and disk point to the segmentation kept for further analysis.

(A)

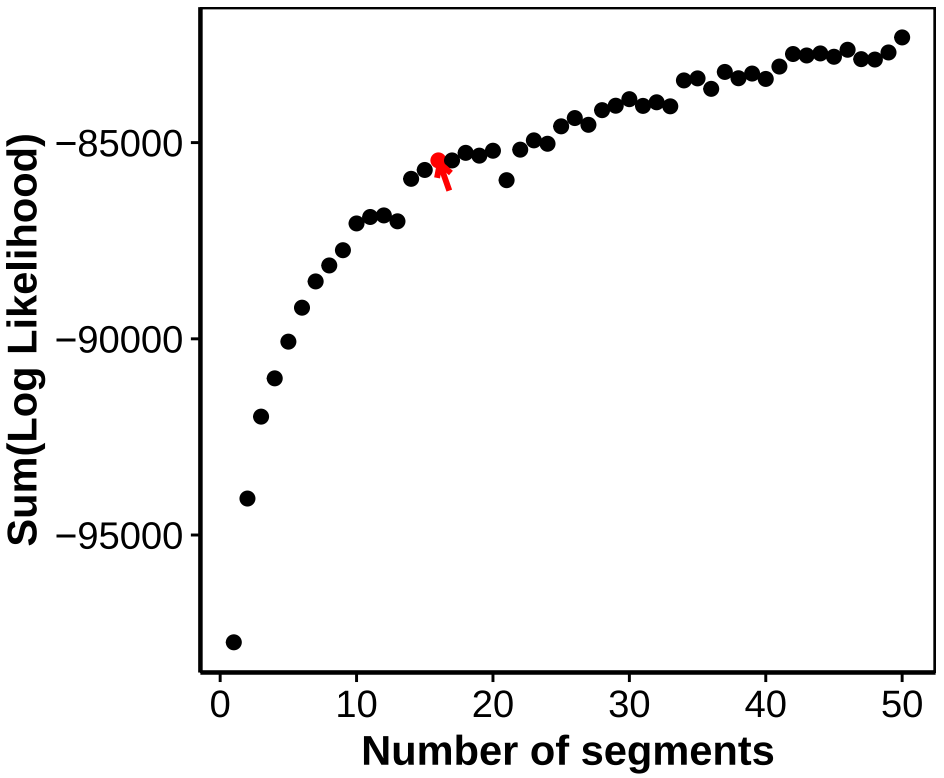

Sum(Log Likelihood)
−85000
−90000
−95000
0
10
20
30
40
50
Number of segments

(B)

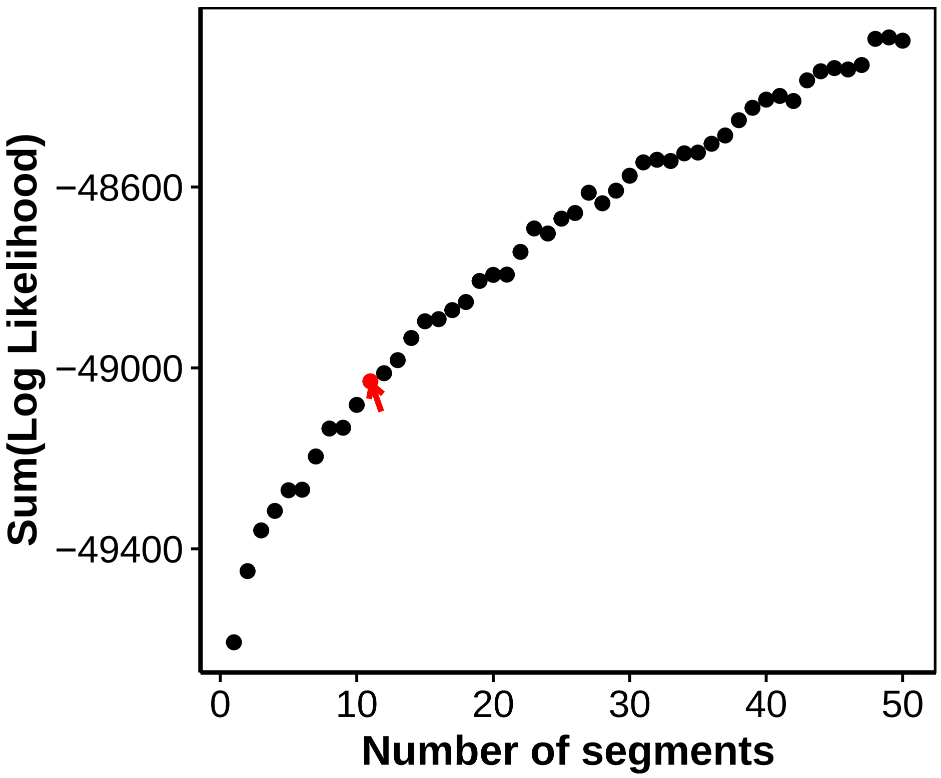

Sum(Log Likelihood)
−48600
−49000
−49400
0
10
20
30
40
50
Number of segments

(C)

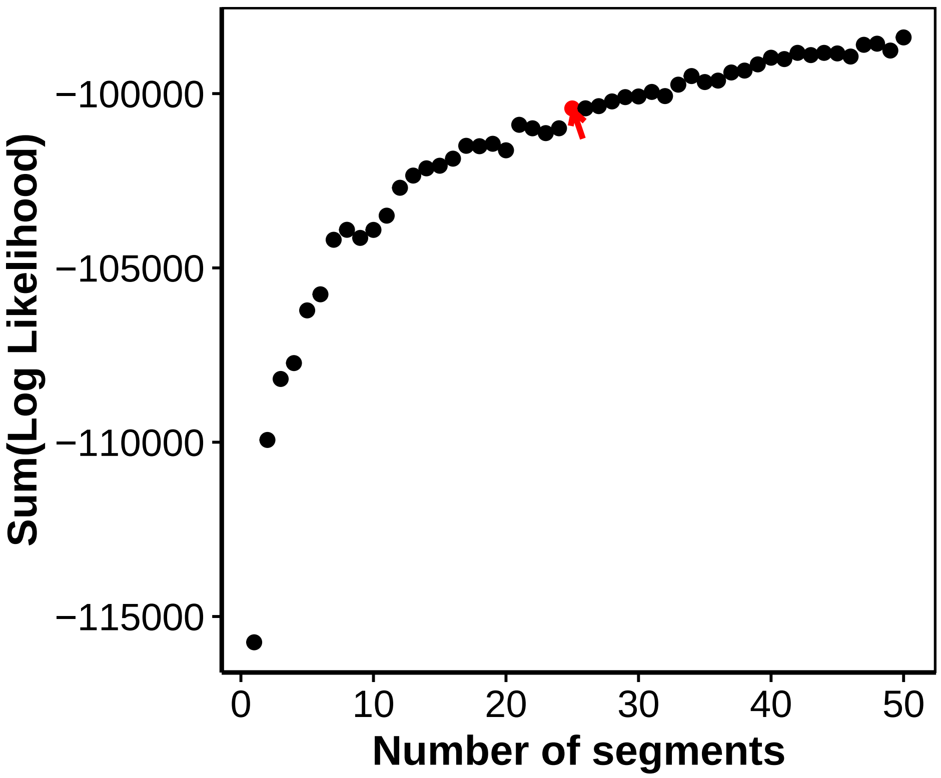

−100000
−105000
−110000
−115000
Sum(Log Likelihood)
0
10
20
30
40
50
Number of segments

Figure S6. Akaike Information Criteria (AIC) computed as a function of the number of segments. (A) SARS-CoV-1, (B) MERS and (C) SARS-CoV-2. The red arrow and disk point to the segmentation kept for further analysis.

(A)

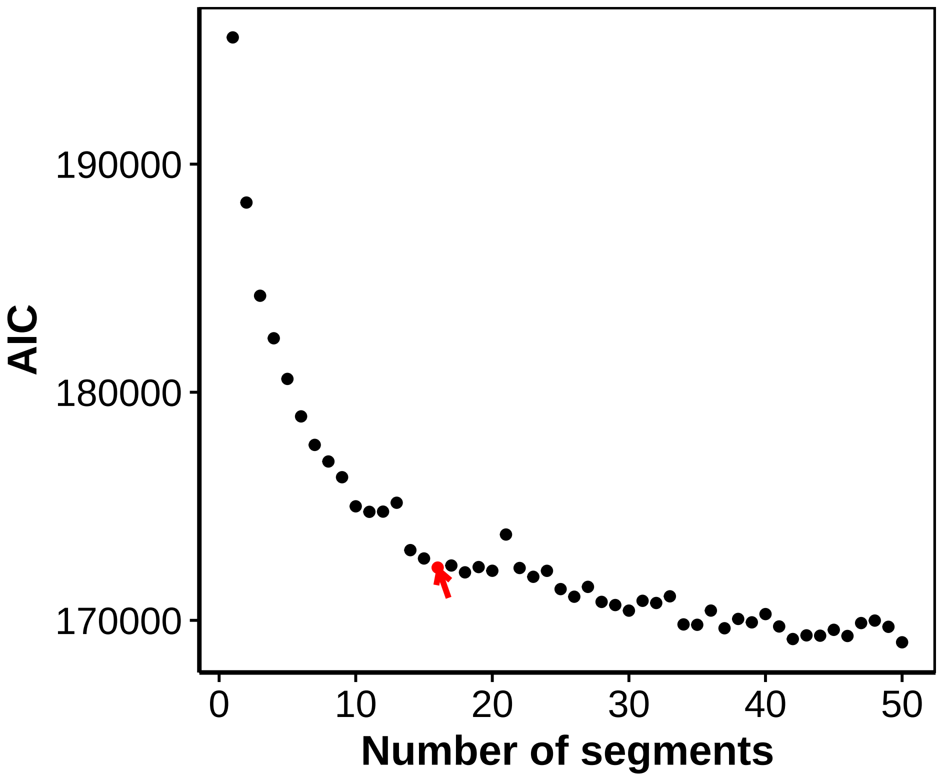

(B)

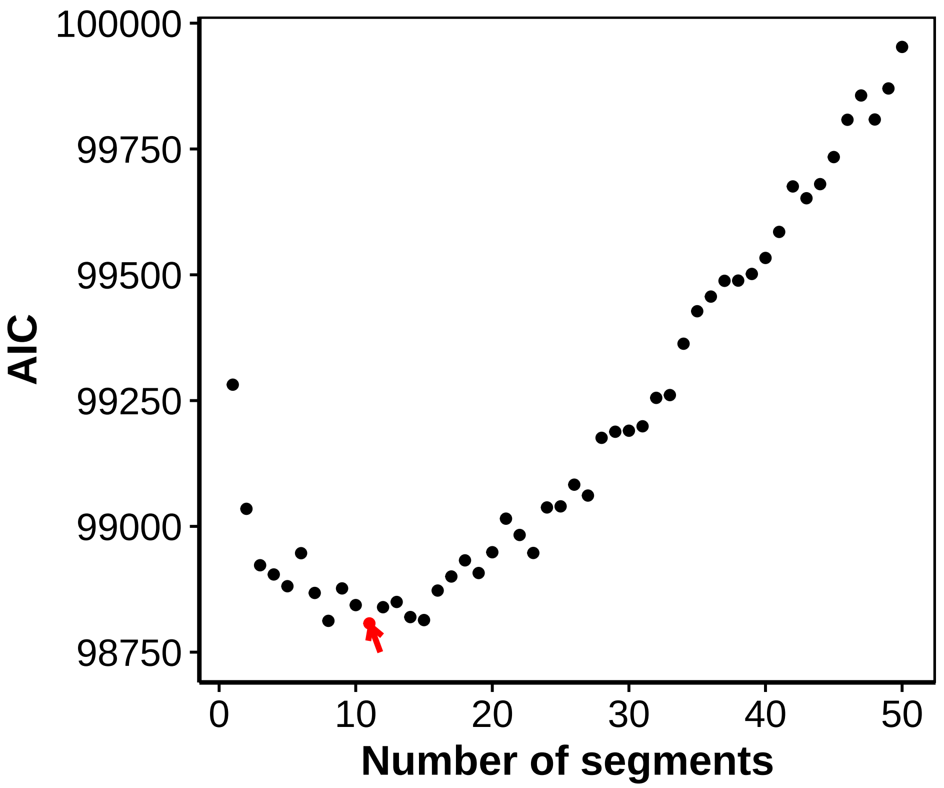
AIC
100000
99750
99500
99250
99000
98750
0
10
20
30
40
50
Number of segments

(C)

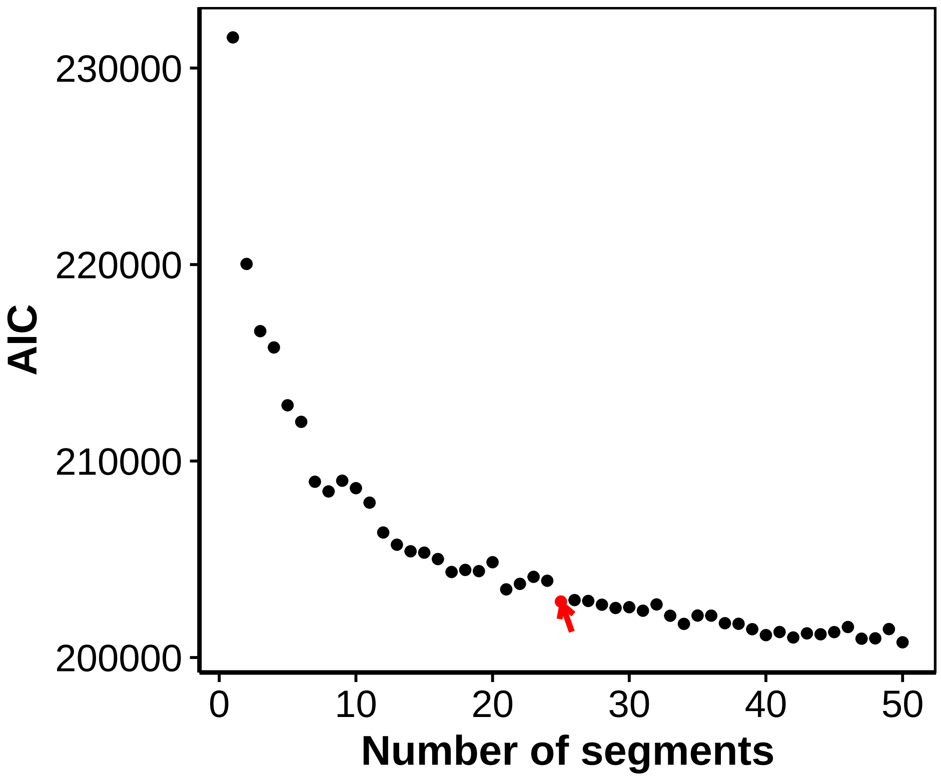
230000
220000
210000
200000
AIC
0
10
20
30
40
50
Number of segments

Figure S7. Bayesian Information Criteria (BIC) computed as a function of the number of segments. (A) SARS-CoV-1, (B) MERS and (C) SARS-CoV-2. The red arrow and disk point to the segmentation kept for further analysis.

(A)

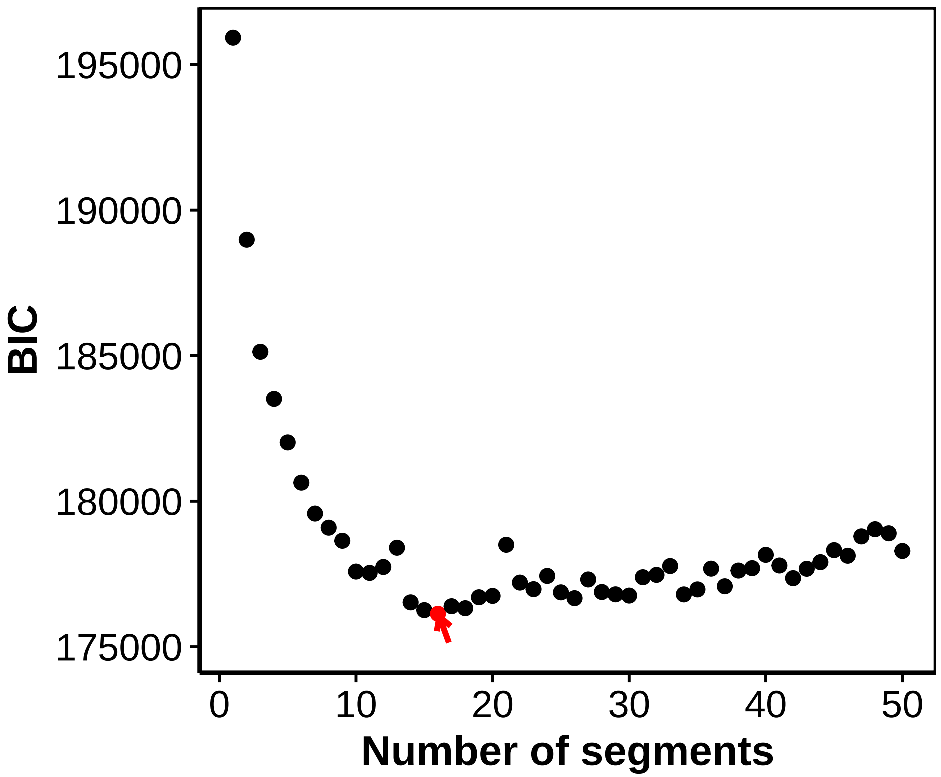

BIC
195000
190000
185000
180000
175000
0
10
20
30
40
50
Number of segments

(B)

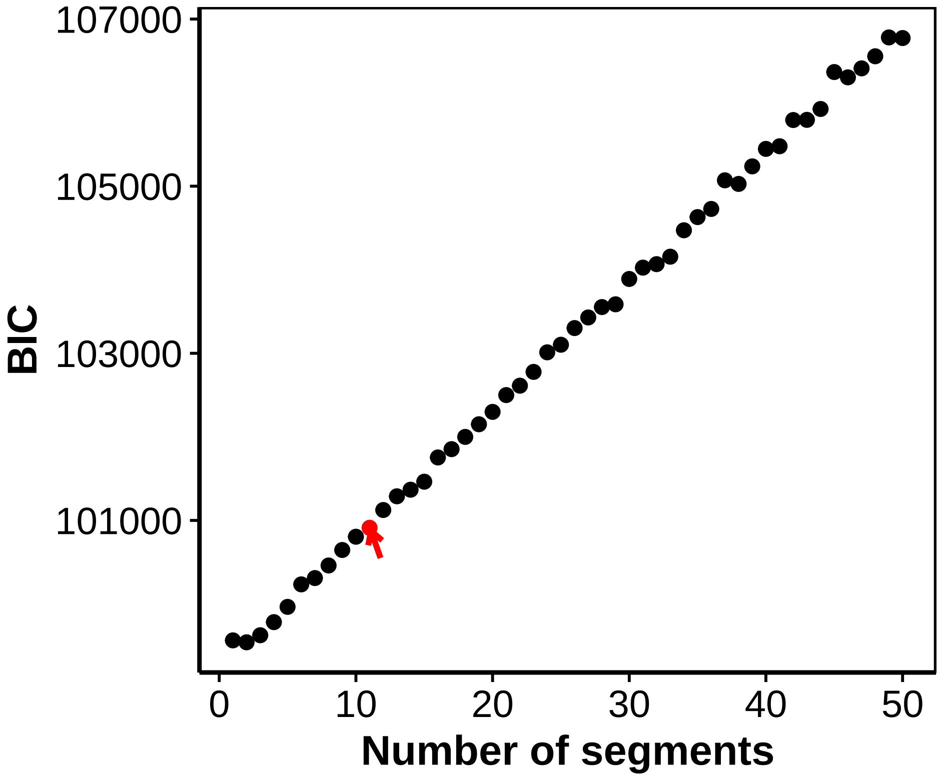

107000
105000
103000
101000
BIC
0
10
20
30
40
50
Number of segments

(C)

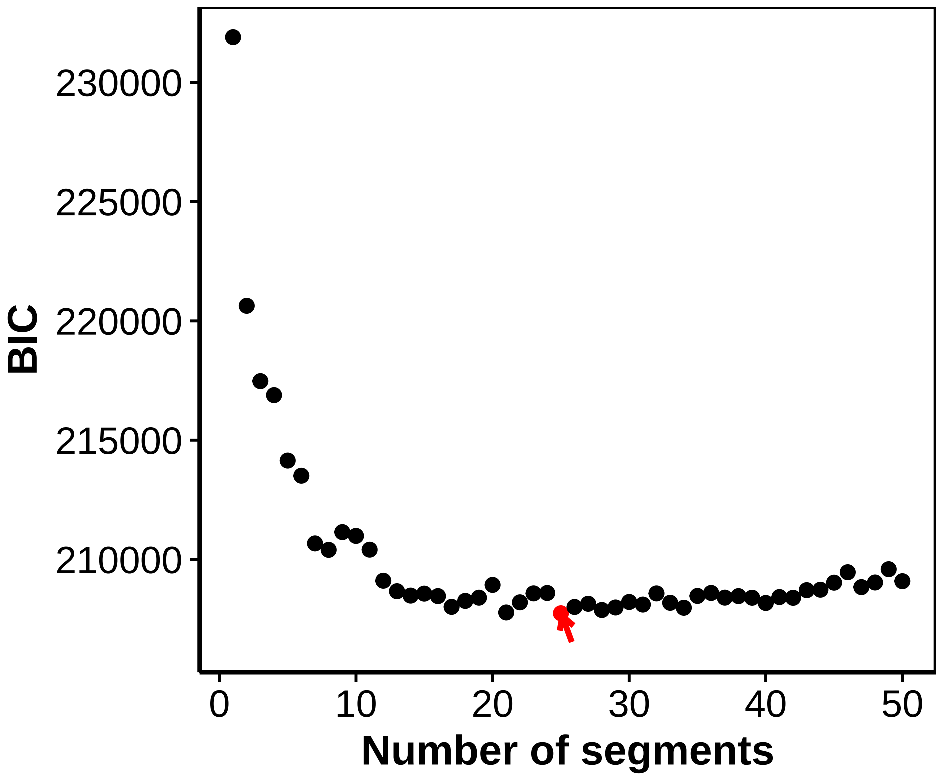

230000
225000
220000
215000
210000
BIC
0
10
20
30
40
50
Number of segments

Figure S8. Homoplasy versus number of segments. (A) SARS-CoV-1, (B) MERS and (C) SARS-CoV-2. The black dot highlights the segmentation selected: 16 segments for SARS-CoV-1, 11 segments for MERS and 25 segments for SARS-CoV-2. The red point and dashed line represent the homoplasy value obtained with the GARD algorithm on the CNCA alignment.

(A)

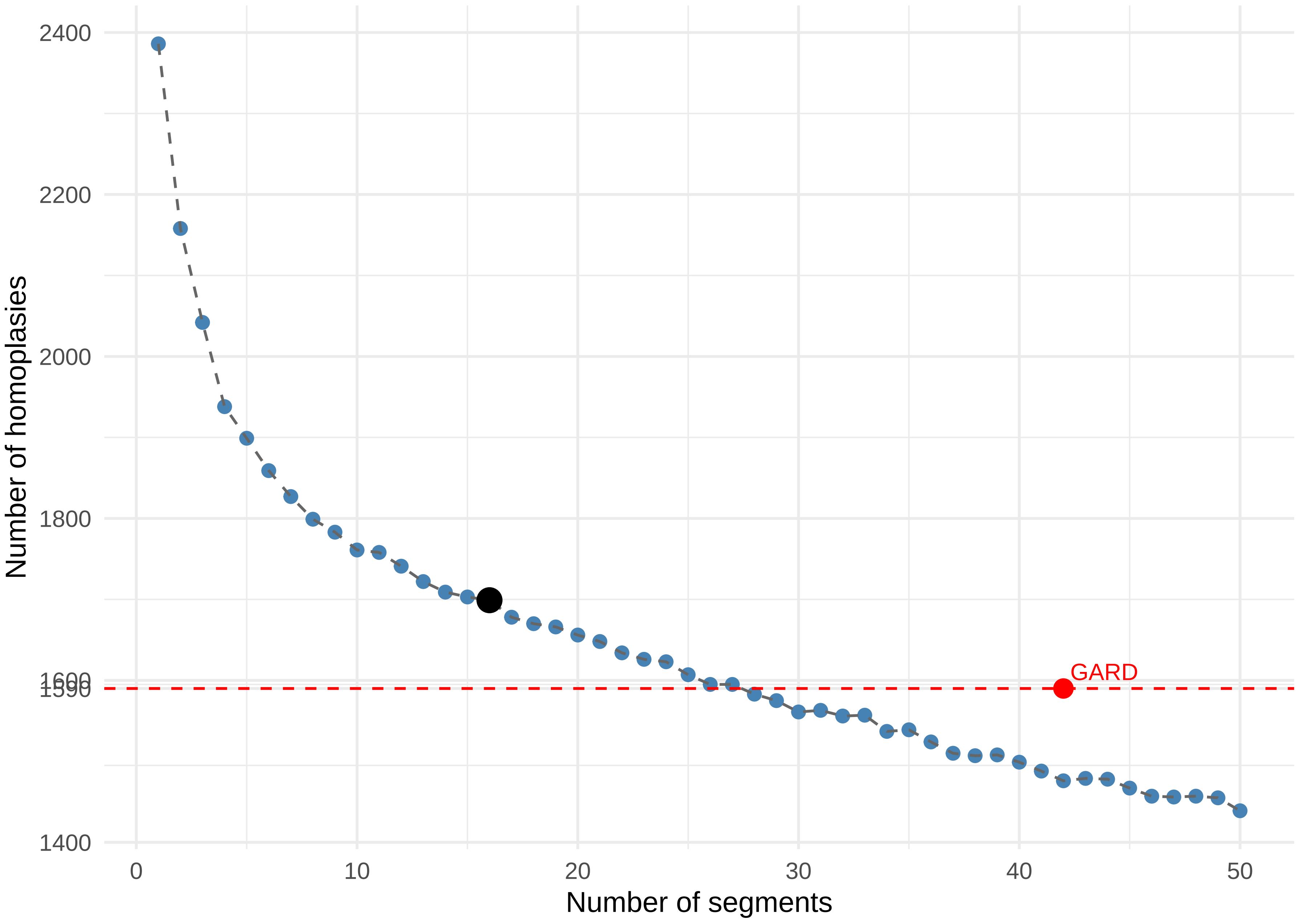

2400
2200
2000
1800
1600
1590
1400
0
10
20
30
40
50
GARD
Number of homoplasies
Number of segments

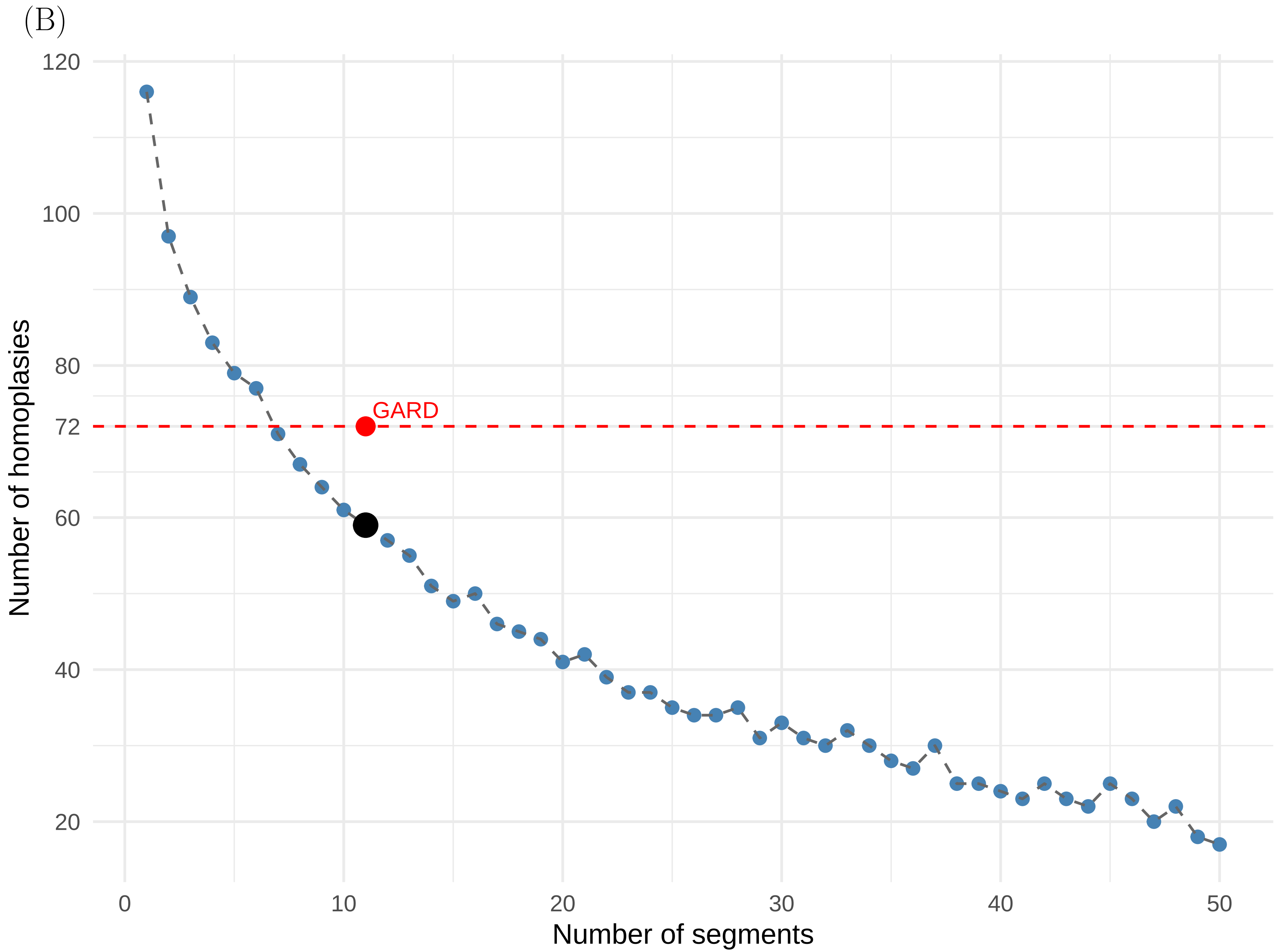

(B)
GARD
Number of homoplasies
Number of segments
120
100
80
72
60
40
20
0
10
20
30
40
50

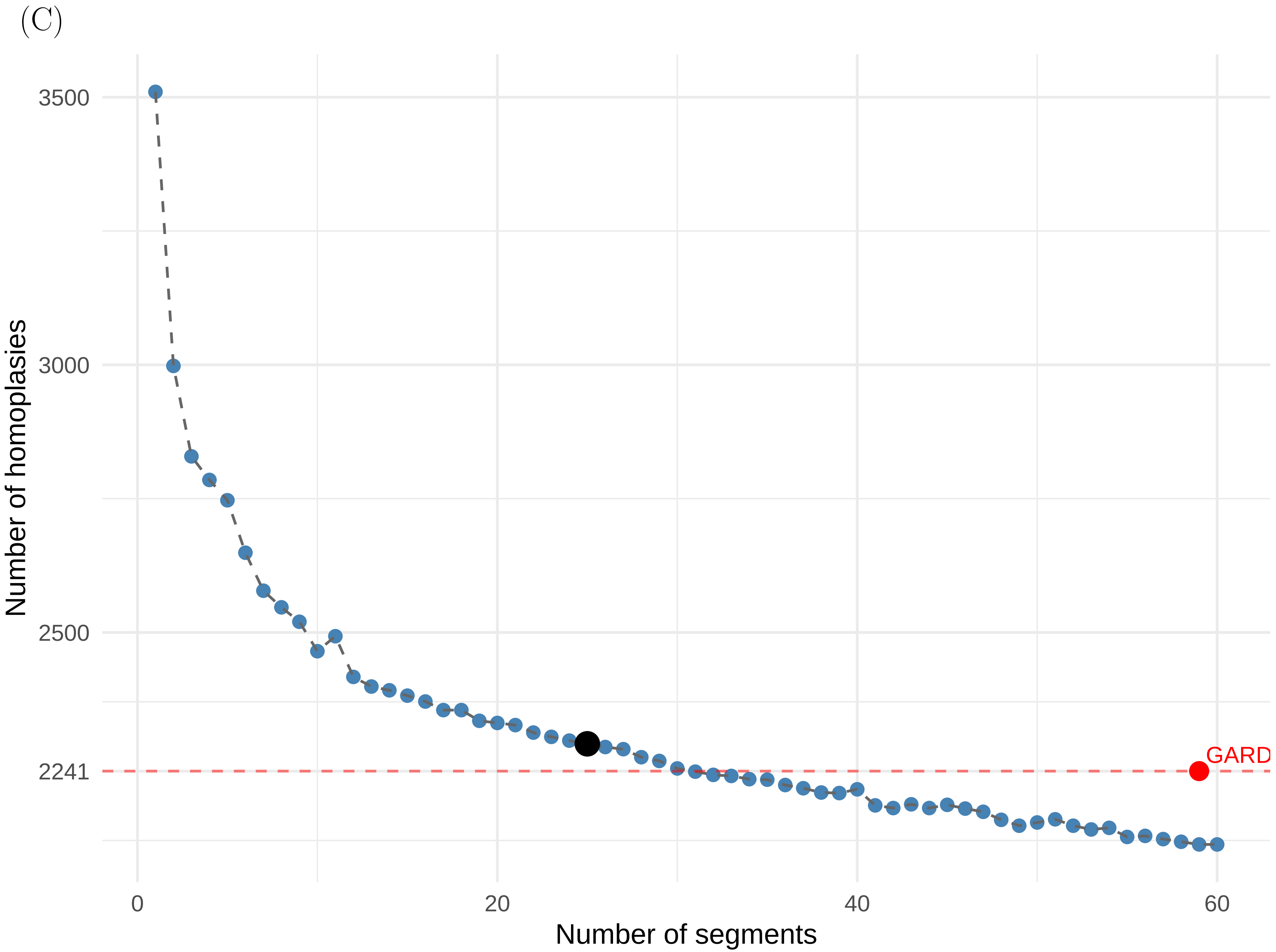
(C)
3500
3000
2500
2241
0
20
40
60
GARD
Number of homoplasies
Number of segments

Figure S9. Cumulative homoplasy values along the CNCA alignment. (A) SARS-CoV-1, (B) MERS and (C) SARS-CoV-2. Segmentations and methods are compared, for example, GARD 59 means segmentation into 59 segments with GARD. The $x$-axis shows nucleotide positions in the multiple sequence alignment. The $y$-axis shows the cumulative homoplasy value, computed as the sum of each segment homoplasy number $H$. Each dot in the plot corresponds to a new segment boundary, where the $H$ value of that segment is added to the cumulative total.

(A)

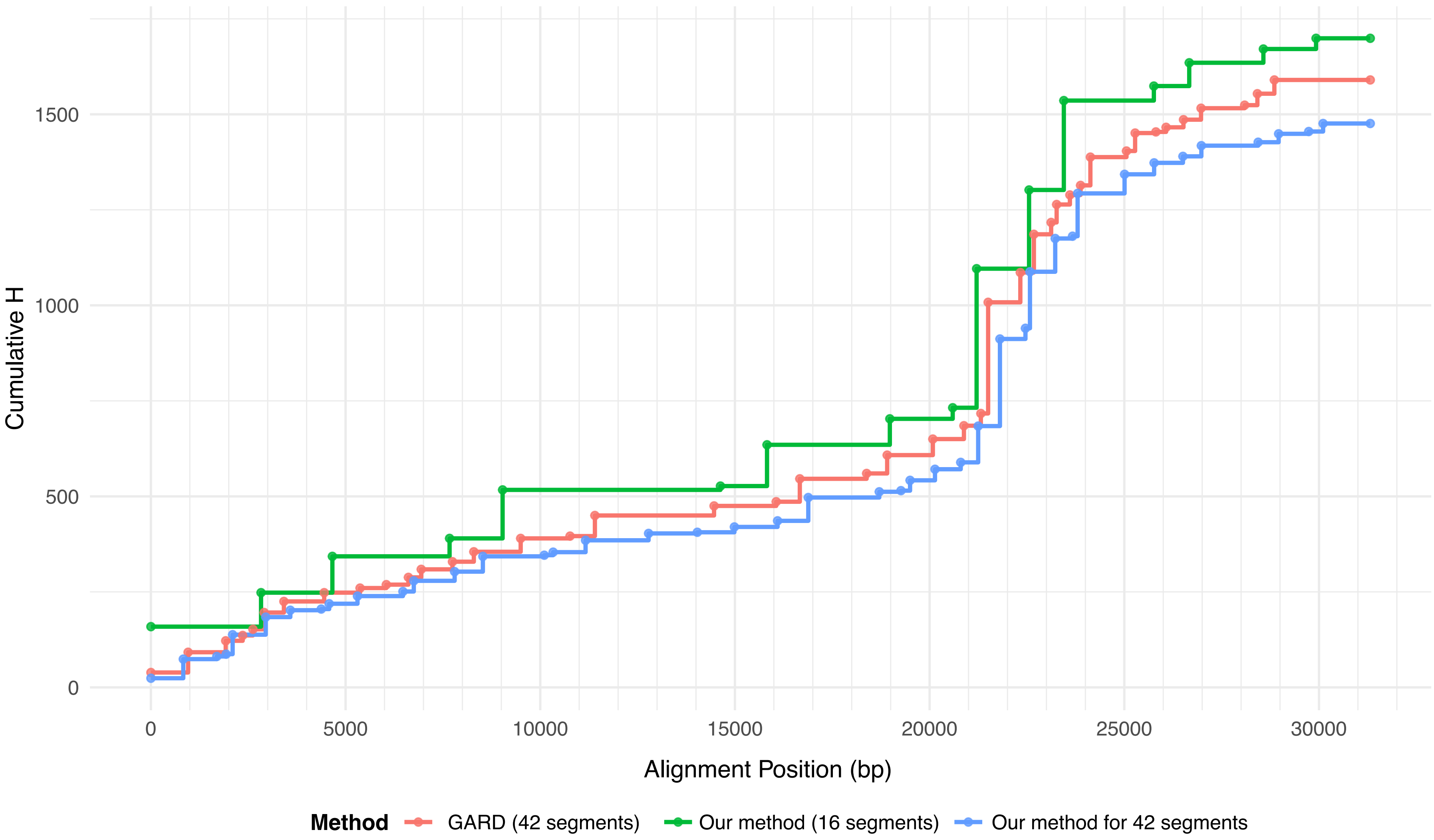

Cumulative H
1500
1000
500
0
0
5000
10000
15000
20000
25000
30000
Alignment Position (bp)
Method
GARD (42 segments)
Our method (16 segments)
Our method for 42 segments

(B)

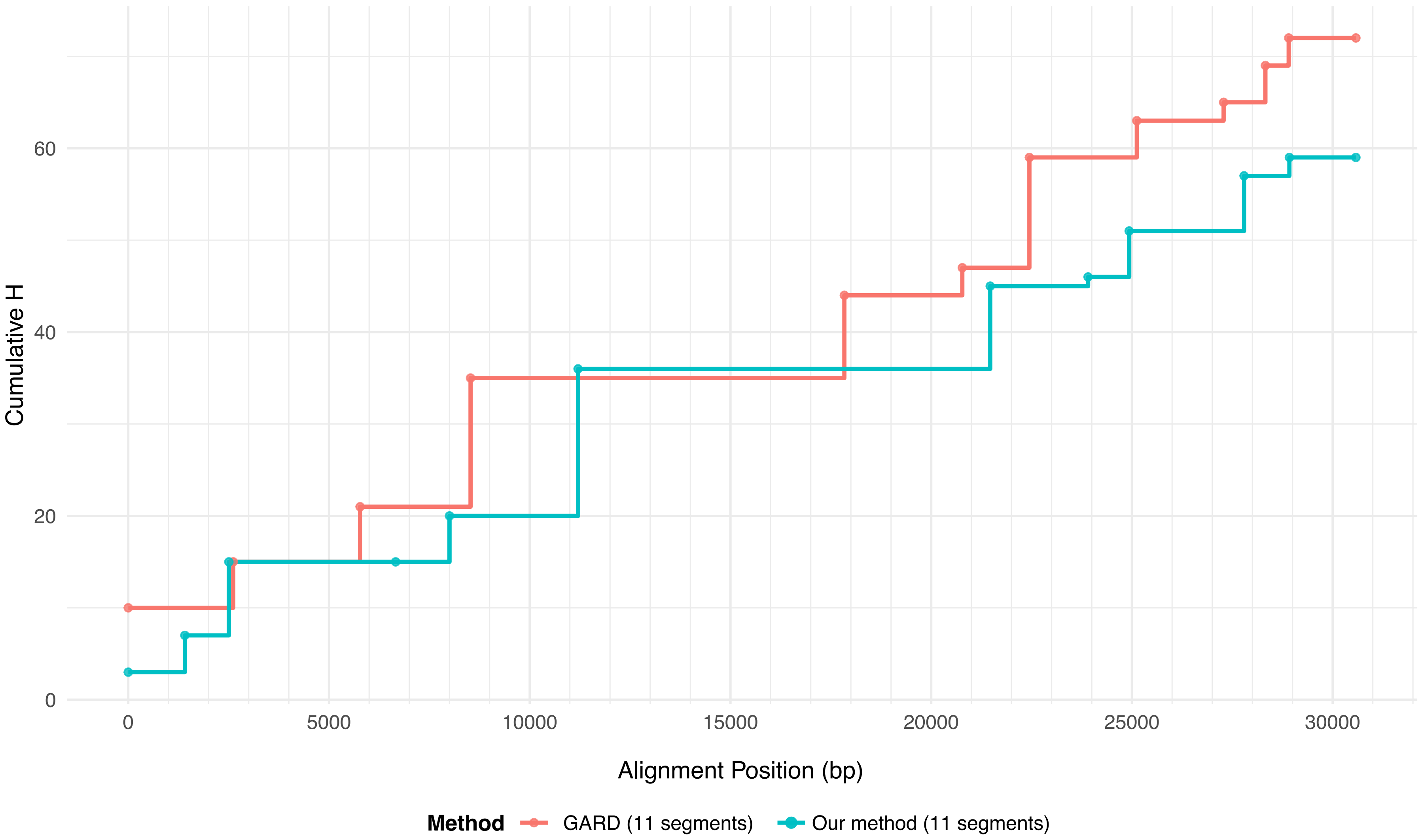

Cumulative H
0
20
40
60
0
5000
10000
15000
20000
25000
30000
Alignment Position (bp)
Method
GARD (11 segments)
Our method (11 segments)

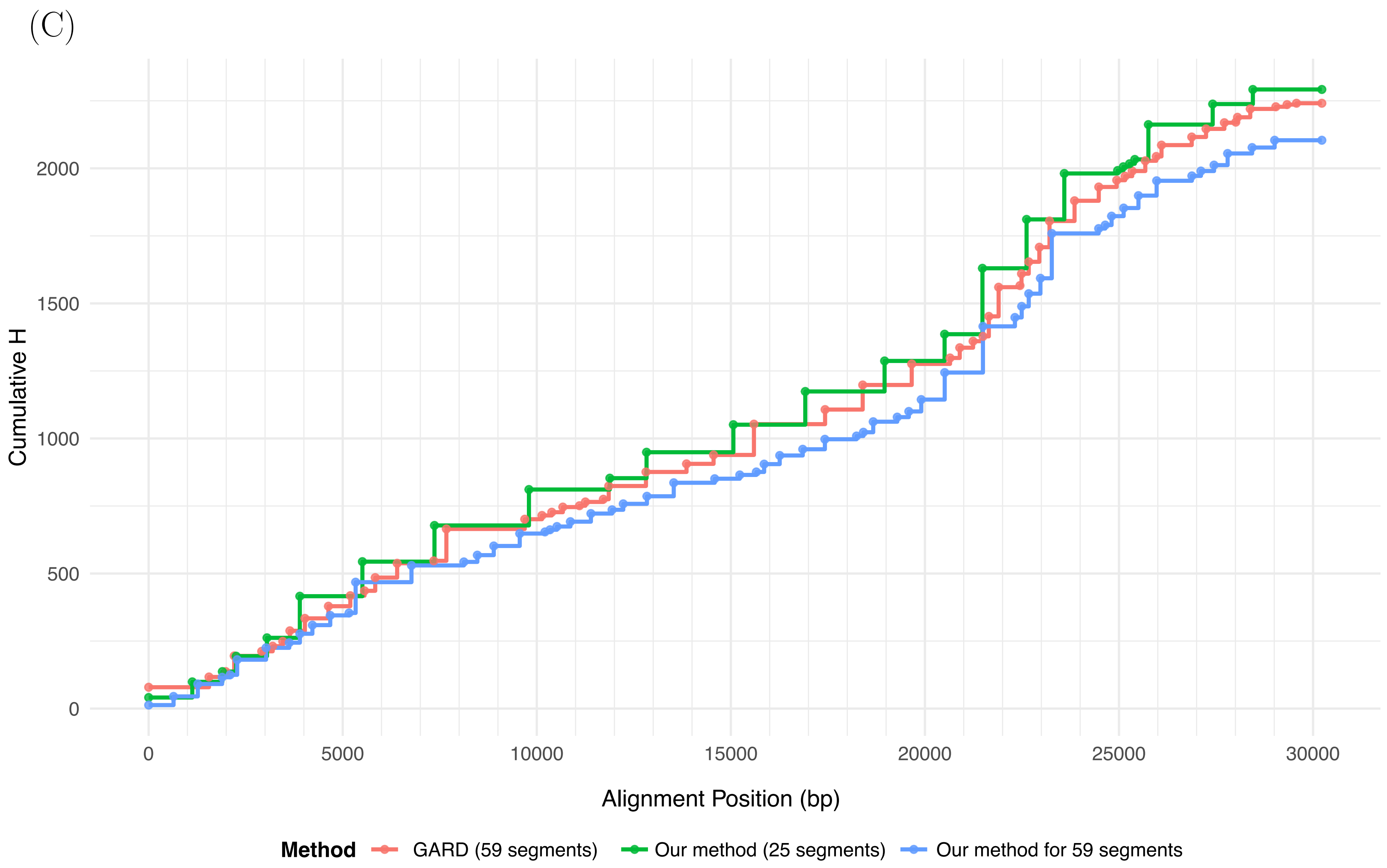
(C)
Cumulative H
0
500
1000
1500
2000
0
5000
10000
15000
20000
25000
30000
Alignment Position (bp)
Method
GARD (59 segments)
Our method (25 segments)
Our method for 59 segments

Figure S10. Phylograms of all genomic segments for the three reference datasets. (A) SARS-CoV-1, (B) MERS, and (C) SARS-CoV-2. Each tree corresponds to one segment, identified by its nucleotide range in the title. Trees are shown as phylograms; branch lengths are proportional to the estimated number of substitutions per site under the GTR+FO+G4m (*ie* RAxML-NG was run with model GTR+G, which was optimized as GTR+FO+G4m.) substitution model. A shared scale bar is displayed for each page to facilitate comparison among the four trees shown together.

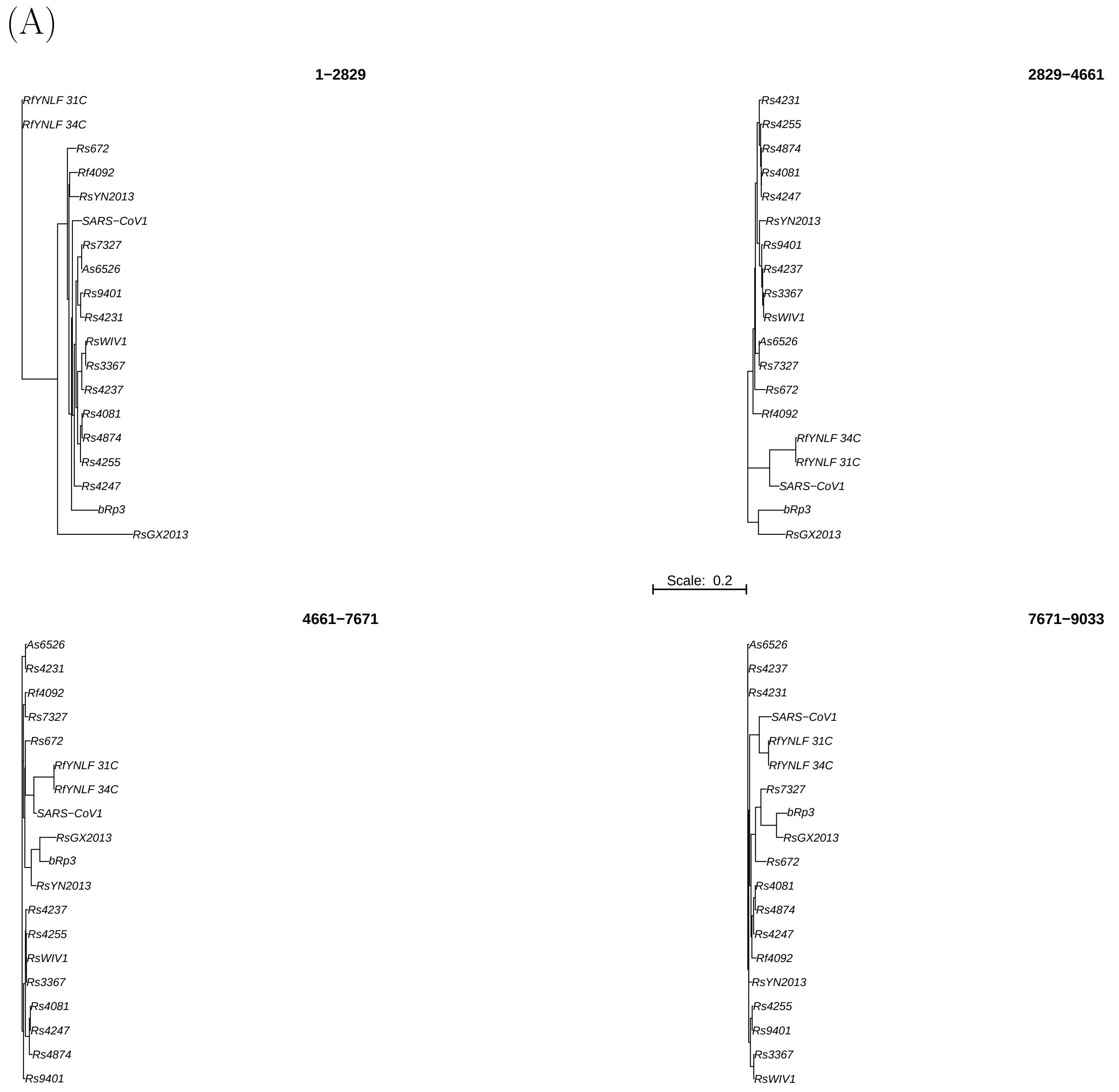
(A)
1−2829
RfYNLF 31C
RfYNLF 34C
Rs672
Rf4092
RsYN2013
SARS−CoV1
Rs7327
As6526
Rs9401
Rs4231
RsWIV1
Rs3367
Rs4237
Rs4081
Rs4874
Rs4255
Rs4247
bRp3
RsGX2013
2829−4661
Rs4231
Rs4255
Rs4874
Rs4081
Rs4247
RsYN2013
Rs9401
Rs4237
Rs3367
RsWIV1
As6526
Rs7327
Rs672
Rf4092
RfYNLF 34C
RfYNLF 31C
SARS−CoV1
bRp3
RsGX2013
Scale: 0.2
4661−7671
As6526
Rs4231
Rf4092
Rs7327
Rs672
RfYNLF 31C
RfYNLF 34C
SARS−CoV1
RsGX2013
bRp3
RsYN2013
Rs4237
Rs4255
RsWIV1
Rs3367
Rs4081
Rs4247
Rs4874
Rs9401
7671−9033
As6526
Rs4237
Rs4231
SARS−CoV1
RfYNLF 31C
RfYNLF 34C
Rs7327
bRp3
RsGX2013
Rs672
Rs4081
Rs4874
Rs4247
Rf4092
RsYN2013
Rs4255
Rs9401
Rs3367
RsWIV1

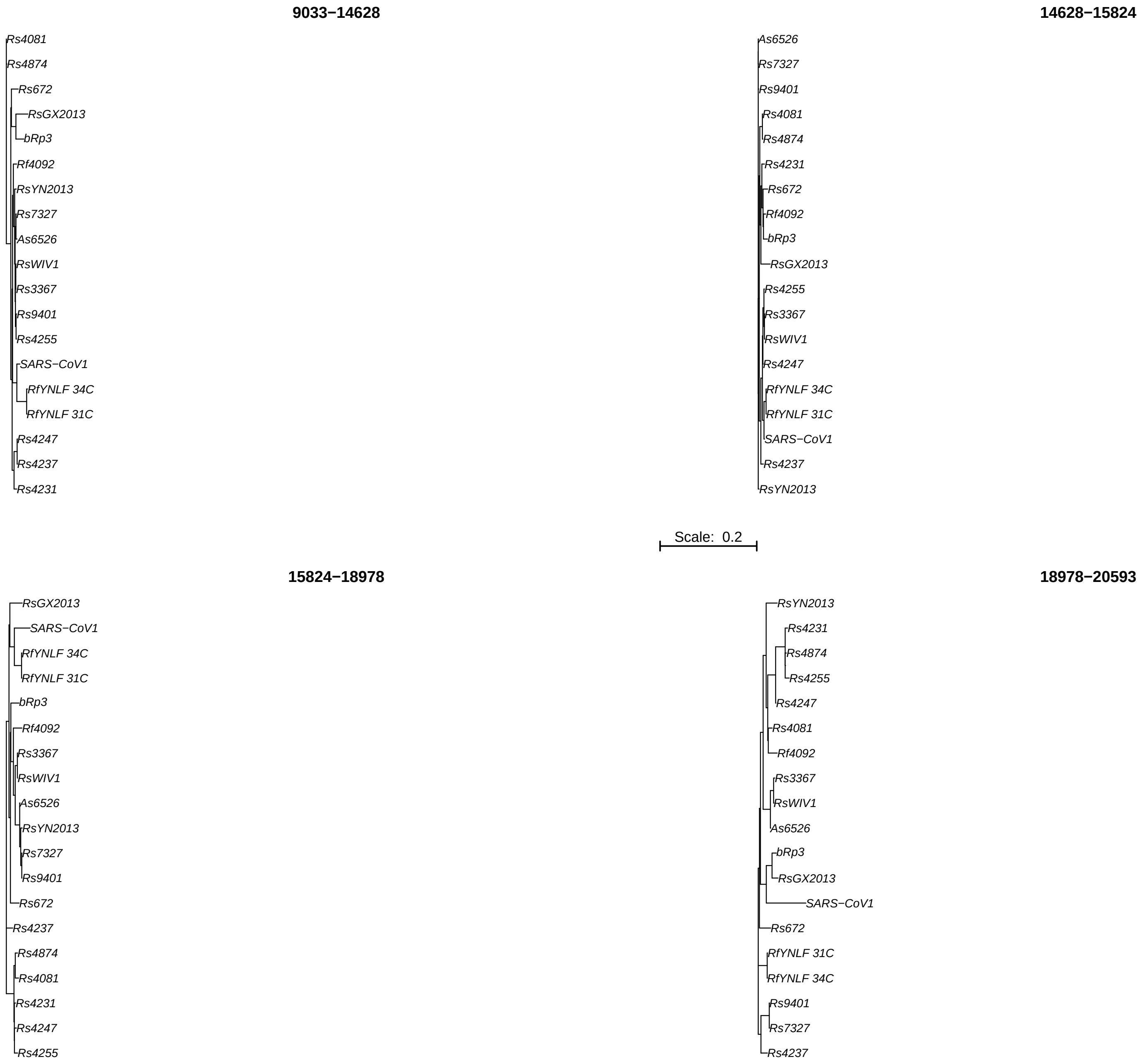
9033−14628
Rs4081
Rs4874
Rs672
RsGX2013
bRp3
Rf4092
RsYN2013
Rs7327
As6526
RsWIV1
Rs3367
Rs9401
Rs4255
SARS−CoV1
RfYNLF 34C
RfYNLF 31C
Rs4247
Rs4237
Rs4231
14628−15824
As6526
Rs7327
Rs9401
Rs4081
Rs4874
Rs4231
Rs672
Rf4092
bRp3
RsGX2013
Rs4255
Rs3367
RsWIV1
Rs4247
RfYNLF 34C
RfYNLF 31C
SARS−CoV1
Rs4237
RsYN2013
Scale: 0.2
15824−18978
RsGX2013
SARS−CoV1
RfYNLF 34C
RfYNLF 31C
bRp3
Rf4092
Rs3367
RsWIV1
As6526
RsYN2013
Rs7327
Rs9401
Rs672
Rs4237
Rs4874
Rs4081
Rs4231
Rs4247
Rs4255
18978−20593
RsYN2013
Rs4231
Rs4874
Rs4255
Rs4247
Rs4081
Rf4092
Rs3367
RsWIV1
As6526
bRp3
RsGX2013
SARS−CoV1
Rs672
RfYNLF 31C
RfYNLF 34C
Rs9401
Rs7327
Rs4237

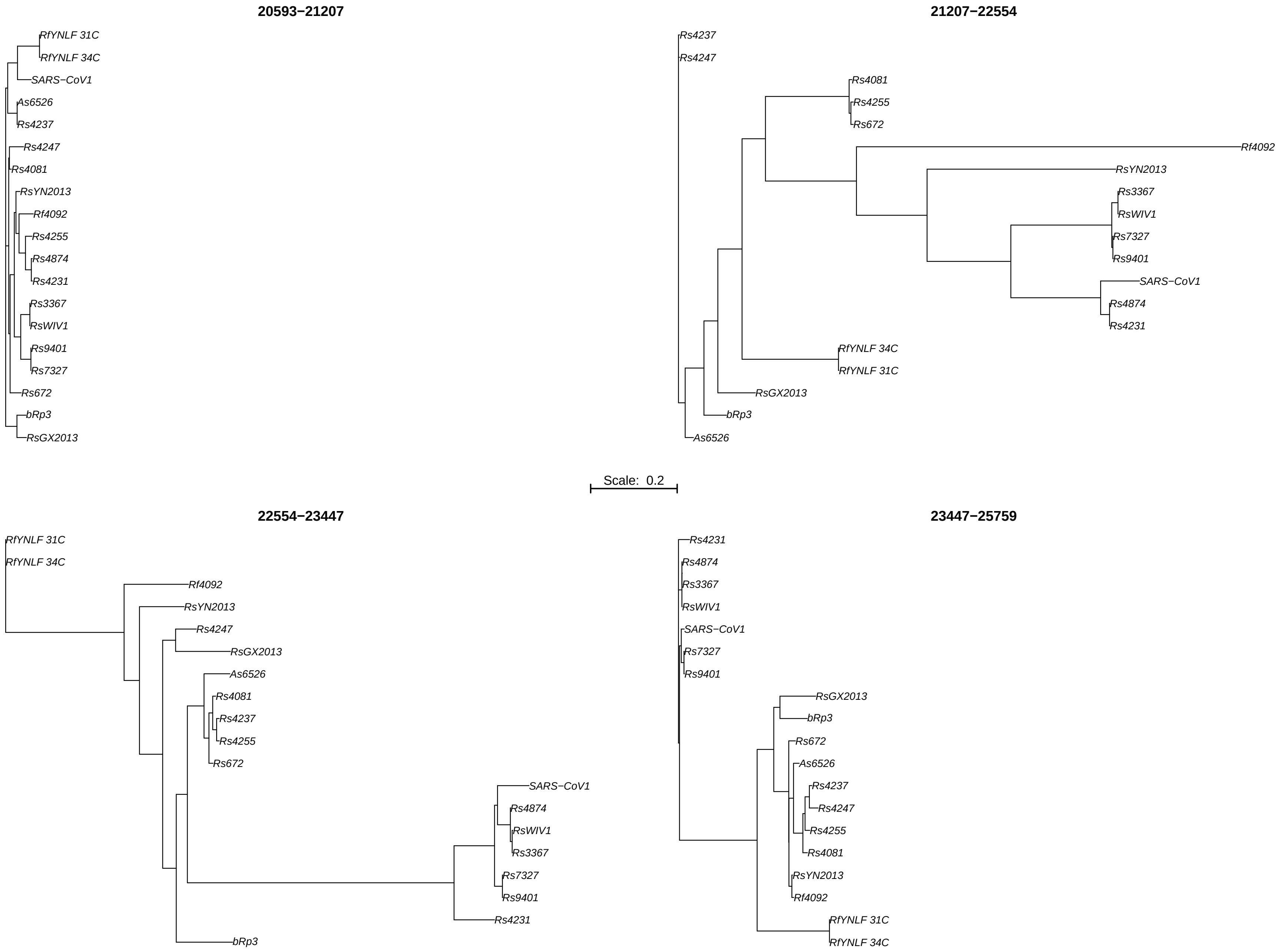
20593−21207
RfYNLF 31C
RfYNLF 34C
SARS−CoV1
As6526
Rs4237
Rs4247
Rs4081
RsYN2013
Rf4092
Rs4255
Rs4874
Rs4231
Rs3367
RsWIV1
Rs9401
Rs7327
Rs672
bRp3
RsGX2013
21207−22554
Rs4237
Rs4247
Rs4081
Rs4255
Rs672
Rf4092
RsYN2013
Rs3367
RsWIV1
Rs7327
Rs9401
SARS−CoV1
Rs4874
Rs4231
RfYNLF 34C
RfYNLF 31C
RsGX2013
bRp3
As6526
Scale: 0.2
22554−23447
RfYNLF 31C
RfYNLF 34C
Rf4092
RsYN2013
Rs4247
RsGX2013
As6526
Rs4081
Rs4237
Rs4255
Rs672
SARS−CoV1
Rs4874
RsWIV1
Rs3367
Rs7327
Rs9401
Rs4231
bRp3
23447−25759
Rs4231
Rs4874
Rs3367
RsWIV1
SARS−CoV1
Rs7327
Rs9401
RsGX2013
bRp3
Rs672
As6526
Rs4237
Rs4247
Rs4255
Rs4081
RsYN2013
Rf4092
RfYNLF 31C
RfYNLF 34C

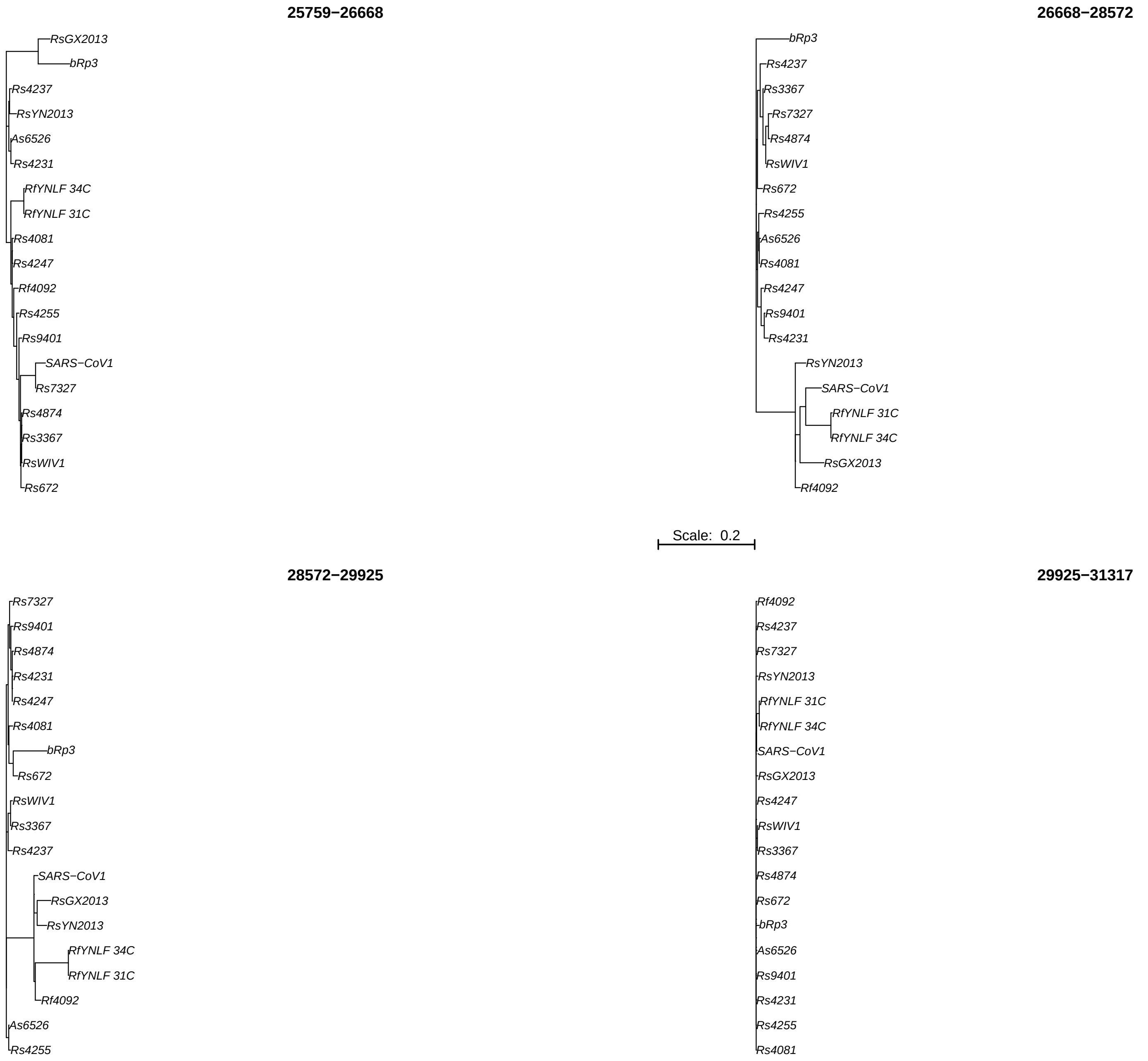
25759−26668
RsGX2013
bRp3
Rs4237
RsYN2013
As6526
Rs4231
RfYNLF 34C
RfYNLF 31C
Rs4081
Rs4247
Rf4092
Rs4255
Rs9401
SARS−CoV1
Rs7327
Rs4874
Rs3367
RsWIV1
Rs672
26668−28572
bRp3
Rs4237
Rs3367
Rs7327
Rs4874
RsWIV1
Rs672
Rs4255
As6526
Rs4081
Rs4247
Rs9401
Rs4231
RsYN2013
SARS−CoV1
RfYNLF 31C
RfYNLF 34C
RsGX2013
Rf4092
Scale: 0.2
28572−29925
Rs7327
Rs9401
Rs4874
Rs4231
Rs4247
Rs4081
bRp3
Rs672
RsWIV1
Rs3367
Rs4237
SARS−CoV1
RsGX2013
RsYN2013
RfYNLF 34C
RfYNLF 31C
Rf4092
As6526
Rs4255
29925−31317
Rf4092
Rs4237
Rs7327
RsYN2013
RfYNLF 31C
RfYNLF 34C
SARS−CoV1
RsGX2013
Rs4247
RsWIV1
Rs3367
Rs4874
Rs672
bRp3
As6526
Rs9401
Rs4231
Rs4255
Rs4081

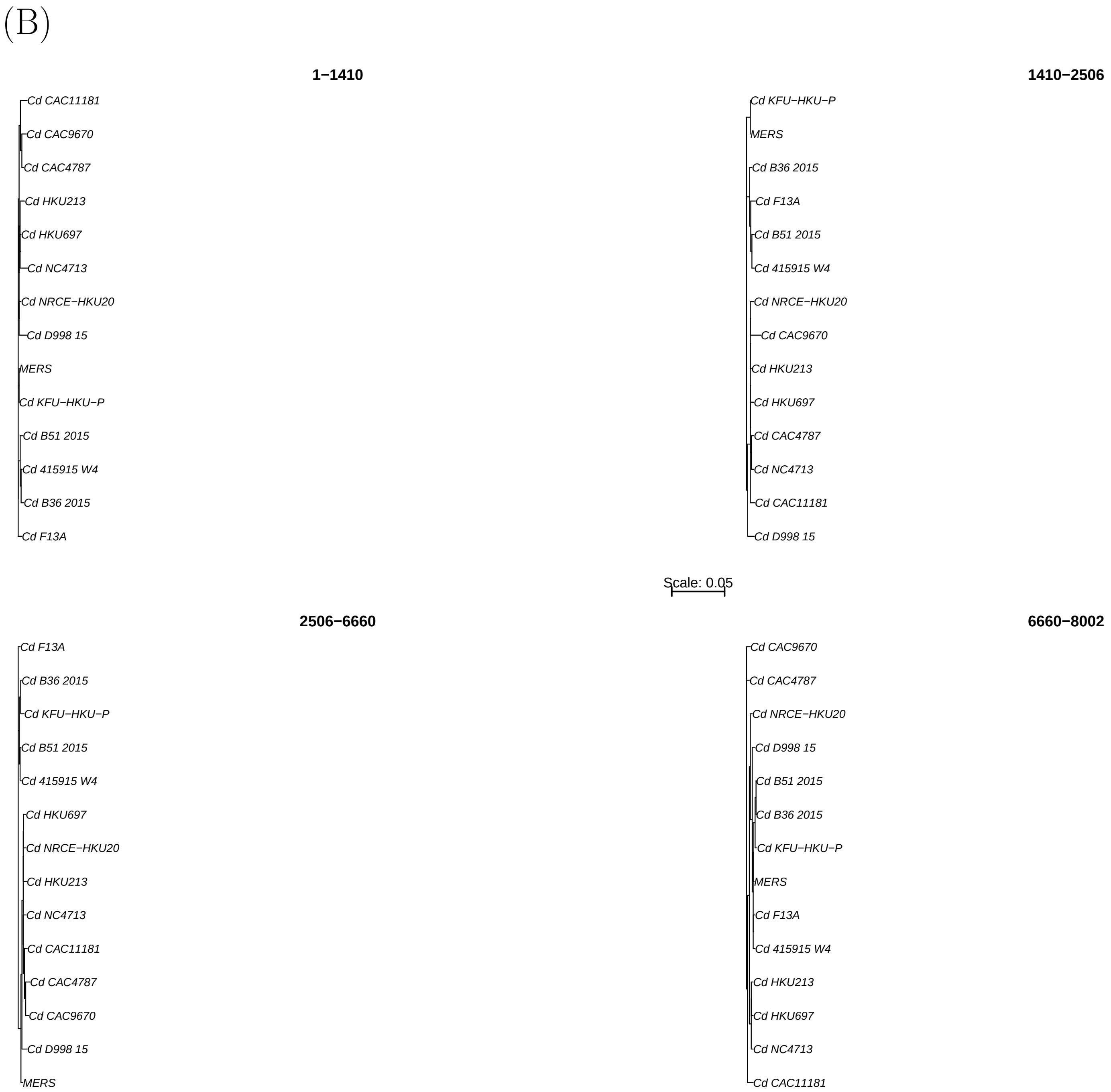
(B)
1−1410
Cd CAC11181
Cd CAC9670
Cd CAC4787
Cd HKU213
Cd HKU697
Cd NC4713
Cd NRCE−HKU20
Cd D998 15
MERS
Cd KFU−HKU−P
Cd B51 2015
Cd 415915 W4
Cd B36 2015
Cd F13A
1410−2506
Cd KFU−HKU−P
MERS
Cd B36 2015
Cd F13A
Cd B51 2015
Cd 415915 W4
Cd NRCE−HKU20
Cd CAC9670
Cd HKU213
Cd HKU697
Cd CAC4787
Cd NC4713
Cd CAC11181
Cd D998 15
Scale: 0.05
2506−6660
Cd F13A
Cd B36 2015
Cd KFU−HKU−P
Cd B51 2015
Cd 415915 W4
Cd HKU697
Cd NRCE−HKU20
Cd HKU213
Cd NC4713
Cd CAC11181
Cd CAC4787
Cd CAC9670
Cd D998 15
MERS
6660−8002
Cd CAC9670
Cd CAC4787
Cd NRCE−HKU20
Cd D998 15
Cd B51 2015
Cd B36 2015
Cd KFU−HKU−P
MERS
Cd F13A
Cd 415915 W4
Cd HKU213
Cd HKU697
Cd NC4713
Cd CAC11181

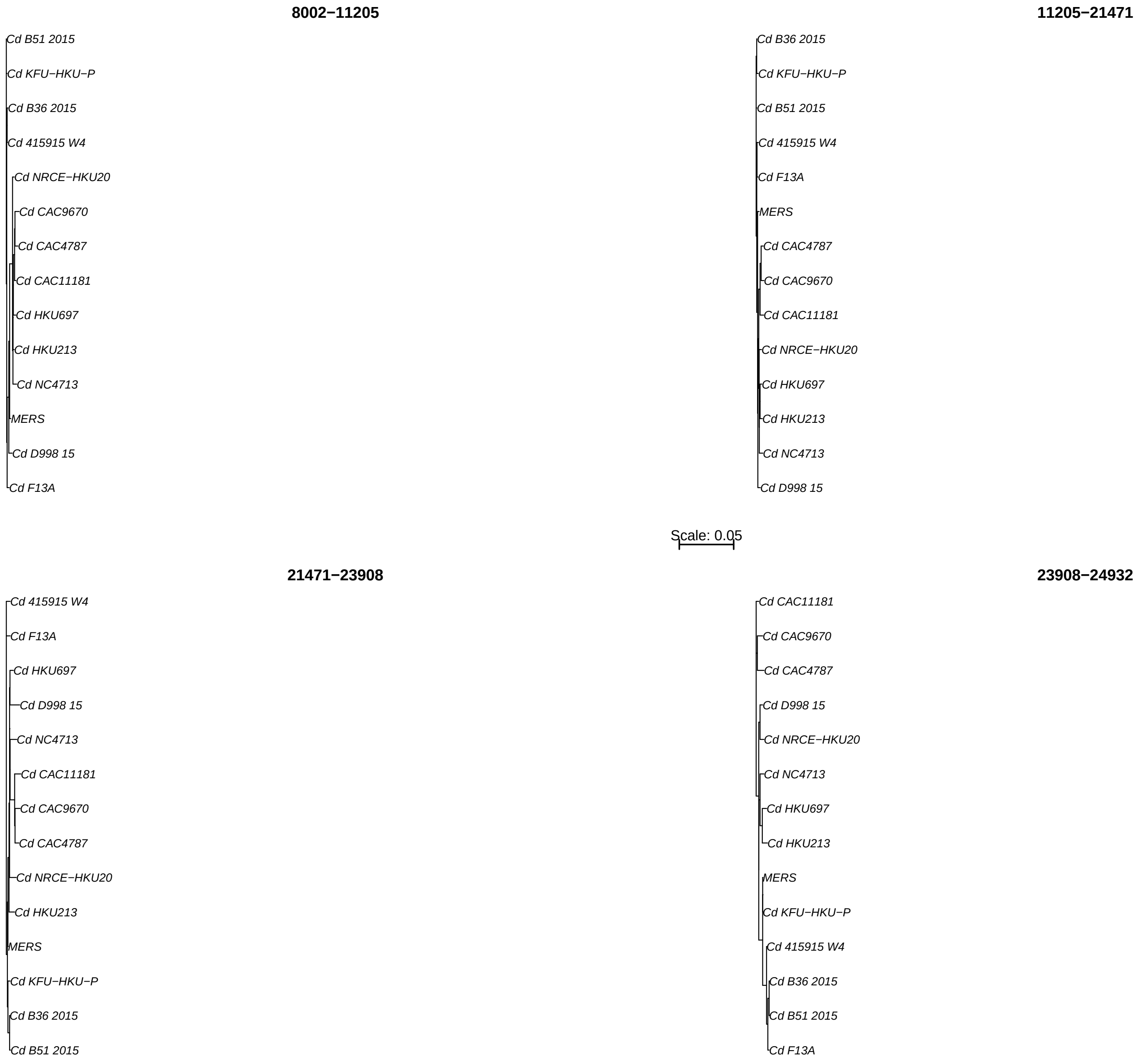
8002−11205
Cd B51 2015
Cd KFU−HKU−P
Cd B36 2015
Cd 415915 W4
Cd NRCE−HKU20
Cd CAC9670
Cd CAC4787
Cd CAC11181
Cd HKU697
Cd HKU213
Cd NC4713
MERS
Cd D998 15
Cd F13A
11205−21471
Cd B36 2015
Cd KFU−HKU−P
Cd B51 2015
Cd 415915 W4
Cd F13A
MERS
Cd CAC4787
Cd CAC9670
Cd CAC11181
Cd NRCE−HKU20
Cd HKU697
Cd HKU213
Cd NC4713
Cd D998 15
Scale: 0.05
21471−23908
Cd 415915 W4
Cd F13A
Cd HKU697
Cd D998 15
Cd NC4713
Cd CAC11181
Cd CAC9670
Cd CAC4787
Cd NRCE−HKU20
Cd HKU213
MERS
Cd KFU−HKU−P
Cd B36 2015
Cd B51 2015
23908−24932
Cd CAC11181
Cd CAC9670
Cd CAC4787
Cd D998 15
Cd NRCE−HKU20
Cd NC4713
Cd HKU697
Cd HKU213
MERS
Cd KFU−HKU−P
Cd 415915 W4
Cd B36 2015
Cd B51 2015
Cd F13A

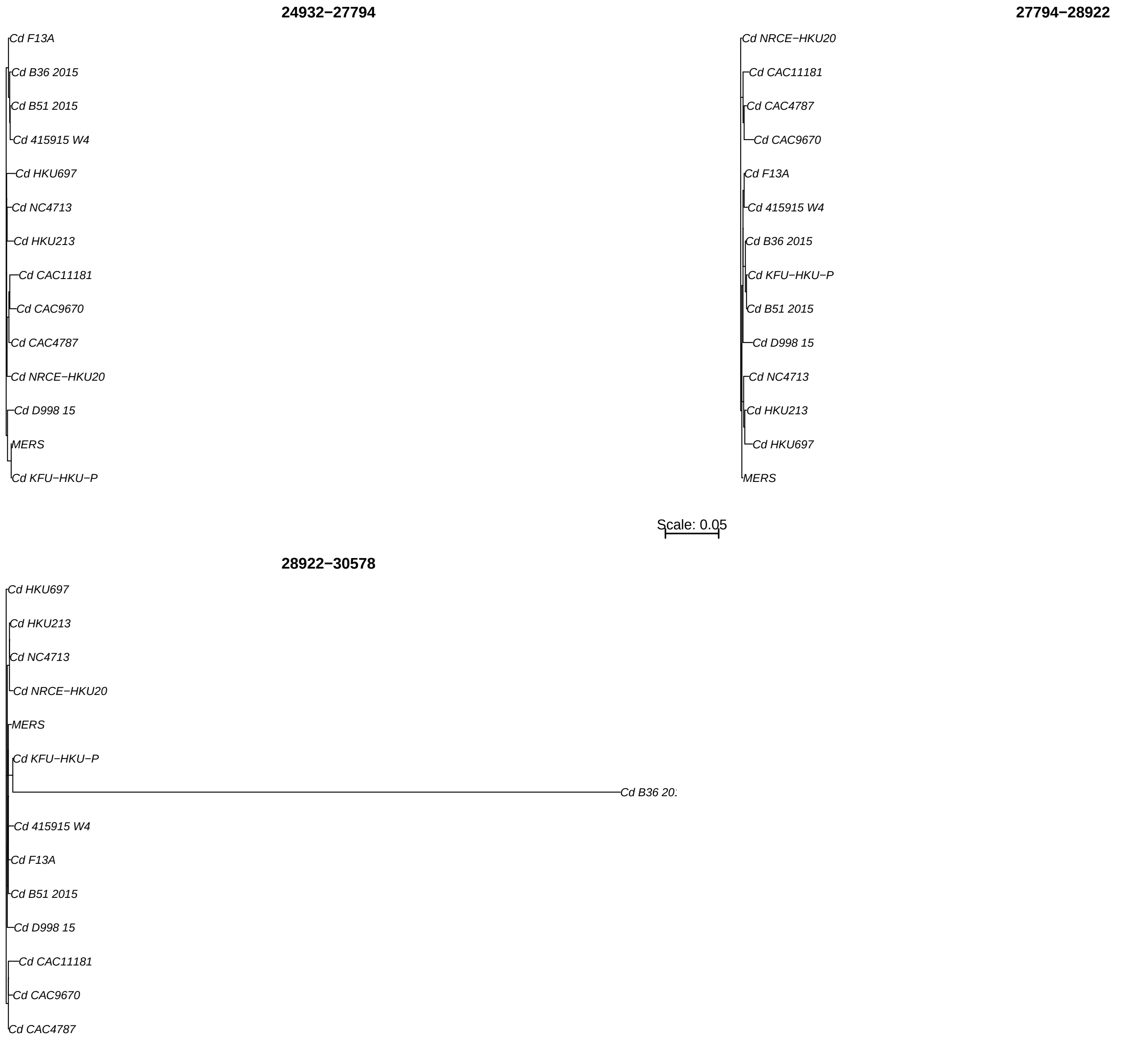
24932−27794
Cd F13A
Cd B36 2015
Cd B51 2015
Cd 415915 W4
Cd HKU697
Cd NC4713
Cd HKU213
Cd CAC11181
Cd CAC9670
Cd CAC4787
Cd NRCE−HKU20
Cd D998 15
MERS
Cd KFU−HKU−P
27794−28922
Cd NRCE−HKU20
Cd CAC11181
Cd CAC4787
Cd CAC9670
Cd F13A
Cd 415915 W4
Cd B36 2015
Cd KFU−HKU−P
Cd B51 2015
Cd D998 15
Cd NC4713
Cd HKU213
Cd HKU697
MERS
Scale: 0.05
28922−30578
Cd HKU697
Cd HKU213
Cd NC4713
Cd NRCE−HKU20
MERS
Cd KFU−HKU−P
Cd B36 20
Cd 415915 W4
Cd F13A
Cd B51 2015
Cd D998 15
Cd CAC11181
Cd CAC9670
Cd CAC4787

(C)

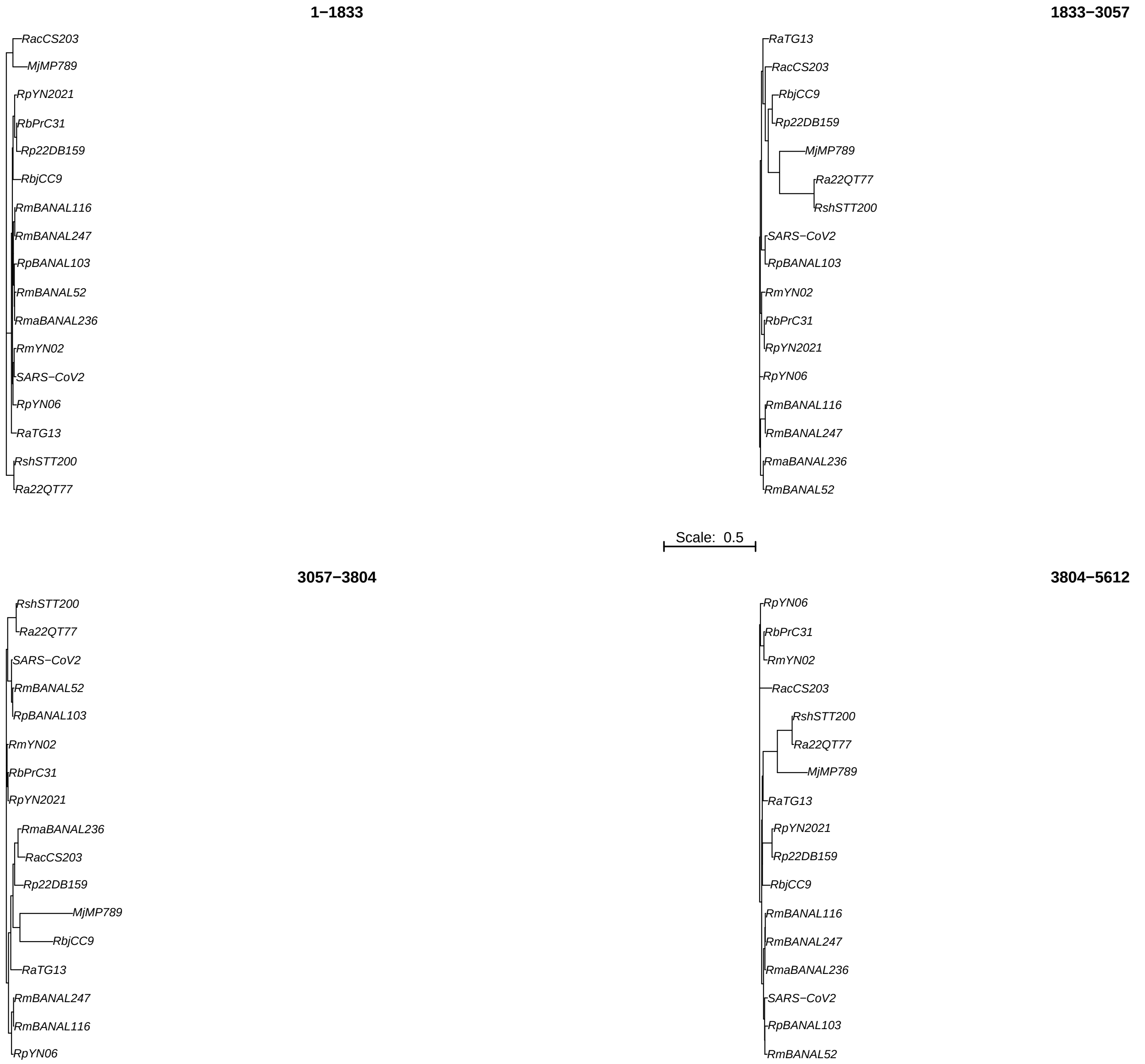

1−1833
RacCS203
MjMP789
RpYN2021
RbPrC31
Rp22DB159
RbjCC9
RmBANAL116
RmBANAL247
RpBANAL103
RmBANAL52
RmaBANAL236
RmYN02
SARS−CoV2
RpYN06
RaTG13
RshSTT200
Ra22QT77
1833−3057
RaTG13
RacCS203
RbjCC9
Rp22DB159
MjMP789
Ra22QT77
RshSTT200
SARS−CoV2
RpBANAL103
RmYN02
RbPrC31
RpYN2021
RpYN06
RmBANAL116
RmBANAL247
RmaBANAL236
RmBANAL52
Scale: 0.5
3057−3804
RshSTT200
Ra22QT77
SARS−CoV2
RmBANAL52
RpBANAL103
RmYN02
RbPrC31
RpYN2021
RmaBANAL236
RacCS203
Rp22DB159
MjMP789
RbjCC9
RaTG13
RmBANAL247
RmBANAL116
RpYN06
3804−5612
RpYN06
RbPrC31
RmYN02
RacCS203
RshSTT200
Ra22QT77
MjMP789
RaTG13
RpYN2021
Rp22DB159
RbjCC9
RmBANAL116
RmBANAL247
RmaBANAL236
SARS−CoV2
RpBANAL103
RmBANAL52

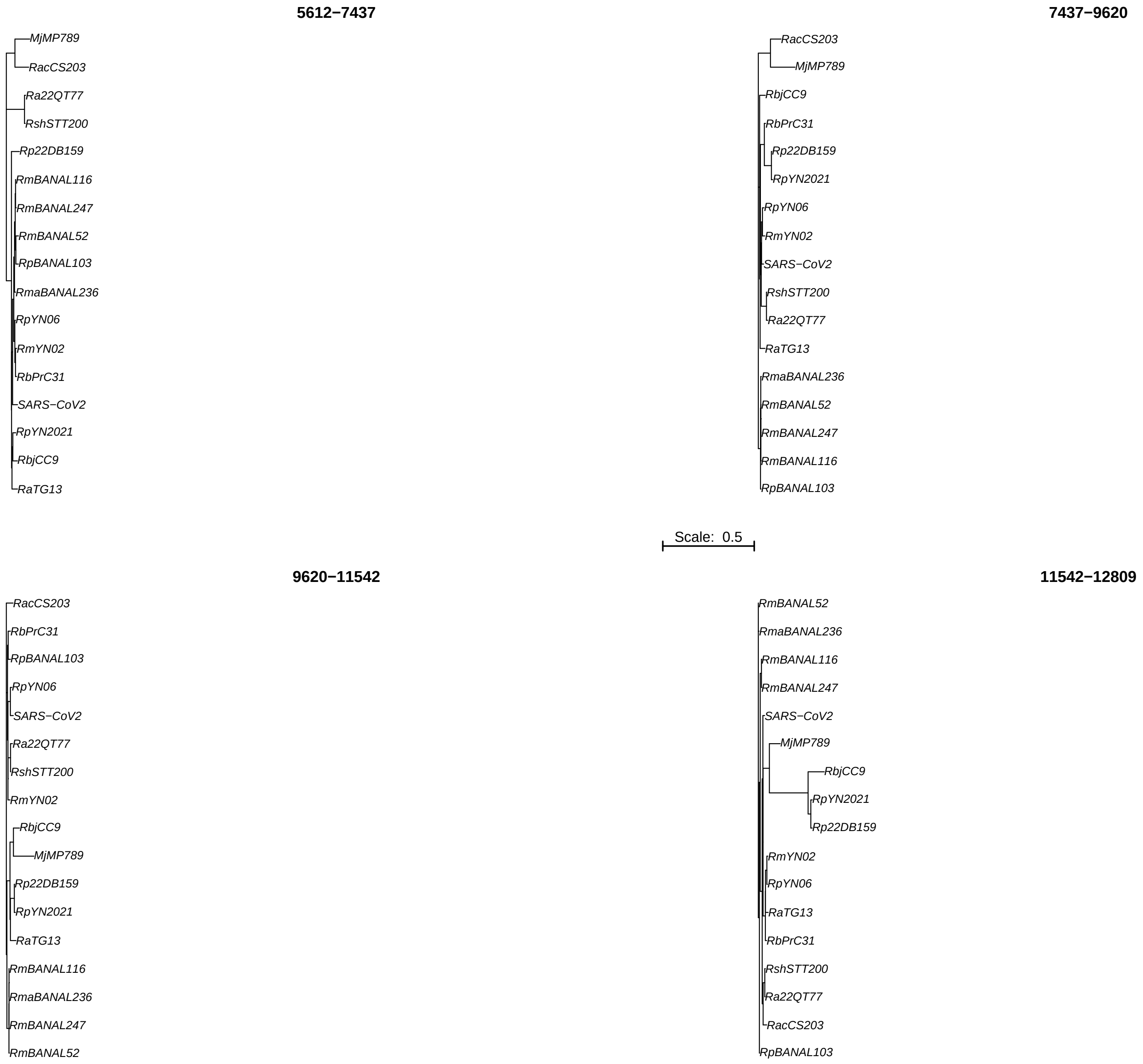
5612−7437
MjMP789
RacCS203
Ra22QT77
RshSTT200
Rp22DB159
RmBANAL116
RmBANAL247
RmBANAL52
RpBANAL103
RmaBANAL236
RpYN06
RmYN02
RbPrC31
SARS−CoV2
RpYN2021
RbjCC9
RaTG13
7437−9620
RacCS203
MjMP789
RbjCC9
RbPrC31
Rp22DB159
RpYN2021
RpYN06
RmYN02
SARS−CoV2
RshSTT200
Ra22QT77
RaTG13
RmaBANAL236
RmBANAL52
RmBANAL247
RmBANAL116
RpBANAL103
Scale: 0.5
9620−11542
RacCS203
RbPrC31
RpBANAL103
RpYN06
SARS−CoV2
Ra22QT77
RshSTT200
RmYN02
RbjCC9
MjMP789
Rp22DB159
RpYN2021
RaTG13
RmBANAL116
RmaBANAL236
RmBANAL247
RmBANAL52
11542−12809
RmBANAL52
RmaBANAL236
RmBANAL116
RmBANAL247
SARS−CoV2
MjMP789
RbjCC9
RpYN2021
Rp22DB159
RmYN02
RpYN06
RaTG13
RbPrC31
RshSTT200
Ra22QT77
RacCS203
RpBANAL103

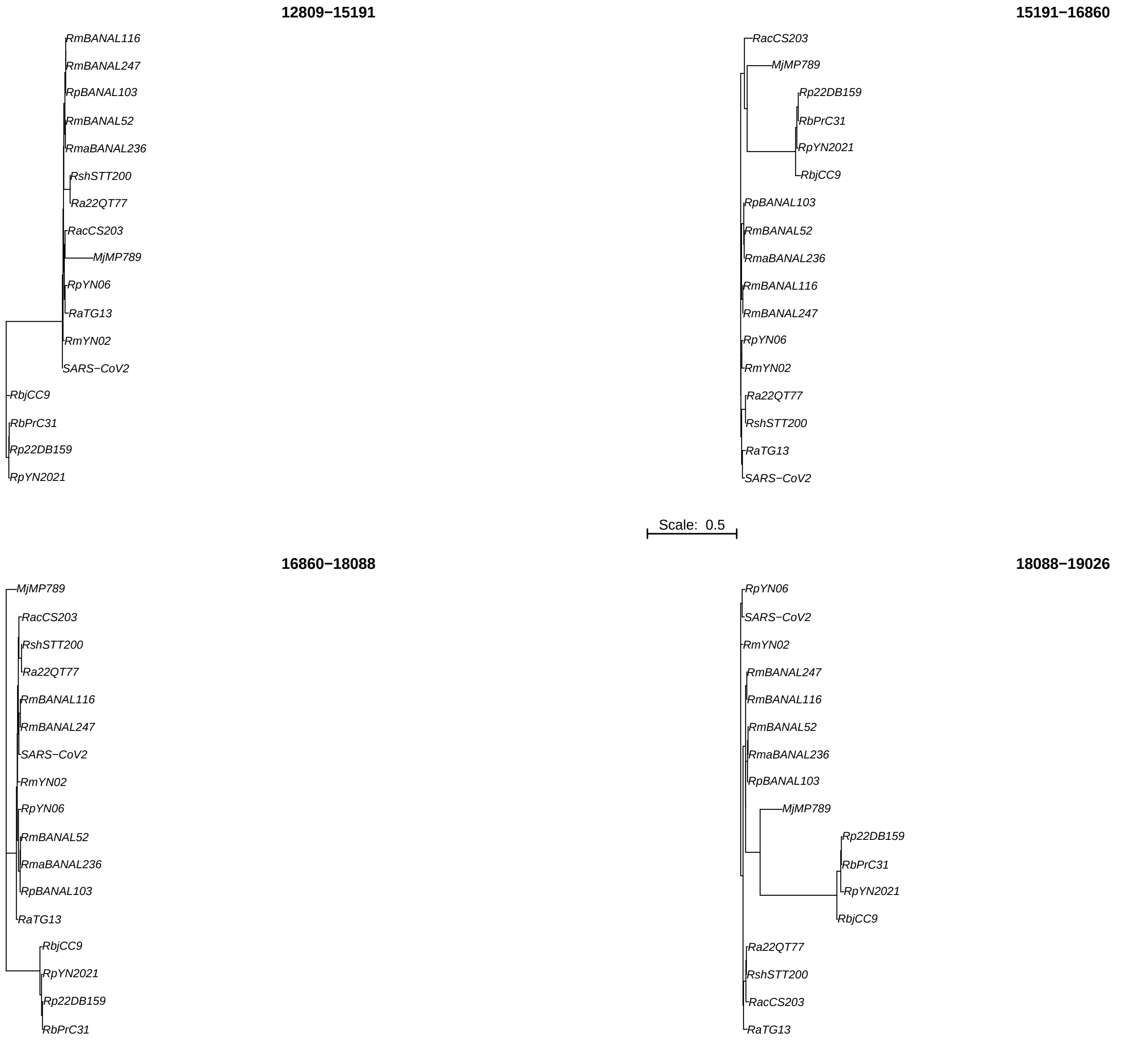
12809−15191
RmBANAL116
RmBANAL247
RpBANAL103
RmBANAL52
RmaBANAL236
RshSTT200
Ra22QT77
RacCS203
MjMP789
RpYN06
RaTG13
RmYN02
SARS−CoV2
RbjCC9
RbPrC31
Rp22DB159
RpYN2021
15191−16860
RacCS203
MjMP789
Rp22DB159
RbPrC31
RpYN2021
RbjCC9
RpBANAL103
RmBANAL52
RmaBANAL236
RmBANAL116
RmBANAL247
RpYN06
RmYN02
Ra22QT77
RshSTT200
RaTG13
SARS−CoV2
Scale: 0.5
16860−18088
MjMP789
RacCS203
RshSTT200
Ra22QT77
RmBANAL116
RmBANAL247
SARS−CoV2
RmYN02
RpYN06
RmBANAL52
RmaBANAL236
RpBANAL103
RaTG13
RbjCC9
RpYN2021
Rp22DB159
RbPrC31
18088−19026
RpYN06
SARS−CoV2
RmYN02
RmBANAL247
RmBANAL116
RmBANAL52
RmaBANAL236
RpBANAL103
MjMP789
Rp22DB159
RbPrC31
RpYN2021
RbjCC9
Ra22QT77
RshSTT200
RacCS203
RaTG13

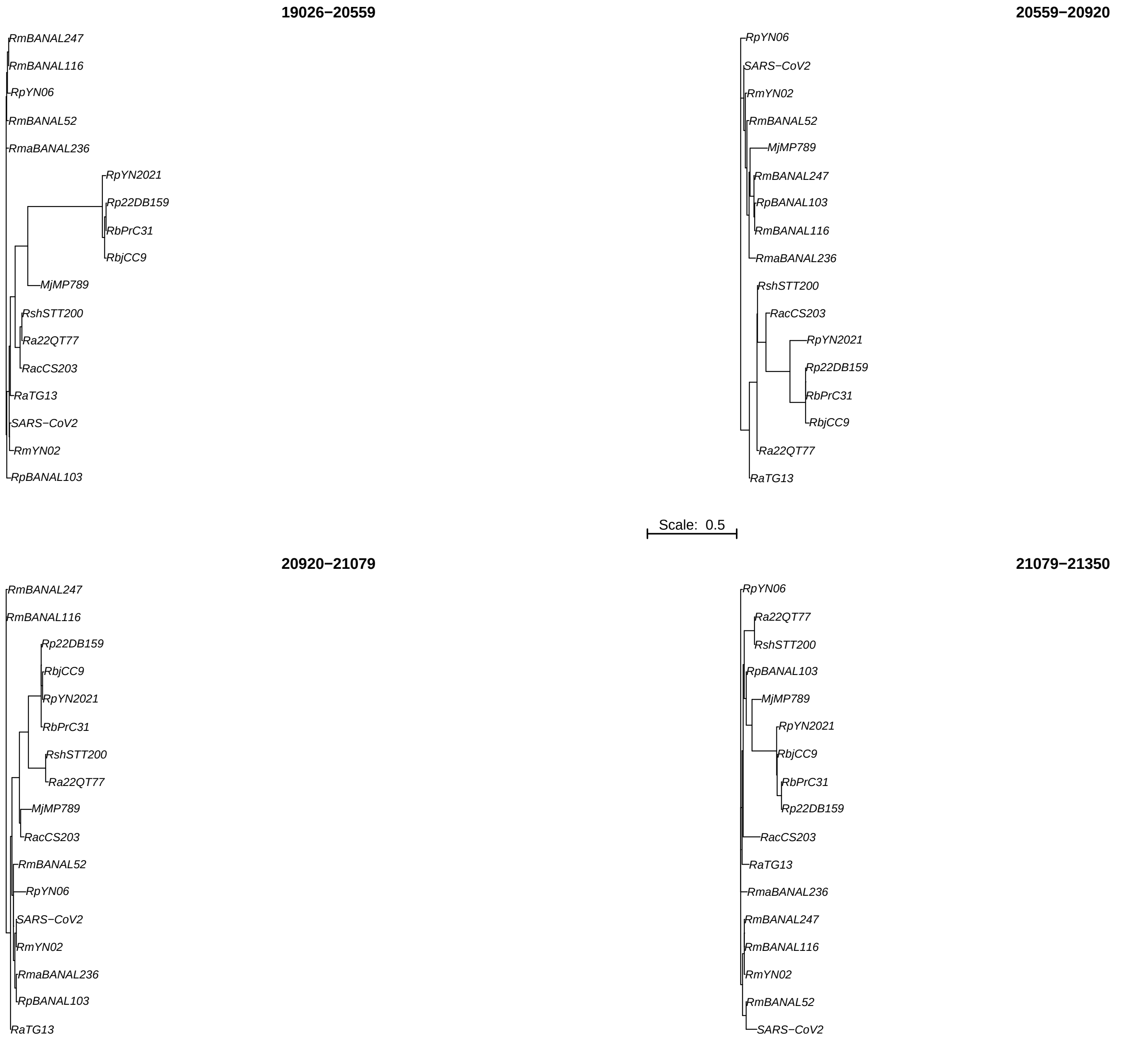
19026−20559
RmBANAL247
RmBANAL116
RpYN06
RmBANAL52
RmaBANAL236
RpYN2021
Rp22DB159
RbPrC31
RbjCC9
MjMP789
RshSTT200
Ra22QT77
RacCS203
RaTG13
SARS−CoV2
RmYN02
RpBANAL103
20559−20920
RpYN06
SARS−CoV2
RmYN02
RmBANAL52
MjMP789
RmBANAL247
RpBANAL103
RmBANAL116
RmaBANAL236
RshSTT200
RacCS203
RpYN2021
Rp22DB159
RbPrC31
RbjCC9
Ra22QT77
RaTG13
Scale: 0.5
20920−21079
RmBANAL247
RmBANAL116
Rp22DB159
RbjCC9
RpYN2021
RbPrC31
RshSTT200
Ra22QT77
MjMP789
RacCS203
RmBANAL52
RpYN06
SARS−CoV2
RmYN02
RmaBANAL236
RpBANAL103
RaTG13
21079−21350
RpYN06
Ra22QT77
RshSTT200
RpBANAL103
MjMP789
RpYN2021
RbjCC9
RbPrC31
Rp22DB159
RacCS203
RaTG13
RmaBANAL236
RmBANAL247
RmBANAL116
RmYN02
RmBANAL52
SARS−CoV2

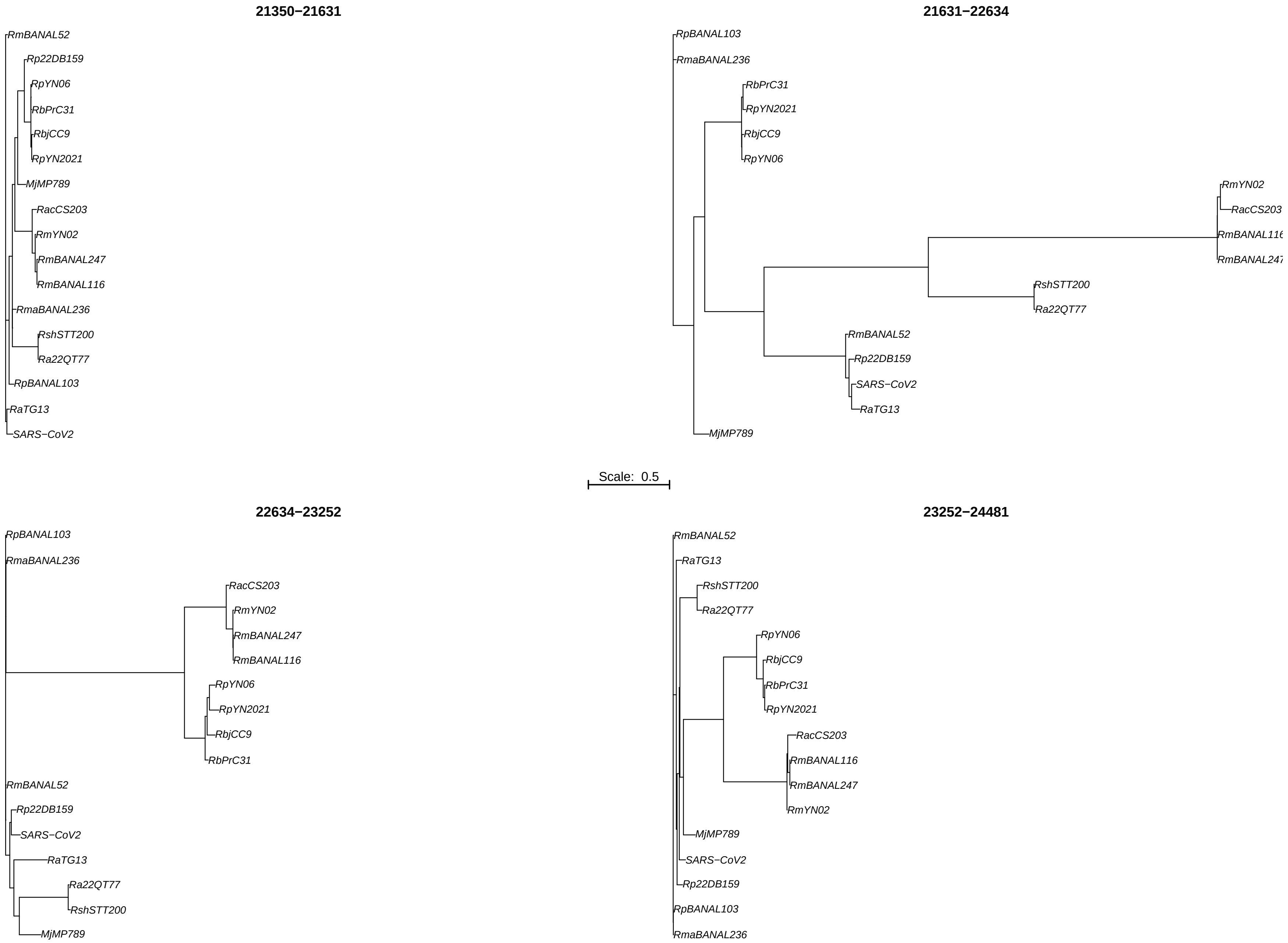
21350−21631
RmBANAL52
Rp22DB159
RpYN06
RbPrC31
RbjCC9
RpYN2021
MjMP789
RacCS203
RmYN02
RmBANAL247
RmBANAL116
RmaBANAL236
RshSTT200
Ra22QT77
RpBANAL103
RaTG13
SARS−CoV2
21631−22634
RpBANAL103
RmaBANAL236
RbPrC31
RpYN2021
RbjCC9
RpYN06
RmYN02
RacCS203
RshSTT200
Ra22QT77
RmBANAL52
Rp22DB159
SARS−CoV2
RaTG13
MjMP789
Scale: 0.5
22634−23252
RpBANAL103
RmaBANAL236
RacCS203
RmYN02
RmBANAL247
RmBANAL116
RpYN06
RpYN2021
RbjCC9
RbPrC31
RmBANAL52
Rp22DB159
SARS−CoV2
RaTG13
Ra22QT77
RshSTT200
MjMP789
23252−24481
RmBANAL52
RaTG13
RshSTT200
Ra22QT77
RpYN06
RbjCC9
RbPrC31
RpYN2021
RacCS203
RmBANAL116
RmBANAL247
RmYN02
MjMP789
SARS−CoV2
Rp22DB159
RpBANAL103
RmaBANAL236

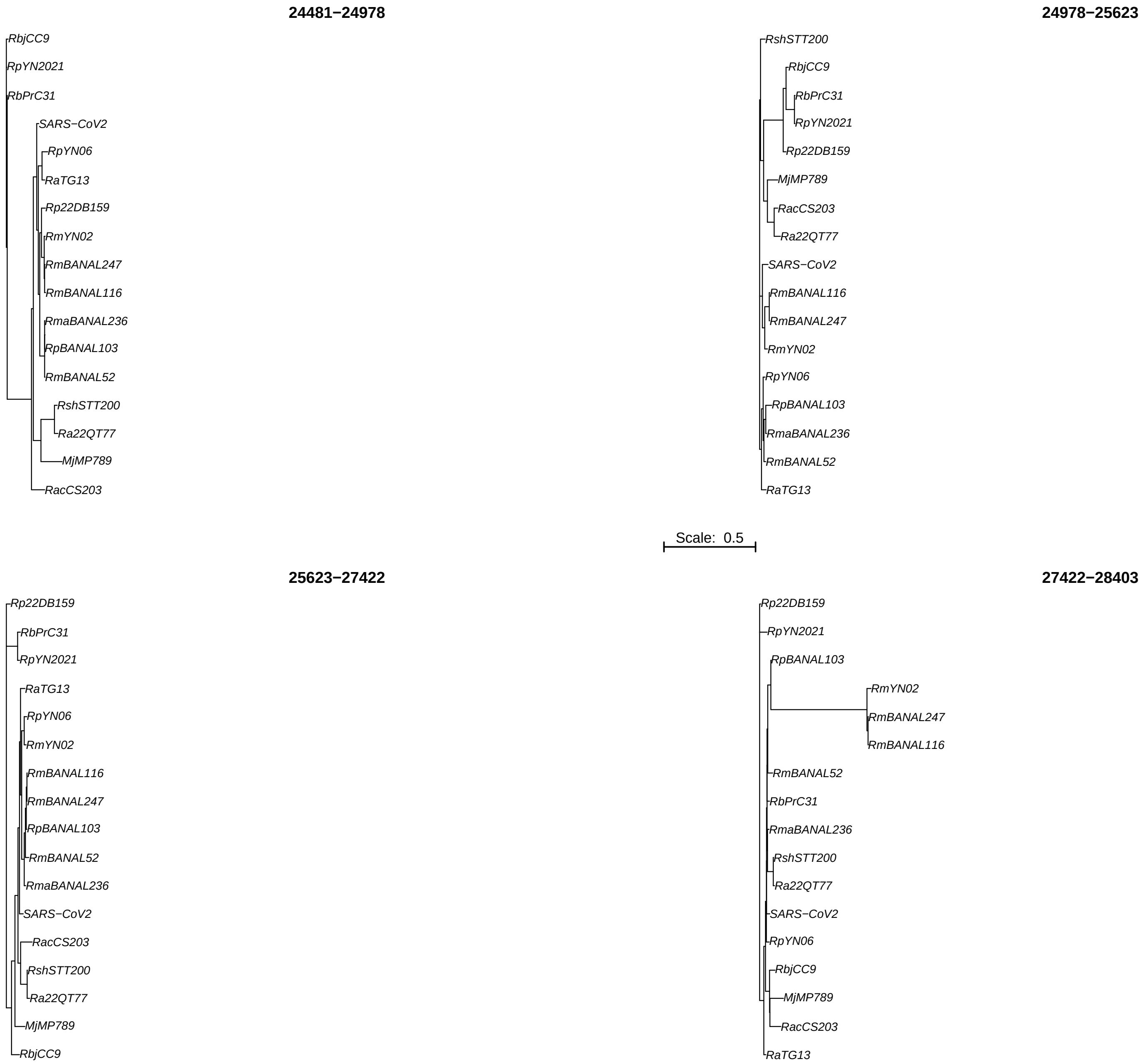
24481−24978
RbjCC9
RpYN2021
RbPrC31
SARS−CoV2
RpYN06
RaTG13
Rp22DB159
RmYN02
RmBANAL247
RmBANAL116
RmaBANAL236
RpBANAL103
RmBANAL52
RshSTT200
Ra22QT77
MjMP789
RacCS203
24978−25623
RshSTT200
RbjCC9
RbPrC31
RpYN2021
Rp22DB159
MjMP789
RacCS203
Ra22QT77
SARS−CoV2
RmBANAL116
RmBANAL247
RmYN02
RpYN06
RpBANAL103
RmaBANAL236
RmBANAL52
RaTG13
Scale: 0.5
25623−27422
Rp22DB159
RbPrC31
RpYN2021
RaTG13
RpYN06
RmYN02
RmBANAL116
RmBANAL247
RpBANAL103
RmBANAL52
RmaBANAL236
SARS−CoV2
RacCS203
RshSTT200
Ra22QT77
MjMP789
RbjCC9
27422−28403
Rp22DB159
RpYN2021
RpBANAL103
RmYN02
RmBANAL247
RmBANAL116
RmBANAL52
RbPrC31
RmaBANAL236
RshSTT200
Ra22QT77
SARS−CoV2
RpYN06
RbjCC9
MjMP789
RacCS203
RaTG13

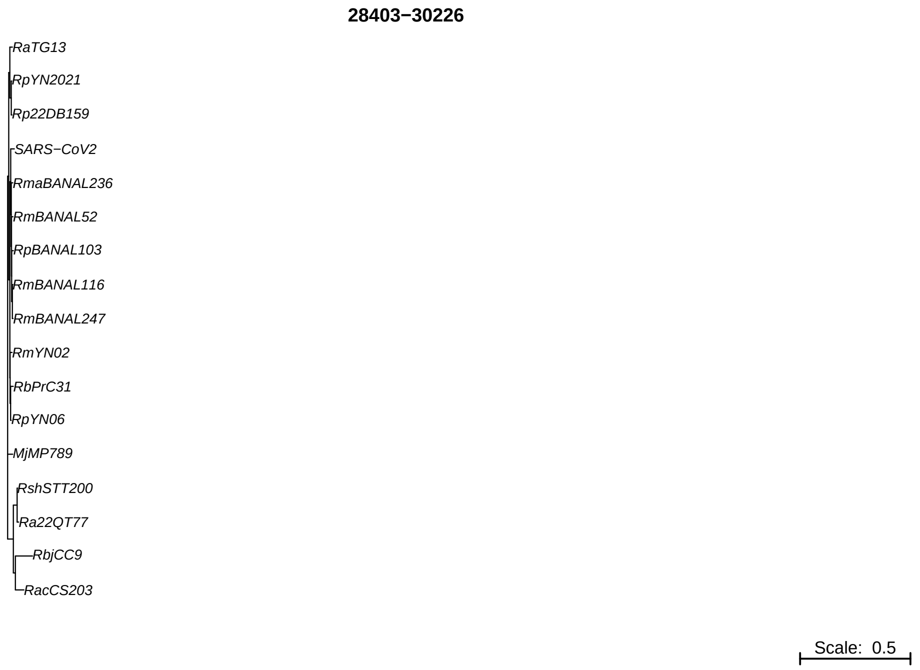
28403−30226
RaTG13
RpYN2021
Rp22DB159
SARS−CoV2
RmaBANAL236
RmBANAL52
RpBANAL103
RmBANAL116
RmBANAL247
RmYN02
RbPrC31
RpYN06
MjMP789
RshSTT200
Ra22QT77
RbjCC9
RacCS203
Scale: 0.5

Figure S11. Cladograms of all genomic segments for the three reference datasets. (A) SARS-CoV-1, (B) MERS, and (C) SARS-CoV-2. Each tree corresponds to one segment, identified by its nucleotide range in the title. Trees are shown as cladograms to emphasize branching structure and sequence grouping; branch lengths are therefore not interpreted as phylogenetic distances. Bootstrap support values are displayed at internal nodes.

(A)

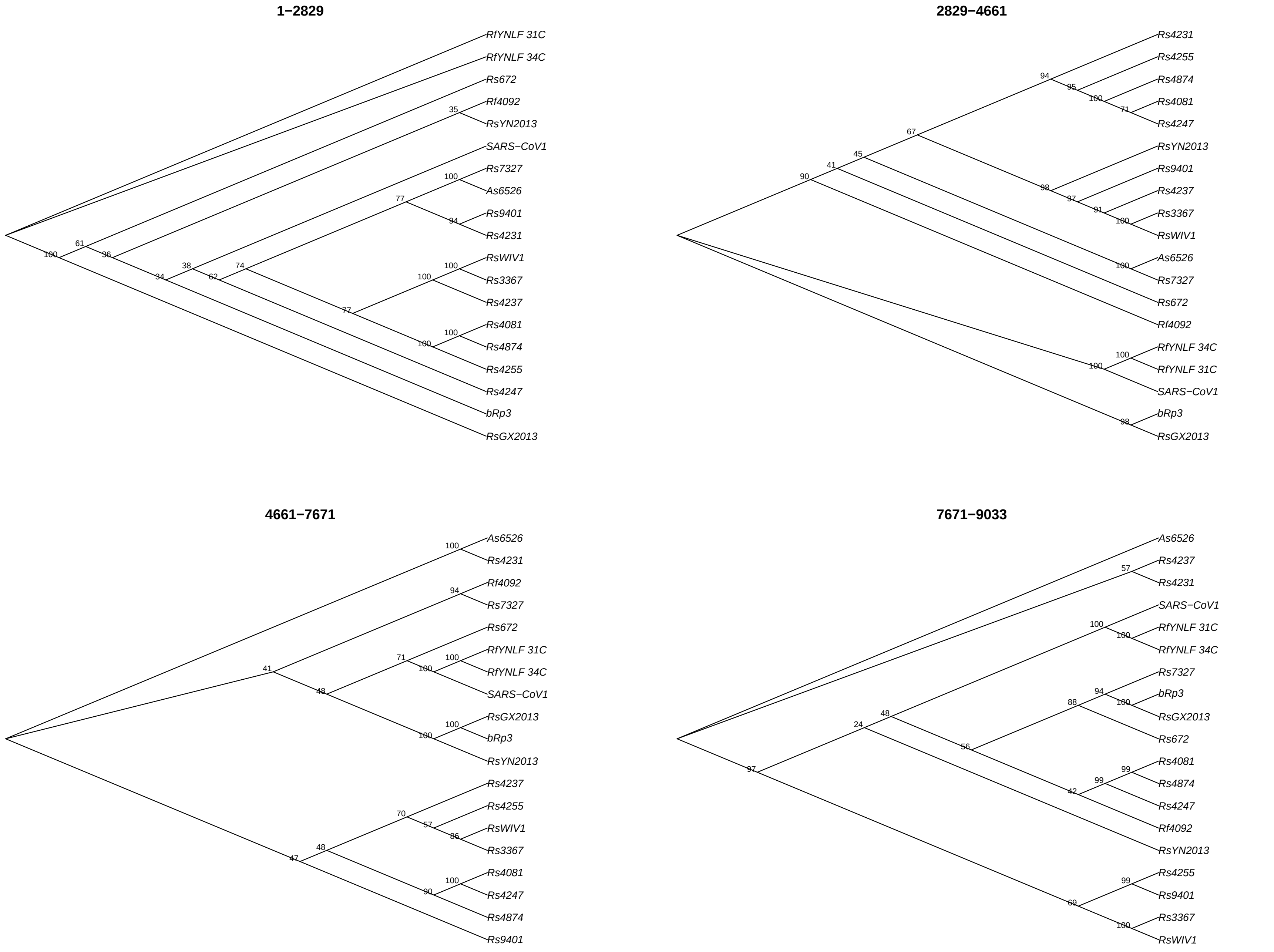
1−2829
RfYNLF 31C
RfYNLF 34C
Rs672
Rf4092
RsYN2013
SARS−CoV1
Rs7327
As6526
Rs9401
Rs4231
RsWIV1
Rs3367
Rs4237
Rs4081
Rs4874
Rs4255
Rs4247
bRp3
RsGX2013
2829−4661
Rs4231
Rs4255
Rs4874
Rs4081
Rs4247
RsYN2013
Rs9401
Rs4237
Rs3367
RsWIV1
As6526
Rs7327
Rs672
Rf4092
RfYNLF 34C
RfYNLF 31C
SARS−CoV1
bRp3
RsGX2013
4661−7671
As6526
Rs4231
Rf4092
Rs7327
Rs672
RfYNLF 31C
RfYNLF 34C
SARS−CoV1
RsGX2013
bRp3
RsYN2013
Rs4237
Rs4255
RsWIV1
Rs3367
Rs4081
Rs4247
Rs4874
Rs9401
7671−9033
As6526
Rs4237
Rs4231
SARS−CoV1
RfYNLF 31C
RfYNLF 34C
Rs7327
bRp3
RsGX2013
Rs672
Rs4081
Rs4874
Rs4247
Rf4092
RsYN2013
Rs4255
Rs9401
Rs3367
RsWIV1

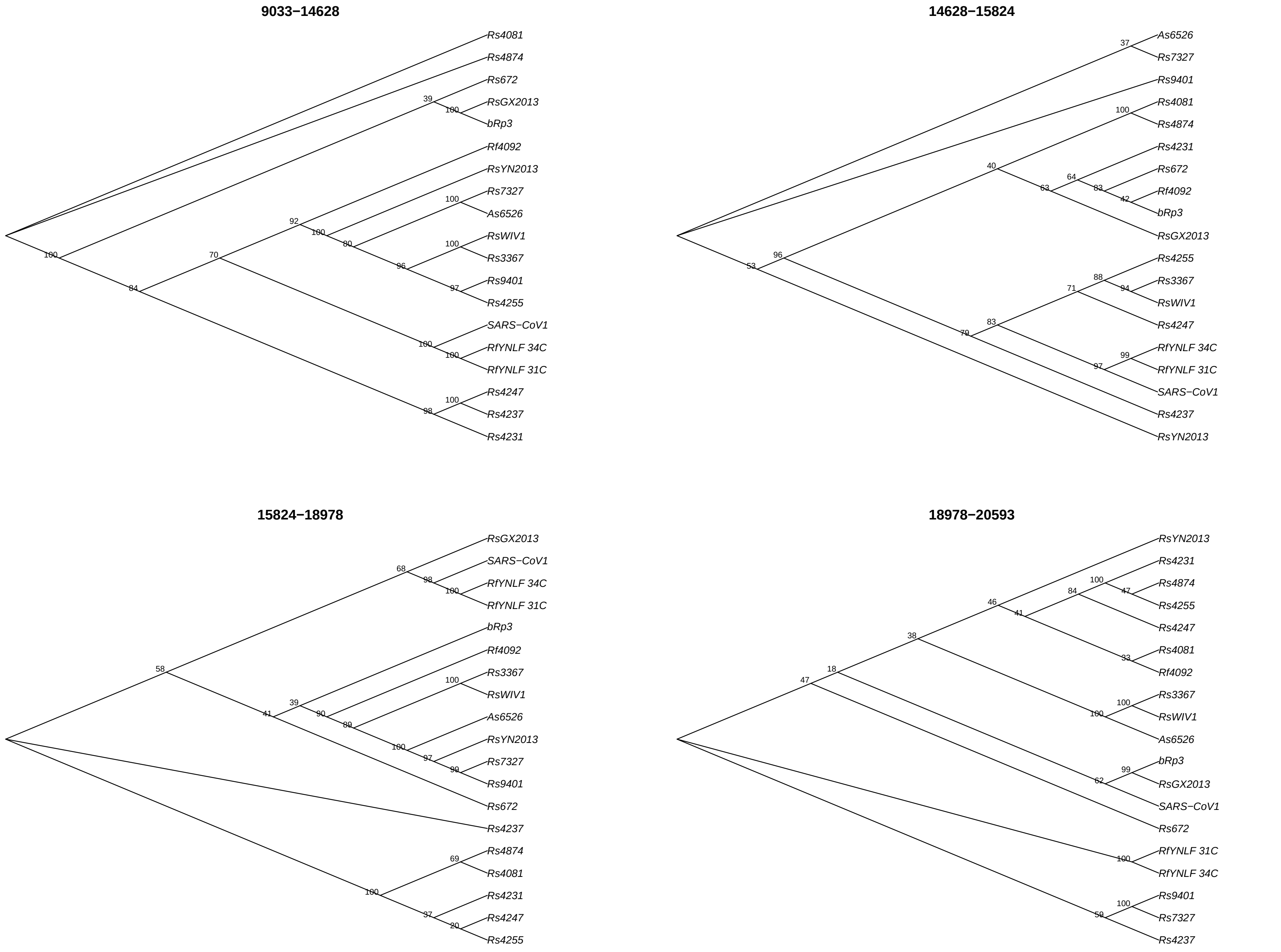
9033−14628
Rs4081
Rs4874
Rs672
RsGX2013
bRp3
Rf4092
RsYN2013
Rs7327
As6526
RsWIV1
Rs3367
Rs9401
Rs4255
SARS−CoV1
RfYNLF 34C
RfYNLF 31C
Rs4247
Rs4237
Rs4231
14628−15824
As6526
Rs7327
Rs9401
Rs4081
Rs4874
Rs4231
Rs672
Rf4092
bRp3
RsGX2013
Rs4255
Rs3367
RsWIV1
Rs4247
RfYNLF 34C
RfYNLF 31C
SARS−CoV1
Rs4237
RsYN2013
15824−18978
RsGX2013
SARS−CoV1
RfYNLF 34C
RfYNLF 31C
bRp3
Rf4092
Rs3367
RsWIV1
As6526
RsYN2013
Rs7327
Rs9401
Rs672
Rs4237
Rs4874
Rs4081
Rs4231
Rs4247
Rs4255
18978−20593
RsYN2013
Rs4231
Rs4874
Rs4255
Rs4247
Rs4081
Rf4092
Rs3367
RsWIV1
As6526
bRp3
RsGX2013
SARS−CoV1
Rs672
RfYNLF 31C
RfYNLF 34C
Rs9401
Rs7327
Rs4237

**20593−21207**

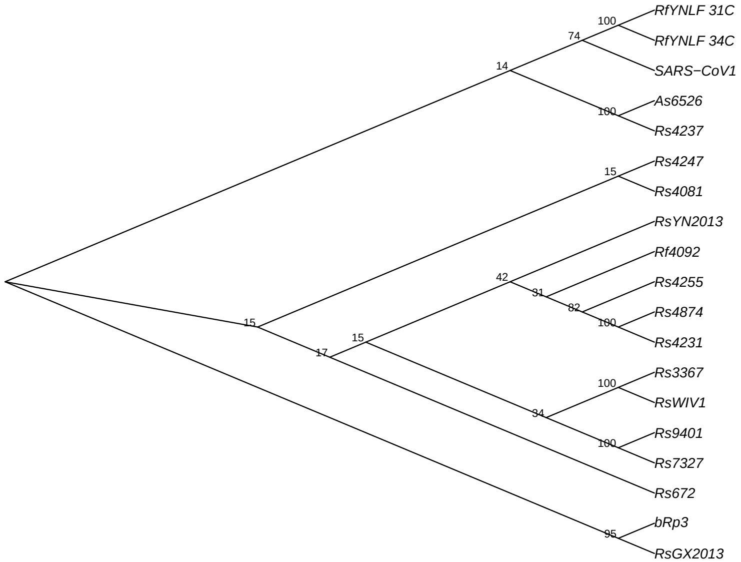

**21207−22554**

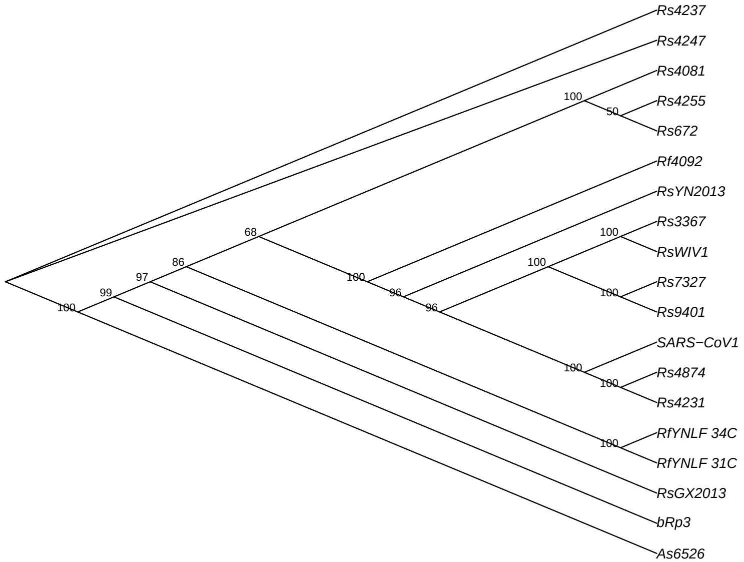

**22554−23447**

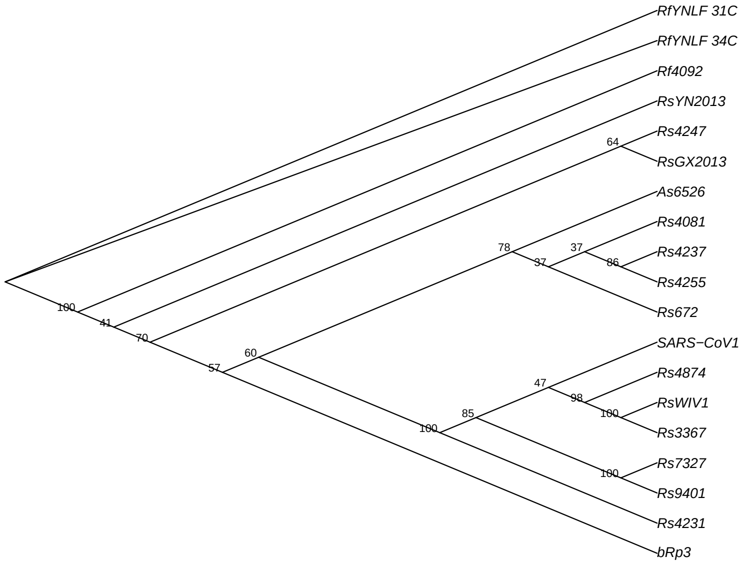

**23447−25759**

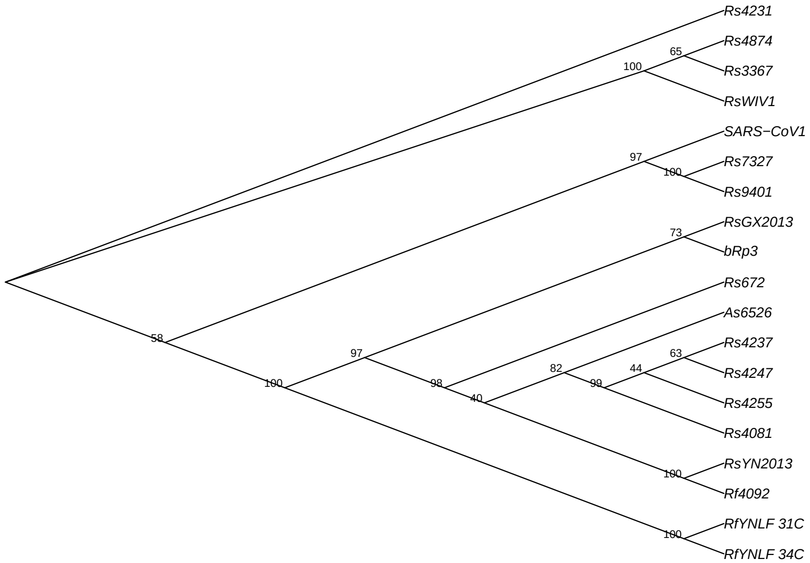

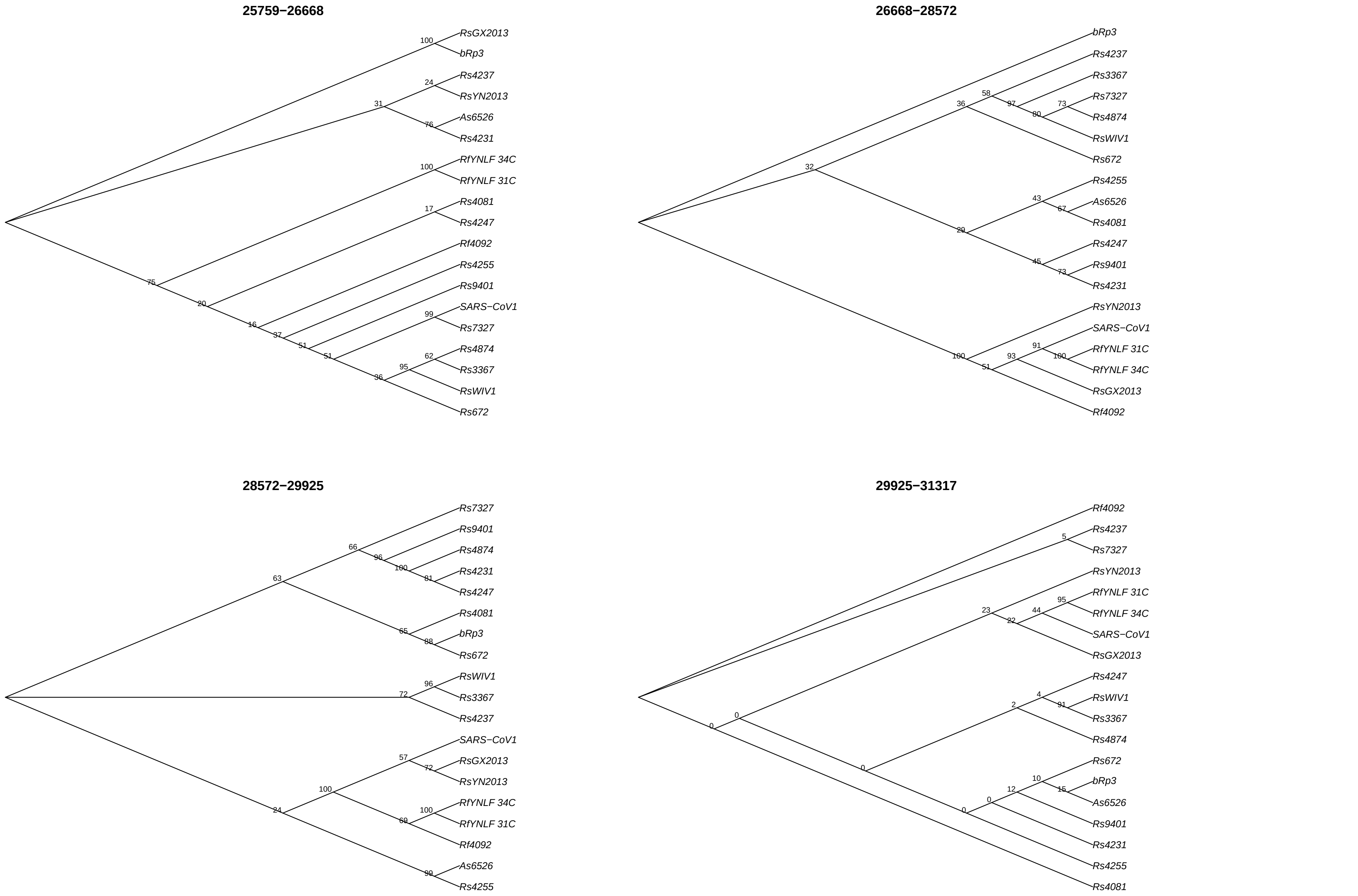
25759−26668
RsGX2013
bRp3
Rs4237
RsYN2013
As6526
Rs4231
RfYNLF 34C
RfYNLF 31C
Rs4081
Rs4247
Rf4092
Rs4255
Rs9401
SARS−CoV1
Rs7327
Rs4874
Rs3367
RsWIV1
Rs672
26668−28572
bRp3
Rs4237
Rs3367
Rs7327
Rs4874
RsWIV1
Rs672
Rs4255
As6526
Rs4081
Rs4247
Rs9401
Rs4231
RsYN2013
SARS−CoV1
RfYNLF 31C
RfYNLF 34C
RsGX2013
Rf4092
28572−29925
Rs7327
Rs9401
Rs4874
Rs4231
Rs4247
Rs4081
bRp3
Rs672
RsWIV1
Rs3367
Rs4237
SARS−CoV1
RsGX2013
RsYN2013
RfYNLF 34C
RfYNLF 31C
Rf4092
As6526
Rs4255
29925−31317
Rf4092
Rs4237
Rs7327
RsYN2013
RfYNLF 31C
RfYNLF 34C
SARS−CoV1
RsGX2013
Rs4247
RsWIV1
Rs3367
Rs4874
Rs672
bRp3
As6526
Rs9401
Rs4231
Rs4255
Rs4081

(B)

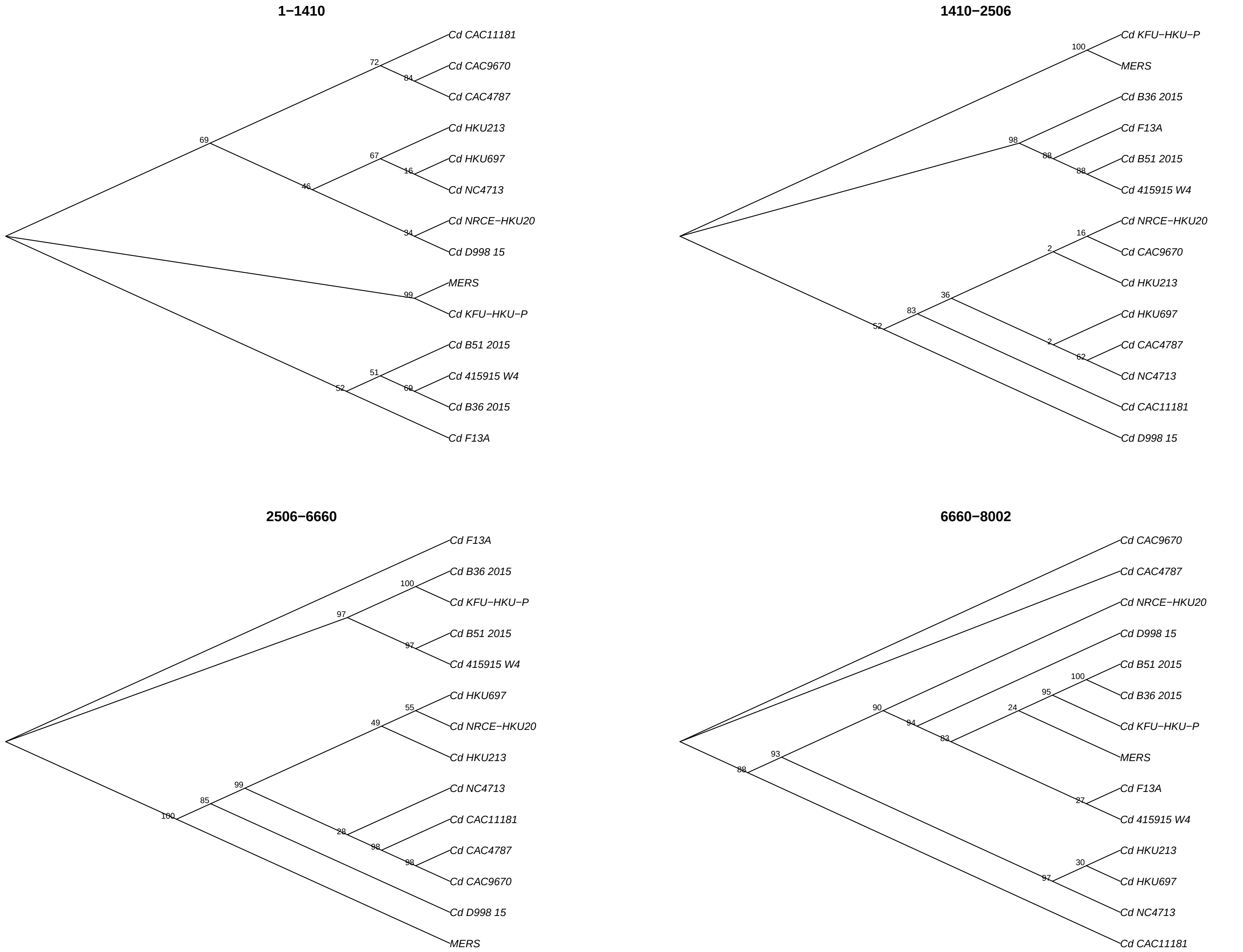
1−1410
Cd CAC11181
Cd CAC9670
Cd CAC4787
Cd HKU213
Cd HKU697
Cd NC4713
Cd NRCE−HKU20
Cd D998 15
MERS
Cd KFU−HKU−P
Cd B51 2015
Cd 415915 W4
Cd B36 2015
Cd F13A
1410−2506
Cd KFU−HKU−P
MERS
Cd B36 2015
Cd F13A
Cd B51 2015
Cd 415915 W4
Cd NRCE−HKU20
Cd CAC9670
Cd HKU213
Cd HKU697
Cd CAC4787
Cd NC4713
Cd CAC11181
Cd D998 15
2506−6660
Cd F13A
Cd B36 2015
Cd KFU−HKU−P
Cd B51 2015
Cd 415915 W4
Cd HKU697
Cd NRCE−HKU20
Cd HKU213
Cd NC4713
Cd CAC11181
Cd CAC4787
Cd CAC9670
Cd D998 15
MERS
6660−8002
Cd CAC9670
Cd CAC4787
Cd NRCE−HKU20
Cd D998 15
Cd B51 2015
Cd B36 2015
Cd KFU−HKU−P
MERS
Cd F13A
Cd 415915 W4
Cd HKU213
Cd HKU697
Cd NC4713
Cd CAC11181

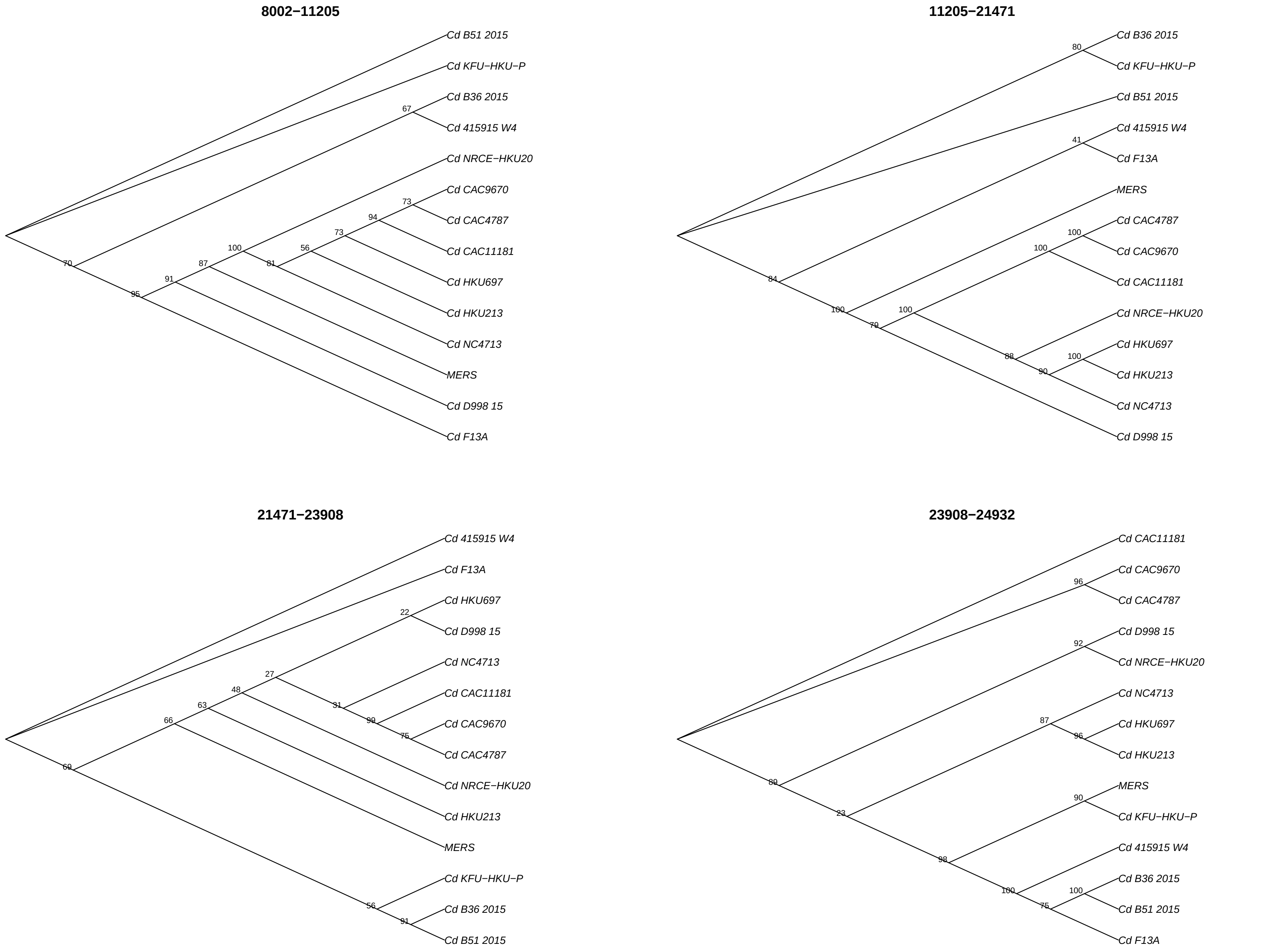
8002−11205
Cd B51 2015
Cd KFU−HKU−P
Cd B36 2015
Cd 415915 W4
Cd NRCE−HKU20
Cd CAC9670
Cd CAC4787
Cd CAC11181
Cd HKU697
Cd HKU213
Cd NC4713
MERS
Cd D998 15
Cd F13A
11205−21471
Cd B36 2015
Cd KFU−HKU−P
Cd B51 2015
Cd 415915 W4
Cd F13A
MERS
Cd CAC4787
Cd CAC9670
Cd CAC11181
Cd NRCE−HKU20
Cd HKU697
Cd HKU213
Cd NC4713
Cd D998 15
21471−23908
Cd 415915 W4
Cd F13A
Cd HKU697
Cd D998 15
Cd NC4713
Cd CAC11181
Cd CAC9670
Cd CAC4787
Cd NRCE−HKU20
Cd HKU213
MERS
Cd KFU−HKU−P
Cd B36 2015
Cd B51 2015
23908−24932
Cd CAC11181
Cd CAC9670
Cd CAC4787
Cd D998 15
Cd NRCE−HKU20
Cd NC4713
Cd HKU697
Cd HKU213
MERS
Cd KFU−HKU−P
Cd 415915 W4
Cd B36 2015
Cd B51 2015
Cd F13A

## 24932−27794

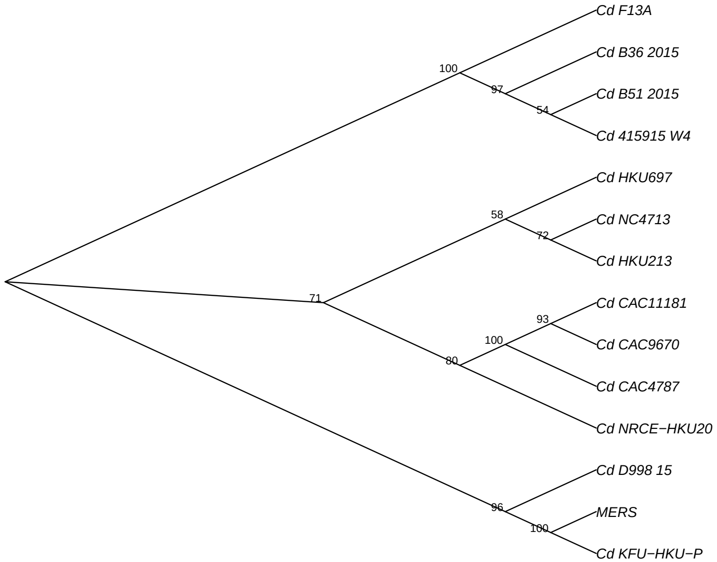

## 27794−28922

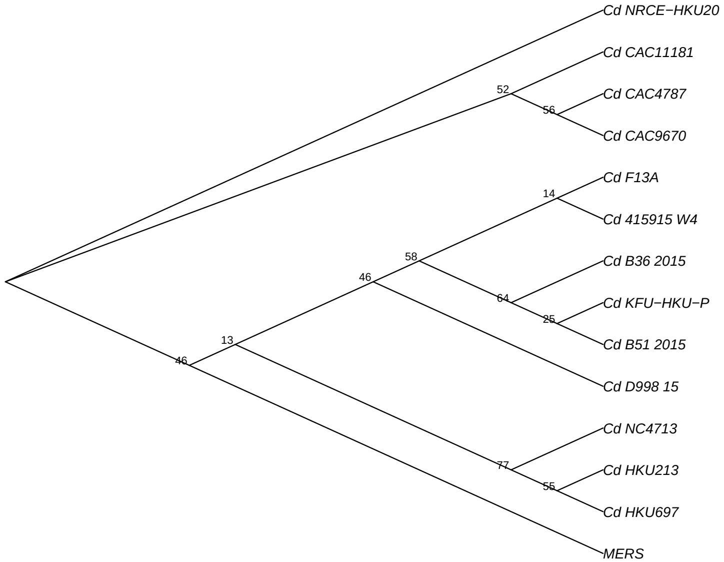

## 28922−30578

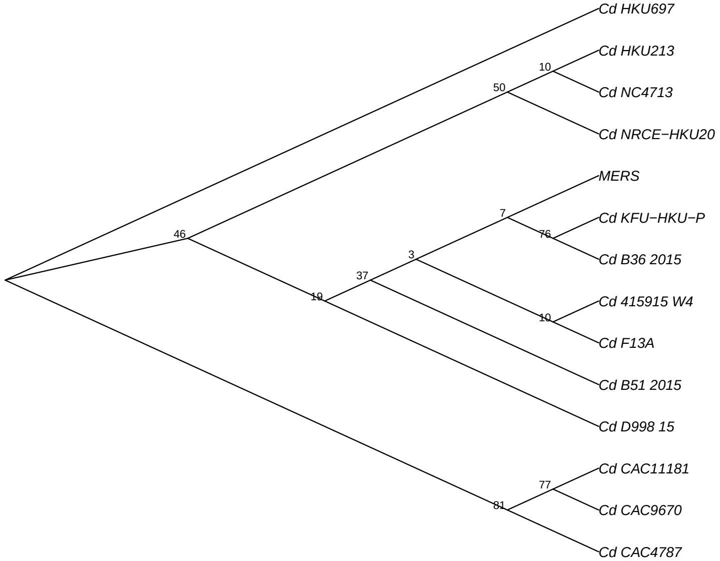

(C)

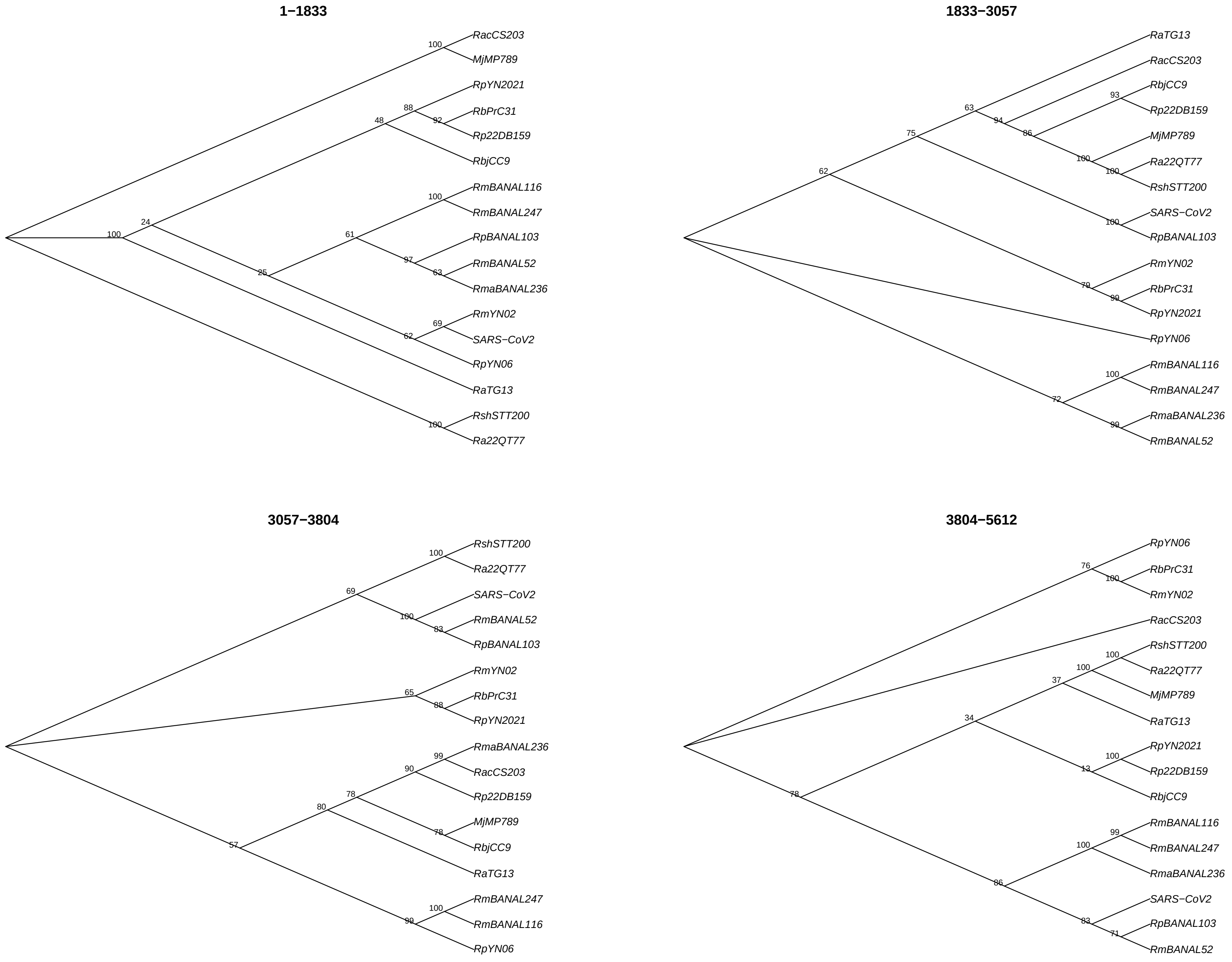
1−1833
RacCS203
MjMP789
RpYN2021
RbPrC31
Rp22DB159
RbjCC9
RmBANAL116
RmBANAL247
RpBANAL103
RmBANAL52
RmaBANAL236
RmYN02
SARS−CoV2
RpYN06
RaTG13
RshSTT200
Ra22QT77
1833−3057
RaTG13
RacCS203
RbjCC9
Rp22DB159
MjMP789
Ra22QT77
RshSTT200
SARS−CoV2
RpBANAL103
RmYN02
RbPrC31
RpYN2021
RpYN06
RmBANAL116
RmBANAL247
RmaBANAL236
RmBANAL52
3057−3804
RshSTT200
Ra22QT77
SARS−CoV2
RmBANAL52
RpBANAL103
RmYN02
RbPrC31
RpYN2021
RmaBANAL236
RacCS203
Rp22DB159
MjMP789
RbjCC9
RaTG13
RmBANAL247
RmBANAL116
RpYN06
3804−5612
RpYN06
RbPrC31
RmYN02
RacCS203
RshSTT200
Ra22QT77
MjMP789
RaTG13
RpYN2021
Rp22DB159
RbjCC9
RmBANAL116
RmBANAL247
RmaBANAL236
SARS−CoV2
RpBANAL103
RmBANAL52

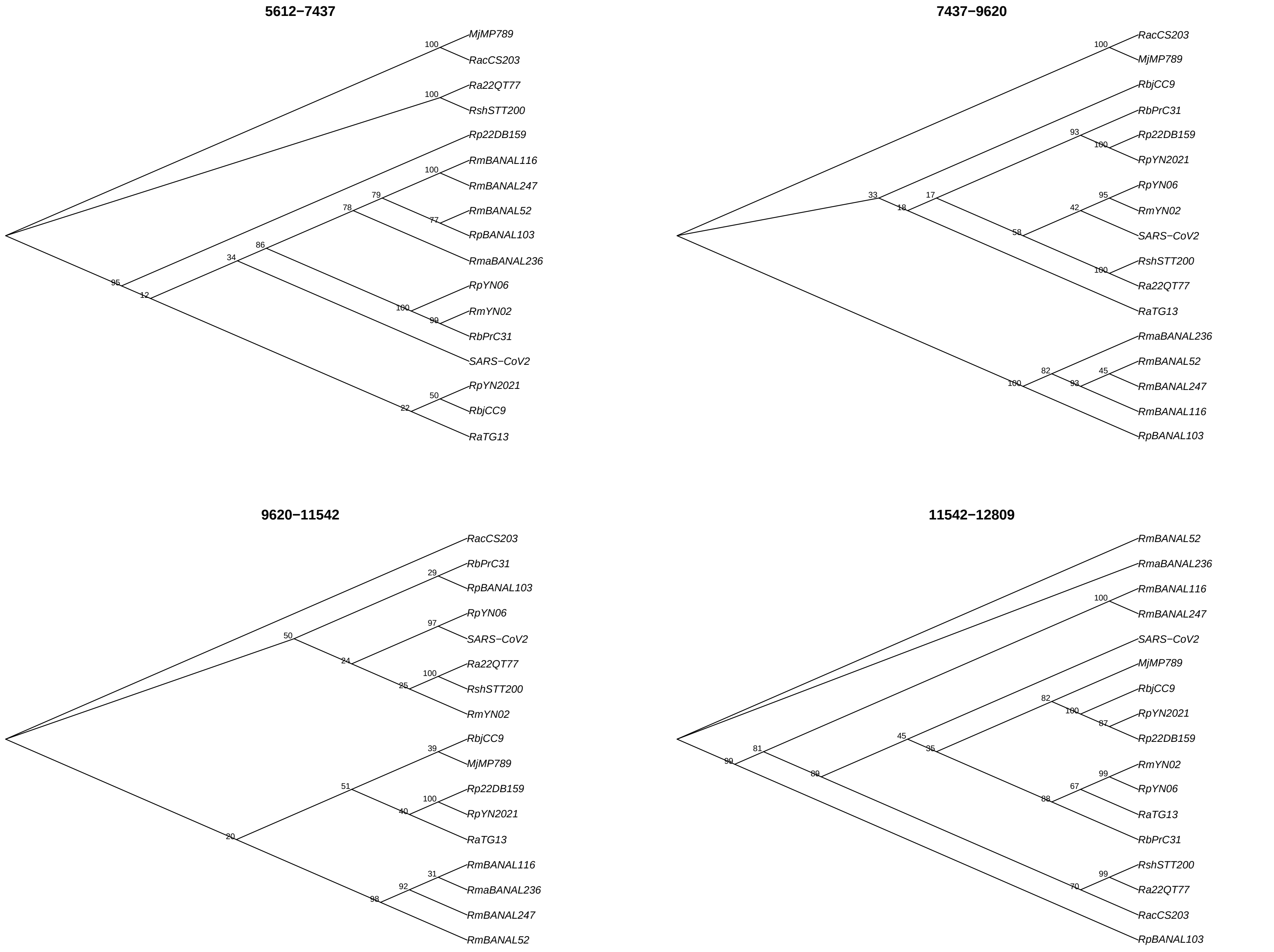
5612−7437
MjMP789
RacCS203
Ra22QT77
RshSTT200
Rp22DB159
RmBANAL116
RmBANAL247
RmBANAL52
RpBANAL103
RmaBANAL236
RpYN06
RmYN02
RbPrC31
SARS−CoV2
RpYN2021
RbjCC9
RaTG13
7437−9620
RacCS203
MjMP789
RbjCC9
RbPrC31
Rp22DB159
RpYN2021
RpYN06
RmYN02
SARS−CoV2
RshSTT200
Ra22QT77
RaTG13
RmaBANAL236
RmBANAL52
RmBANAL247
RmBANAL116
RpBANAL103
9620−11542
RacCS203
RbPrC31
RpBANAL103
RpYN06
SARS−CoV2
Ra22QT77
RshSTT200
RmYN02
RbjCC9
MjMP789
Rp22DB159
RpYN2021
RaTG13
RmBANAL116
RmaBANAL236
RmBANAL247
RmBANAL52
11542−12809
RmBANAL52
RmaBANAL236
RmBANAL116
RmBANAL247
SARS−CoV2
MjMP789
RbjCC9
RpYN2021
Rp22DB159
RmYN02
RpYN06
RaTG13
RbPrC31
RshSTT200
Ra22QT77
RacCS203
RpBANAL103

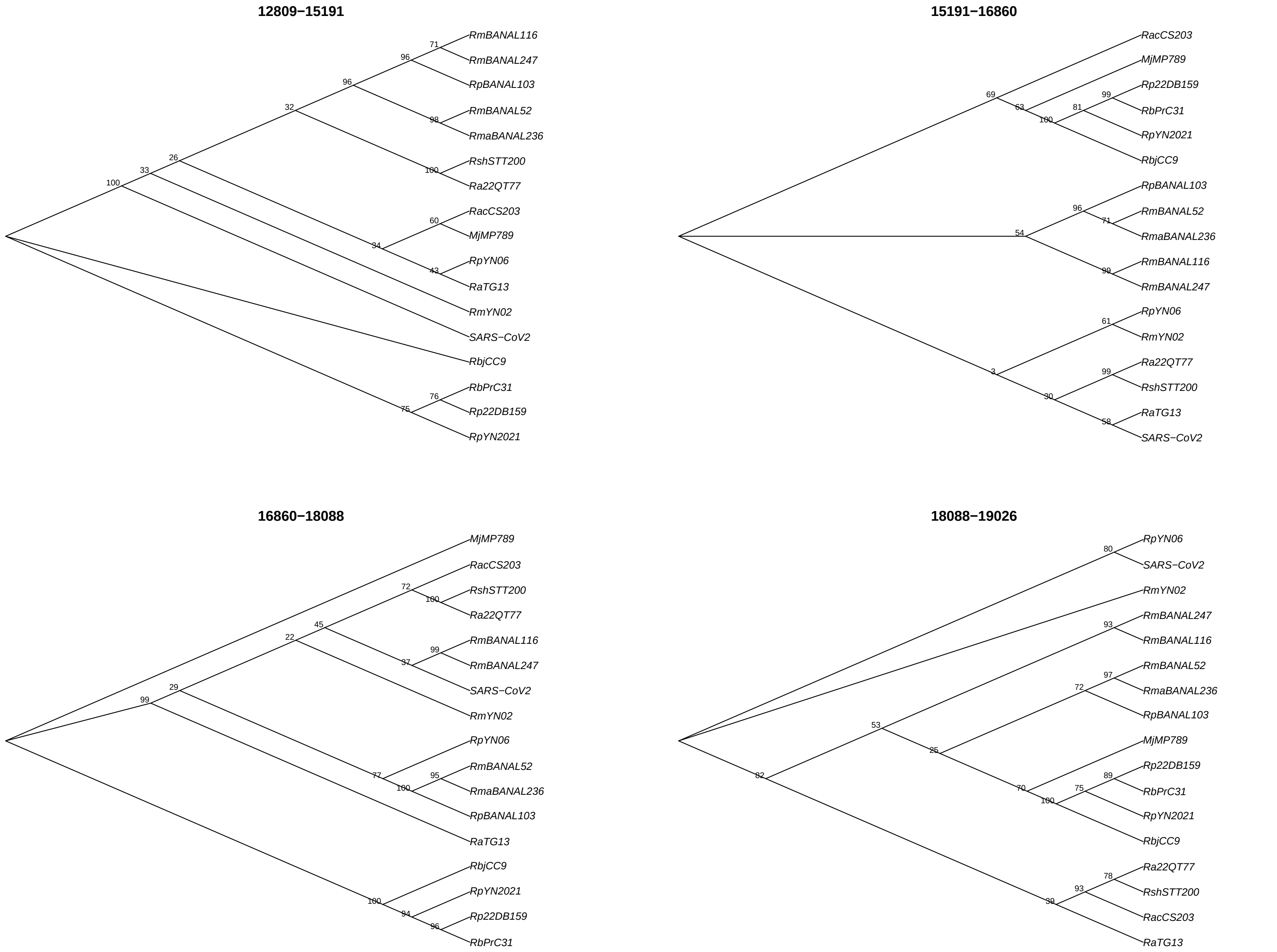
12809−15191
RmBANAL116
RmBANAL247
RpBANAL103
RmBANAL52
RmaBANAL236
RshSTT200
Ra22QT77
RacCS203
MjMP789
RpYN06
RaTG13
RmYN02
SARS−CoV2
RbjCC9
RbPrC31
Rp22DB159
RpYN2021
15191−16860
RacCS203
MjMP789
Rp22DB159
RbPrC31
RpYN2021
RbjCC9
RpBANAL103
RmBANAL52
RmaBANAL236
RmBANAL116
RmBANAL247
RpYN06
RmYN02
Ra22QT77
RshSTT200
RaTG13
SARS−CoV2
16860−18088
MjMP789
RacCS203
RshSTT200
Ra22QT77
RmBANAL116
RmBANAL247
SARS−CoV2
RmYN02
RpYN06
RmBANAL52
RmaBANAL236
RpBANAL103
RaTG13
RbjCC9
RpYN2021
Rp22DB159
RbPrC31
18088−19026
RpYN06
SARS−CoV2
RmYN02
RmBANAL247
RmBANAL116
RmBANAL52
RmaBANAL236
RpBANAL103
MjMP789
Rp22DB159
RbPrC31
RpYN2021
RbjCC9
Ra22QT77
RshSTT200
RacCS203
RaTG13

## 19026−20559

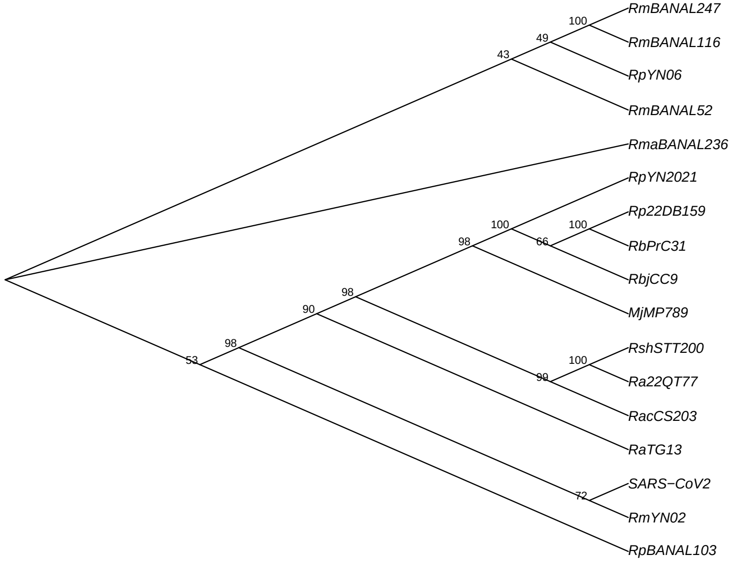

## 20559−20920

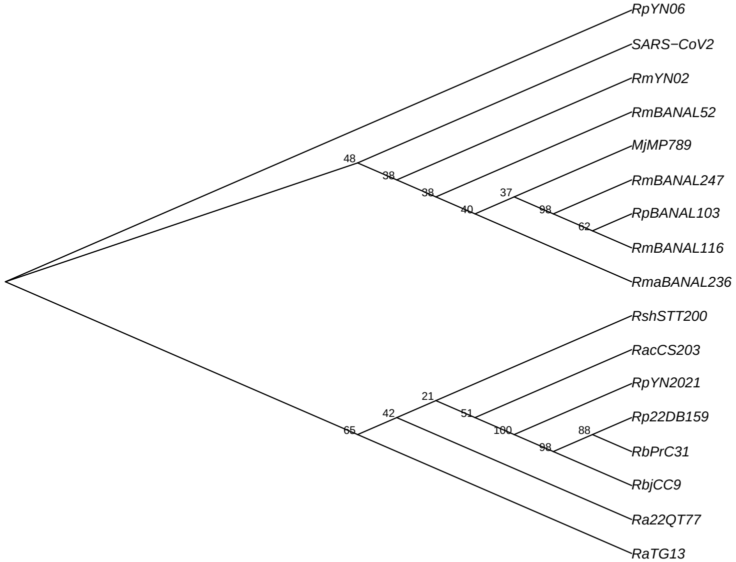

## 20920−21079

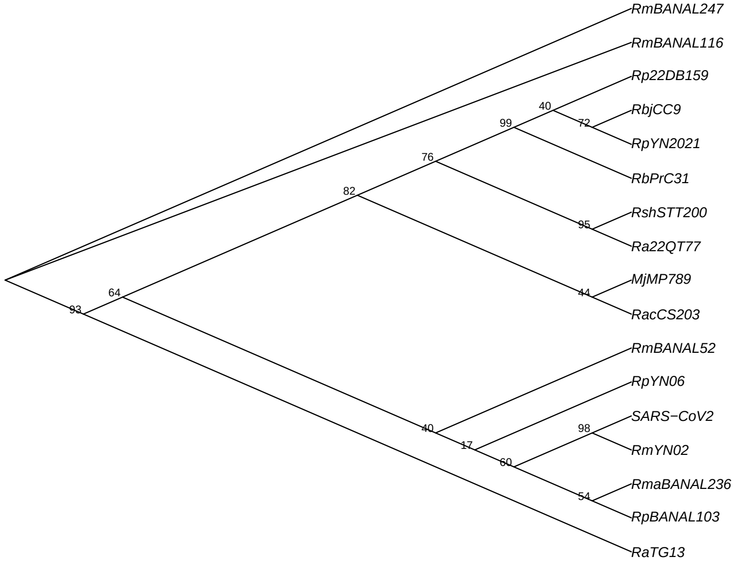

## 21079−21350

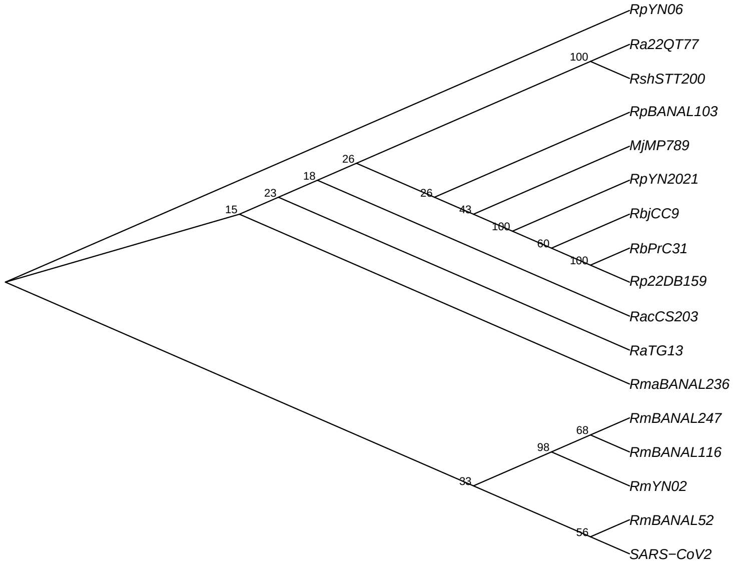

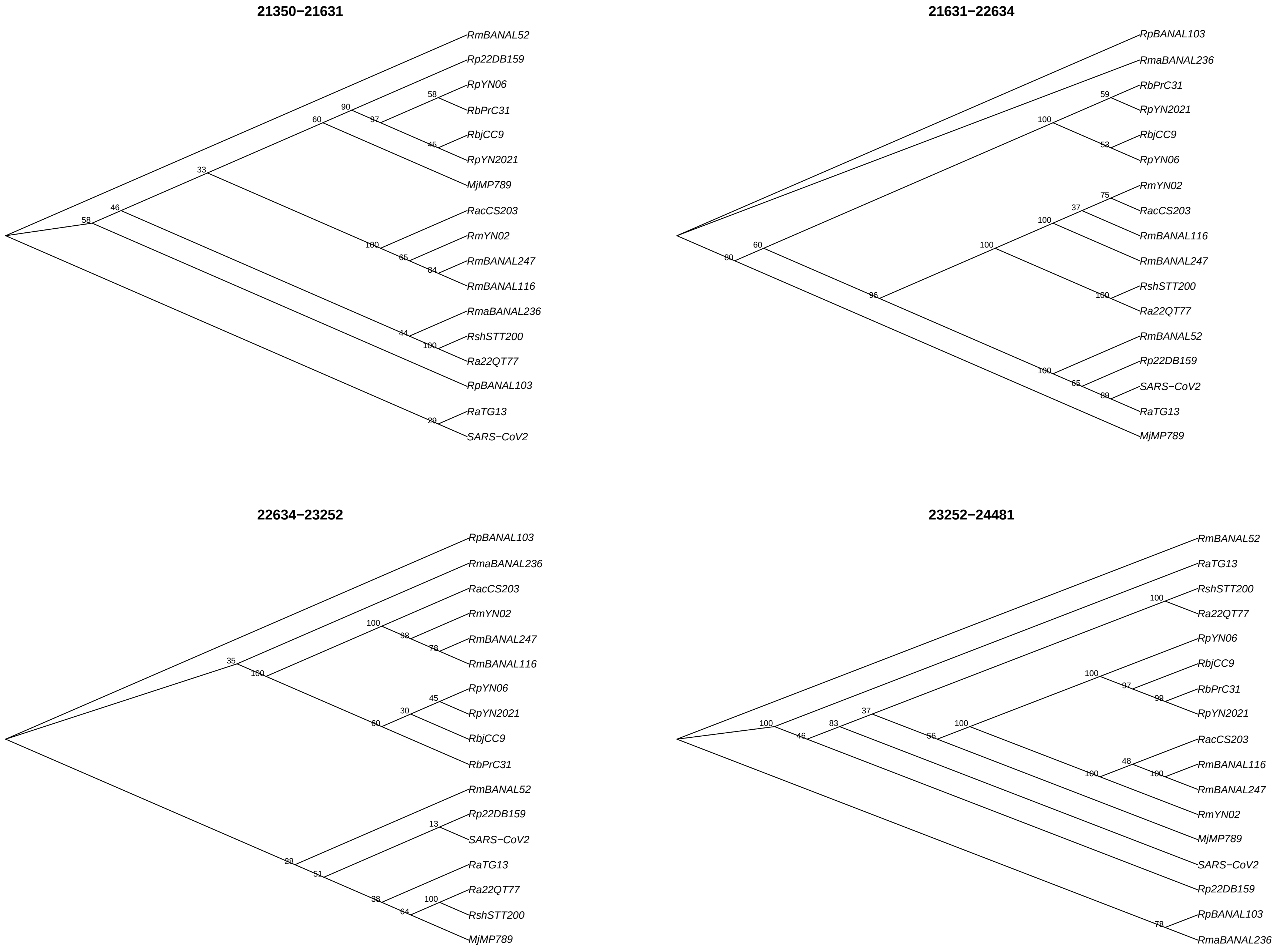
21350−21631
RmBANAL52
Rp22DB159
RpYN06
RbPrC31
RbjCC9
RpYN2021
MjMP789
RacCS203
RmYN02
RmBANAL247
RmBANAL116
RmaBANAL236
RshSTT200
Ra22QT77
RpBANAL103
RaTG13
SARS−CoV2
21631−22634
RpBANAL103
RmaBANAL236
RbPrC31
RpYN2021
RbjCC9
RpYN06
RmYN02
RacCS203
RmBANAL116
RmBANAL247
RshSTT200
Ra22QT77
RmBANAL52
Rp22DB159
SARS−CoV2
RaTG13
MjMP789
22634−23252
RpBANAL103
RmaBANAL236
RacCS203
RmYN02
RmBANAL247
RmBANAL116
RpYN06
RpYN2021
RbjCC9
RbPrC31
RmBANAL52
Rp22DB159
SARS−CoV2
RaTG13
Ra22QT77
RshSTT200
MjMP789
23252−24481
RmBANAL52
RaTG13
RshSTT200
Ra22QT77
RpYN06
RbjCC9
RbPrC31
RpYN2021
RacCS203
RmBANAL116
RmBANAL247
RmYN02
MjMP789
SARS−CoV2
Rp22DB159
RpBANAL103
RmaBANAL236

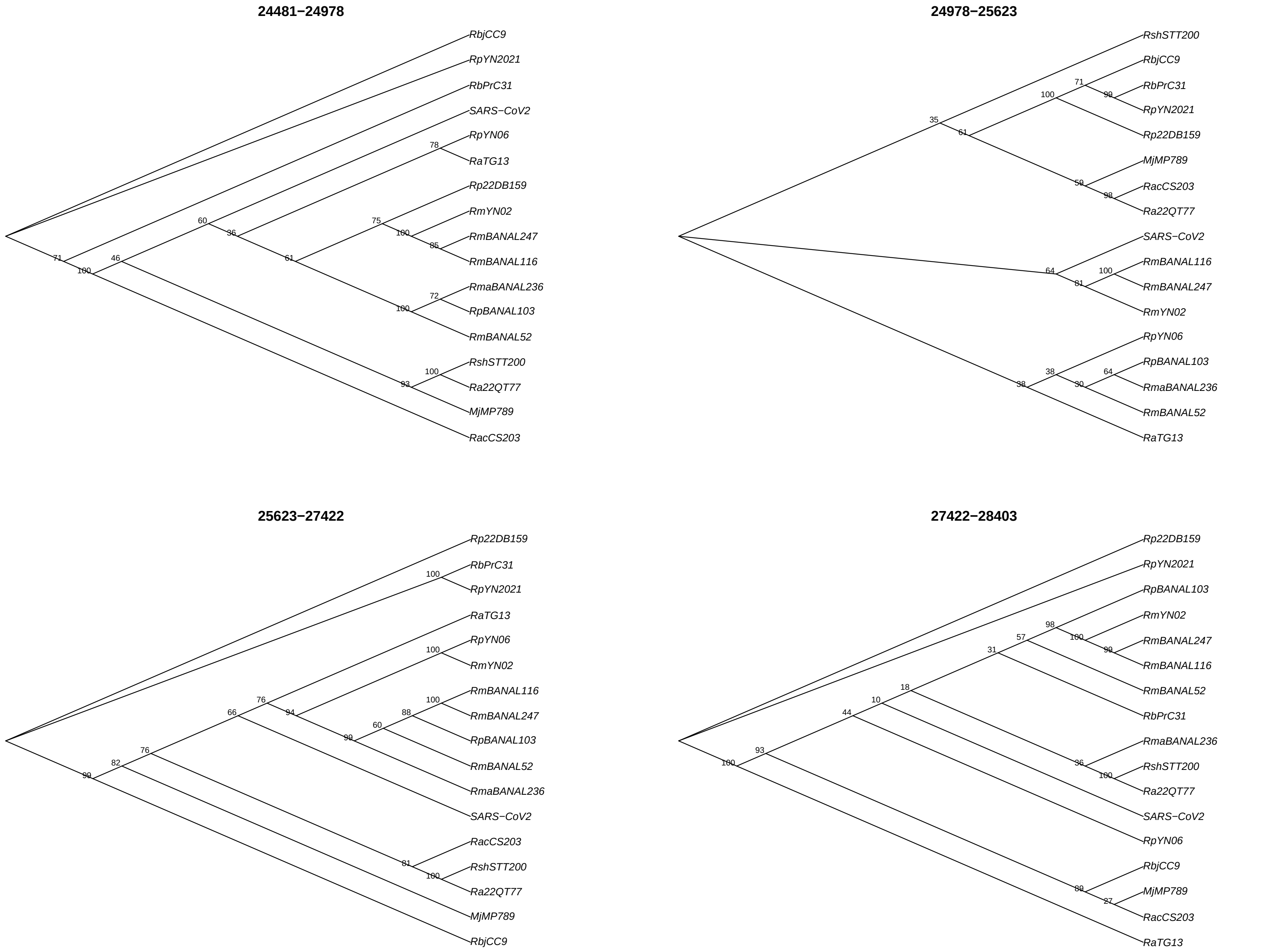
24481−24978
RbjCC9
RpYN2021
RbPrC31
SARS−CoV2
RpYN06
RaTG13
Rp22DB159
RmYN02
RmBANAL247
RmBANAL116
RmaBANAL236
RpBANAL103
RmBANAL52
RshSTT200
Ra22QT77
MjMP789
RacCS203
24978−25623
RshSTT200
RbjCC9
RbPrC31
RpYN2021
Rp22DB159
MjMP789
RacCS203
Ra22QT77
SARS−CoV2
RmBANAL116
RmBANAL247
RmYN02
RpYN06
RpBANAL103
RmaBANAL236
RmBANAL52
RaTG13
25623−27422
Rp22DB159
RbPrC31
RpYN2021
RaTG13
RpYN06
RmYN02
RmBANAL116
RmBANAL247
RpBANAL103
RmBANAL52
RmaBANAL236
SARS−CoV2
RacCS203
RshSTT200
Ra22QT77
MjMP789
RbjCC9
27422−28403
Rp22DB159
RpYN2021
RpBANAL103
RmYN02
RmBANAL247
RmBANAL116
RmBANAL52
RbPrC31
RmaBANAL236
RshSTT200
Ra22QT77
SARS−CoV2
RpYN06
RbjCC9
MjMP789
RacCS203
RaTG13

## 28403−30226

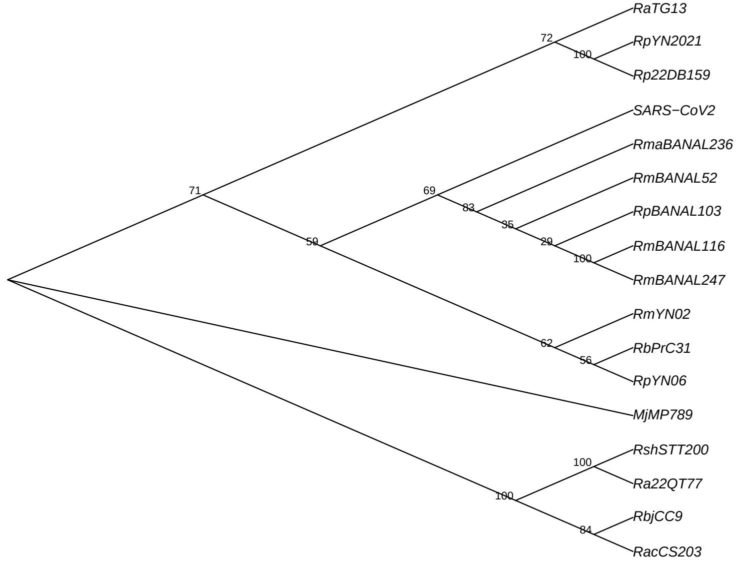
RaTG13
RpYN2021
Rp22DB159
SARS−CoV2
RmaBANAL236
RmBANAL52
RpBANAL103
RmBANAL116
RmBANAL247
RmYN02
RbPrC31
RpYN06
MjMP789
RshSTT200
Ra22QT77
RbjCC9
RacCS203
72
100
71
69
83
35
29
100
59
62
56
100
100
84

Figure S12. Estimation of d$N$ across all the segments of the three datasets. (A) SARS-CoV-1, (B) MERS and (C) SARS-CoV-2, (D) SARS-CoV-2 with only 14 segments (larger segments), (E) CoV-2-S$^{\mathrm{edit}}$, (F) WIV-BANAL-20-236$^{\mathrm{opt}}$, and (G) WIV-BANAL-20-236$^{\mathrm{nat}}$. Virus sequences are listed on the left. The $x$-axis shows consecutive segments and their corresponding nucleotide positions in the multiple sequence alignment. Segment lengths (in number of codons) are indicated in bold in the first row of the table. For each segment and each terminal branch of the phylogeny (i.e., each leaf sequence), the estimated d$N$ values are reported numerically and visualized using a color-coded heatmap.

(A)

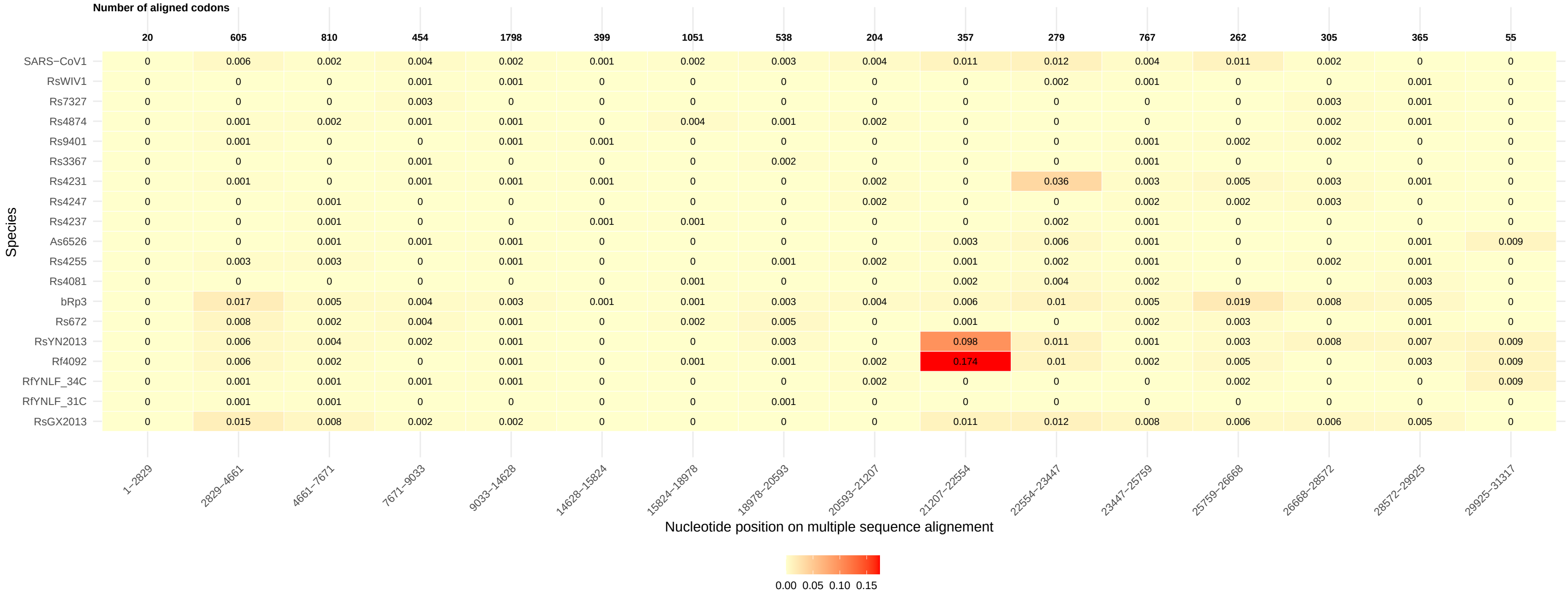
Number of aligned codons
20 605 810 454 1798 399 1051 538 204 357 279 767 262 305 365 55
Species
SARS−CoV1 0 0.006 0.002 0.004 0.002 0.001 0.002 0.003 0.004 0.011 0.012 0.004 0.011 0.002 0 0
RsWIV1 0 0 0 0.001 0.001 0 0 0 0 0 0.002 0.001 0 0 0.001 0
Rs7327 0 0 0 0.003 0 0 0 0 0 0 0 0 0 0.003 0.001 0
Rs4874 0 0.001 0.002 0.001 0.001 0 0.004 0.001 0.002 0 0 0 0 0.002 0.001 0
Rs9401 0 0.001 0 0 0.001 0.001 0 0 0 0 0 0.001 0.002 0.002 0 0
Rs3367 0 0 0 0.001 0 0 0 0.002 0 0 0 0.001 0 0 0 0
Rs4231 0 0.001 0 0.001 0.001 0.001 0 0 0.002 0 0.036 0.003 0.005 0.003 0.001 0
Rs4247 0 0 0.001 0 0 0 0 0 0.002 0 0 0.002 0.002 0.003 0 0
Rs4237 0 0 0.001 0 0 0.001 0.001 0 0 0 0.002 0.001 0 0 0 0
As6526 0 0 0.001 0.001 0.001 0 0 0 0 0.003 0.006 0.001 0 0 0.001 0.009
Rs4255 0 0.003 0.003 0 0.001 0 0 0.001 0.002 0.001 0.002 0.001 0 0.002 0.001 0
Rs4081 0 0 0 0 0 0 0.001 0 0 0.002 0.004 0.002 0 0 0.003 0
bRp3 0 0.017 0.005 0.004 0.003 0.001 0.001 0.003 0.004 0.006 0.01 0.005 0.019 0.008 0.005 0
Rs672 0 0.008 0.002 0.004 0.001 0 0.002 0.005 0 0.001 0 0.002 0.003 0 0.001 0
RsYN2013 0 0.006 0.004 0.002 0.001 0 0 0.003 0 0.098 0.011 0.001 0.003 0.008 0.007 0.009
Rf4092 0 0.006 0.002 0 0.001 0 0.001 0.001 0.002 0.174 0.01 0.002 0.005 0 0.003 0.009
RfYNLF_34C 0 0.001 0.001 0.001 0.001 0 0 0 0.002 0 0 0 0.002 0 0 0.009
RfYNLF_31C 0 0.001 0.001 0 0 0 0 0.001 0 0 0 0 0 0 0 0
RsGX2013 0 0.015 0.008 0.002 0.002 0 0 0 0 0.011 0.012 0.008 0.006 0.006 0.005 0
1−2829 2829−4661 4661−7671 7671−9033 9033−14628 14628−15824 15824−18978 18978−20593 20593−21207 21207−22554 22554−23447 23447−25759 25759−26668 26668−28572 28572−29925 29925−31317
Nucleotide position on multiple sequence alignement
0.00 0.05 0.10 0.15

(B)

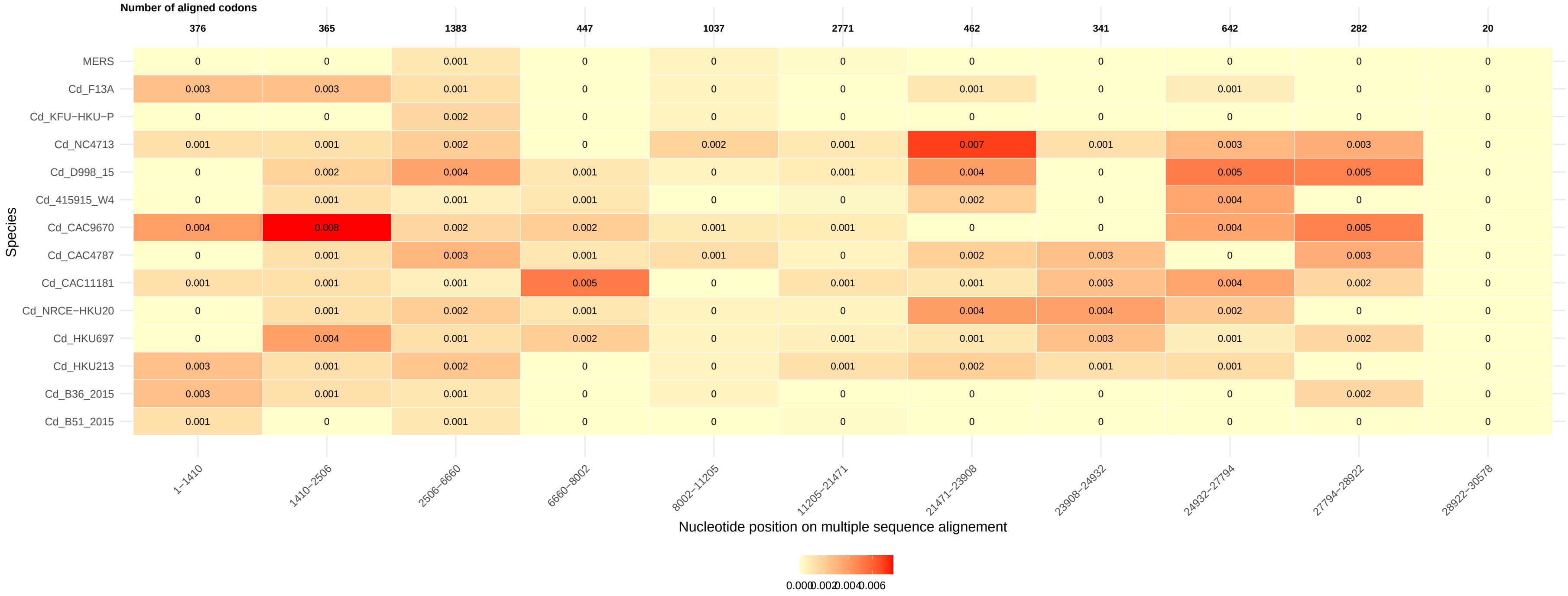

Number of aligned codons
376
365
1383
447
1037
2771
462
341
642
282
20
MERS
Cd_F13A
Cd_KFU−HKU−P
Cd_NC4713
Cd_D998_15
Cd_415915_W4
Cd_CAC9670
Cd_CAC4787
Cd_CAC11181
Cd_NRCE−HKU20
Cd_HKU697
Cd_HKU213
Cd_B36_2015
Cd_B51_2015
Species
1−1410
1410−2506
2506−6660
6660−8002
8002−11205
11205−21471
21471−23908
23908−24932
24932−27794
27794−28922
28922−30578
Nucleotide position on multiple sequence alignement
0.000
0.002
0.004
0.006

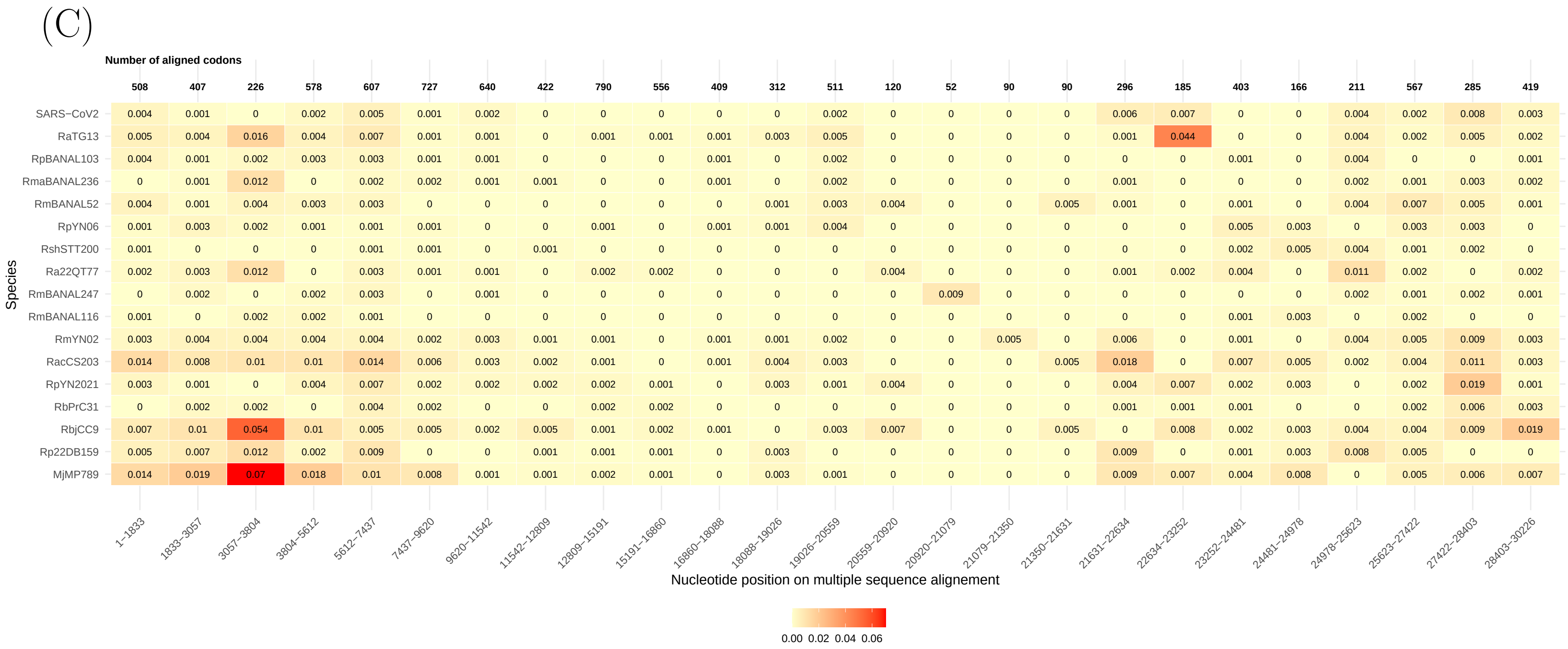
(C)
Number of aligned codons
508 407 226 578 607 727 640 422 790 556 409 312 511 120 52 90 90 296 185 403 166 211 567 285 419
SARS−CoV2 0.004 0.001 0 0.002 0.005 0.001 0.002 0 0 0 0 0 0.002 0 0 0 0 0.006 0.007 0 0 0.004 0.002 0.008 0.003
RaTG13 0.005 0.004 0.016 0.004 0.007 0.001 0.001 0 0.001 0.001 0.001 0.003 0.005 0 0 0 0 0.001 0.044 0 0 0.004 0.002 0.005 0.002
RpBANAL103 0.004 0.001 0.002 0.003 0.003 0.001 0.001 0 0 0 0.001 0 0.002 0 0 0 0 0 0 0.001 0 0.004 0 0 0.001
RmaBANAL236 0 0.001 0.012 0 0.002 0.002 0.001 0.001 0 0 0.001 0 0.002 0 0 0 0 0.001 0 0 0 0.002 0.001 0.003 0.002
RmBANAL52 0.004 0.001 0.004 0.003 0.003 0 0 0 0 0 0 0.001 0.003 0.004 0 0 0.005 0.001 0 0.001 0 0.004 0.007 0.005 0.001
RpYN06 0.001 0.003 0.002 0.001 0.001 0.001 0 0 0.001 0 0.001 0.001 0.004 0 0 0 0 0 0 0.005 0.003 0 0.003 0.003 0
RshSTT200 0.001 0 0 0 0.001 0.001 0 0.001 0 0 0 0 0 0 0 0 0 0 0 0.002 0.005 0.004 0.001 0.002 0
Ra22QT77 0.002 0.003 0.012 0 0.003 0.001 0.001 0 0.002 0.002 0 0 0 0.004 0 0 0 0.001 0.002 0.004 0 0.011 0.002 0 0.002
RmBANAL247 0 0.002 0 0.002 0.003 0 0.001 0 0 0 0 0 0 0 0.009 0 0 0 0 0 0 0.002 0.001 0.002 0.001
RmBANAL116 0.001 0 0.002 0.002 0.001 0 0 0 0 0 0 0 0 0 0 0 0 0 0 0.001 0.003 0 0.002 0 0
RmYN02 0.003 0.004 0.004 0.004 0.004 0.002 0.003 0.001 0.001 0 0.001 0.001 0.002 0 0 0.005 0 0.006 0 0.001 0 0.004 0.005 0.009 0.003
RacCS203 0.014 0.008 0.01 0.01 0.014 0.006 0.003 0.002 0.001 0 0.001 0.004 0.003 0 0 0 0.005 0.018 0 0.007 0.005 0.002 0.004 0.011 0.003
RpYN2021 0.003 0.001 0 0.004 0.007 0.002 0.002 0.002 0.002 0.001 0 0.003 0.001 0.004 0 0 0 0.004 0.007 0.002 0.003 0 0.002 0.019 0.001
RbPrC31 0 0.002 0.002 0 0.004 0.002 0 0 0.002 0.002 0 0 0 0 0 0 0 0.001 0.001 0.001 0 0 0.002 0.006 0.003
RbjCC9 0.007 0.01 0.054 0.01 0.005 0.005 0.002 0.005 0.001 0.002 0.001 0 0.003 0.007 0 0 0.005 0 0.008 0.002 0.003 0.004 0.004 0.009 0.019
Rp22DB159 0.005 0.007 0.012 0.002 0.009 0 0 0.001 0.001 0.001 0 0.003 0 0 0 0 0 0.009 0 0.001 0.003 0.008 0.005 0 0
MjMP789 0.014 0.019 0.07 0.018 0.01 0.008 0.001 0.001 0.002 0.001 0 0.003 0.001 0 0 0 0 0.009 0.007 0.004 0.008 0 0.005 0.006 0.007
Species
1−1833 1833−3057 3057−3804 3804−5612 5612−7437 7437−9620 9620−11542 11542−12809 12809−15191 15191−16860 16860−18088 18088−19026 19026−20559 20559−20920 20920−21079 21079−21350 21350−21631 21631−22634 22634−23252 23252−24481 24481−24978 24978−25623 25623−27422 27422−28403 28403−30226
Nucleotide position on multiple sequence alignement
0.00 0.02 0.04 0.06

(D)

Number of aligned codons

| Species | 1–1846 | 1846–3116 | 3116–3807 | 3807–5867 | 5867–7225 | 7225–9614 | 9614–11386 | 11386–12828 | 12828–18449 | 18449–21351 | 21351–22633 | 22633–24995 | 24995–27393 | 27393–30226 |
|---|---|---|---|---|---|---|---|---|---|---|---|---|---|---|
| Number of aligned codons | 513 | 423 | 208 | 662 | 452 | 796 | 590 | 480 | 1869 | 967 | 386 | 761 | 762 | 714 |
| SARS−CoV2 | 0.004 | 0.001 | 0 | 0.004 | 0.005 | 0.001 | 0.002 | 0 | 0 | 0.001 | 0.005 | 0.002 | 0.003 | 0.005 |
| RaTG13 | 0.005 | 0.004 | 0.018 | 0.004 | 0.01 | 0.001 | 0.002 | 0 | 0.001 | 0.003 | 0.001 | 0.01 | 0.002 | 0.003 |
| RpBANAL103 | 0.004 | 0.001 | 0.003 | 0.003 | 0.002 | 0.002 | 0.002 | 0 | 0 | 0.001 | 0 | 0.001 | 0.001 | 0.001 |
| RmaBANAL236 | 0 | 0.001 | 0.013 | 0 | 0.001 | 0.002 | 0.001 | 0.001 | 0 | 0.001 | 0.001 | 0.001 | 0.002 | 0.003 |
| RmBANAL52 | 0.004 | 0.002 | 0.004 | 0.003 | 0.003 | 0 | 0 | 0 | 0 | 0.003 | 0.002 | 0.001 | 0.006 | 0.003 |
| RpYN06 | 0.002 | 0.003 | 0.002 | 0.001 | 0.001 | 0.001 | 0 | 0 | 0.001 | 0.002 | 0 | 0.003 | 0.002 | 0.001 |
| RshSTT200 | 0.001 | 0 | 0 | 0.001 | 0 | 0.001 | 0 | 0.001 | 0 | 0 | 0 | 0.002 | 0.002 | 0.001 |
| Ra22QT77 | 0.002 | 0.003 | 0.013 | 0 | 0.003 | 0.002 | 0.002 | 0 | 0.001 | 0 | 0.001 | 0.003 | 0.004 | 0.001 |
| RmBANAL247 | 0 | 0.002 | 0 | 0.001 | 0.002 | 0.001 | 0.001 | 0 | 0 | 0 | 0 | 0 | 0.001 | 0.001 |
| RmBANAL116 | 0.001 | 0.001 | 0 | 0.002 | 0.001 | 0 | 0 | 0 | 0 | 0 | 0 | 0.001 | 0.002 | 0 |
| RmYN02 | 0.003 | 0.004 | 0.004 | 0.003 | 0.005 | 0.002 | 0.004 | 0.001 | 0.001 | 0.001 | 0.004 | 0.001 | 0.005 | 0.006 |
| RacCS203 | 0.014 | 0.009 | 0.011 | 0.011 | 0.014 | 0.007 | 0.003 | 0.002 | 0.001 | 0.002 | 0.016 | 0.002 | 0.003 | 0.007 |
| RpYN2021 | 0.004 | 0.001 | 0 | 0.003 | 0.009 | 0.002 | 0.002 | 0.002 | 0.001 | 0.001 | 0.003 | 0.004 | 0.001 | 0.008 |
| RbPrC31 | 0 | 0.002 | 0.002 | 0 | 0.006 | 0.002 | 0 | 0 | 0.001 | 0 | 0.001 | 0.002 | 0.001 | 0.004 |
| RbjCC9 | 0.009 | 0.01 | 0.062 | 0.012 | 0.003 | 0.005 | 0.003 | 0.005 | 0.001 | 0.002 | 0.001 | 0.003 | 0.004 | 0.014 |
| Rp22DB159 | 0.005 | 0.007 | 0.014 | 0.002 | 0.011 | 0 | 0 | 0.001 | 0.001 | 0 | 0.006 | 0.002 | 0.005 | 0 |
| MjMP789 | 0.013 | 0.023 | 0.067 | 0.018 | 0.011 | 0.008 | 0.002 | 0.001 | 0.001 | 0.001 | 0.006 | 0.007 | 0.004 | 0.008 |

Species

Nucleotide position on multiple sequence alignement

0.00 0.02 0.04 0.06

(E)

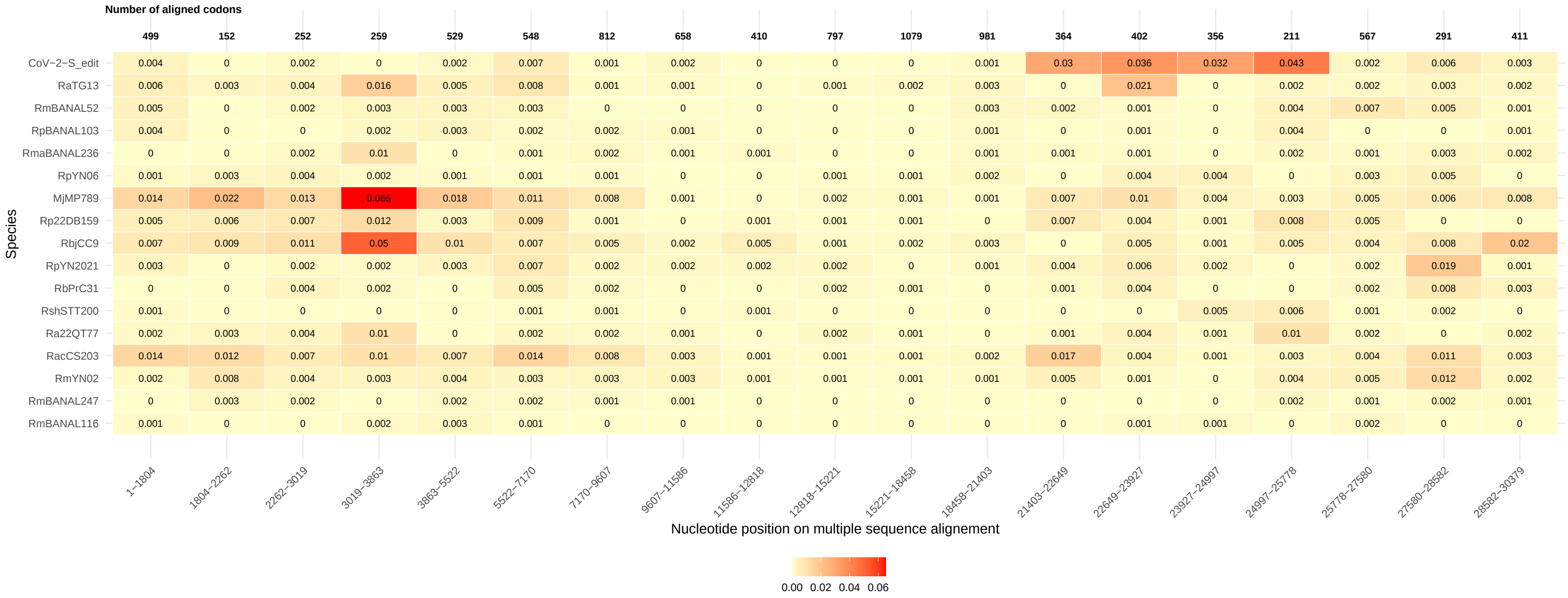
Number of aligned codons
499 152 252 259 529 548 812 658 410 797 1079 981 364 402 356 211 567 291 411
CoV−2−S_edit 0.004 0 0.002 0 0.002 0.007 0.001 0.002 0 0 0 0.001 0.03 0.036 0.032 0.043 0.002 0.006 0.003
RaTG13 0.006 0.003 0.004 0.016 0.005 0.008 0.001 0.001 0 0.001 0.002 0.003 0 0.021 0 0.002 0.002 0.003 0.002
RmBANAL52 0.005 0 0.002 0.003 0.003 0.003 0 0 0 0 0 0.003 0.002 0.001 0 0.004 0.007 0.005 0.001
RpBANAL103 0.004 0 0 0.002 0.003 0.002 0.002 0.001 0 0 0 0.001 0 0.001 0 0.004 0 0 0.001
RmaBANAL236 0 0 0.002 0.01 0 0.001 0.002 0.001 0.001 0 0 0.001 0.001 0.001 0 0.002 0.001 0.003 0.002
RpYN06 0.001 0.003 0.004 0.002 0.001 0.001 0.001 0 0 0.001 0.001 0.002 0 0.004 0.004 0 0.003 0.005 0
MjMP789 0.014 0.022 0.013 0.065 0.018 0.011 0.008 0.001 0 0.002 0.001 0.001 0.007 0.01 0.004 0.003 0.005 0.006 0.008
Rp22DB159 0.005 0.006 0.007 0.012 0.003 0.009 0.001 0 0.001 0.001 0.001 0 0.007 0.004 0.001 0.008 0.005 0 0
RbjCC9 0.007 0.009 0.011 0.05 0.01 0.007 0.005 0.002 0.005 0.001 0.002 0.003 0 0.005 0.001 0.005 0.004 0.008 0.02
RpYN2021 0.003 0 0.002 0.002 0.003 0.007 0.002 0.002 0.002 0.002 0 0.001 0.004 0.006 0.002 0 0.002 0.019 0.001
RbPrC31 0 0 0.004 0.002 0 0.005 0.002 0 0 0.002 0.001 0 0.001 0.004 0 0 0.002 0.008 0.003
RshSTT200 0.001 0 0 0 0 0.001 0.001 0 0.001 0 0 0 0 0 0.005 0.006 0.001 0.002 0
Ra22QT77 0.002 0.003 0.004 0.01 0 0.002 0.002 0.001 0 0.002 0.001 0 0.001 0.004 0.001 0.01 0.002 0 0.002
RacCS203 0.014 0.012 0.007 0.01 0.007 0.014 0.008 0.003 0.001 0.001 0.001 0.002 0.017 0.004 0.001 0.003 0.004 0.011 0.003
RmYN02 0.002 0.008 0.004 0.003 0.004 0.003 0.003 0.003 0.001 0.001 0.001 0.001 0.005 0.001 0 0.004 0.005 0.012 0.002
RmBANAL247 0 0.003 0.002 0 0.002 0.002 0.001 0.001 0 0 0 0 0 0 0 0.002 0.001 0.002 0.001
RmBANAL116 0.001 0 0 0.002 0.003 0.001 0 0 0 0 0 0 0 0.001 0.001 0 0.002 0 0
Species
1−1804
1804−2262
2262−3019
3019−3863
3863−5522
5522−7170
7170−9607
9607−11586
11586−12818
12818−15221
15221−18458
18458−21403
21403−22649
22649−23927
23927−24997
24997−25778
25778−27580
27580−28582
28582−30379
Nucleotide position on multiple sequence alignement
0.00 0.02 0.04 0.06

(F)

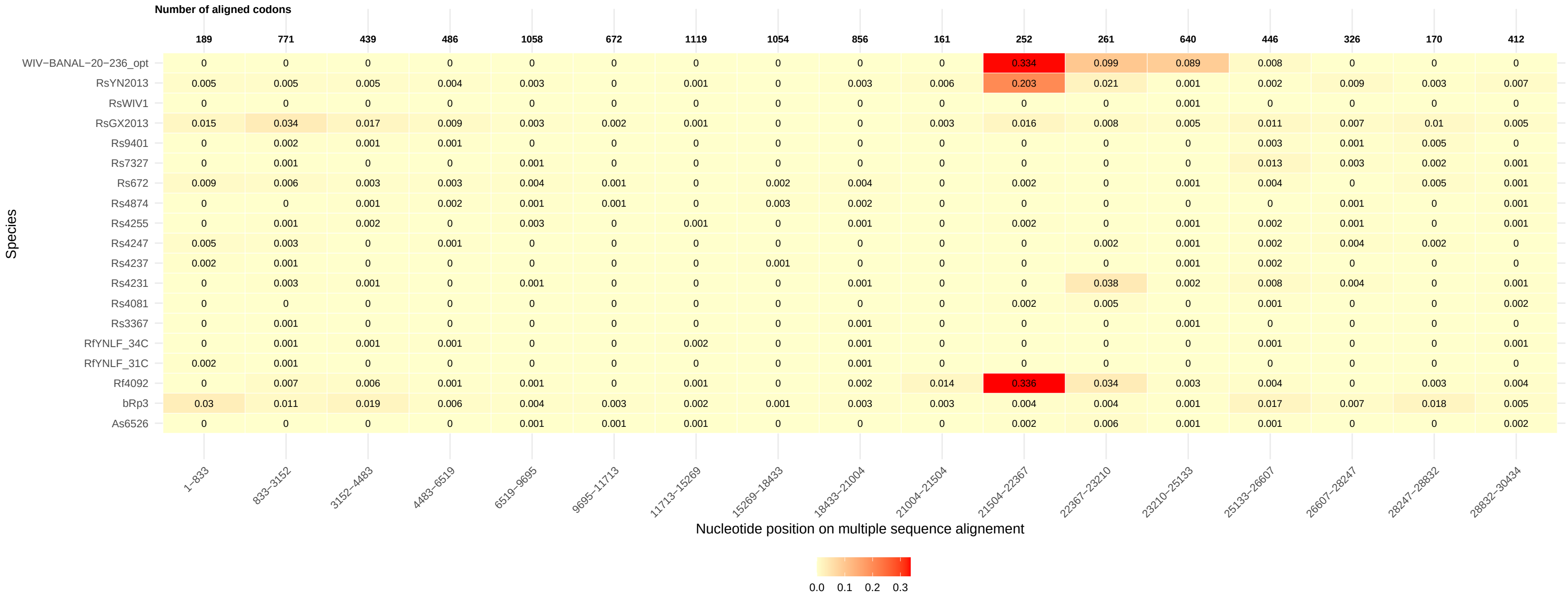
Number of aligned codons
189 771 439 486 1058 672 1119 1054 856 161 252 261 640 446 326 170 412
WIV−BANAL−20−236_opt 0 0 0 0 0 0 0 0 0 0 0.334 0.099 0.089 0.008 0 0 0
RsYN2013 0.005 0.005 0.005 0.004 0.003 0 0.001 0 0.003 0.006 0.203 0.021 0.001 0.002 0.009 0.003 0.007
RsWIV1 0 0 0 0 0 0 0 0 0 0 0 0 0.001 0 0 0 0
RsGX2013 0.015 0.034 0.017 0.009 0.003 0.002 0.001 0 0 0.003 0.016 0.008 0.005 0.011 0.007 0.01 0.005
Rs9401 0 0.002 0.001 0.001 0 0 0 0 0 0 0 0 0 0.003 0.001 0.005 0
Rs7327 0 0.001 0 0 0.001 0 0 0 0 0 0 0 0 0.013 0.003 0.002 0.001
Rs672 0.009 0.006 0.003 0.003 0.004 0.001 0 0.002 0.004 0 0.002 0 0.001 0.004 0 0.005 0.001
Rs4874 0 0 0.001 0.002 0.001 0.001 0 0.003 0.002 0 0 0 0 0 0.001 0 0.001
Rs4255 0 0.001 0.002 0 0.003 0 0.001 0 0.001 0 0.002 0 0.001 0.002 0.001 0 0.001
Rs4247 0.005 0.003 0 0.001 0 0 0 0 0 0 0 0.002 0.001 0.002 0.004 0.002 0
Rs4237 0.002 0.001 0 0 0 0 0 0.001 0 0 0 0 0.001 0.002 0 0 0
Rs4231 0 0.003 0.001 0 0.001 0 0 0 0.001 0 0 0.038 0.002 0.008 0.004 0 0.001
Rs4081 0 0 0 0 0 0 0 0 0 0 0.002 0.005 0 0.001 0 0 0.002
Rs3367 0 0.001 0 0 0 0 0 0 0.001 0 0 0 0.001 0 0 0 0
RfYNLF_34C 0 0.001 0.001 0.001 0 0 0.002 0 0.001 0 0 0 0 0.001 0 0 0.001
RfYNLF_31C 0.002 0.001 0 0 0 0 0 0 0.001 0 0 0 0 0 0 0 0
Rf4092 0 0.007 0.006 0.001 0.001 0 0.001 0 0.002 0.014 0.336 0.034 0.003 0.004 0 0.003 0.004
bRp3 0.03 0.011 0.019 0.006 0.004 0.003 0.002 0.001 0.003 0.003 0.004 0.004 0.001 0.017 0.007 0.018 0.005
As6526 0 0 0 0 0.001 0.001 0.001 0 0 0 0.002 0.006 0.001 0.001 0 0 0.002
Species
1−833 833−3152 3152−4483 4483−6519 6519−9695 9695−11713 11713−15269 15269−18433 18433−21004 21004−21504 21504−22367 22367−23210 23210−25133 25133−26607 26607−28247 28247−28832 28832−30434
Nucleotide position on multiple sequence alignement
0.0 0.1 0.2 0.3

(G)

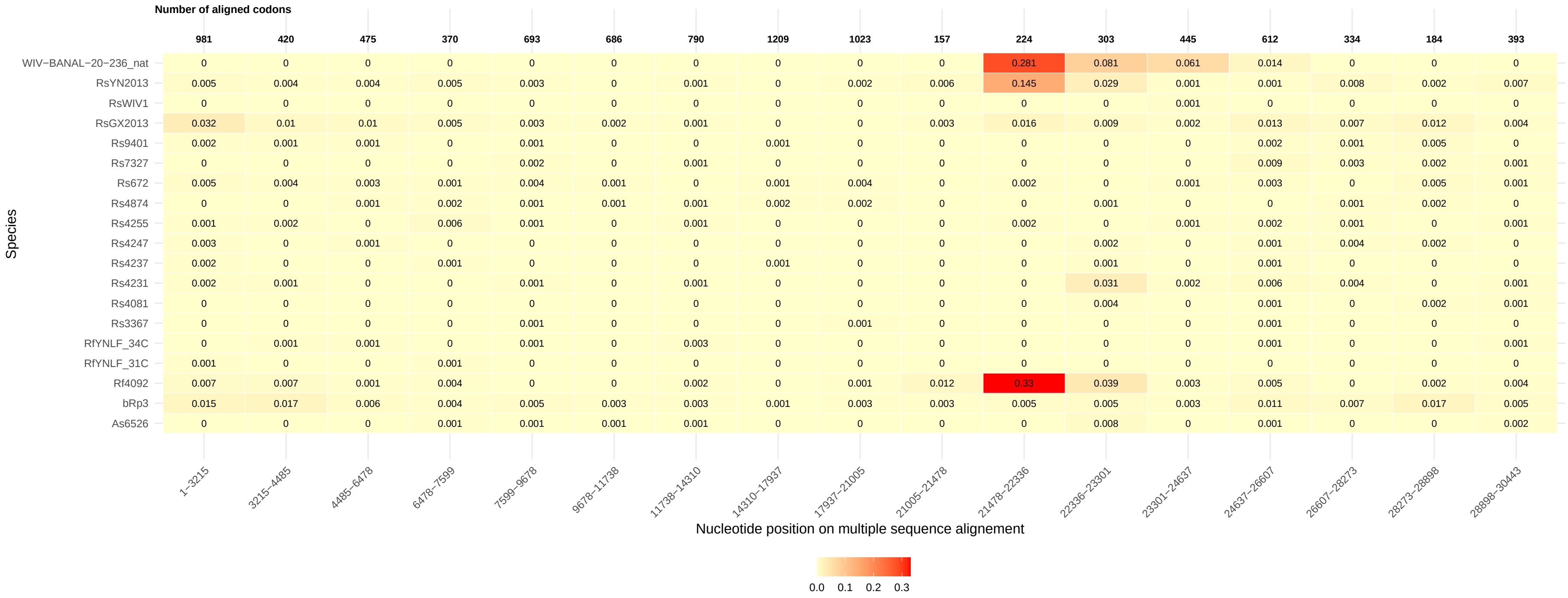

Number of aligned codons
981 420 475 370 693 686 790 1209 1023 157 224 303 445 612 334 184 393
Species
WIV−BANAL−20−236_nat 0 0 0 0 0 0 0 0 0 0 0.281 0.081 0.061 0.014 0 0 0
RsYN2013 0.005 0.004 0.004 0.005 0.003 0 0.001 0 0.002 0.006 0.145 0.029 0.001 0.001 0.008 0.002 0.007
RsWIV1 0 0 0 0 0 0 0 0 0 0 0 0 0.001 0 0 0 0
RsGX2013 0.032 0.01 0.01 0.005 0.003 0.002 0.001 0 0 0.003 0.016 0.009 0.002 0.013 0.007 0.012 0.004
Rs9401 0.002 0.001 0.001 0 0.001 0 0 0.001 0 0 0 0 0 0.002 0.001 0.005 0
Rs7327 0 0 0 0 0.002 0 0.001 0 0 0 0 0 0 0.009 0.003 0.002 0.001
Rs672 0.005 0.004 0.003 0.001 0.004 0.001 0 0.001 0.004 0 0.002 0 0.001 0.003 0 0.005 0.001
Rs4874 0 0 0.001 0.002 0.001 0.001 0.001 0.002 0.002 0 0 0.001 0 0 0.001 0.002 0
Rs4255 0.001 0.002 0 0.006 0.001 0 0.001 0 0 0 0.002 0 0.001 0.002 0.001 0 0.001
Rs4247 0.003 0 0.001 0 0 0 0 0 0 0 0 0.002 0 0.001 0.004 0.002 0
Rs4237 0.002 0 0 0.001 0 0 0 0.001 0 0 0 0.001 0 0.001 0 0 0
Rs4231 0.002 0.001 0 0 0.001 0 0.001 0 0 0 0 0.031 0.002 0.006 0.004 0 0.001
Rs4081 0 0 0 0 0 0 0 0 0 0 0 0.004 0 0.001 0 0.002 0.001
Rs3367 0 0 0 0 0.001 0 0 0 0.001 0 0 0 0 0.001 0 0 0
RfYNLF_34C 0 0.001 0.001 0 0.001 0 0.003 0 0 0 0 0 0 0.001 0 0 0.001
RfYNLF_31C 0.001 0 0 0.001 0 0 0 0 0 0 0 0 0 0 0 0 0
Rf4092 0.007 0.007 0.001 0.004 0 0 0.002 0 0.001 0.012 0.33 0.039 0.003 0.005 0 0.002 0.004
bRp3 0.015 0.017 0.006 0.004 0.005 0.003 0.003 0.001 0.003 0.003 0.005 0.005 0.003 0.011 0.007 0.017 0.005
As6526 0 0 0 0.001 0.001 0.001 0.001 0 0 0 0 0.008 0 0.001 0 0 0.002
1−3215 3215−4485 4485−6478 6478−7599 7599−9678 9678−11738 11738−14310 14310−17937 17937−21005 21005−21478 21478−22336 22336−23301 23301−24637 24637−26607 26607−28273 28273−28898 28898−30443
Nucleotide position on multiple sequence alignement
0.0 0.1 0.2 0.3

Figure S13. Estimation of d$S$ across all the segments of the three datasets. (A) SARS-CoV-1, (B) MERS and (C) SARS-CoV-2, (D) SARS-CoV-2 with only 14 segments (larger segments), (E) CoV-2-S$^{\text{edit}}$, (F) WIV-BANAL-20-236$^{\text{opt}}$, and (G) WIV-BANAL-20-236$^{\text{nat}}$. Same legend as Fig. S12.

(A)

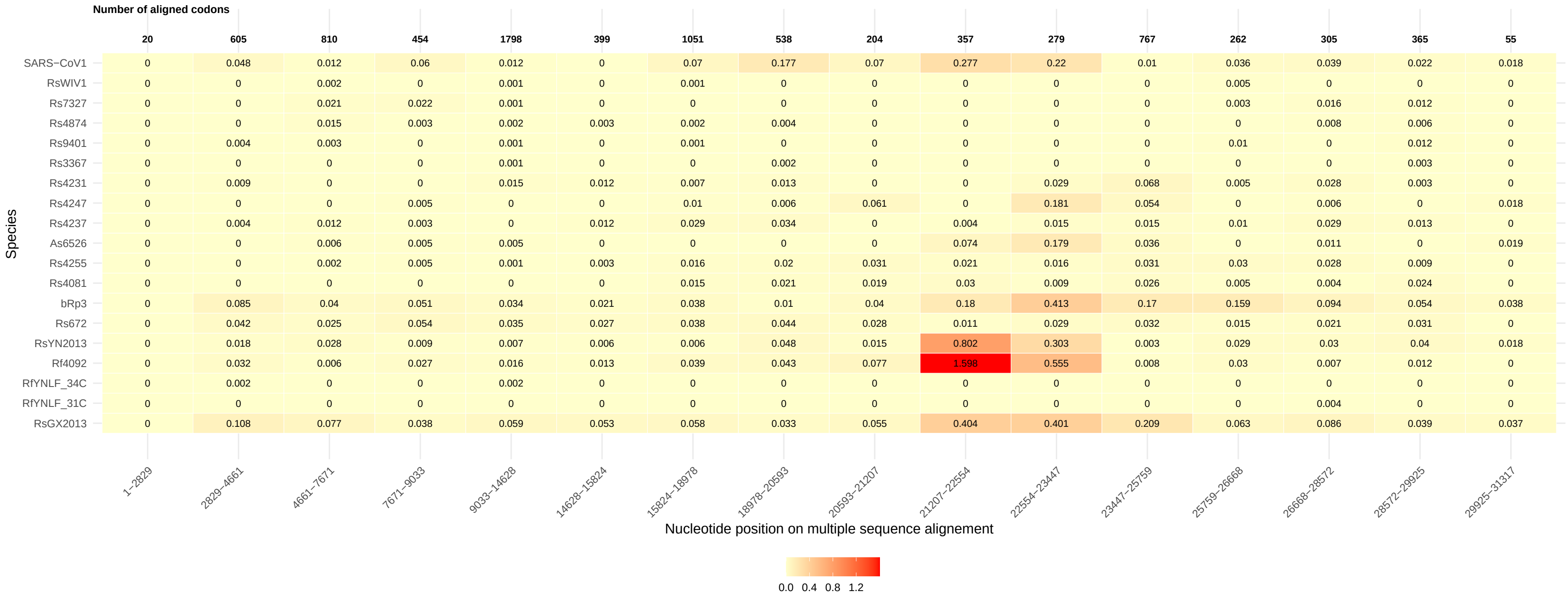
Number of aligned codons
20 605 810 454 1798 399 1051 538 204 357 279 767 262 305 365 55
Species
SARS−CoV1 0 0.048 0.012 0.06 0.012 0 0.07 0.177 0.07 0.277 0.22 0.01 0.036 0.039 0.022 0.018
RsWIV1 0 0 0.002 0 0.001 0 0.001 0 0 0 0 0 0.005 0 0 0
Rs7327 0 0 0.021 0.022 0.001 0 0 0 0 0 0 0 0.003 0.016 0.012 0
Rs4874 0 0 0.015 0.003 0.002 0.003 0.002 0.004 0 0 0 0 0 0.008 0.006 0
Rs9401 0 0.004 0.003 0 0.001 0 0.001 0 0 0 0 0 0.01 0 0.012 0
Rs3367 0 0 0 0 0.001 0 0 0.002 0 0 0 0 0 0 0.003 0
Rs4231 0 0.009 0 0 0.015 0.012 0.007 0.013 0 0 0.029 0.068 0.005 0.028 0.003 0
Rs4247 0 0 0 0.005 0 0 0.01 0.006 0.061 0 0.181 0.054 0 0.006 0 0.018
Rs4237 0 0.004 0.012 0.003 0 0.012 0.029 0.034 0 0.004 0.015 0.015 0.01 0.029 0.013 0
As6526 0 0 0.006 0.005 0.005 0 0 0 0 0.074 0.179 0.036 0 0.011 0 0.019
Rs4255 0 0 0.002 0.005 0.001 0.003 0.016 0.02 0.031 0.021 0.016 0.031 0.03 0.028 0.009 0
Rs4081 0 0 0 0 0 0 0.015 0.021 0.019 0.03 0.009 0.026 0.005 0.004 0.024 0
bRp3 0 0.085 0.04 0.051 0.034 0.021 0.038 0.01 0.04 0.18 0.413 0.17 0.159 0.094 0.054 0.038
Rs672 0 0.042 0.025 0.054 0.035 0.027 0.038 0.044 0.028 0.011 0.029 0.032 0.015 0.021 0.031 0
RsYN2013 0 0.018 0.028 0.009 0.007 0.006 0.006 0.048 0.015 0.802 0.303 0.003 0.029 0.03 0.04 0.018
Rf4092 0 0.032 0.006 0.027 0.016 0.013 0.039 0.043 0.077 1.598 0.555 0.008 0.03 0.007 0.012 0
RfYNLF_34C 0 0.002 0 0 0.002 0 0 0 0 0 0 0 0 0 0 0
RfYNLF_31C 0 0 0 0 0 0 0 0 0 0 0 0 0 0.004 0 0
RsGX2013 0 0.108 0.077 0.038 0.059 0.053 0.058 0.033 0.055 0.404 0.401 0.209 0.063 0.086 0.039 0.037
1−2829 2829−4661 4661−7671 7671−9033 9033−14628 14628−15824 15824−18978 18978−20593 20593−21207 21207−22554 22554−23447 23447−25759 25759−26668 26668−28572 28572−29925 29925−31317
Nucleotide position on multiple sequence alignement
0.0 0.4 0.8 1.2

(B)

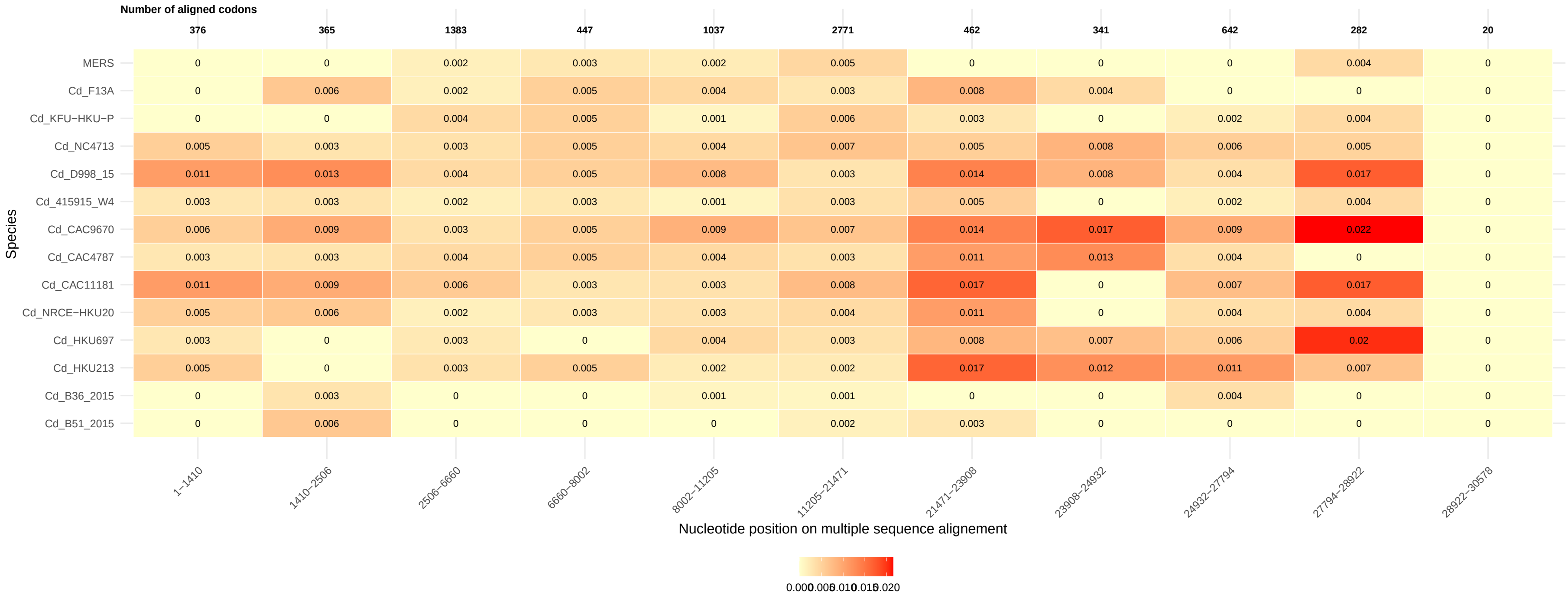
Number of aligned codons
376 365 1383 447 1037 2771 462 341 642 282 20
MERS 0 0 0.002 0.003 0.002 0.005 0 0 0 0.004 0
Cd_F13A 0 0.006 0.002 0.005 0.004 0.003 0.008 0.004 0 0 0
Cd_KFU−HKU−P 0 0 0.004 0.005 0.001 0.006 0.003 0 0.002 0.004 0
Cd_NC4713 0.005 0.003 0.003 0.005 0.004 0.007 0.005 0.008 0.006 0.005 0
Cd_D998_15 0.011 0.013 0.004 0.005 0.008 0.003 0.014 0.008 0.004 0.017 0
Cd_415915_W4 0.003 0.003 0.002 0.003 0.001 0.003 0.005 0 0.002 0.004 0
Cd_CAC9670 0.006 0.009 0.003 0.005 0.009 0.007 0.014 0.017 0.009 0.022 0
Cd_CAC4787 0.003 0.003 0.004 0.005 0.004 0.003 0.011 0.013 0.004 0 0
Cd_CAC11181 0.011 0.009 0.006 0.003 0.003 0.008 0.017 0 0.007 0.017 0
Cd_NRCE−HKU20 0.005 0.006 0.002 0.003 0.003 0.004 0.011 0 0.004 0.004 0
Cd_HKU697 0.003 0 0.003 0 0.004 0.003 0.008 0.007 0.006 0.02 0
Cd_HKU213 0.005 0 0.003 0.005 0.002 0.002 0.017 0.012 0.011 0.007 0
Cd_B36_2015 0 0.003 0 0 0.001 0.001 0 0 0.004 0 0
Cd_B51_2015 0 0.006 0 0 0 0.002 0.003 0 0 0 0
Species
1−1410 1410−2506 2506−6660 6660−8002 8002−11205 11205−21471 21471−23908 23908−24932 24932−27794 27794−28922 28922−30578
Nucleotide position on multiple sequence alignement
0.000 0.005 0.010 0.015 0.020

(C)

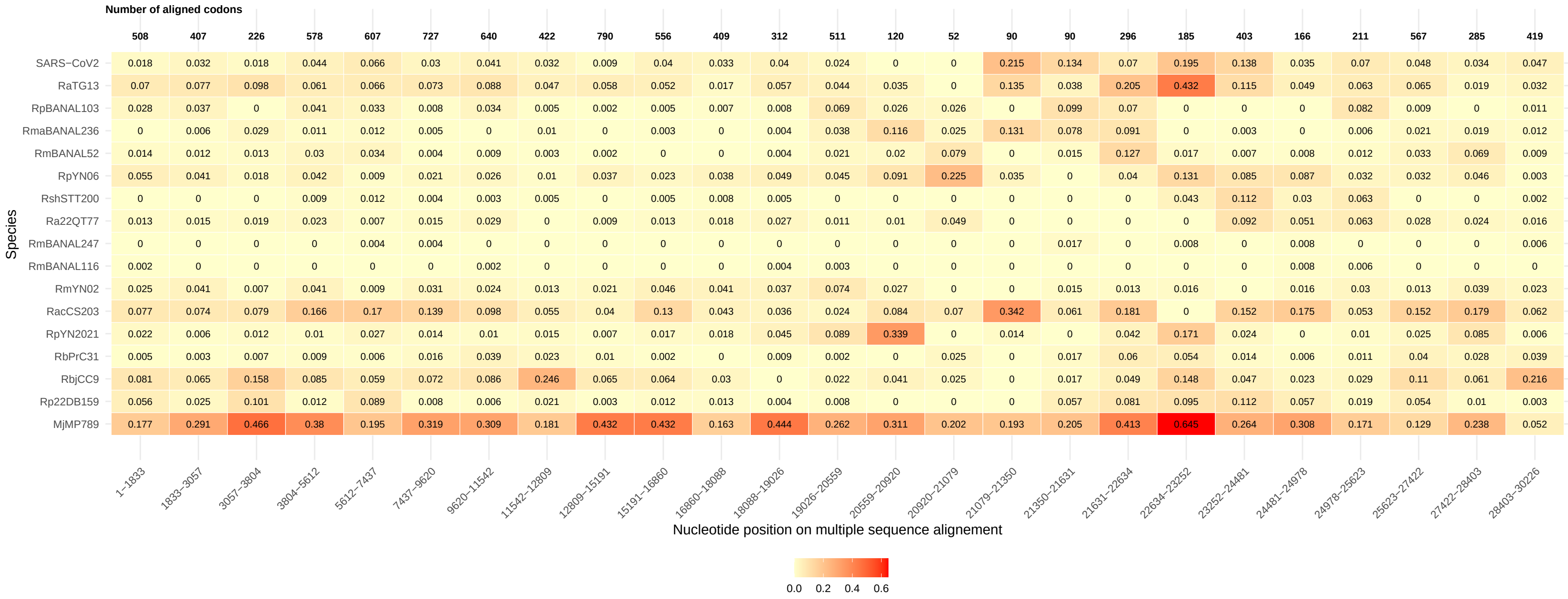
Number of aligned codons
508 407 226 578 607 727 640 422 790 556 409 312 511 120 52 90 90 296 185 403 166 211 567 285 419
SARS−CoV2 0.018 0.032 0.018 0.044 0.066 0.03 0.041 0.032 0.009 0.04 0.033 0.04 0.024 0 0 0.215 0.134 0.07 0.195 0.138 0.035 0.07 0.048 0.034 0.047
RaTG13 0.07 0.077 0.098 0.061 0.066 0.073 0.088 0.047 0.058 0.052 0.017 0.057 0.044 0.035 0 0.135 0.038 0.205 0.432 0.115 0.049 0.063 0.065 0.019 0.032
RpBANAL103 0.028 0.037 0 0.041 0.033 0.008 0.034 0.005 0.002 0.005 0.007 0.008 0.069 0.026 0.026 0 0.099 0.07 0 0 0 0.082 0.009 0 0.011
RmaBANAL236 0 0.006 0.029 0.011 0.012 0.005 0 0.01 0 0.003 0 0.004 0.038 0.116 0.025 0.131 0.078 0.091 0 0.003 0 0.006 0.021 0.019 0.012
RmBANAL52 0.014 0.012 0.013 0.03 0.034 0.004 0.009 0.003 0.002 0 0 0.004 0.021 0.02 0.079 0 0.015 0.127 0.017 0.007 0.008 0.012 0.033 0.069 0.009
RpYN06 0.055 0.041 0.018 0.042 0.009 0.021 0.026 0.01 0.037 0.023 0.038 0.049 0.045 0.091 0.225 0.035 0 0.04 0.131 0.085 0.087 0.032 0.032 0.046 0.003
RshSTT200 0 0 0 0.009 0.012 0.004 0.003 0.005 0 0.005 0.008 0.005 0 0 0 0 0 0 0.043 0.112 0.03 0.063 0 0 0.002
Ra22QT77 0.013 0.015 0.019 0.023 0.007 0.015 0.029 0 0.009 0.013 0.018 0.027 0.011 0.01 0.049 0 0 0 0 0.092 0.051 0.063 0.028 0.024 0.016
RmBANAL247 0 0 0 0 0.004 0.004 0 0 0 0 0 0 0 0 0 0 0.017 0 0.008 0 0.008 0 0 0 0.006
RmBANAL116 0.002 0 0 0 0 0 0.002 0 0 0 0 0.004 0.003 0 0 0 0 0 0 0 0.008 0.006 0 0 0
RmYN02 0.025 0.041 0.007 0.041 0.009 0.031 0.024 0.013 0.021 0.046 0.041 0.037 0.074 0.027 0 0 0.015 0.013 0.016 0 0.016 0.03 0.013 0.039 0.023
RacCS203 0.077 0.074 0.079 0.166 0.17 0.139 0.098 0.055 0.04 0.13 0.043 0.036 0.024 0.084 0.07 0.342 0.061 0.181 0 0.152 0.175 0.053 0.152 0.179 0.062
RpYN2021 0.022 0.006 0.012 0.01 0.027 0.014 0.01 0.015 0.007 0.017 0.018 0.045 0.089 0.339 0 0.014 0 0.042 0.171 0.024 0 0.01 0.025 0.085 0.006
RbPrC31 0.005 0.003 0.007 0.009 0.006 0.016 0.039 0.023 0.01 0.002 0 0.009 0.002 0 0.025 0 0.017 0.06 0.054 0.014 0.006 0.011 0.04 0.028 0.039
RbjCC9 0.081 0.065 0.158 0.085 0.059 0.072 0.086 0.246 0.065 0.064 0.03 0 0.022 0.041 0.025 0 0.017 0.049 0.148 0.047 0.023 0.029 0.11 0.061 0.216
Rp22DB159 0.056 0.025 0.101 0.012 0.089 0.008 0.006 0.021 0.003 0.012 0.013 0.004 0.008 0 0 0 0.057 0.081 0.095 0.112 0.057 0.019 0.054 0.01 0.003
MjMP789 0.177 0.291 0.466 0.38 0.195 0.319 0.309 0.181 0.432 0.432 0.163 0.444 0.262 0.311 0.202 0.193 0.205 0.413 0.645 0.264 0.308 0.171 0.129 0.238 0.052
Species
1−1833 1833−3057 3057−3804 3804−5612 5612−7437 7437−9620 9620−11542 11542−12809 12809−15191 15191−16860 16860−18088 18088−19026 19026−20559 20559−20920 20920−21079 21079−21350 21350−21631 21631−22634 22634−23252 23252−24481 24481−24978 24978−25623 25623−27422 27422−28403 28403−30226
Nucleotide position on multiple sequence alignement
0.0 0.2 0.4 0.6

(D)

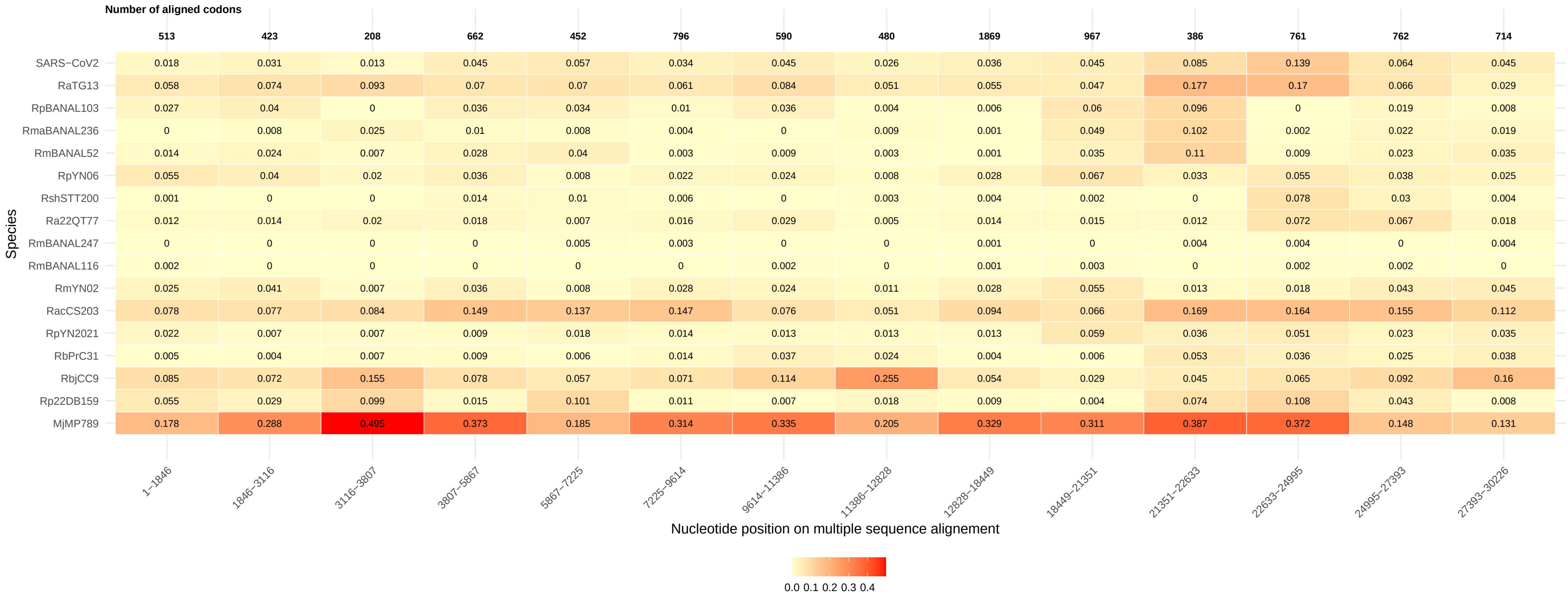
Number of aligned codons
513 423 208 662 452 796 590 480 1869 967 386 761 762 714
SARS−CoV2 0.018 0.031 0.013 0.045 0.057 0.034 0.045 0.026 0.036 0.045 0.085 0.139 0.064 0.045
RaTG13 0.058 0.074 0.093 0.07 0.07 0.061 0.084 0.051 0.055 0.047 0.177 0.17 0.066 0.029
RpBANAL103 0.027 0.04 0 0.036 0.034 0.01 0.036 0.004 0.006 0.06 0.096 0 0.019 0.008
RmaBANAL236 0 0.008 0.025 0.01 0.008 0.004 0 0.009 0.001 0.049 0.102 0.002 0.022 0.019
RmBANAL52 0.014 0.024 0.007 0.028 0.04 0.003 0.009 0.003 0.001 0.035 0.11 0.009 0.023 0.035
RpYN06 0.055 0.04 0.02 0.036 0.008 0.022 0.024 0.008 0.028 0.067 0.033 0.055 0.038 0.025
RshSTT200 0.001 0 0 0.014 0.01 0.006 0 0.003 0.004 0.002 0 0.078 0.03 0.004
Ra22QT77 0.012 0.014 0.02 0.018 0.007 0.016 0.029 0.005 0.014 0.015 0.012 0.072 0.067 0.018
RmBANAL247 0 0 0 0 0.005 0.003 0 0 0.001 0 0.004 0.004 0 0.004
RmBANAL116 0.002 0 0 0 0 0 0.002 0 0.001 0.003 0 0.002 0.002 0
RmYN02 0.025 0.041 0.007 0.036 0.008 0.028 0.024 0.011 0.028 0.055 0.013 0.018 0.043 0.045
RacCS203 0.078 0.077 0.084 0.149 0.137 0.147 0.076 0.051 0.094 0.066 0.169 0.164 0.155 0.112
RpYN2021 0.022 0.007 0.007 0.009 0.018 0.014 0.013 0.013 0.013 0.059 0.036 0.051 0.023 0.035
RbPrC31 0.005 0.004 0.007 0.009 0.006 0.014 0.037 0.024 0.004 0.006 0.053 0.036 0.025 0.038
RbjCC9 0.085 0.072 0.155 0.078 0.057 0.071 0.114 0.255 0.054 0.029 0.045 0.065 0.092 0.16
Rp22DB159 0.055 0.029 0.099 0.015 0.101 0.011 0.007 0.018 0.009 0.004 0.074 0.108 0.043 0.008
MjMP789 0.178 0.288 0.495 0.373 0.185 0.314 0.335 0.205 0.329 0.311 0.387 0.372 0.148 0.131
Species
1–1846 1846–3116 3116–3807 3807–5867 5867–7225 7225–9614 9614–11386 11386–12828 12828–18449 18449–21351 21351–22633 22633–24995 24995–27393 27393–30226
Nucleotide position on multiple sequence alignement
0.0 0.1 0.2 0.3 0.4

(E)

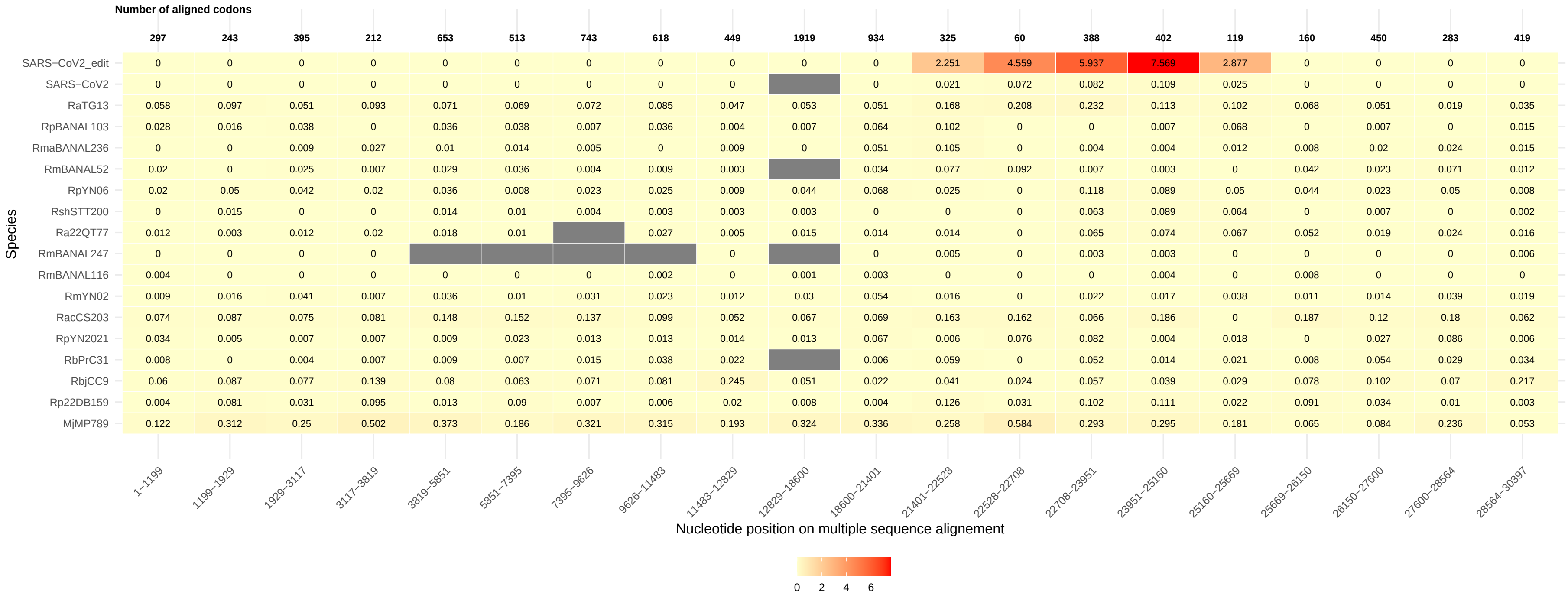
Number of aligned codons
297 243 395 212 653 513 743 618 449 1919 934 325 60 388 402 119 160 450 283 419
SARS−CoV2_edit 0 0 0 0 0 0 0 0 0 0 0 2.251 4.559 5.937 7.569 2.877 0 0 0 0
SARS−CoV2 0 0 0 0 0 0 0 0 0 0 0.021 0.072 0.082 0.109 0.025 0 0 0 0
RaTG13 0.058 0.097 0.051 0.093 0.071 0.069 0.072 0.085 0.047 0.053 0.051 0.168 0.208 0.232 0.113 0.102 0.068 0.051 0.019 0.035
RpBANAL103 0.028 0.016 0.038 0 0.036 0.038 0.007 0.036 0.004 0.007 0.064 0.102 0 0 0.007 0.068 0 0.007 0 0.015
RmaBANAL236 0 0 0.009 0.027 0.01 0.014 0.005 0 0.009 0 0.051 0.105 0 0.004 0.004 0.012 0.008 0.02 0.024 0.015
RmBANAL52 0.02 0 0.025 0.007 0.029 0.036 0.004 0.009 0.003 0.034 0.077 0.092 0.007 0.003 0 0.042 0.023 0.071 0.012
RpYN06 0.02 0.05 0.042 0.02 0.036 0.008 0.023 0.025 0.009 0.044 0.068 0.025 0 0.118 0.089 0.05 0.044 0.023 0.05 0.008
RshSTT200 0 0.015 0 0 0.014 0.01 0.004 0.003 0.003 0.003 0 0 0 0.063 0.089 0.064 0 0.007 0 0.002
Ra22QT77 0.012 0.003 0.012 0.02 0.018 0.01 0.027 0.005 0.015 0.014 0.014 0 0.065 0.074 0.067 0.052 0.019 0.024 0.016
RmBANAL247 0 0 0 0 0 0 0.005 0 0.003 0.003 0 0 0 0 0.006
RmBANAL116 0.004 0 0 0 0 0 0 0.002 0 0.001 0.003 0 0 0 0.004 0 0.008 0 0 0
RmYN02 0.009 0.016 0.041 0.007 0.036 0.01 0.031 0.023 0.012 0.03 0.054 0.016 0 0.022 0.017 0.038 0.011 0.014 0.039 0.019
RacCS203 0.074 0.087 0.075 0.081 0.148 0.152 0.137 0.099 0.052 0.067 0.069 0.163 0.162 0.066 0.186 0 0.187 0.12 0.18 0.062
RpYN2021 0.034 0.005 0.007 0.007 0.009 0.023 0.013 0.013 0.014 0.013 0.067 0.006 0.076 0.082 0.004 0.018 0 0.027 0.086 0.006
RbPrC31 0.008 0 0.004 0.007 0.009 0.007 0.015 0.038 0.022 0.006 0.059 0 0.052 0.014 0.021 0.008 0.054 0.029 0.034
RbjCC9 0.06 0.087 0.077 0.139 0.08 0.063 0.071 0.081 0.245 0.051 0.022 0.041 0.024 0.057 0.039 0.029 0.078 0.102 0.07 0.217
Rp22DB159 0.004 0.081 0.031 0.095 0.013 0.09 0.007 0.006 0.02 0.008 0.004 0.126 0.031 0.102 0.111 0.022 0.091 0.034 0.01 0.003
MjMP789 0.122 0.312 0.25 0.502 0.373 0.186 0.321 0.315 0.193 0.324 0.336 0.258 0.584 0.293 0.295 0.181 0.065 0.084 0.236 0.053
Species
1−1199 1199−1929 1929−3117 3117−3819 3819−5851 5851−7395 7395−9626 9626−11483 11483−12829 12829−18600 18600−21401 21401−22528 22528−22708 22708−23951 23951−25160 25160−25669 25669−26150 26150−27600 27600−28564 28564−30397
Nucleotide position on multiple sequence alignement
0 2 4 6

(F)

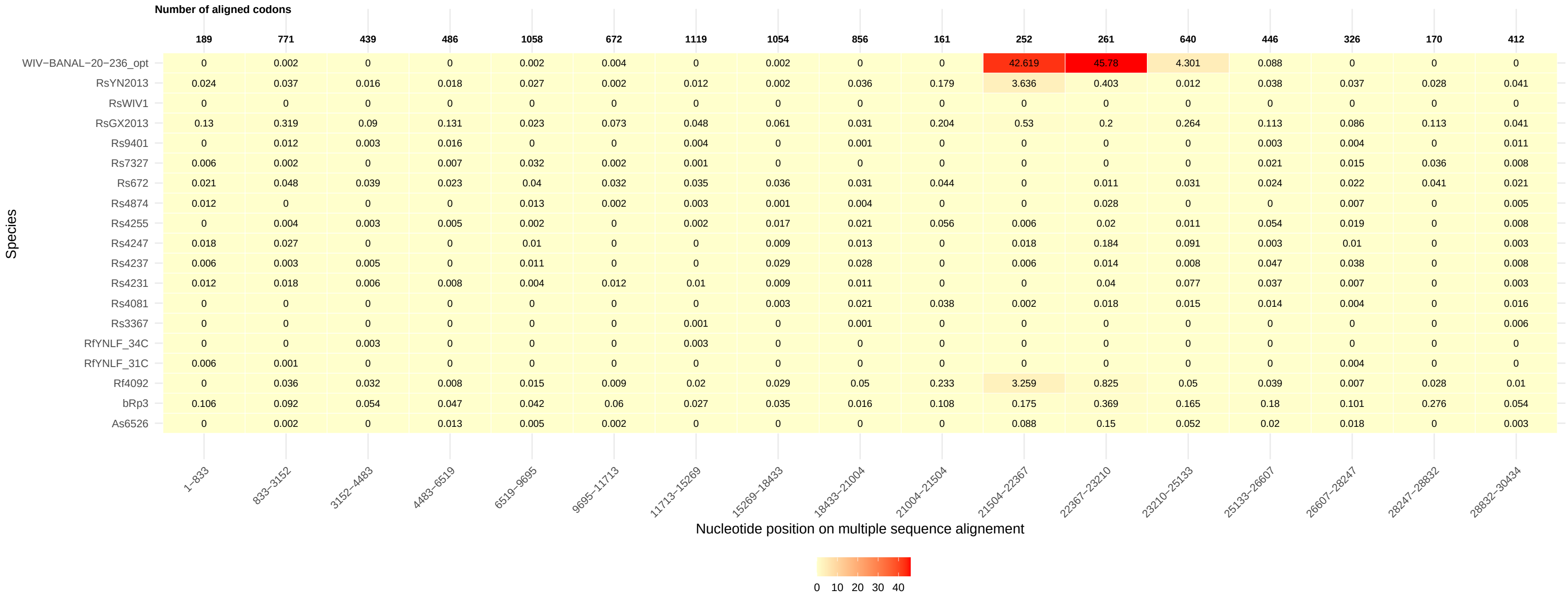
Number of aligned codons
189 771 439 486 1058 672 1119 1054 856 161 252 261 640 446 326 170 412
WIV−BANAL−20−236_opt 0 0.002 0 0 0.002 0.004 0 0.002 0 0 42.619 45.78 4.301 0.088 0 0 0
RsYN2013 0.024 0.037 0.016 0.018 0.027 0.002 0.012 0.002 0.036 0.179 3.636 0.403 0.012 0.038 0.037 0.028 0.041
RsWIV1 0 0 0 0 0 0 0 0 0 0 0 0 0 0 0 0 0
RsGX2013 0.13 0.319 0.09 0.131 0.023 0.073 0.048 0.061 0.031 0.204 0.53 0.2 0.264 0.113 0.086 0.113 0.041
Rs9401 0 0.012 0.003 0.016 0 0 0.004 0 0.001 0 0 0 0 0.003 0.004 0 0.011
Rs7327 0.006 0.002 0 0.007 0.032 0.002 0.001 0 0 0 0 0 0 0.021 0.015 0.036 0.008
Rs672 0.021 0.048 0.039 0.023 0.04 0.032 0.035 0.036 0.031 0.044 0 0.011 0.031 0.024 0.022 0.041 0.021
Rs4874 0.012 0 0 0 0.013 0.002 0.003 0.001 0.004 0 0 0.028 0 0 0.007 0 0.005
Rs4255 0 0.004 0.003 0.005 0.002 0 0.002 0.017 0.021 0.056 0.006 0.02 0.011 0.054 0.019 0 0.008
Rs4247 0.018 0.027 0 0 0.01 0 0 0.009 0.013 0 0.018 0.184 0.091 0.003 0.01 0 0.003
Rs4237 0.006 0.003 0.005 0 0.011 0 0 0.029 0.028 0 0.006 0.014 0.008 0.047 0.038 0 0.008
Rs4231 0.012 0.018 0.006 0.008 0.004 0.012 0.01 0.009 0.011 0 0 0.04 0.077 0.037 0.007 0 0.003
Rs4081 0 0 0 0 0 0 0 0.003 0.021 0.038 0.002 0.018 0.015 0.014 0.004 0 0.016
Rs3367 0 0 0 0 0 0 0.001 0 0.001 0 0 0 0 0 0 0 0.006
RfYNLF_34C 0 0 0.003 0 0 0 0.003 0 0 0 0 0 0 0 0 0 0
RfYNLF_31C 0.006 0.001 0 0 0 0 0 0 0 0 0 0 0 0 0.004 0 0
Rf4092 0 0.036 0.032 0.008 0.015 0.009 0.02 0.029 0.05 0.233 3.259 0.825 0.05 0.039 0.007 0.028 0.01
bRp3 0.106 0.092 0.054 0.047 0.042 0.06 0.027 0.035 0.016 0.108 0.175 0.369 0.165 0.18 0.101 0.276 0.054
As6526 0 0.002 0 0.013 0.005 0.002 0 0 0 0 0.088 0.15 0.052 0.02 0.018 0 0.003
Species
1−833 833−3152 3152−4483 4483−6519 6519−9695 9695−11713 11713−15269 15269−18433 18433−21004 21004−21504 21504−22367 22367−23210 23210−25133 25133−26607 26607−28247 28247−28832 28832−30434
Nucleotide position on multiple sequence alignement
0 10 20 30 40

(G)

Number of aligned codons

| Species | 1–3215 | 3215–4485 | 4485–6478 | 6478–7599 | 7599–9678 | 9678–11738 | 11738–14310 | 14310–17937 | 17937–21005 | 21005–21478 | 21478–22336 | 22336–23301 | 23301–24637 | 24637–26607 | 26607–28273 | 28273–28898 | 28898–30443 |
|---|---|---|---|---|---|---|---|---|---|---|---|---|---|---|---|---|---|
| Number of aligned codons | **981** | **420** | **475** | **370** | **693** | **686** | **790** | **1209** | **1023** | **157** | **224** | **303** | **445** | **612** | **334** | **184** | **393** |
| WIV−BANAL−20−236_nat | 0.001 | 0 | 0 | 0 | 0.003 | 0.003 | 0 | 0.002 | 0 | 0 | 1.747 | 1.476 | 0.934 | 0.275 | 0 | 0 | 0 |
| RsYN2013 | 0.04 | 0.014 | 0.013 | 0.028 | 0.017 | 0.003 | 0.011 | 0.003 | 0.037 | 0.182 | 4.722 | 0.399 | 0.008 | 0.013 | 0.036 | 0.025 | 0.043 |
| RsWIV1 | 0 | 0 | 0 | 0 | 0 | 0 | 0 | 0 | 0 | 0 | 0 | 0 | 0 | 0 | 0 | 0 | 0 |
| RsGX2013 | 0.271 | 0.083 | 0.136 | 0.014 | 0.03 | 0.071 | 0.046 | 0.062 | 0.031 | 0.195 | 0.525 | 0.284 | 0.231 | 0.146 | 0.084 | 0.1 | 0.043 |
| Rs9401 | 0.01 | 0.003 | 0.014 | 0 | 0 | 0 | 0.001 | 0.001 | 0.001 | 0 | 0 | 0 | 0 | 0.002 | 0.004 | 0 | 0.011 |
| Rs7327 | 0.002 | 0 | 0.005 | 0.042 | 0.013 | 0.002 | 0 | 0.001 | 0 | 0 | 0 | 0 | 0 | 0.015 | 0.014 | 0.032 | 0.008 |
| Rs672 | 0.036 | 0.04 | 0.024 | 0.015 | 0.042 | 0.031 | 0.039 | 0.035 | 0.032 | 0.045 | 0 | 0.01 | 0.025 | 0.027 | 0.021 | 0.038 | 0.022 |
| Rs4874 | 0.002 | 0 | 0 | 0.028 | 0.005 | 0.002 | 0.003 | 0.002 | 0.006 | 0 | 0 | 0.021 | 0 | 0 | 0.007 | 0 | 0.006 |
| Rs4255 | 0.005 | 0.003 | 0.005 | 0 | 0.003 | 0 | 0.001 | 0.016 | 0.018 | 0.057 | 0.007 | 0.023 | 0 | 0.052 | 0.018 | 0 | 0.008 |
| Rs4247 | 0.023 | 0 | 0 | 0 | 0.009 | 0 | 0 | 0.008 | 0.012 | 0 | 0.021 | 0.189 | 0.045 | 0.013 | 0.01 | 0 | 0.003 |
| Rs4237 | 0.008 | 0.006 | 0 | 0.02 | 0.004 | 0 | 0 | 0.028 | 0.027 | 0 | 0.007 | 0.018 | 0.008 | 0.045 | 0.037 | 0 | 0.009 |
| Rs4231 | 0.016 | 0.006 | 0.008 | 0 | 0.005 | 0.012 | 0.01 | 0.01 | 0.011 | 0 | 0 | 0.05 | 0.057 | 0.049 | 0.007 | 0 | 0.003 |
| Rs4081 | 0 | 0 | 0 | 0 | 0 | 0 | 0 | 0.001 | 0.02 | 0.038 | 0 | 0.015 | 0.027 | 0.024 | 0.004 | 0 | 0.017 |
| Rs3367 | 0 | 0 | 0 | 0 | 0 | 0 | 0 | 0.001 | 0.001 | 0 | 0 | 0 | 0 | 0 | 0 | 0 | 0.006 |
| RfYNLF_34C | 0 | 0.003 | 0 | 0 | 0 | 0 | 0.004 | 0 | 0 | 0 | 0 | 0 | 0 | 0 | 0 | 0 | 0 |
| RfYNLF_31C | 0.002 | 0 | 0 | 0 | 0 | 0 | 0 | 0 | 0 | 0 | 0 | 0 | 0 | 0 | 0.004 | 0 | 0 |
| Rf4092 | 0.031 | 0.033 | 0.005 | 0.006 | 0.019 | 0.009 | 0.022 | 0.019 | 0.048 | 0.225 | 3.416 | 0.817 | 0.051 | 0.024 | 0.007 | 0.026 | 0.01 |
| bRp3 | 0.089 | 0.05 | 0.045 | 0.04 | 0.041 | 0.058 | 0.029 | 0.031 | 0.023 | 0.1 | 0.139 | 0.351 | 0.152 | 0.188 | 0.098 | 0.27 | 0.051 |
| As6526 | 0.001 | 0 | 0.013 | 0 | 0.007 | 0.002 | 0 | 0 | 0.005 | 0 | 0.095 | 0.143 | 0.069 | 0.019 | 0.018 | 0 | 0.003 |

Species

Nucleotide position on multiple sequence alignement

Colour scale: 0 1 2 3 4

Figure S14. Estimation of $\mathrm{d}N/\mathrm{d}S$ across all the segments of the three datasets. (A) SARS-CoV-1, (B) MERS and (C) SARS-CoV-2, (D) SARS-CoV-2 with only 14 segments (larger segments), (E) CoV-2-S$^{\mathrm{edit}}$, (F) WIV-BANAL-20-236$^{\mathrm{opt}}$, and (G) WIV-BANAL-20-236$^{\mathrm{nat}}$. Virus sequences are listed on the left. The $x$-axis shows consecutive segments and their corresponding nucleotide positions in the multiple sequence alignment. Segment length (in number of codons) is indicated in bold in the first row of the table. The $\mathrm{d}N/\mathrm{d}S$ ratio obtained with CODEML model 0 for each segment is shown in bold in the last row of the table. For each segment and each terminal branch of the phylogeny (i.e., each leaf sequence), the estimated $\mathrm{d}N/\mathrm{d}S$ values are reported numerically. Colors represent the $p$-value of the likelihood ratio test (LRT) for the presence of a different rate of amino-acid evolution in the terminal branch leading to each virus for each segment (see Materials and Methods for details). Red: $\mathrm{d}N/\mathrm{d}S$ is significantly ($p$-value $< 0.001$) higher in the terminal branch leading to the corresponding virus than in the remaining branches of the tree, green: $\mathrm{d}N/\mathrm{d}S$ is significantly ($p$-value $< 0.001$) lower than in the remaining branches of the tree, blue: no significant difference, grey: test not done due to limited data (number of informative codons too small according to CODEML, leading to $\mathrm{d}S = 0$).

(A)

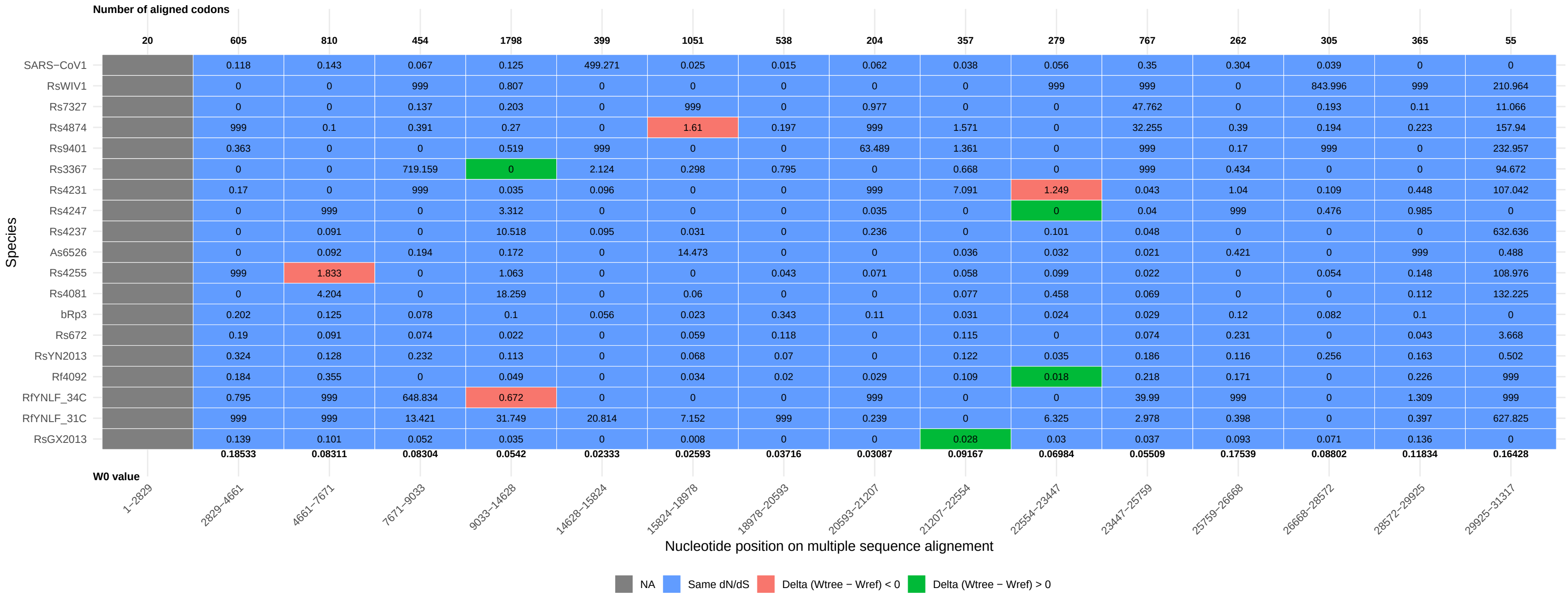
Number of aligned codons
20 605 810 454 1798 399 1051 538 204 357 279 767 262 305 365 55
SARS−CoV1 0.118 0.143 0.067 0.125 499.271 0.025 0.015 0.062 0.038 0.056 0.35 0.304 0.039 0 0
RsWIV1 0 0 999 0.807 0 0 0 0 0 999 999 0 843.996 999 210.964
Rs7327 0 0 0.137 0.203 0 999 0 0.977 0 0 47.762 0 0.193 0.11 11.066
Rs4874 999 0.1 0.391 0.27 0 1.61 0.197 999 1.571 0 32.255 0.39 0.194 0.223 157.94
Rs9401 0.363 0 0 0.519 999 0 0 63.489 1.361 0 999 0.17 999 0 232.957
Rs3367 0 0 719.159 0 2.124 0.298 0.795 0 0.668 0 999 0.434 0 0 94.672
Rs4231 0.17 0 999 0.035 0.096 0 0 999 7.091 1.249 0.043 1.04 0.109 0.448 107.042
Rs4247 0 999 0 3.312 0 0 0 0.035 0 0 0.04 999 0.476 0.985 0
Rs4237 0 0.091 0 10.518 0.095 0.031 0 0.236 0 0.101 0.048 0 0 0 632.636
As6526 0 0.092 0.194 0.172 0 14.473 0 0 0.036 0.032 0.021 0.421 0 999 0.488
Rs4255 999 1.833 0 1.063 0 0 0.043 0.071 0.058 0.099 0.022 0 0.054 0.148 108.976
Rs4081 0 4.204 0 18.259 0 0.06 0 0 0.077 0.458 0.069 0 0 0.112 132.225
bRp3 0.202 0.125 0.078 0.1 0.056 0.023 0.343 0.11 0.031 0.024 0.029 0.12 0.082 0.1 0
Rs672 0.19 0.091 0.074 0.022 0 0.059 0.118 0 0.115 0 0.074 0.231 0 0.043 3.668
RsYN2013 0.324 0.128 0.232 0.113 0 0.068 0.07 0 0.122 0.035 0.186 0.116 0.256 0.163 0.502
Rf4092 0.184 0.355 0 0.049 0 0.034 0.02 0.029 0.109 0.018 0.218 0.171 0 0.226 999
RfYNLF_34C 0.795 999 648.834 0.672 0 0 0 999 0 0 39.99 999 0 1.309 999
RfYNLF_31C 999 999 13.421 31.749 20.814 7.152 999 0.239 0 6.325 2.978 0.398 0 0.397 627.825
RsGX2013 0.139 0.101 0.052 0.035 0 0.008 0 0 0.028 0.03 0.037 0.093 0.071 0.136 0
Species
0.18533 0.08311 0.08304 0.0542 0.02333 0.02593 0.03716 0.03087 0.09167 0.06984 0.05509 0.17539 0.08802 0.11834 0.16428
W0 value
1−2829 2829−4661 4661−7671 7671−9033 9033−14628 14628−15824 15824−18978 18978−20593 20593−21207 21207−22554 22554−23447 23447−25759 25759−26668 26668−28572 28572−29925 29925−31317
Nucleotide position on multiple sequence alignement
NA
Same dN/dS
Delta (Wtree − Wref) < 0
Delta (Wtree − Wref) > 0

(B)

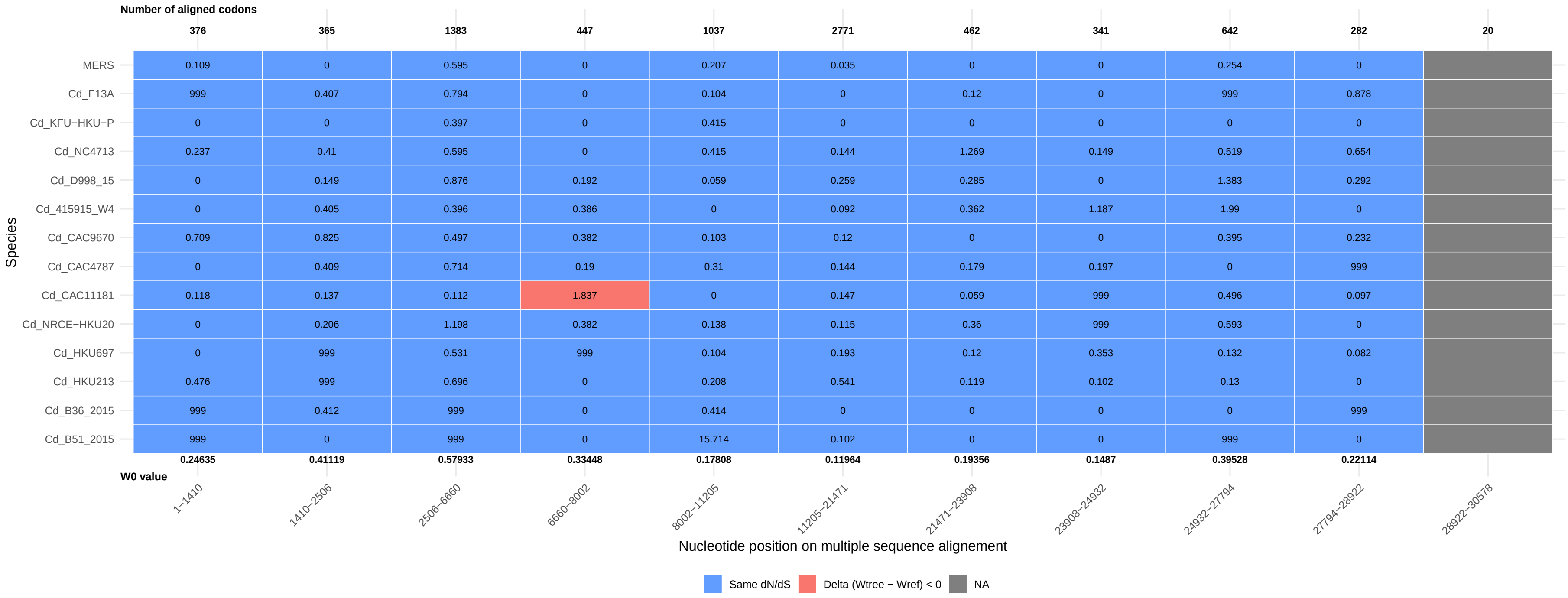
Number of aligned codons
376 365 1383 447 1037 2771 462 341 642 282 20
MERS 0.109 0 0.595 0 0.207 0.035 0 0 0.254 0
Cd_F13A 999 0.407 0.794 0 0.104 0 0.12 0 999 0.878
Cd_KFU−HKU−P 0 0 0.397 0 0.415 0 0 0 0 0
Cd_NC4713 0.237 0.41 0.595 0 0.415 0.144 1.269 0.149 0.519 0.654
Cd_D998_15 0 0.149 0.876 0.192 0.059 0.259 0.285 0 1.383 0.292
Cd_415915_W4 0 0.405 0.396 0.386 0 0.092 0.362 1.187 1.99 0
Cd_CAC9670 0.709 0.825 0.497 0.382 0.103 0.12 0 0 0.395 0.232
Cd_CAC4787 0 0.409 0.714 0.19 0.31 0.144 0.179 0.197 0 999
Cd_CAC11181 0.118 0.137 0.112 1.837 0 0.147 0.059 999 0.496 0.097
Cd_NRCE−HKU20 0 0.206 1.198 0.382 0.138 0.115 0.36 999 0.593 0
Cd_HKU697 0 999 0.531 999 0.104 0.193 0.12 0.353 0.132 0.082
Cd_HKU213 0.476 999 0.696 0 0.208 0.541 0.119 0.102 0.13 0
Cd_B36_2015 999 0.412 999 0 0.414 0 0 0 0 999
Cd_B51_2015 999 0 999 0 15.714 0.102 0 0 999 0
Species
0.24635 0.41119 0.57933 0.33448 0.17808 0.11964 0.19356 0.1487 0.39528 0.22114
W0 value
1−1410 1410−2506 2506−6660 6660−8002 8002−11205 11205−21471 21471−23908 23908−24932 24932−27794 27794−28922 28922−30578
Nucleotide position on multiple sequence alignement
Same dN/dS
Delta (Wtree − Wref) < 0
NA

(C)

Number of aligned codons / W0 value / Nucleotide position on multiple sequence alignement

| Species | 1−1833 | 1833−3057 | 3057−3804 | 3804−5612 | 5612−7437 | 7437−9620 | 9620−11542 | 11542−12809 | 12809−15191 | 15191−16860 | 16860−18088 | 18088−19026 | 19026−20559 |
|---|---|---|---|---|---|---|---|---|---|---|---|---|---|
| Number of aligned codons | 508 | 407 | 226 | 578 | 607 | 727 | 640 | 422 | 790 | 556 | 409 | 312 | 511 |
| SARS−CoV2 | 0.204 | 0.035 | 0 | 0.053 | 0.078 | 0.02 | 0.051 | 0 | 0 | 0 | 0 | 0 | 0.07 |
| RaTG13 | 0.078 | 0.056 | 0.164 | 0.072 | 0.112 | 0.017 | 0.016 | 0 | 0.019 | 0.015 | 0.063 | 0.049 | 0.116 |
| RpBANAL103 | 0.13 | 0.029 | 999 | 0.076 | 0.09 | 0.08 | 0.04 | 0 | 0 | 0 | 0.163 | 0 | 0.035 |
| RmaBANAL236 | 16.023 | 0.196 | 0.4 | 0 | 0.161 | 0.351 | 999 | 0.105 | 0 | 0 | 999 | 0 | 0.065 |
| RmBANAL52 | 0.317 | 0.094 | 0.288 | 0.105 | 0.087 | 0 | 0 | 0 | 0 | 0 | 0 | 0.312 | 0.163 |
| RpYN06 | 0.023 | 0.081 | 0.107 | 0.02 | 0.084 | 0.057 | 0 | 0 | 0.015 | 0 | 0.028 | 0.028 | 0.092 |
| RshSTT200 | 999 | 27.795 | 7.647 | 0.043 | 0.063 | 0.142 | 0 | 0.205 | 0 | 0 | 0 | 0 | 2.257 |
| Ra22QT77 | 0.137 | 0.22 | 0.615 | 0 | 0.441 | 0.079 | 0.049 | 0 | 0.257 | 0.118 | 0 | 0 | 0 |
| RmBANAL247 | 2.643 | 999 | 0 | 999 | 0.69 | 0 | 999 | 0 | 1.815 | 121.649 | 0 | 0 | 0 |
| RmBANAL116 | 0.382 | 21.73 | 999 | 999 | 999 | 0 | 0 | 0 | 10.704 | 0 | 0.08 | 0 | 0 |
| RmYN02 | 0.107 | 0.108 | 0.619 | 0.094 | 0.429 | 0.077 | 0.144 | 0.08 | 0.027 | 0 | 0.026 | 0.037 | 0.023 |
| RacCS203 | 0.174 | 0.113 | 0.123 | 0.058 | 0.085 | 0.04 | 0.028 | 0.038 | 0.028 | 0 | 0.025 | 0.118 | 0.107 |
| RpYN2021 | 0.152 | 0.189 | 0 | 0.381 | 0.247 | 0.169 | 0.206 | 0.137 | 0.324 | 0.047 | 0 | 0.064 | 0.01 |
| RbPrC31 | 0 | 0.771 | 0.269 | 0 | 0.714 | 0.096 | 0 | 0 | 0.164 | 0.65 | 0 | 0 | 0 |
| RbjCC9 | 0.088 | 0.154 | 0.342 | 0.12 | 0.088 | 0.065 | 0.024 | 0.022 | 0.015 | 0.037 | 0.028 | 10.395 | 0.12 |
| Rp22DB159 | 0.093 | 0.278 | 0.121 | 0.194 | 0.099 | 0 | 0 | 0.051 | 0.165 | 0.065 | 0 | 0.671 | 0 |
| MjMP789 | 0.077 | 0.066 | 0.15 | 0.049 | 0.053 | 0.027 | 0.002 | 0.006 | 0.004 | 0.002 | 0 | 0.006 | 0.003 |
| W0 value | 0.10177 | 0.09448 | 0.20058 | 0.07095 | 0.10161 | 0.0375 | 0.02177 | 0.01583 | 0.02316 | 0.00931 | 0.0075 | 0.02155 | 0.03403 |

| Species | 20559−20920 | 20920−21079 | 21079−21350 | 21350−21631 | 21631−22634 | 22634−23252 | 23252−24481 | 24481−24978 | 24978−25623 | 25623−27422 | 27422−28403 | 28403−30226 |
|---|---|---|---|---|---|---|---|---|---|---|---|---|
| Number of aligned codons | 120 | 52 | 90 | 90 | 296 | 185 | 403 | 166 | 211 | 567 | 285 | 419 |
| SARS−CoV2 | 0 | 0 | 0 | 0 | 0.086 | 0.035 | 0 | 0 | 0.06 | 0.051 | 0.232 | 0.071 |
| RaTG13 | 0 | 0 | 0 | 0 | 0.007 | 0.102 | 0 | 0 | 0.065 | 0.025 | 0.252 | 0.07 |
| RpBANAL103 | 0 | 0 | 0 | 0 | 0 | 0.102 | 999 | 0 | 0.051 | 0 | 0 | 0.101 |
| RmaBANAL236 | 0 | 0 | 0 | 0 | 0.015 | 533.959 | 0 | 0 | 0.339 | 0.047 | 0.164 | 0.179 |
| RmBANAL52 | 0.177 | 0 | 0 | 0.322 | 0.011 | 0 | 0.157 | 0 | 0.346 | 0.223 | 0.067 | 0.126 |
| RpYN06 | 0 | 0 | 0 | 0 | 0 | 0 | 0.056 | 0.031 | 0 | 0.102 | 0.068 | 0 |
| RshSTT200 | 0 | 0 | 0 | 0 | 10.329 | 0 | 0.019 | 0.179 | 0.066 | 16.627 | 999 | 0 |
| Ra22QT77 | 0.356 | 0 | 0 | 0 | 15.995 | 999 | 0.047 | 0 | 0.177 | 0.058 | 0 | 0.137 |
| RmBANAL247 | 0 | 999 | 0 | 0 | 0 | 0 | 5.184 | 0 | 999 | 999 | 999 | 0.191 |
| RmBANAL116 | 0.025 | 0 | 0.21 | 0 | 0 | 0 | 999 | 0.344 | 0 | 999 | 0 | 0.614 |
| RmYN02 | 0 | 0 | 999 | 0 | 0.444 | 0 | 8.155 | 0 | 0.139 | 0.376 | 0.243 | 0.144 |
| RacCS203 | 0 | 0 | 0 | 0.077 | 0.098 | 0.126 | 0.044 | 0.031 | 0.031 | 0.027 | 0.061 | 0.05 |
| RpYN2021 | 0.011 | 0 | 0 | 7.106 | 0.106 | 0.044 | 0.09 | 999 | 0 | 0.065 | 0.223 | 0.188 |
| RbPrC31 | 0 | 0 | 0 | 0 | 0.025 | 0.017 | 0.079 | 0 | 0 | 0.041 | 0.221 | 0.086 |
| RbjCC9 | 0.172 | 0 | 0.32 | 0.265 | 0 | 0.051 | 0.042 | 0.118 | 0.153 | 0.033 | 0.152 | 0.089 |
| Rp22DB159 | 0.011 | 0 | 0 | 0 | 0.106 | 0 | 0.013 | 0.047 | 0.443 | 0.086 | 0 | 0 |
| MjMP789 | 0 | 0 | 0 | 0 | 0.021 | 0.011 | 0.015 | 0.026 | 0 | 0.038 | 0.026 | 0.14 |
| W0 value | 0.01052 | 0.00469 | 0.00663 | 0.04543 | 0.04362 | 0.03577 | 0.05323 | 0.02237 | 0.05765 | 0.05985 | 0.2019 | 0.0983 |

Same dN/dS | Delta (Wtree − Wref) < 0 | Delta (Wtree − Wref) > 0

(D)

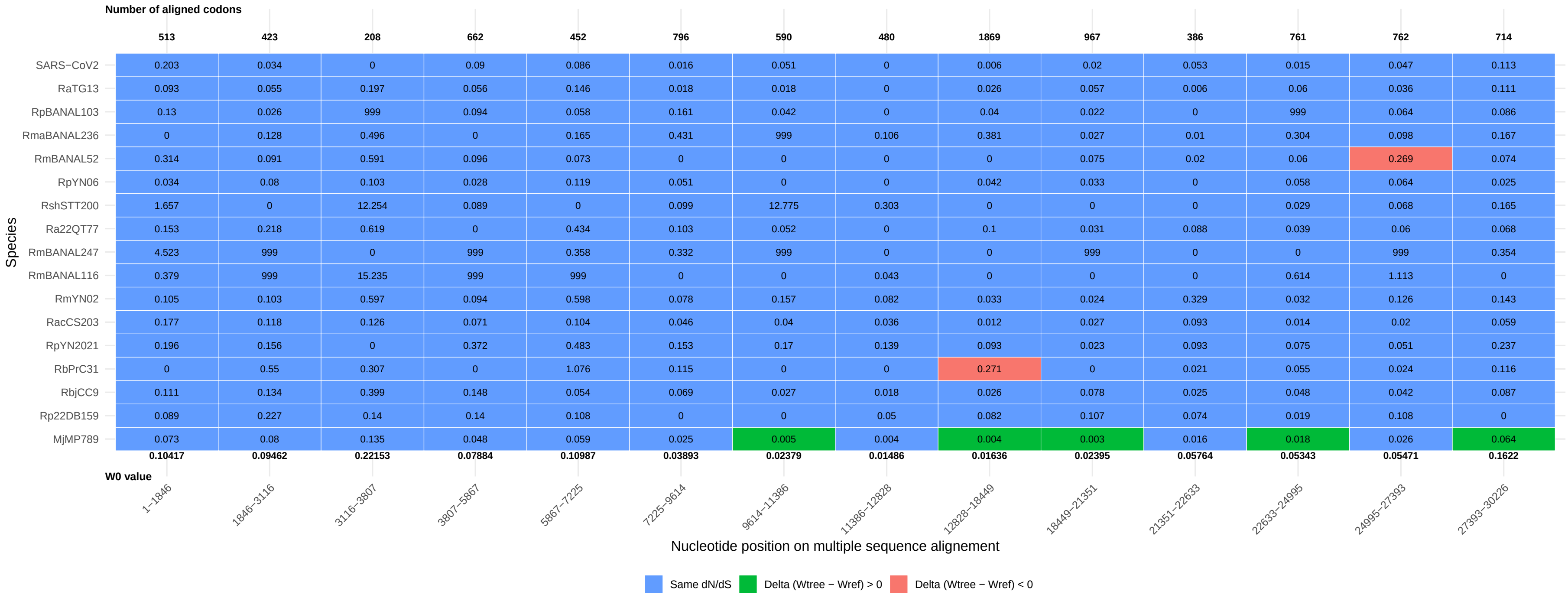
Number of aligned codons
513 423 208 662 452 796 590 480 1869 967 386 761 762 714
Species
SARS−CoV2 0.203 0.034 0 0.09 0.086 0.016 0.051 0 0.006 0.02 0.053 0.015 0.047 0.113
RaTG13 0.093 0.055 0.197 0.056 0.146 0.018 0.018 0 0.026 0.057 0.006 0.06 0.036 0.111
RpBANAL103 0.13 0.026 999 0.094 0.058 0.161 0.042 0 0.04 0.022 0 999 0.064 0.086
RmaBANAL236 0 0.128 0.496 0 0.165 0.431 999 0.106 0.381 0.027 0.01 0.304 0.098 0.167
RmBANAL52 0.314 0.091 0.591 0.096 0.073 0 0 0 0 0.075 0.02 0.06 0.269 0.074
RpYN06 0.034 0.08 0.103 0.028 0.119 0.051 0 0 0.042 0.033 0 0.058 0.064 0.025
RshSTT200 1.657 0 12.254 0.089 0 0.099 12.775 0.303 0 0 0 0.029 0.068 0.165
Ra22QT77 0.153 0.218 0.619 0 0.434 0.103 0.052 0 0.1 0.031 0.088 0.039 0.06 0.068
RmBANAL247 4.523 999 0 999 0.358 0.332 999 0 0 999 0 0 999 0.354
RmBANAL116 0.379 999 15.235 999 999 0 0 0.043 0 0 0 0.614 1.113 0
RmYN02 0.105 0.103 0.597 0.094 0.598 0.078 0.157 0.082 0.033 0.024 0.329 0.032 0.126 0.143
RacCS203 0.177 0.118 0.126 0.071 0.104 0.046 0.04 0.036 0.012 0.027 0.093 0.014 0.02 0.059
RpYN2021 0.196 0.156 0 0.372 0.483 0.153 0.17 0.139 0.093 0.023 0.093 0.075 0.051 0.237
RbPrC31 0 0.55 0.307 0 1.076 0.115 0 0 0.271 0 0.021 0.055 0.024 0.116
RbjCC9 0.111 0.134 0.399 0.148 0.054 0.069 0.027 0.018 0.026 0.078 0.025 0.048 0.042 0.087
Rp22DB159 0.089 0.227 0.14 0.14 0.108 0 0 0.05 0.082 0.107 0.074 0.019 0.108 0
MjMP789 0.073 0.08 0.135 0.048 0.059 0.025 0.005 0.004 0.004 0.003 0.016 0.018 0.026 0.064
0.10417 0.09462 0.22153 0.07884 0.10987 0.03893 0.02379 0.01486 0.01636 0.02395 0.05764 0.05343 0.05471 0.1622
W0 value
1−1846 1846−3116 3116−3807 3807−5867 5867−7225 7225−9614 9614−11386 11386−12828 12828−18449 18449−21351 21351−22633 22633−24995 24995−27393 27393−30226
Nucleotide position on multiple sequence alignement
Same dN/dS
Delta (Wtree − Wref) > 0
Delta (Wtree − Wref) < 0

(E)

Number of aligned codons

| Species | 1–1804 | 1804–2262 | 2262–3019 | 3019–3863 | 3863–5522 | 5522–7170 | 7170–9607 | 9607–11586 | 11586–12818 | 12818–15221 | 15221–18458 | 18458–21403 | 21403–22649 | 22649–23927 | 23927–24997 | 24997–25778 | 25778–27580 | 27580–28582 | 28582–30379 |
|---|---|---|---|---|---|---|---|---|---|---|---|---|---|---|---|---|---|---|---|
| Number of aligned codons | 499 | 152 | 252 | 259 | 529 | 548 | 812 | 658 | 410 | 797 | 1079 | 981 | 364 | 402 | 356 | 211 | 567 | 291 | 411 |
| CoV−2−S_edit | 0.203 | 0 | 0.094 | 0 | 0.04 | 0.089 | 0.016 | 0.051 | 0 | 0 | 0 | 0.017 | 0.01 | 0.007 | 0.005 | 0.026 | 0.051 | 0.181 | 0.071 |
| RaTG13 | 0.084 | 0.018 | 0.089 | 0.201 | 0.075 | 0.12 | 0.018 | 0.016 | 0 | 0.018 | 0.031 | 0.054 | 0 | 0.107 | 0 | 0.039 | 0.025 | 0.163 | 0.065 |
| RmBANAL52 | 0.318 | 0 | 0.131 | 0.188 | 0.114 | 0.086 | 0 | 0 | 0 | 0 | 0 | 0.075 | 0.029 | 0.079 | 0 | 0.293 | 0.223 | 0.066 | 0.096 |
| RpBANAL103 | 0.13 | 0 | 0 | 0.375 | 0.086 | 0.054 | 0.161 | 0.04 | 0 | 0 | 0.059 | 0.021 | 0 | 999 | 0.29 | 0.049 | 0 | 0 | 0.076 |
| RmaBANAL236 | 0 | 0 | 999 | 0.278 | 0 | 0.173 | 0.301 | 999 | 0.105 | 0 | 0.321 | 0.027 | 0.012 | 0.302 | 0.292 | 0.333 | 0.046 | 0.128 | 0.152 |
| RpYN06 | 0.024 | 0.081 | 0.091 | 0.089 | 0.021 | 0.118 | 0.051 | 0 | 0 | 0.015 | 0.033 | 0.031 | 0 | 0.033 | 0.044 | 0 | 0.103 | 0.094 | 0 |
| MjMP789 | 0.08 | 0.07 | 0.047 | 0.137 | 0.049 | 0.051 | 0.024 | 0.002 | 0 | 0.004 | 0.005 | 0.003 | 0.024 | 0.024 | 0.012 | 0.015 | 0.038 | 0.025 | 0.154 |
| Rp22DB159 | 0.094 | 0.867 | 0.233 | 0.113 | 0.292 | 0.102 | 0.088 | 0 | 0.061 | 0.164 | 0.066 | 0.107 | 0.048 | 0.036 | 0.018 | 0.253 | 0.086 | 0 | 0 |
| RbjCC9 | 0.088 | 0.226 | 0.139 | 0.351 | 0.117 | 0.137 | 0.069 | 0.023 | 0.022 | 0.015 | 0.034 | 0.094 | 0 | 0.094 | 0.046 | 0.163 | 0.033 | 0.108 | 0.091 |
| RpYN2021 | 0.153 | 0 | 0.394 | 0.152 | 0.304 | 0.283 | 0.156 | 0.172 | 0.134 | 0.323 | 0.024 | 0.023 | 0.099 | 0.067 | 0.624 | 0 | 0.065 | 0.221 | 0.189 |
| RbPrC31 | 0 | 0 | 0.794 | 0.314 | 0 | 1.072 | 0.117 | 0 | 0 | 0.196 | 0.673 | 0 | 0.021 | 0.07 | 0 | 0 | 0.041 | 0.271 | 0.089 |
| RshSTT200 | 999 | 49.318 | 0 | 0 | 0.039 | 0.079 | 0.101 | 0 | 0.202 | 0 | 0 | 0 | 5.392 | 0 | 0.057 | 0.085 | 0.151 | 999 | 0 |
| Ra22QT77 | 0.137 | 0.111 | 0.386 | 0.633 | 0 | 0.495 | 0.103 | 0.049 | 0.088 | 0.255 | 0.044 | 0.031 | 0.09 | 0.068 | 0.017 | 0.163 | 0.058 | 0 | 0.136 |
| RacCS203 | 0.177 | 0.196 | 0.119 | 0.137 | 0.047 | 0.079 | 0.057 | 0.035 | 0.019 | 0.026 | 0.01 | 0.026 | 0.1 | 0.048 | 0.006 | 0.045 | 0.027 | 0.06 | 0.051 |
| RmYN02 | 0.071 | 0.133 | 0.117 | 0.304 | 0.098 | 0.72 | 0.091 | 0.133 | 0.08 | 0.023 | 0.021 | 0.024 | 0.354 | 0.052 | 0 | 0.128 | 0.376 | 0.321 | 0.119 |
| RmBANAL247 | 0.069 | 999 | 999 | 4.711 | 999 | 0.359 | 0.331 | 999 | 0 | 0 | 0 | 999 | 0 | 0 | 0 | 999 | 999 | 999 | 0.192 |
| RmBANAL116 | 0.383 | 0 | 0 | 999 | 999 | 999 | 6.351 | 0 | 0 | 0 | 0 | 0 | 0 | 999 | 0.305 | 0 | 999 | 0 | 0.465 |
| W0 value | 0.10156 | 0.08678 | 0.10375 | 0.19297 | 0.07036 | 0.10573 | 0.04115 | 0.02201 | 0.01441 | 0.02276 | 0.01017 | 0.02409 | 0.03781 | 0.04417 | 0.01948 | 0.0453 | 0.0599 | 0.19655 | 0.09651 |

Species

Nucleotide position on multiple sequence alignement

Same dN/dS | Delta (Wtree − Wref) < 0 | Delta (Wtree − Wref) > 0

(F)

| Species | 1–833 | 833–3152 | 3152–4483 | 4483–6519 | 6519–9695 | 9695–11713 | 11713–15269 | 15269–18433 | 18433–21004 | 21004–21504 | 21504–22367 | 22367–23210 | 23210–25133 | 25133–26607 | 26607–28247 | 28247–28832 | 28832–30434 |
|---|---|---|---|---|---|---|---|---|---|---|---|---|---|---|---|---|---|
| **Number of aligned codons** | **189** | **771** | **439** | **486** | **1058** | **672** | **1119** | **1054** | **856** | **161** | **252** | **261** | **640** | **446** | **326** | **170** | **412** |
| WIV−BANAL−20−236_opt | 2.529 | 0 | 0 | 0.218 | 0 | 0 | 16.831 | 0 | 3.928 | 0.329 | 0.008 | 0.002 | 0.021 | 0.095 | 0 | 0 | 0.333 |
| RsYN2013 | 0.211 | 0.127 | 0.314 | 0.209 | 0.094 | 0 | 0.099 | 0 | 0.069 | 0.032 | 0.056 | 0.053 | 0.11 | 0.055 | 0.231 | 0.089 | 0.173 |
| RsWIV1 | 23.087 | 0 | 0 | 1.775 | 8.081 | 0 | 0 | 0.47 | 0 | 0.115 | 1.979 | 0 | 999 | 0.429 | 0 | 2 | 0.45 |
| RsGX2013 | 0.115 | 0.106 | 0.185 | 0.072 | 0.151 | 0.028 | 0.017 | 0.007 | 0 | 0.014 | 0.031 | 0.039 | 0.019 | 0.094 | 0.083 | 0.09 | 0.117 |
| Rs9401 | 0.502 | 0.13 | 0.376 | 0.057 | 707.013 | 0 | 0.112 | 999 | 0 | 0 | 10.394 | 0 | 0 | 0.909 | 0.389 | 999 | 0 |
| Rs7327 | 0 | 0.395 | 0 | 0 | 0.04 | 0 | 0.408 | 999 | 0 | 0.217 | 0 | 0 | 0 | 0.627 | 0.194 | 0.068 | 0.154 |
| Rs672 | 0.412 | 0.132 | 0.086 | 0.12 | 0.11 | 0.043 | 0 | 0.049 | 0.12 | 0 | 999 | 0 | 0.022 | 0.169 | 0 | 0.12 | 0.057 |
| Rs4874 | 0 | 0.35 | 999 | 999 | 0.101 | 0.398 | 0.135 | 2.53 | 0.509 | 0 | 0 | 0 | 0 | 0.466 | 0.194 | 0 | 0.225 |
| Rs4255 | 23.859 | 0.294 | 0.757 | 0 | 1.132 | 0 | 0.37 | 0 | 0.025 | 0 | 0.281 | 0 | 0.064 | 0.038 | 0.075 | 0 | 0.15 |
| Rs4247 | 0.278 | 0.127 | 0 | 999 | 0.042 | 0 | 0 | 0 | 0 | 0.233 | 0 | 0.009 | 0.008 | 0.79 | 0.409 | 999 | 0 |
| Rs4237 | 0.437 | 0.257 | 0 | 0 | 0.038 | 0.22 | 0.121 | 0.045 | 0 | 0 | 0 | 0 | 0.088 | 0.044 | 0 | 0 | 0 |
| Rs4231 | 0 | 0.151 | 0.193 | 0 | 0.197 | 0 | 0.04 | 0.049 | 0.047 | 0.239 | 0 | 0.943 | 0.027 | 0.205 | 0.583 | 28.911 | 0.452 |
| Rs4081 | 0.214 | 0 | 1.158 | 0 | 205.863 | 0.027 | 5.115 | 0.136 | 0.021 | 0 | 1.005 | 0.274 | 0 | 0.074 | 0 | 0 | 0.149 |
| Rs3367 | 0.458 | 999 | 0 | 0 | 999 | 3.03 | 0 | 0 | 0.776 | 0 | 2.353 | 0 | 256.582 | 0 | 0 | 0 | 0 |
| RfYNLF_34C | 0.681 | 999 | 0.377 | 999 | 999 | 0 | 0.67 | 1.125 | 999 | 0 | 0 | 6.633 | 18.288 | 999 | 0 | 0 | 999 |
| RfYNLF_31C | 0.429 | 0.465 | 0 | 0 | 999 | 0 | 19.568 | 0 | 999 | 0.794 | 0 | 14.542 | 0 | 0.455 | 0 | 4.421 | 50.526 |
| Rf4092 | 15.411 | 0.203 | 0.193 | 0.123 | 0.086 | 0 | 0.062 | 0.015 | 0.032 | 0.06 | 0.103 | 0.042 | 0.056 | 0.106 | 0 | 0.092 | 0.358 |
| bRp3 | 0.286 | 0.12 | 0.348 | 0.127 | 0.102 | 0.046 | 0.076 | 0.037 | 0.199 | 0.024 | 0.023 | 0.011 | 0.007 | 0.097 | 0.071 | 0.067 | 0.088 |
| As6526 | 0.536 | 0 | 0 | 0 | 0.187 | 0.396 | 999 | 0 | 0 | 0 | 0.02 | 0.037 | 0.013 | 0.052 | 0 | 0 | 0.887 |
| **W0 value** | **0.20498** | **0.12942** | **0.21076** | **0.0703** | **0.08647** | **0.03182** | **0.05573** | **0.02321** | **0.03845** | **0.03785** | **0.05782** | **0.06373** | **0.03452** | **0.12856** | **0.09454** | **0.17981** | **0.1276** |

Nucleotide position on multiple sequence alignement

Same dN/dS
Delta (Wtree − Wref) < 0
Delta (Wtree − Wref) > 0

(G)

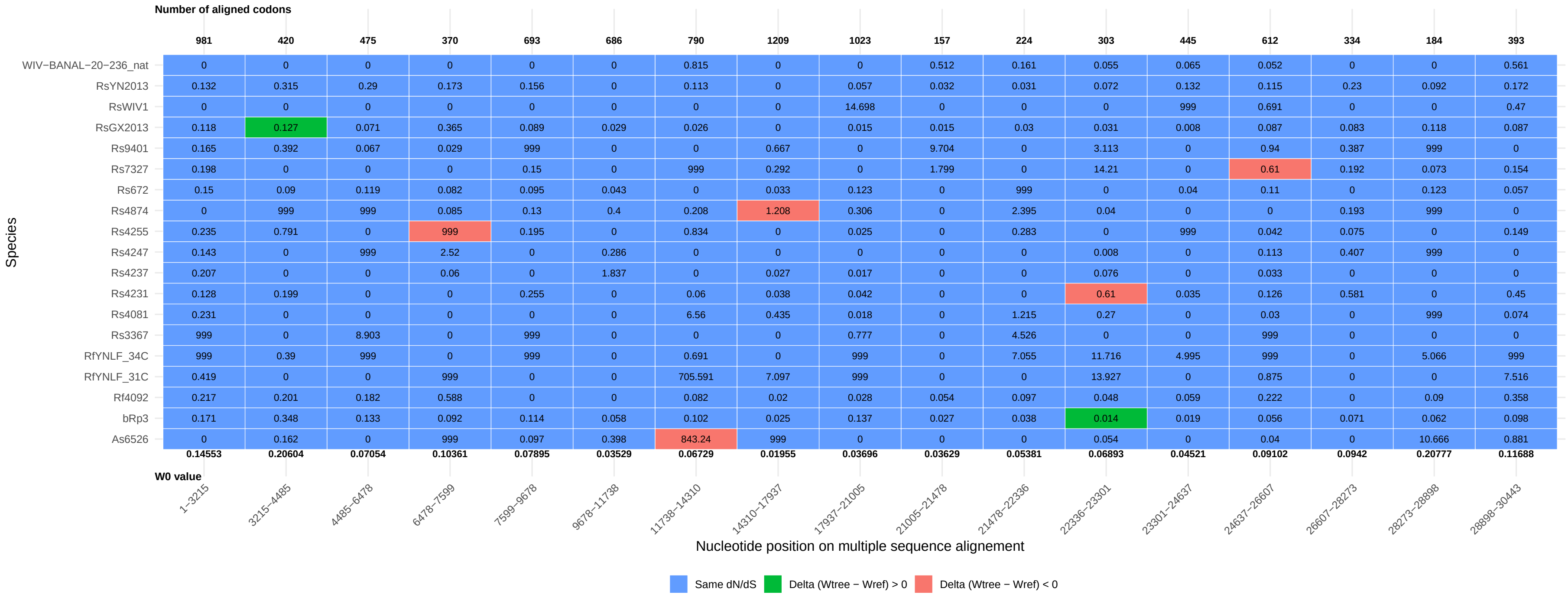
Number of aligned codons
981 420 475 370 693 686 790 1209 1023 157 224 303 445 612 334 184 393
WIV−BANAL−20−236_nat 0 0 0 0 0 0 0.815 0 0 0.512 0.161 0.055 0.065 0.052 0 0 0.561
RsYN2013 0.132 0.315 0.29 0.173 0.156 0 0.113 0 0.057 0.032 0.031 0.072 0.132 0.115 0.23 0.092 0.172
RsWIV1 0 0 0 0 0 0 0 0 14.698 0 0 0 999 0.691 0 0 0.47
RsGX2013 0.118 0.127 0.071 0.365 0.089 0.029 0.026 0 0.015 0.015 0.03 0.031 0.008 0.087 0.083 0.118 0.087
Rs9401 0.165 0.392 0.067 0.029 999 0 0 0.667 0 9.704 0 3.113 0 0.94 0.387 999 0
Rs7327 0.198 0 0 0 0.15 0 999 0.292 0 1.799 0 14.21 0 0.61 0.192 0.073 0.154
Rs672 0.15 0.09 0.119 0.082 0.095 0.043 0 0.033 0.123 0 999 0 0.04 0.11 0 0.123 0.057
Rs4874 0 999 999 0.085 0.13 0.4 0.208 1.208 0.306 0 2.395 0.04 0 0 0.193 999 0
Rs4255 0.235 0.791 0 999 0.195 0 0.834 0 0.025 0 0.283 0 999 0.042 0.075 0 0.149
Rs4247 0.143 0 999 2.52 0 0.286 0 0 0 0 0 0.008 0 0.113 0.407 999 0
Rs4237 0.207 0 0 0.06 0 1.837 0 0.027 0.017 0 0 0.076 0 0.033 0 0 0
Rs4231 0.128 0.199 0 0 0.255 0 0.06 0.038 0.042 0 0 0.61 0.035 0.126 0.581 0 0.45
Rs4081 0.231 0 0 0 0 0 6.56 0.435 0.018 0 1.215 0.27 0 0.03 0 999 0.074
Rs3367 999 0 8.903 0 999 0 0 0 0.777 0 4.526 0 0 999 0 0 0
RfYNLF_34C 999 0.39 999 0 999 0 0.691 0 999 0 7.055 11.716 4.995 999 0 5.066 999
RfYNLF_31C 0.419 0 0 999 0 0 705.591 7.097 999 0 0 13.927 0 0.875 0 0 7.516
Rf4092 0.217 0.201 0.182 0.588 0 0 0.082 0.02 0.028 0.054 0.097 0.048 0.059 0.222 0 0.09 0.358
bRp3 0.171 0.348 0.133 0.092 0.114 0.058 0.102 0.025 0.137 0.027 0.038 0.014 0.019 0.056 0.071 0.062 0.098
As6526 0 0.162 0 999 0.097 0.398 843.24 999 0 0 0 0.054 0 0.04 0 10.666 0.881
0.14553 0.20604 0.07054 0.10361 0.07895 0.03529 0.06729 0.01955 0.03696 0.03629 0.05381 0.06893 0.04521 0.09102 0.0942 0.20777 0.11688
W0 value
1−3215 3215−4485 4485−6478 6478−7599 7599−9678 9678−11738 11738−14310 14310−17937 17937−21005 21005−21478 21478−22336 22336−23301 23301−24637 24637−26607 26607−28273 28273−28898 28898−30443
Species
Nucleotide position on multiple sequence alignement
Same dN/dS
Delta (Wtree − Wref) > 0
Delta (Wtree − Wref) < 0

Figure S15. Graphical representation of the d$N$/d$S$ ratio across all the segments of the three datasets. The average value of the local tree is represented by triangles connected by a straight line. dN/dS ratio inferred for each external branch is represented by a circle (or a red disk for the virus of interest) or a star, if the value significantly differ from the rest of the phylogeny. When the number of Non-synonymous and/or of Synonymous mutations in the external branch equals 0, the value is reported in one of the three grey bars. (A) SARS-CoV-1, (B) MERS and (C) SARS-CoV-2.

(A)

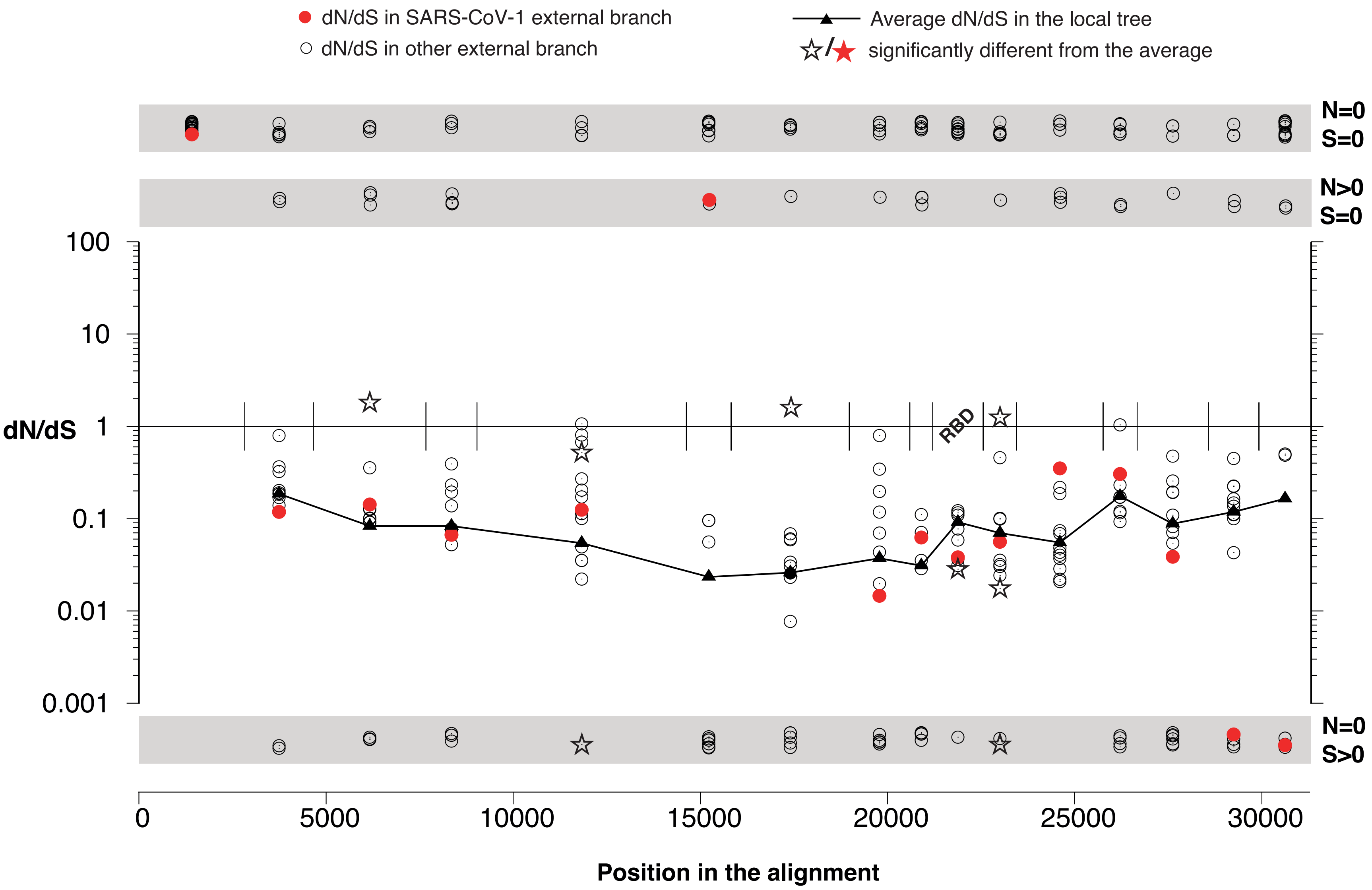
dN/dS in SARS-CoV-1 external branch
dN/dS in other external branch
Average dN/dS in the local tree
significantly different from the average
N=0
S=0
N>0
S=0
100
10
dN/dS 1
0.1
0.01
0.001
RBD
N=0
S>0
0
5000
10000
15000
20000
25000
30000
Position in the alignment

(B)

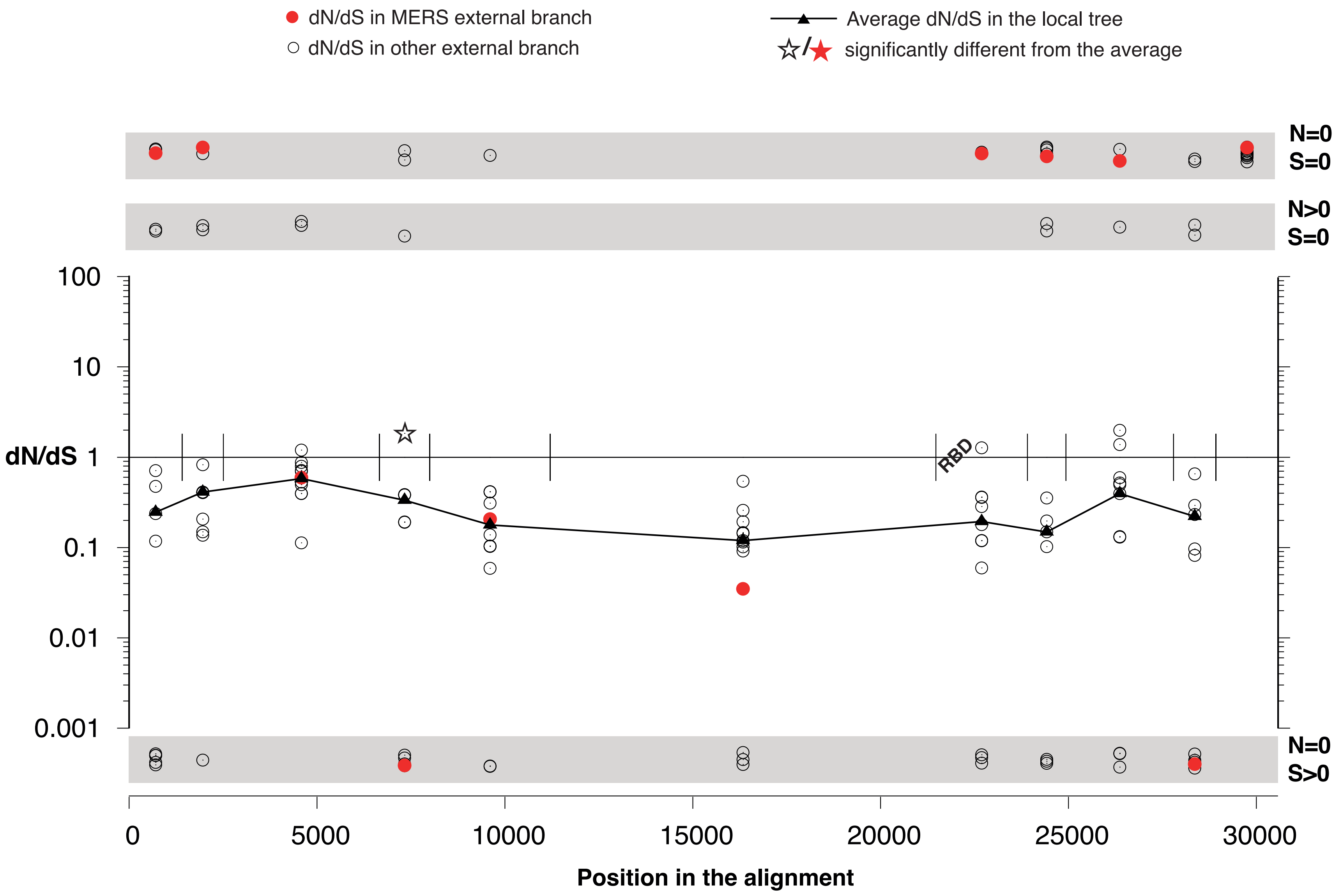

dN/dS in MERS external branch
dN/dS in other external branch
Average dN/dS in the local tree
significantly different from the average
N=0
S=0
N>0
S=0
100
10
dN/dS 1
0.1
0.01
0.001
RBD
N=0
S>0
0
5000
10000
15000
20000
25000
30000
Position in the alignment

(C)

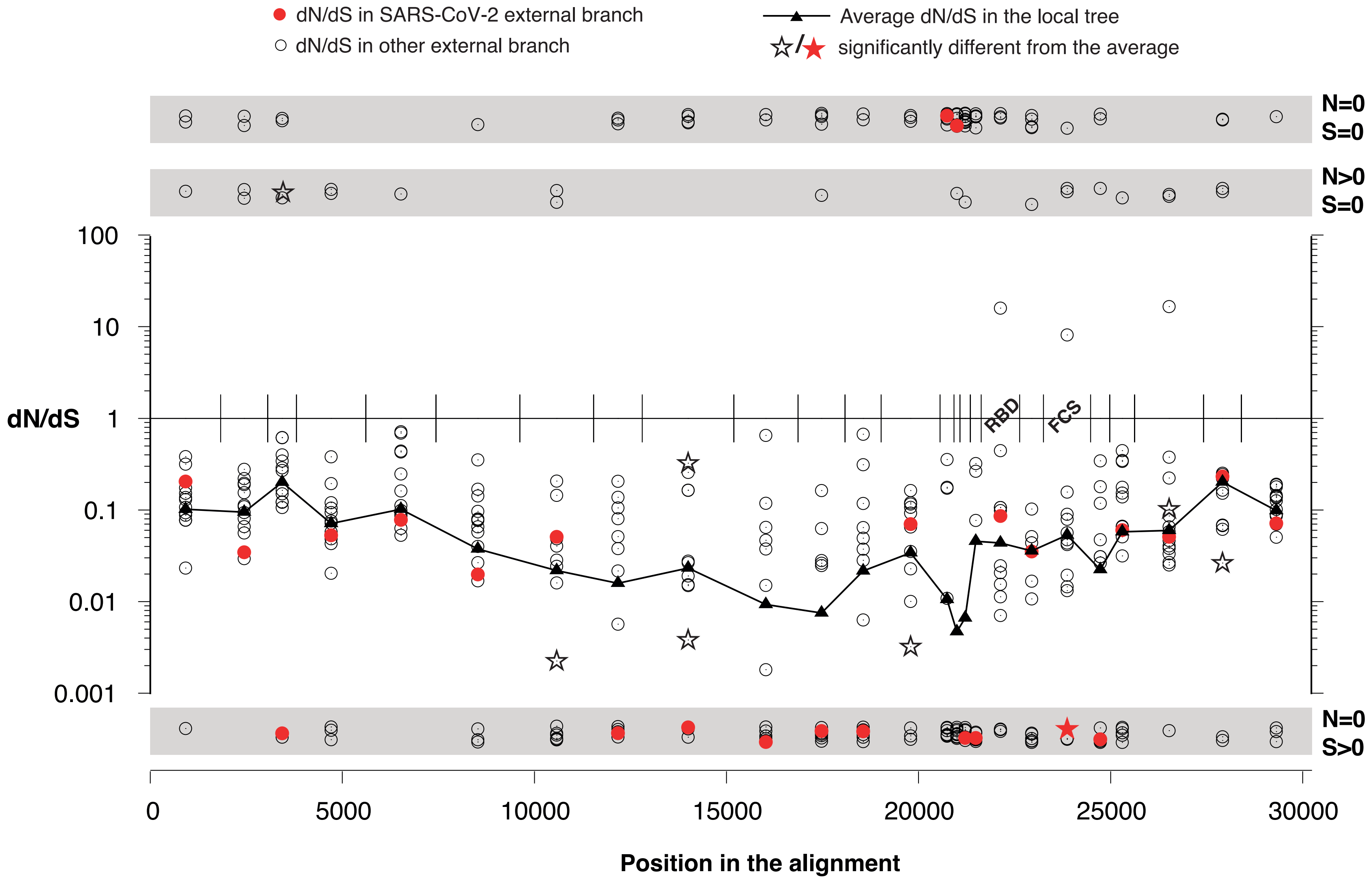
dN/dS in SARS-CoV-2 external branch
dN/dS in other external branch
Average dN/dS in the local tree
significantly different from the average
N=0
S=0
N>0
S=0
N=0
S>0
100
10
dN/dS
1
0.1
0.01
0.001
RBD
FCS
0
5000
10000
15000
20000
25000
30000
Position in the alignment

Figure S16. Estimation of Codon Adaptation Index (CAI) across all the segments of the three datasets. (A) SARS-CoV-1, (B) MERS and (C) SARS-CoV-2, (D) SARS-CoV-2 with only 14 segments (larger segments), (E) CoV-2-S$^{edit}$, (F) WIV-BANAL-20-236$^{opt}$, and (G) WIV-BANAL-20-236$^{nat}$.

(A)

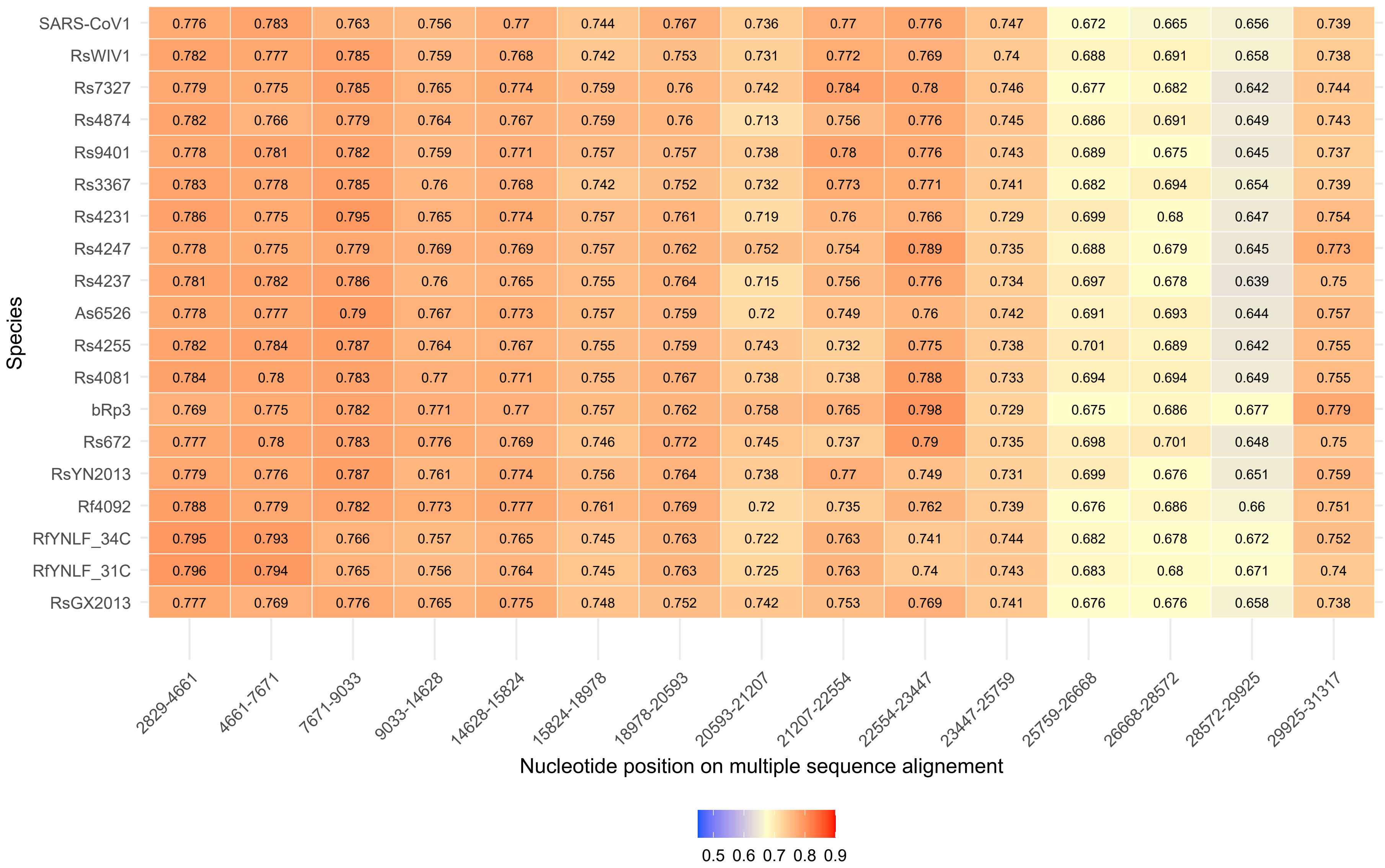

Species
SARS-CoV1 0.776 0.783 0.763 0.756 0.77 0.744 0.767 0.736 0.77 0.776 0.747 0.672 0.665 0.656 0.739
RsWIV1 0.782 0.777 0.785 0.759 0.768 0.742 0.753 0.731 0.772 0.769 0.74 0.688 0.691 0.658 0.738
Rs7327 0.779 0.775 0.785 0.765 0.774 0.759 0.76 0.742 0.784 0.78 0.746 0.677 0.682 0.642 0.744
Rs4874 0.782 0.766 0.779 0.764 0.767 0.759 0.76 0.713 0.756 0.776 0.745 0.686 0.691 0.649 0.743
Rs9401 0.778 0.781 0.782 0.759 0.771 0.757 0.757 0.738 0.78 0.776 0.743 0.689 0.675 0.645 0.737
Rs3367 0.783 0.778 0.785 0.76 0.768 0.742 0.752 0.732 0.773 0.771 0.741 0.682 0.694 0.654 0.739
Rs4231 0.786 0.775 0.795 0.765 0.774 0.757 0.761 0.719 0.76 0.766 0.729 0.699 0.68 0.647 0.754
Rs4247 0.778 0.775 0.779 0.769 0.769 0.757 0.762 0.752 0.754 0.789 0.735 0.688 0.679 0.645 0.773
Rs4237 0.781 0.782 0.786 0.76 0.765 0.755 0.764 0.715 0.756 0.776 0.734 0.697 0.678 0.639 0.75
As6526 0.778 0.777 0.79 0.767 0.773 0.757 0.759 0.72 0.749 0.76 0.742 0.691 0.693 0.644 0.757
Rs4255 0.782 0.784 0.787 0.764 0.767 0.755 0.759 0.743 0.732 0.775 0.738 0.701 0.689 0.642 0.755
Rs4081 0.784 0.78 0.783 0.77 0.771 0.755 0.767 0.738 0.738 0.788 0.733 0.694 0.694 0.649 0.755
bRp3 0.769 0.775 0.782 0.771 0.77 0.757 0.762 0.758 0.765 0.798 0.729 0.675 0.686 0.677 0.779
Rs672 0.777 0.78 0.783 0.776 0.769 0.746 0.772 0.745 0.737 0.79 0.735 0.698 0.701 0.648 0.75
RsYN2013 0.779 0.776 0.787 0.761 0.774 0.756 0.764 0.738 0.77 0.749 0.731 0.699 0.676 0.651 0.759
Rf4092 0.788 0.779 0.782 0.773 0.777 0.761 0.769 0.72 0.735 0.762 0.739 0.676 0.686 0.66 0.751
RfYNLF_34C 0.795 0.793 0.766 0.757 0.765 0.745 0.763 0.722 0.763 0.741 0.744 0.682 0.678 0.672 0.752
RfYNLF_31C 0.796 0.794 0.765 0.756 0.764 0.745 0.763 0.725 0.763 0.74 0.743 0.683 0.68 0.671 0.74
RsGX2013 0.777 0.769 0.776 0.765 0.775 0.748 0.752 0.742 0.753 0.769 0.741 0.676 0.676 0.658 0.738
2829-4661
4661-7671
7671-9033
9033-14628
14628-15824
15824-18978
18978-20593
20593-21207
21207-22554
22554-23447
23447-25759
25759-26668
26668-28572
28572-29925
29925-31317
Nucleotide position on multiple sequence alignement
0.5 0.6 0.7 0.8 0.9

(B)

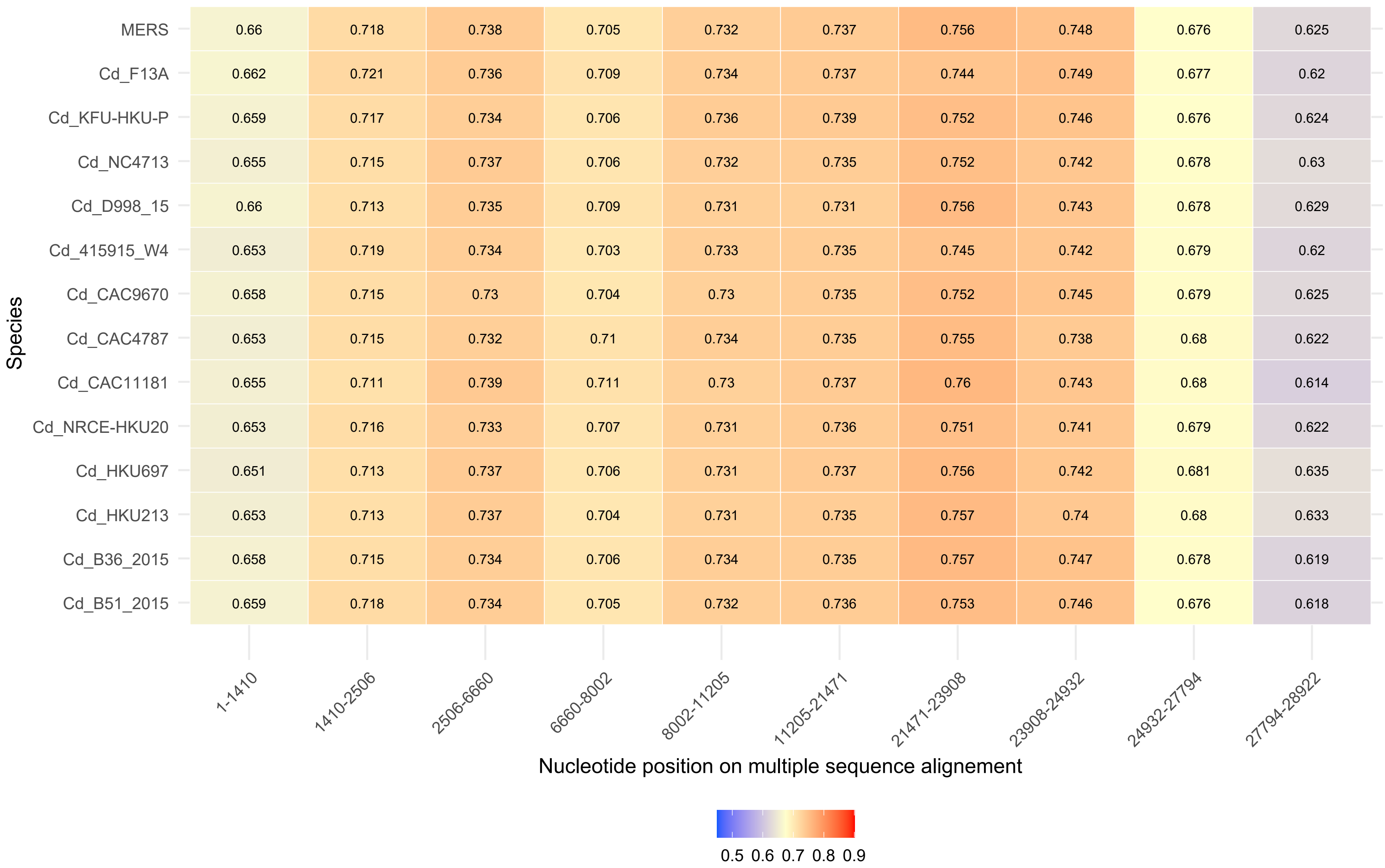

Species
MERS 0.66 0.718 0.738 0.705 0.732 0.737 0.756 0.748 0.676 0.625
Cd_F13A 0.662 0.721 0.736 0.709 0.734 0.737 0.744 0.749 0.677 0.62
Cd_KFU-HKU-P 0.659 0.717 0.734 0.706 0.736 0.739 0.752 0.746 0.676 0.624
Cd_NC4713 0.655 0.715 0.737 0.706 0.732 0.735 0.752 0.742 0.678 0.63
Cd_D998_15 0.66 0.713 0.735 0.709 0.731 0.731 0.756 0.743 0.678 0.629
Cd_415915_W4 0.653 0.719 0.734 0.703 0.733 0.735 0.745 0.742 0.679 0.62
Cd_CAC9670 0.658 0.715 0.73 0.704 0.73 0.735 0.752 0.745 0.679 0.625
Cd_CAC4787 0.653 0.715 0.732 0.71 0.734 0.735 0.755 0.738 0.68 0.622
Cd_CAC11181 0.655 0.711 0.739 0.711 0.73 0.737 0.76 0.743 0.68 0.614
Cd_NRCE-HKU20 0.653 0.716 0.733 0.707 0.731 0.736 0.751 0.741 0.679 0.622
Cd_HKU697 0.651 0.713 0.737 0.706 0.731 0.737 0.756 0.742 0.681 0.635
Cd_HKU213 0.653 0.713 0.737 0.704 0.731 0.735 0.757 0.74 0.68 0.633
Cd_B36_2015 0.658 0.715 0.734 0.706 0.734 0.735 0.757 0.747 0.678 0.619
Cd_B51_2015 0.659 0.718 0.734 0.705 0.732 0.736 0.753 0.746 0.676 0.618
1-1410
1410-2506
2506-6660
6660-8002
8002-11205
11205-21471
21471-23908
23908-24932
24932-27794
27794-28922
Nucleotide position on multiple sequence alignement
0.5 0.6 0.7 0.8 0.9

(C)

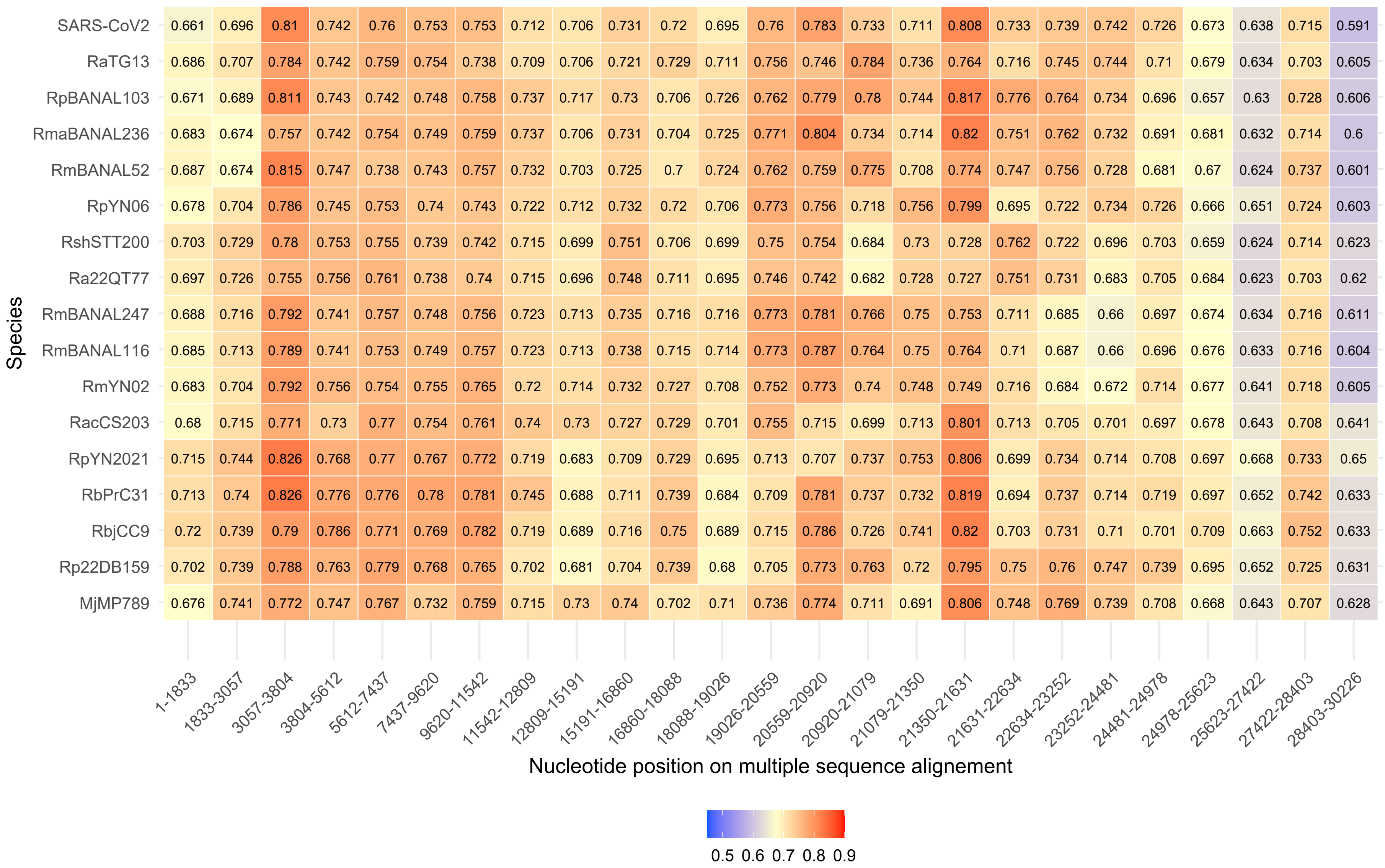
SARS-CoV2 0.661 0.696 0.81 0.742 0.76 0.753 0.753 0.712 0.706 0.731 0.72 0.695 0.76 0.783 0.733 0.711 0.808 0.733 0.739 0.742 0.726 0.673 0.638 0.715 0.591
RaTG13 0.686 0.707 0.784 0.742 0.759 0.754 0.738 0.709 0.706 0.721 0.729 0.711 0.756 0.746 0.784 0.736 0.764 0.716 0.745 0.744 0.71 0.679 0.634 0.703 0.605
RpBANAL103 0.671 0.689 0.811 0.743 0.742 0.748 0.758 0.737 0.717 0.73 0.706 0.726 0.762 0.779 0.78 0.744 0.817 0.776 0.764 0.734 0.696 0.657 0.63 0.728 0.606
RmaBANAL236 0.683 0.674 0.757 0.742 0.754 0.749 0.759 0.737 0.706 0.731 0.704 0.725 0.771 0.804 0.734 0.714 0.82 0.751 0.762 0.732 0.691 0.681 0.632 0.714 0.6
RmBANAL52 0.687 0.674 0.815 0.747 0.738 0.743 0.757 0.732 0.703 0.725 0.7 0.724 0.762 0.759 0.775 0.708 0.774 0.747 0.756 0.728 0.681 0.67 0.624 0.737 0.601
RpYN06 0.678 0.704 0.786 0.745 0.753 0.74 0.743 0.722 0.712 0.732 0.72 0.706 0.773 0.756 0.718 0.756 0.799 0.695 0.722 0.734 0.726 0.666 0.651 0.724 0.603
RshSTT200 0.703 0.729 0.78 0.753 0.755 0.739 0.742 0.715 0.699 0.751 0.706 0.699 0.75 0.754 0.684 0.73 0.728 0.762 0.722 0.696 0.703 0.659 0.624 0.714 0.623
Ra22QT77 0.697 0.726 0.755 0.756 0.761 0.738 0.74 0.715 0.696 0.748 0.711 0.695 0.746 0.742 0.682 0.728 0.727 0.751 0.731 0.683 0.705 0.684 0.623 0.703 0.62
RmBANAL247 0.688 0.716 0.792 0.741 0.757 0.748 0.756 0.723 0.713 0.735 0.716 0.716 0.773 0.781 0.766 0.75 0.753 0.711 0.685 0.66 0.697 0.674 0.634 0.716 0.611
RmBANAL116 0.685 0.713 0.789 0.741 0.753 0.749 0.757 0.723 0.713 0.738 0.715 0.714 0.773 0.787 0.764 0.75 0.764 0.71 0.687 0.66 0.696 0.676 0.633 0.716 0.604
RmYN02 0.683 0.704 0.792 0.756 0.754 0.755 0.765 0.72 0.714 0.732 0.727 0.708 0.752 0.773 0.74 0.748 0.749 0.716 0.684 0.672 0.714 0.677 0.641 0.718 0.605
RacCS203 0.68 0.715 0.771 0.73 0.77 0.754 0.761 0.74 0.73 0.727 0.729 0.701 0.755 0.715 0.699 0.713 0.801 0.713 0.705 0.701 0.697 0.678 0.643 0.708 0.641
RpYN2021 0.715 0.744 0.826 0.768 0.77 0.767 0.772 0.719 0.683 0.709 0.729 0.695 0.713 0.707 0.737 0.753 0.806 0.699 0.734 0.714 0.708 0.697 0.668 0.733 0.65
RbPrC31 0.713 0.74 0.826 0.776 0.776 0.78 0.781 0.745 0.688 0.711 0.739 0.684 0.709 0.781 0.737 0.732 0.819 0.694 0.737 0.714 0.719 0.697 0.652 0.742 0.633
RbjCC9 0.72 0.739 0.79 0.786 0.771 0.769 0.782 0.719 0.689 0.716 0.75 0.689 0.715 0.786 0.726 0.741 0.82 0.703 0.731 0.71 0.701 0.709 0.663 0.752 0.633
Rp22DB159 0.702 0.739 0.788 0.763 0.779 0.768 0.765 0.702 0.681 0.704 0.739 0.68 0.705 0.773 0.763 0.72 0.795 0.75 0.76 0.747 0.739 0.695 0.652 0.725 0.631
MjMP789 0.676 0.741 0.772 0.747 0.767 0.732 0.759 0.715 0.73 0.74 0.702 0.71 0.736 0.774 0.711 0.691 0.806 0.748 0.769 0.739 0.708 0.668 0.643 0.707 0.628
Species
1-1833 1833-3057 3057-3804 3804-5612 5612-7437 7437-9620 9620-11542 11542-12809 12809-15191 15191-16860 16860-18088 18088-19026 19026-20559 20559-20920 20920-21079 21079-21350 21350-21631 21631-22634 22634-23252 23252-24481 24481-24978 24978-25623 25623-27422 27422-28403 28403-30226
Nucleotide position on multiple sequence alignement
0.5 0.6 0.7 0.8 0.9

(D)

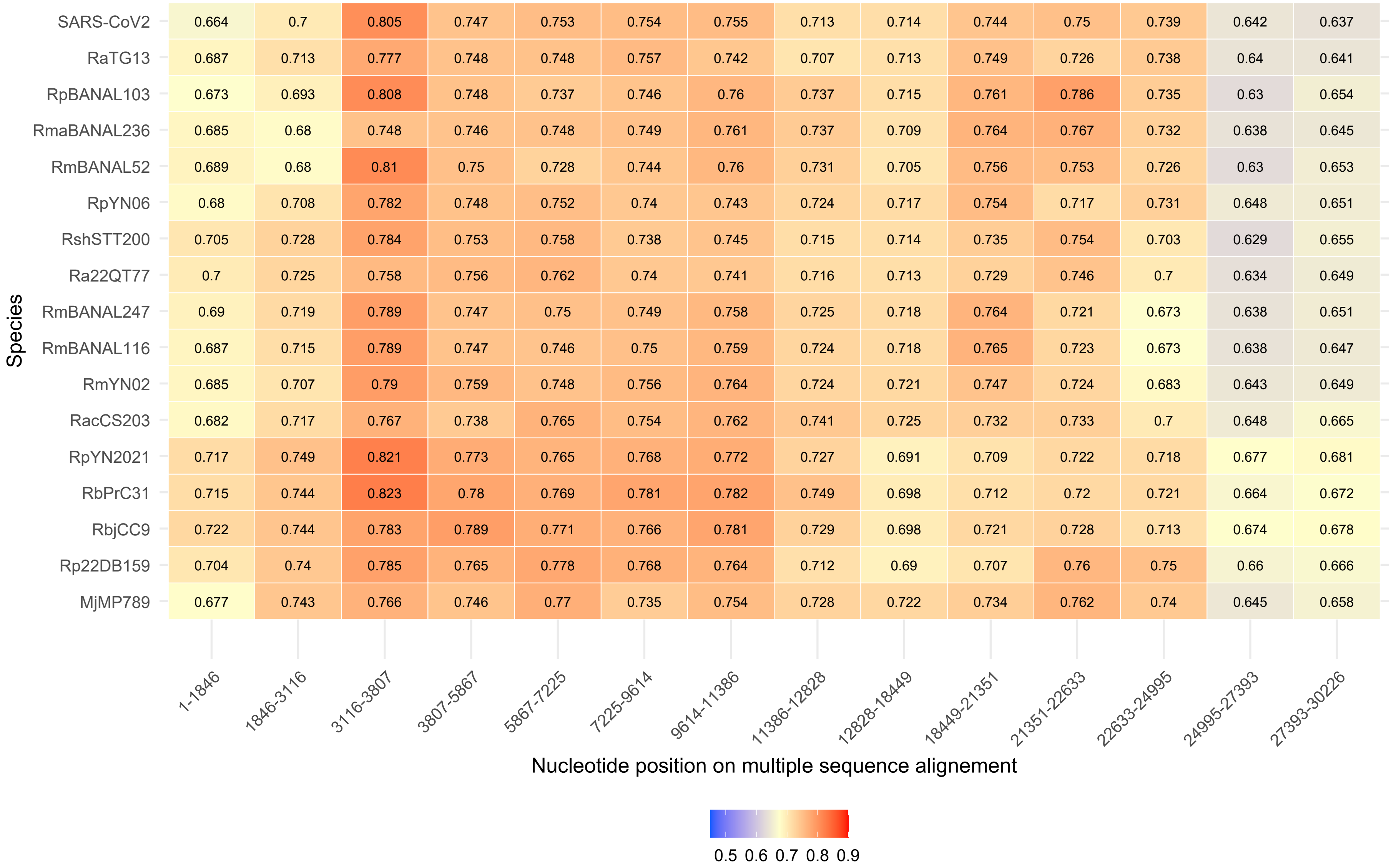
Species
SARS-CoV2 0.664 0.7 0.805 0.747 0.753 0.754 0.755 0.713 0.714 0.744 0.75 0.739 0.642 0.637
RaTG13 0.687 0.713 0.777 0.748 0.748 0.757 0.742 0.707 0.713 0.749 0.726 0.738 0.64 0.641
RpBANAL103 0.673 0.693 0.808 0.748 0.737 0.746 0.76 0.737 0.715 0.761 0.786 0.735 0.63 0.654
RmaBANAL236 0.685 0.68 0.748 0.746 0.748 0.749 0.761 0.737 0.709 0.764 0.767 0.732 0.638 0.645
RmBANAL52 0.689 0.68 0.81 0.75 0.728 0.744 0.76 0.731 0.705 0.756 0.753 0.726 0.63 0.653
RpYN06 0.68 0.708 0.782 0.748 0.752 0.74 0.743 0.724 0.717 0.754 0.717 0.731 0.648 0.651
RshSTT200 0.705 0.728 0.784 0.753 0.758 0.738 0.745 0.715 0.714 0.735 0.754 0.703 0.629 0.655
Ra22QT77 0.7 0.725 0.758 0.756 0.762 0.74 0.741 0.716 0.713 0.729 0.746 0.7 0.634 0.649
RmBANAL247 0.69 0.719 0.789 0.747 0.75 0.749 0.758 0.725 0.718 0.764 0.721 0.673 0.638 0.651
RmBANAL116 0.687 0.715 0.789 0.747 0.746 0.75 0.759 0.724 0.718 0.765 0.723 0.673 0.638 0.647
RmYN02 0.685 0.707 0.79 0.759 0.748 0.756 0.764 0.724 0.721 0.747 0.724 0.683 0.643 0.649
RacCS203 0.682 0.717 0.767 0.738 0.765 0.754 0.762 0.741 0.725 0.732 0.733 0.7 0.648 0.665
RpYN2021 0.717 0.749 0.821 0.773 0.765 0.768 0.772 0.727 0.691 0.709 0.722 0.718 0.677 0.681
RbPrC31 0.715 0.744 0.823 0.78 0.769 0.781 0.782 0.749 0.698 0.712 0.72 0.721 0.664 0.672
RbjCC9 0.722 0.744 0.783 0.789 0.771 0.766 0.781 0.729 0.698 0.721 0.728 0.713 0.674 0.678
Rp22DB159 0.704 0.74 0.785 0.765 0.778 0.768 0.764 0.712 0.69 0.707 0.76 0.75 0.66 0.666
MjMP789 0.677 0.743 0.766 0.746 0.77 0.735 0.754 0.728 0.722 0.734 0.762 0.74 0.645 0.658
1-1846
1846-3116
3116-3807
3807-5867
5867-7225
7225-9614
9614-11386
11386-12828
12828-18449
18449-21351
21351-22633
22633-24995
24995-27393
27393-30226
Nucleotide position on multiple sequence alignement
0.5 0.6 0.7 0.8 0.9

(E)

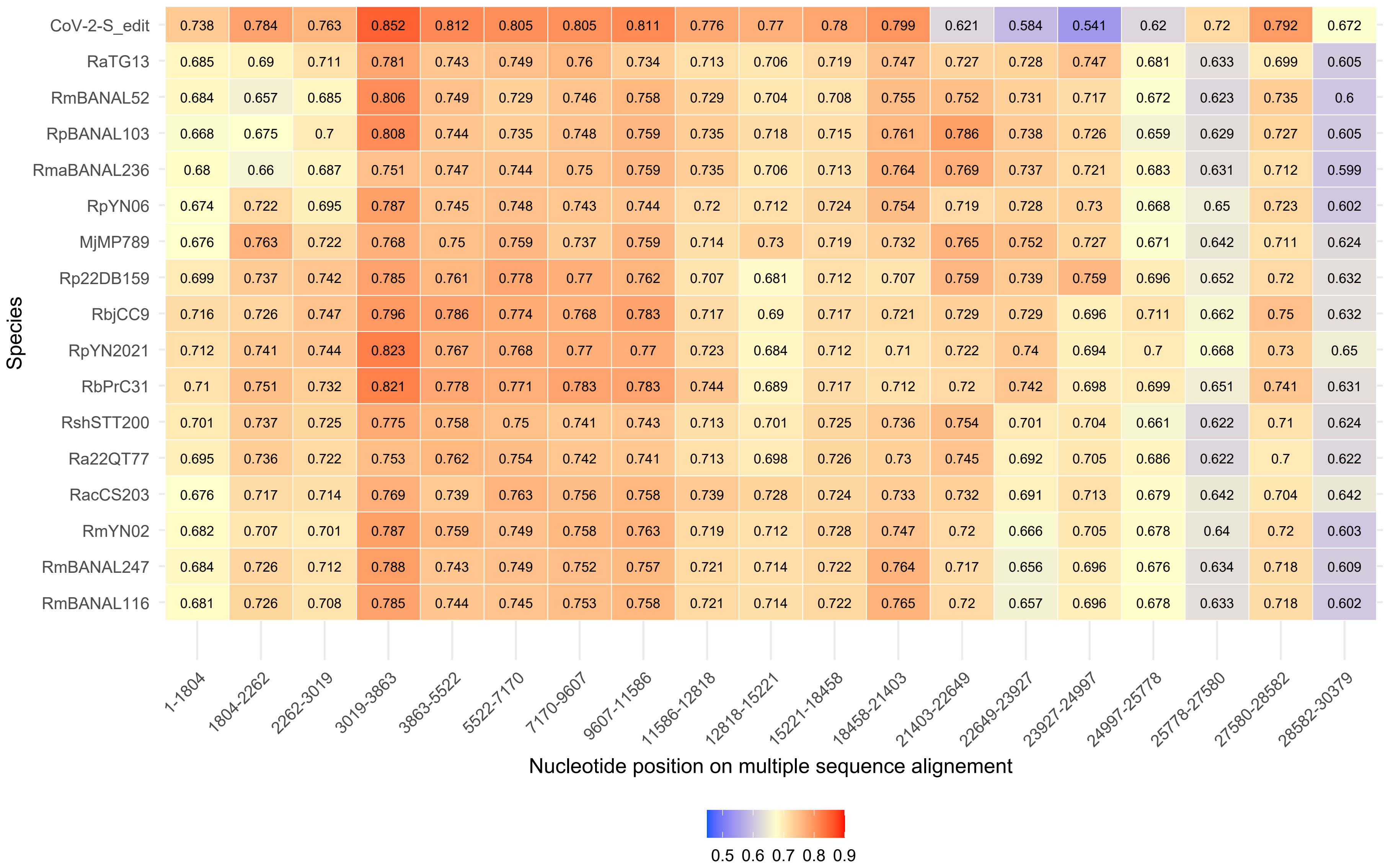
CoV-2-S_edit 0.738 0.784 0.763 0.852 0.812 0.805 0.805 0.811 0.776 0.77 0.78 0.799 0.621 0.584 0.541 0.62 0.72 0.792 0.672
RaTG13 0.685 0.69 0.711 0.781 0.743 0.749 0.76 0.734 0.713 0.706 0.719 0.747 0.727 0.728 0.747 0.681 0.633 0.699 0.605
RmBANAL52 0.684 0.657 0.685 0.806 0.749 0.729 0.746 0.758 0.729 0.704 0.708 0.755 0.752 0.731 0.717 0.672 0.623 0.735 0.6
RpBANAL103 0.668 0.675 0.7 0.808 0.744 0.735 0.748 0.759 0.735 0.718 0.715 0.761 0.786 0.738 0.726 0.659 0.629 0.727 0.605
RmaBANAL236 0.68 0.66 0.687 0.751 0.747 0.744 0.75 0.759 0.735 0.706 0.713 0.764 0.769 0.737 0.721 0.683 0.631 0.712 0.599
RpYN06 0.674 0.722 0.695 0.787 0.745 0.748 0.743 0.744 0.72 0.712 0.724 0.754 0.719 0.728 0.73 0.668 0.65 0.723 0.602
MjMP789 0.676 0.763 0.722 0.768 0.75 0.759 0.737 0.759 0.714 0.73 0.719 0.732 0.765 0.752 0.727 0.671 0.642 0.711 0.624
Rp22DB159 0.699 0.737 0.742 0.785 0.761 0.778 0.77 0.762 0.707 0.681 0.712 0.707 0.759 0.739 0.759 0.696 0.652 0.72 0.632
RbjCC9 0.716 0.726 0.747 0.796 0.786 0.774 0.768 0.783 0.717 0.69 0.717 0.721 0.729 0.729 0.696 0.711 0.662 0.75 0.632
RpYN2021 0.712 0.741 0.744 0.823 0.767 0.768 0.77 0.77 0.723 0.684 0.712 0.71 0.722 0.74 0.694 0.7 0.668 0.73 0.65
RbPrC31 0.71 0.751 0.732 0.821 0.778 0.771 0.783 0.783 0.744 0.689 0.717 0.712 0.72 0.742 0.698 0.699 0.651 0.741 0.631
RshSTT200 0.701 0.737 0.725 0.775 0.758 0.75 0.741 0.743 0.713 0.701 0.725 0.736 0.754 0.701 0.704 0.661 0.622 0.71 0.624
Ra22QT77 0.695 0.736 0.722 0.753 0.762 0.754 0.742 0.741 0.713 0.698 0.726 0.73 0.745 0.692 0.705 0.686 0.622 0.7 0.622
RacCS203 0.676 0.717 0.714 0.769 0.739 0.763 0.756 0.758 0.739 0.728 0.724 0.733 0.732 0.691 0.713 0.679 0.642 0.704 0.642
RmYN02 0.682 0.707 0.701 0.787 0.759 0.749 0.758 0.763 0.719 0.712 0.728 0.747 0.72 0.666 0.705 0.678 0.64 0.72 0.603
RmBANAL247 0.684 0.726 0.712 0.788 0.743 0.749 0.752 0.757 0.721 0.714 0.722 0.764 0.717 0.656 0.696 0.676 0.634 0.718 0.609
RmBANAL116 0.681 0.726 0.708 0.785 0.744 0.745 0.753 0.758 0.721 0.714 0.722 0.765 0.72 0.657 0.696 0.678 0.633 0.718 0.602
Species
1-1804
1804-2262
2262-3019
3019-3863
3863-5522
5522-7170
7170-9607
9607-11586
11586-12818
12818-15221
15221-18458
18458-21403
21403-22649
22649-23927
23927-24997
24997-25778
25778-27580
27580-28582
28582-30379
Nucleotide position on multiple sequence alignement
0.5 0.6 0.7 0.8 0.9

(F)

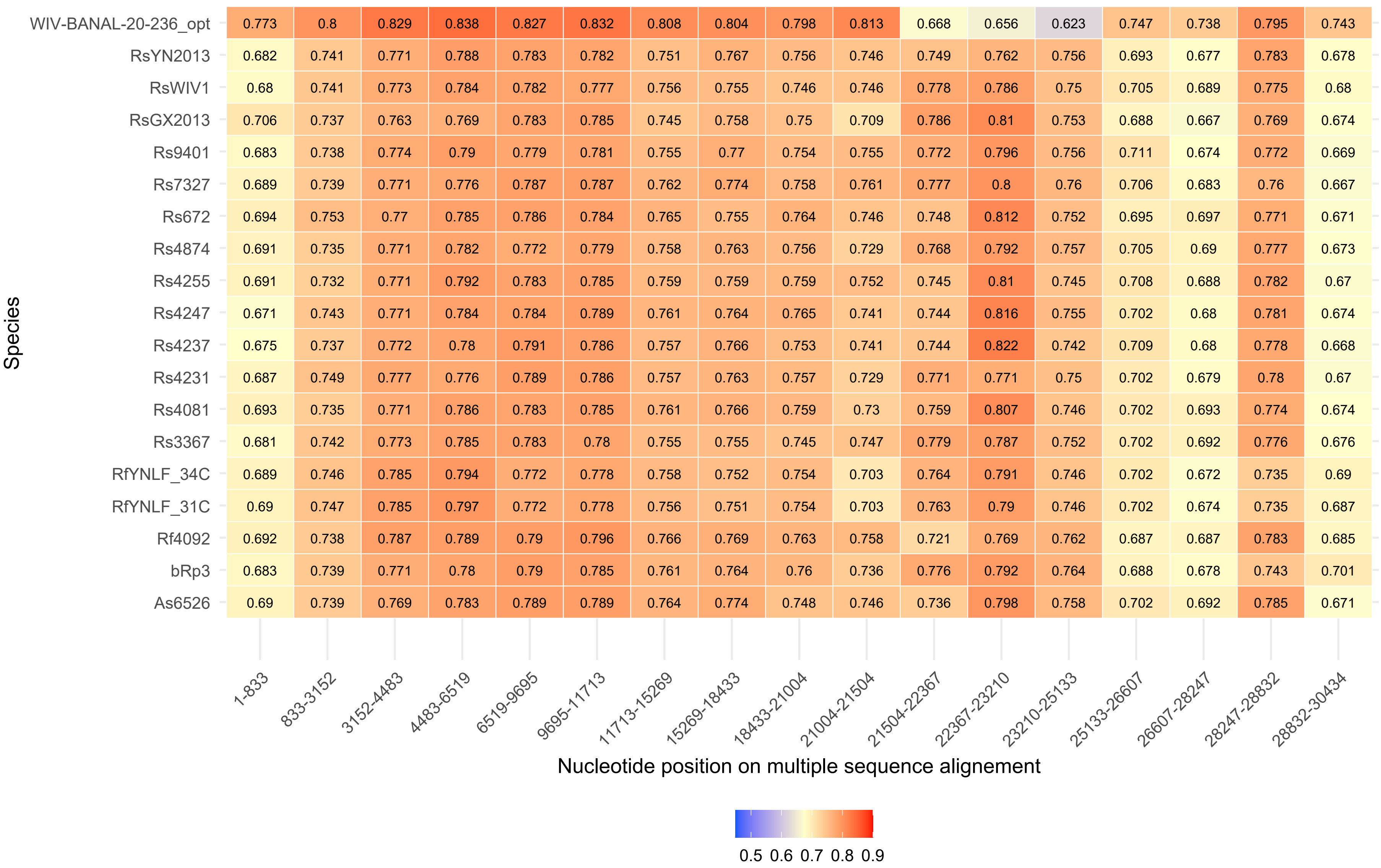
WIV-BANAL-20-236_opt 0.773 0.8 0.829 0.838 0.827 0.832 0.808 0.804 0.798 0.813 0.668 0.656 0.623 0.747 0.738 0.795 0.743
RsYN2013 0.682 0.741 0.771 0.788 0.783 0.782 0.751 0.767 0.756 0.746 0.749 0.762 0.756 0.693 0.677 0.783 0.678
RsWIV1 0.68 0.741 0.773 0.784 0.782 0.777 0.756 0.755 0.746 0.746 0.778 0.786 0.75 0.705 0.689 0.775 0.68
RsGX2013 0.706 0.737 0.763 0.769 0.783 0.785 0.745 0.758 0.75 0.709 0.786 0.81 0.753 0.688 0.667 0.769 0.674
Rs9401 0.683 0.738 0.774 0.79 0.779 0.781 0.755 0.77 0.754 0.755 0.772 0.796 0.756 0.711 0.674 0.772 0.669
Rs7327 0.689 0.739 0.771 0.776 0.787 0.787 0.762 0.774 0.758 0.761 0.777 0.8 0.76 0.706 0.683 0.76 0.667
Rs672 0.694 0.753 0.77 0.785 0.786 0.784 0.765 0.755 0.764 0.746 0.748 0.812 0.752 0.695 0.697 0.771 0.671
Rs4874 0.691 0.735 0.771 0.782 0.772 0.779 0.758 0.763 0.756 0.729 0.768 0.792 0.757 0.705 0.69 0.777 0.673
Rs4255 0.691 0.732 0.771 0.792 0.783 0.785 0.759 0.759 0.759 0.752 0.745 0.81 0.745 0.708 0.688 0.782 0.67
Rs4247 0.671 0.743 0.771 0.784 0.784 0.789 0.761 0.764 0.765 0.741 0.744 0.816 0.755 0.702 0.68 0.781 0.674
Rs4237 0.675 0.737 0.772 0.78 0.791 0.786 0.757 0.766 0.753 0.741 0.744 0.822 0.742 0.709 0.68 0.778 0.668
Rs4231 0.687 0.749 0.777 0.776 0.789 0.786 0.757 0.763 0.757 0.729 0.771 0.771 0.75 0.702 0.679 0.78 0.67
Rs4081 0.693 0.735 0.771 0.786 0.783 0.785 0.761 0.766 0.759 0.73 0.759 0.807 0.746 0.702 0.693 0.774 0.674
Rs3367 0.681 0.742 0.773 0.785 0.783 0.78 0.755 0.755 0.745 0.747 0.779 0.787 0.752 0.702 0.692 0.776 0.676
RfYNLF_34C 0.689 0.746 0.785 0.794 0.772 0.778 0.758 0.752 0.754 0.703 0.764 0.791 0.746 0.702 0.672 0.735 0.69
RfYNLF_31C 0.69 0.747 0.785 0.797 0.772 0.778 0.756 0.751 0.754 0.703 0.763 0.79 0.746 0.702 0.674 0.735 0.687
Rf4092 0.692 0.738 0.787 0.789 0.79 0.796 0.766 0.769 0.763 0.758 0.721 0.769 0.762 0.687 0.687 0.783 0.685
bRp3 0.683 0.739 0.771 0.78 0.79 0.785 0.761 0.764 0.76 0.736 0.776 0.792 0.764 0.688 0.678 0.743 0.701
As6526 0.69 0.739 0.769 0.783 0.789 0.789 0.764 0.774 0.748 0.746 0.736 0.798 0.758 0.702 0.692 0.785 0.671
Species
1-833 833-3152 3152-4483 4483-6519 6519-9695 9695-11713 11713-15269 15269-18433 18433-21004 21004-21504 21504-22367 22367-23210 23210-25133 25133-26607 26607-28247 28247-28832 28832-30434
Nucleotide position on multiple sequence alignement
0.5 0.6 0.7 0.8 0.9

(F)

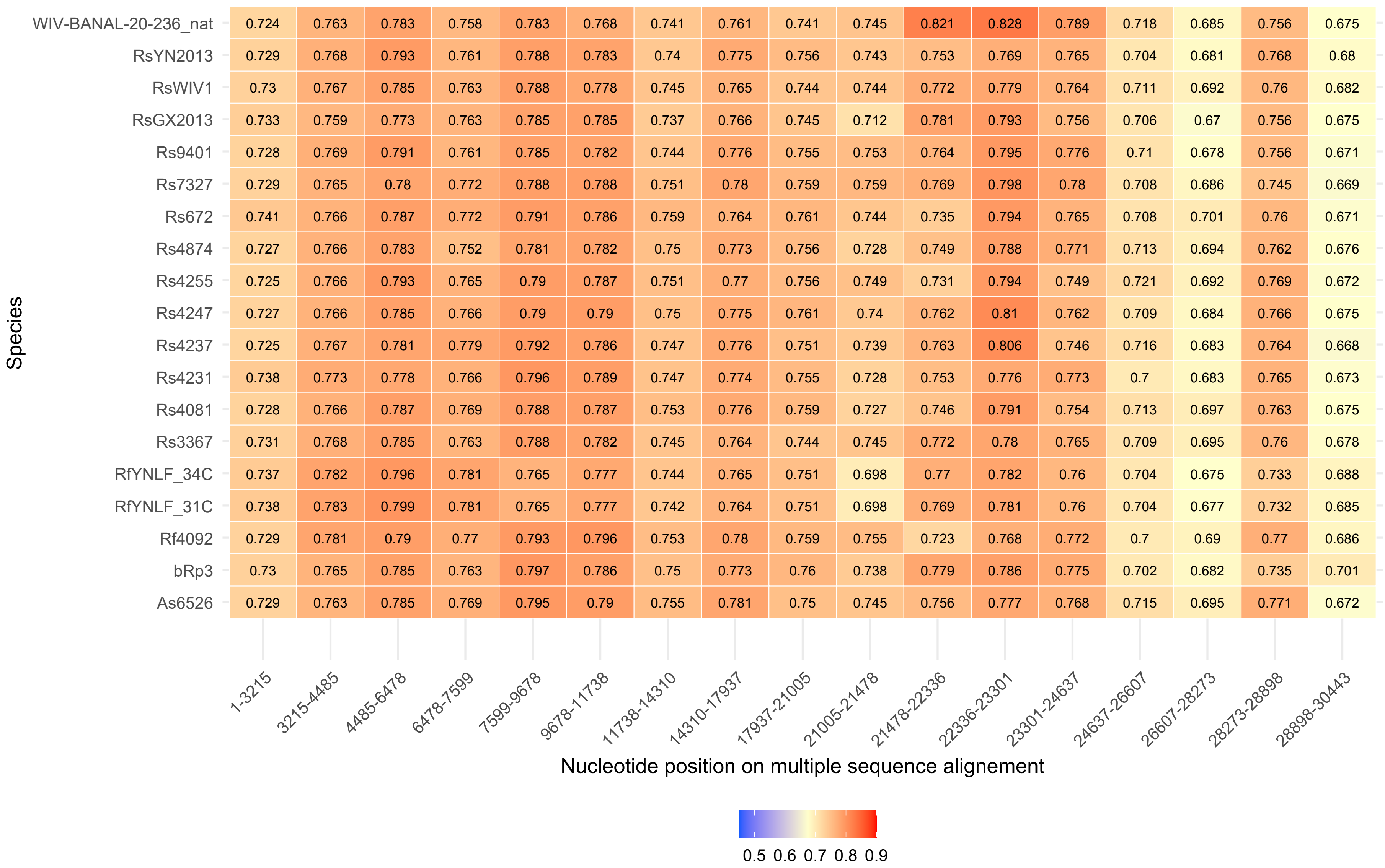
Species
WIV-BANAL-20-236_nat 0.724 0.763 0.783 0.758 0.783 0.768 0.741 0.761 0.741 0.745 0.821 0.828 0.789 0.718 0.685 0.756 0.675
RsYN2013 0.729 0.768 0.793 0.761 0.788 0.783 0.74 0.775 0.756 0.743 0.753 0.769 0.765 0.704 0.681 0.768 0.68
RsWIV1 0.73 0.767 0.785 0.763 0.788 0.778 0.745 0.765 0.744 0.744 0.772 0.779 0.764 0.711 0.692 0.76 0.682
RsGX2013 0.733 0.759 0.773 0.763 0.785 0.785 0.737 0.766 0.745 0.712 0.781 0.793 0.756 0.706 0.67 0.756 0.675
Rs9401 0.728 0.769 0.791 0.761 0.785 0.782 0.744 0.776 0.755 0.753 0.764 0.795 0.776 0.71 0.678 0.756 0.671
Rs7327 0.729 0.765 0.78 0.772 0.788 0.788 0.751 0.78 0.759 0.759 0.769 0.798 0.78 0.708 0.686 0.745 0.669
Rs672 0.741 0.766 0.787 0.772 0.791 0.786 0.759 0.764 0.761 0.744 0.735 0.794 0.765 0.708 0.701 0.76 0.671
Rs4874 0.727 0.766 0.783 0.752 0.781 0.782 0.75 0.773 0.756 0.728 0.749 0.788 0.771 0.713 0.694 0.762 0.676
Rs4255 0.725 0.766 0.793 0.765 0.79 0.787 0.751 0.77 0.756 0.749 0.731 0.794 0.749 0.721 0.692 0.769 0.672
Rs4247 0.727 0.766 0.785 0.766 0.79 0.79 0.75 0.775 0.761 0.74 0.762 0.81 0.762 0.709 0.684 0.766 0.675
Rs4237 0.725 0.767 0.781 0.779 0.792 0.786 0.747 0.776 0.751 0.739 0.763 0.806 0.746 0.716 0.683 0.764 0.668
Rs4231 0.738 0.773 0.778 0.766 0.796 0.789 0.747 0.774 0.755 0.728 0.753 0.776 0.773 0.7 0.683 0.765 0.673
Rs4081 0.728 0.766 0.787 0.769 0.788 0.787 0.753 0.776 0.759 0.727 0.746 0.791 0.754 0.713 0.697 0.763 0.675
Rs3367 0.731 0.768 0.785 0.763 0.788 0.782 0.745 0.764 0.744 0.745 0.772 0.78 0.765 0.709 0.695 0.76 0.678
RfYNLF_34C 0.737 0.782 0.796 0.781 0.765 0.777 0.744 0.765 0.751 0.698 0.77 0.782 0.76 0.704 0.675 0.733 0.688
RfYNLF_31C 0.738 0.783 0.799 0.781 0.765 0.777 0.742 0.764 0.751 0.698 0.769 0.781 0.76 0.704 0.677 0.732 0.685
Rf4092 0.729 0.781 0.79 0.77 0.793 0.796 0.753 0.78 0.759 0.755 0.723 0.768 0.772 0.7 0.69 0.77 0.686
bRp3 0.73 0.765 0.785 0.763 0.797 0.786 0.75 0.773 0.76 0.738 0.779 0.786 0.775 0.702 0.682 0.735 0.701
As6526 0.729 0.763 0.785 0.769 0.795 0.79 0.755 0.781 0.75 0.745 0.756 0.777 0.768 0.715 0.695 0.771 0.672
1-3215 3215-4485 4485-6478 6478-7599 7599-9678 9678-11738 11738-14310 14310-17937 17937-21005 21005-21478 21478-22336 22336-23301 23301-24637 24637-26607 26607-28273 28273-28898 28898-30443
Nucleotide position on multiple sequence alignement
0.5 0.6 0.7 0.8 0.9

Figure S17. Graphical representation of Codon Adaptation Index (CAI) comparing the codon usage of the focal segment to the rest of the genome, across the segments. (A) SARS-CoV-1, (B) MERS and (C) SARS-CoV-2.

(A)

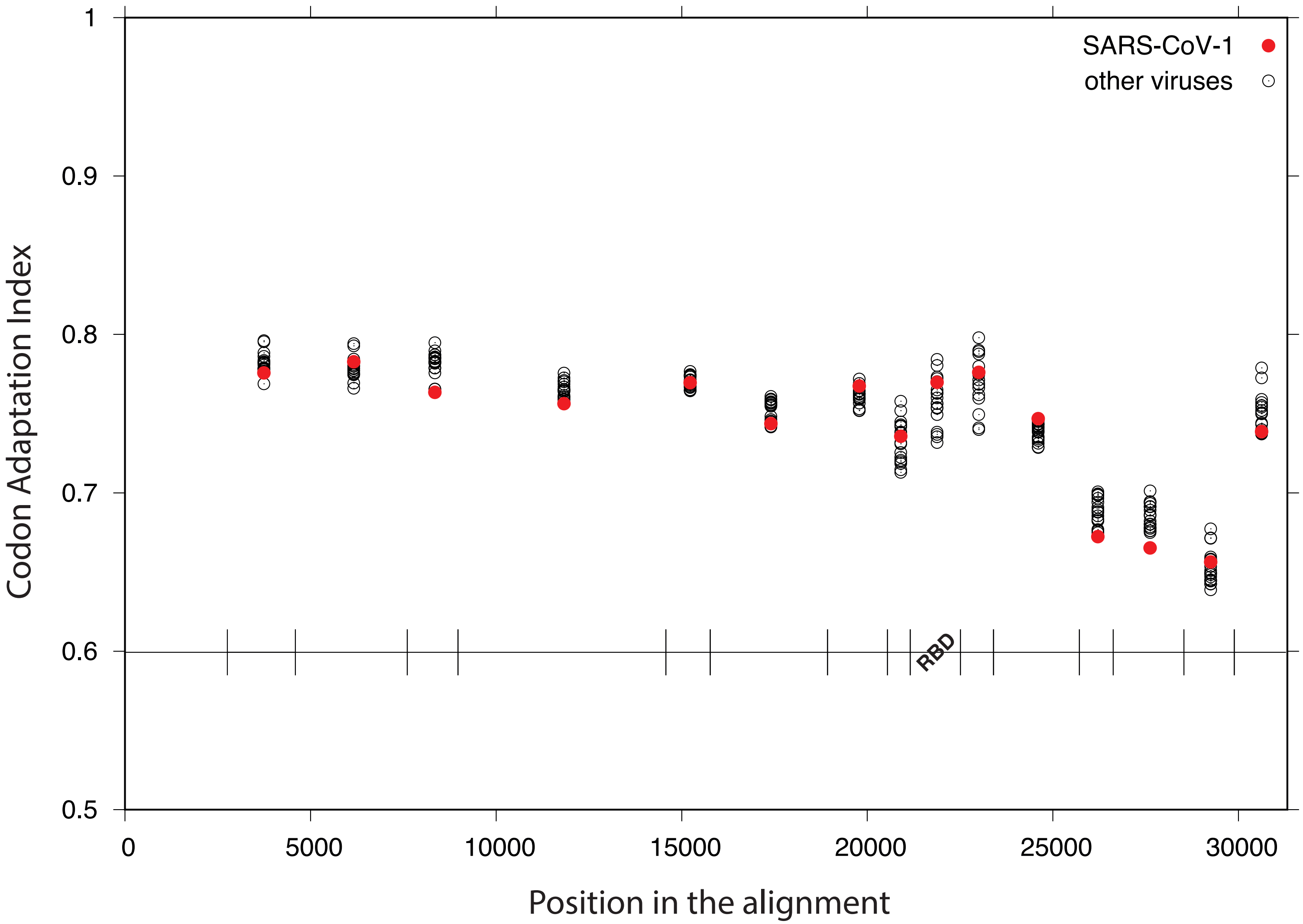
SARS-CoV-1
other viruses
Codon Adaptation Index
1
0.9
0.8
0.7
0.6
0.5
0
5000
10000
15000
20000
25000
30000
RBD
Position in the alignment

(B)

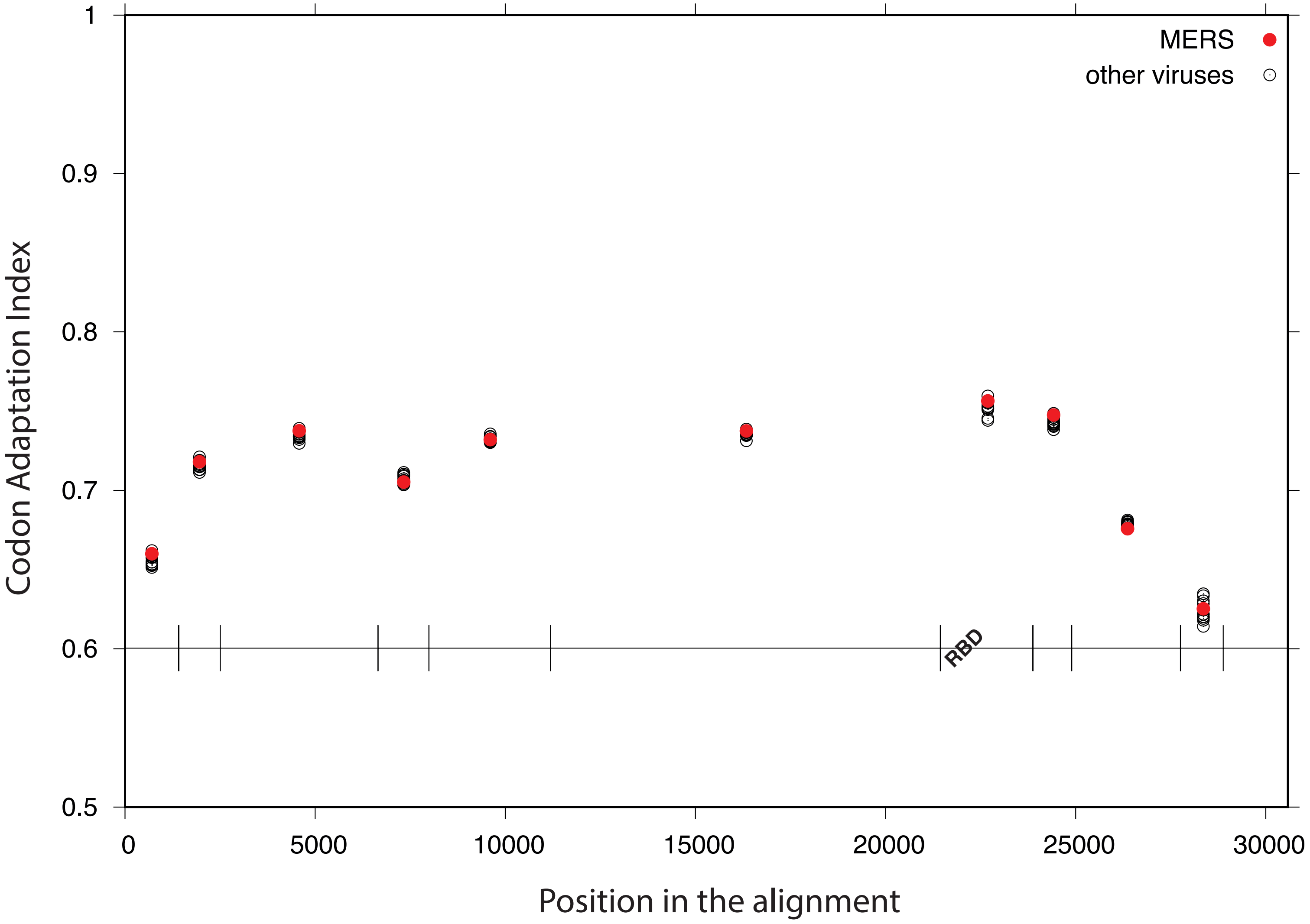

MERS
other viruses
Codon Adaptation Index
1
0.9
0.8
0.7
0.6
0.5
RBD
0
5000
10000
15000
20000
25000
30000
Position in the alignment

(C)

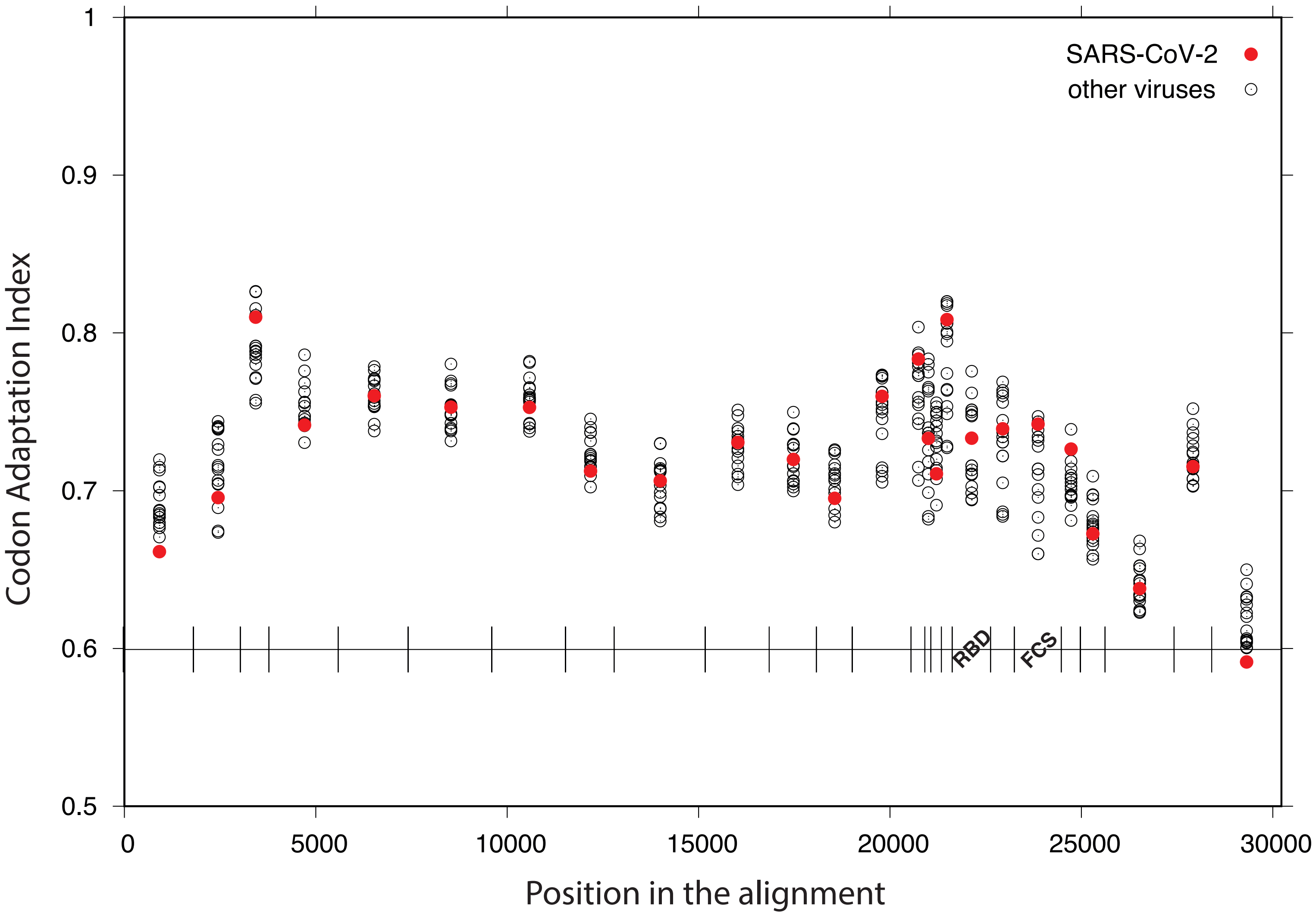
SARS-CoV-2
other viruses
Codon Adaptation Index
1
0.9
0.8
0.7
0.6
0.5
0
5000
10000
15000
20000
25000
30000
Position in the alignment
RBD
FCS

Figure S18. Estimation of the Codon Adaptation Index (CAI) across all the segments of the three datasets in the external branches of the local tree. (A) SARS-CoV-1, (B) MERS and (C) SARS-CoV-2, (D) SARS-CoV-2 with only 14 segments (larger segments), (E) CoV-2-S$^{edit}$, (F) WIV-BANAL-20-236$^{opt}$, and (G) WIV-BANAL-20-236$^{nat}$.

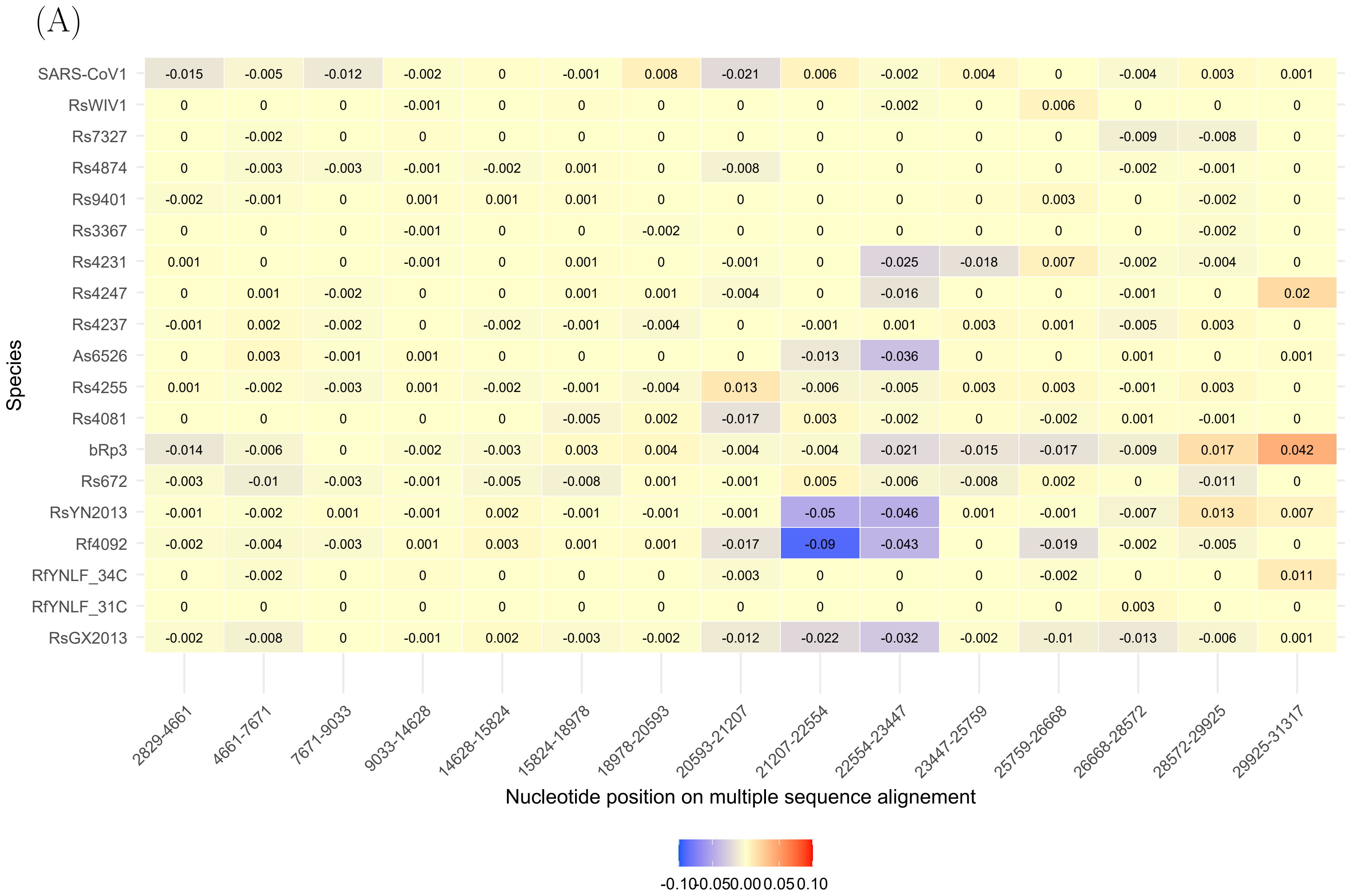
(A)
Species
SARS-CoV1 -0.015 -0.005 -0.012 -0.002 0 -0.001 0.008 -0.021 0.006 -0.002 0.004 0 -0.004 0.003 0.001
RsWIV1 0 0 0 -0.001 0 0 0 0 0 -0.002 0 0.006 0 0 0
Rs7327 0 -0.002 0 0 0 0 0 0 0 0 0 0 -0.009 -0.008 0
Rs4874 0 -0.003 -0.003 -0.001 -0.002 0.001 0 -0.008 0 0 0 0 -0.002 -0.001 0
Rs9401 -0.002 -0.001 0 0.001 0.001 0.001 0 0 0 0 0 0.003 0 -0.002 0
Rs3367 0 0 0 -0.001 0 0 -0.002 0 0 0 0 0 0 -0.002 0
Rs4231 0.001 0 0 -0.001 0 0.001 0 -0.001 0 -0.025 -0.018 0.007 -0.002 -0.004 0
Rs4247 0 0.001 -0.002 0 0 0.001 0.001 -0.004 0 -0.016 0 0 -0.001 0 0.02
Rs4237 -0.001 0.002 -0.002 0 -0.002 -0.001 -0.004 0 -0.001 0.001 0.003 0.001 -0.005 0.003 0
As6526 0 0.003 -0.001 0.001 0 0 0 0 -0.013 -0.036 0 0 0.001 0 0.001
Rs4255 0.001 -0.002 -0.003 0.001 -0.002 -0.001 -0.004 0.013 -0.006 -0.005 0.003 0.003 -0.001 0.003 0
Rs4081 0 0 0 0 0 -0.005 0.002 -0.017 0.003 -0.002 0 -0.002 0.001 -0.001 0
bRp3 -0.014 -0.006 0 -0.002 -0.003 0.003 0.004 -0.004 -0.004 -0.021 -0.015 -0.017 -0.009 0.017 0.042
Rs672 -0.003 -0.01 -0.003 -0.001 -0.005 -0.008 0.001 -0.001 0.005 -0.006 -0.008 0.002 0 -0.011 0
RsYN2013 -0.001 -0.002 0.001 -0.001 0.002 -0.001 -0.001 -0.001 -0.05 -0.046 0.001 -0.001 -0.007 0.013 0.007
Rf4092 -0.002 -0.004 -0.003 0.001 0.003 0.001 0.001 -0.017 -0.09 -0.043 0 -0.019 -0.002 -0.005 0
RfYNLF_34C 0 -0.002 0 0 0 0 0 -0.003 0 0 0 -0.002 0 0 0.011
RfYNLF_31C 0 0 0 0 0 0 0 0 0 0 0 0 0.003 0 0
RsGX2013 -0.002 -0.008 0 -0.001 0.002 -0.003 -0.002 -0.012 -0.022 -0.032 -0.002 -0.01 -0.013 -0.006 0.001
2829-4661
4661-7671
7671-9033
9033-14628
14628-15824
15824-18978
18978-20593
20593-21207
21207-22554
22554-23447
23447-25759
25759-26668
26668-28572
28572-29925
29925-31317
Nucleotide position on multiple sequence alignement
-0.10 -0.05 0.00 0.05 0.10

(B)

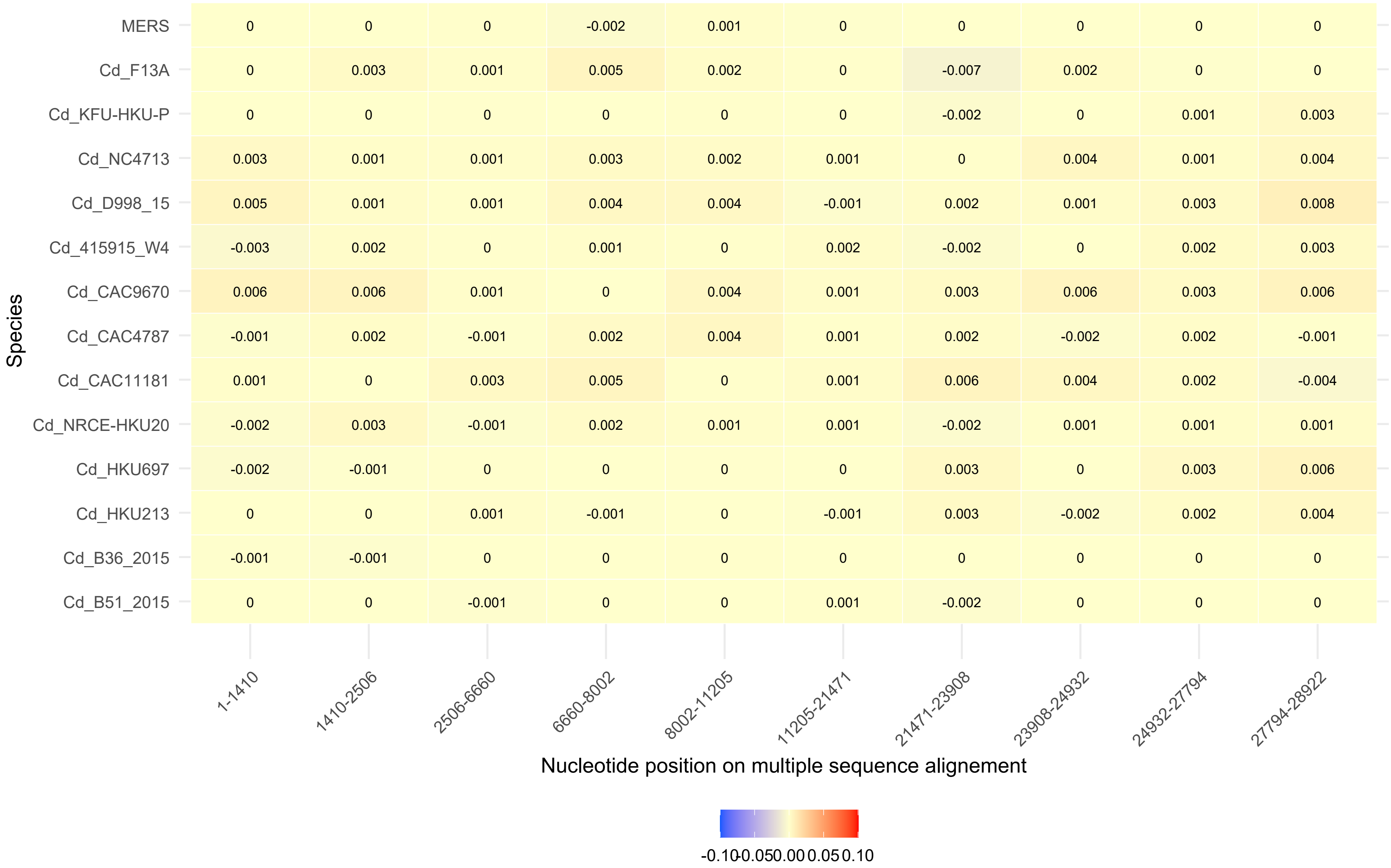

Species
MERS
Cd_F13A
Cd_KFU-HKU-P
Cd_NC4713
Cd_D998_15
Cd_415915_W4
Cd_CAC9670
Cd_CAC4787
Cd_CAC11181
Cd_NRCE-HKU20
Cd_HKU697
Cd_HKU213
Cd_B36_2015
Cd_B51_2015
0 0 0 -0.002 0.001 0 0 0 0 0
0 0.003 0.001 0.005 0.002 0 -0.007 0.002 0 0
0 0 0 0 0 0 -0.002 0 0.001 0.003
0.003 0.001 0.001 0.003 0.002 0.001 0 0.004 0.001 0.004
0.005 0.001 0.001 0.004 0.004 -0.001 0.002 0.001 0.003 0.008
-0.003 0.002 0 0.001 0 0.002 -0.002 0 0.002 0.003
0.006 0.006 0.001 0 0.004 0.001 0.003 0.006 0.003 0.006
-0.001 0.002 -0.001 0.002 0.004 0.001 0.002 -0.002 0.002 -0.001
0.001 0 0.003 0.005 0 0.001 0.006 0.004 0.002 -0.004
-0.002 0.003 -0.001 0.002 0.001 0.001 -0.002 0.001 0.001 0.001
-0.002 -0.001 0 0 0 0 0.003 0 0.003 0.006
0 0 0.001 -0.001 0 -0.001 0.003 -0.002 0.002 0.004
-0.001 -0.001 0 0 0 0 0 0 0 0
0 0 -0.001 0 0 0.001 -0.002 0 0 0
1-1410
1410-2506
2506-6660
6660-8002
8002-11205
11205-21471
21471-23908
23908-24932
24932-27794
27794-28922
Nucleotide position on multiple sequence alignement
-0.10
-0.05
0.00
0.05
0.10

(C)

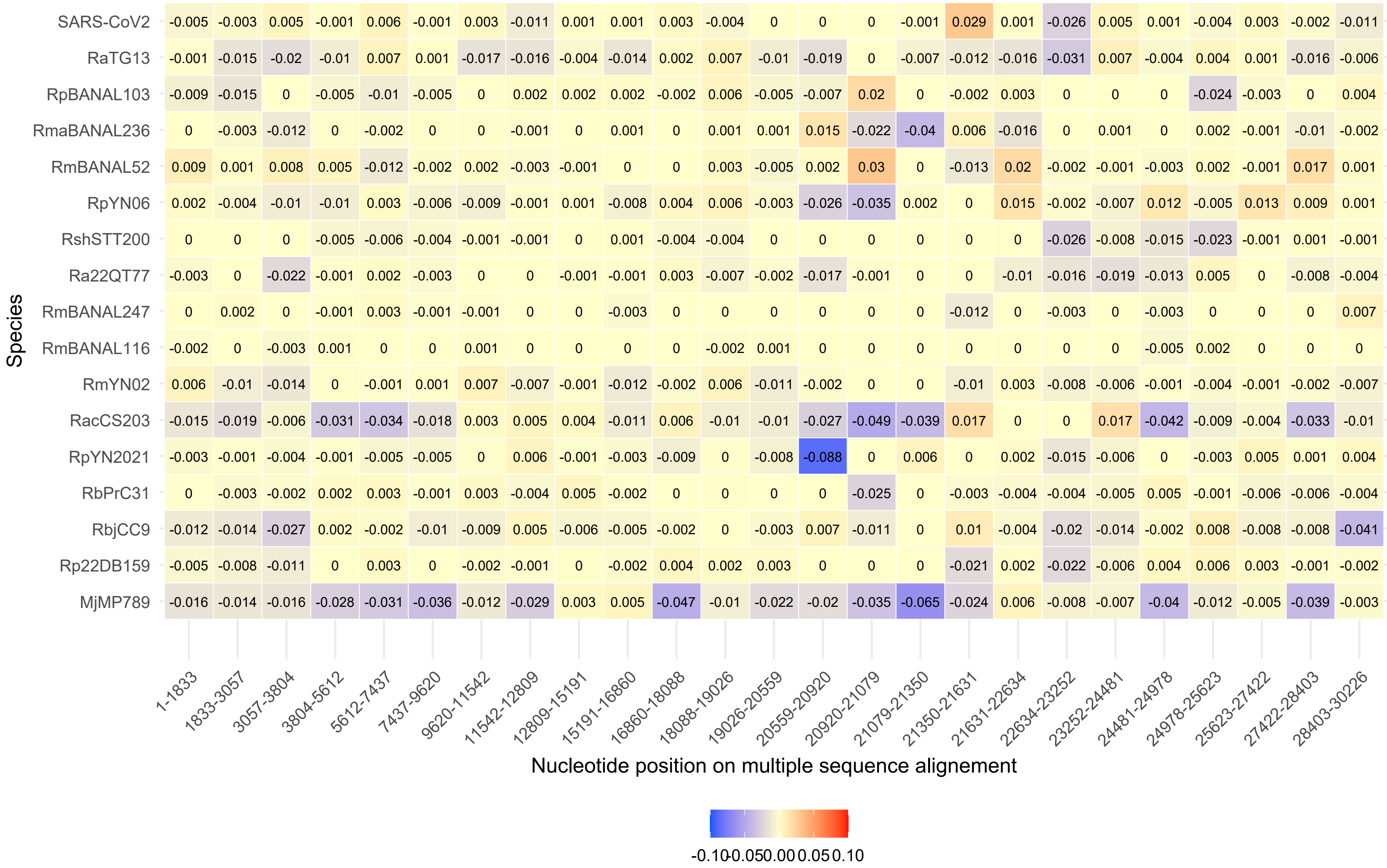
Species
SARS-CoV2 -0.005 -0.003 0.005 -0.001 0.006 -0.001 0.003 -0.011 0.001 0.001 0.003 -0.004 0 0 0 -0.001 0.029 0.001 -0.026 0.005 0.001 -0.004 0.003 -0.002 -0.011
RaTG13 -0.001 -0.015 -0.02 -0.01 0.007 0.001 -0.017 -0.016 -0.004 -0.014 0.002 0.007 -0.01 -0.019 0 -0.007 -0.012 -0.016 -0.031 0.007 -0.004 0.004 0.001 -0.016 -0.006
RpBANAL103 -0.009 -0.015 0 -0.005 -0.01 -0.005 0 0.002 0.002 0.002 -0.002 0.006 -0.005 -0.007 0.02 0 -0.002 0.003 0 0 0 -0.024 -0.003 0 0.004
RmaBANAL236 0 -0.003 -0.012 0 -0.002 0 0 -0.001 0 0.001 0 0.001 0.001 0.015 -0.022 -0.04 0.006 -0.016 0 0.001 0 0.002 -0.001 -0.01 -0.002
RmBANAL52 0.009 0.001 0.008 0.005 -0.012 -0.002 0.002 -0.003 -0.001 0 0 0.003 -0.005 0.002 0.03 0 -0.013 0.02 -0.002 -0.001 -0.003 0.002 -0.001 0.017 0.001
RpYN06 0.002 -0.004 -0.01 -0.01 0.003 -0.006 -0.009 -0.001 0.001 -0.008 0.004 0.006 -0.003 -0.026 -0.035 0.002 0 0.015 -0.002 -0.007 0.012 -0.005 0.013 0.009 0.001
RshSTT200 0 0 0 -0.005 -0.006 -0.004 -0.001 -0.001 0 0.001 -0.004 -0.004 0 0 0 0 0 0 -0.026 -0.008 -0.015 -0.023 -0.001 0.001 -0.001
Ra22QT77 -0.003 0 -0.022 -0.001 0.002 -0.003 0 0 -0.001 -0.001 0.003 -0.007 -0.002 -0.017 -0.001 0 0 -0.01 -0.016 -0.019 -0.013 0.005 0 -0.008 -0.004
RmBANAL247 0 0.002 0 -0.001 0.003 -0.001 -0.001 0 0 -0.003 0 0 0 0 0 0 -0.012 0 -0.003 0 -0.003 0 0 0 0.007
RmBANAL116 -0.002 0 -0.003 0.001 0 0 0.001 0 0 0 0 -0.002 0.001 0 0 0 0 0 0 0 -0.005 0.002 0 0 0
RmYN02 0.006 -0.01 -0.014 0 -0.001 0.001 0.007 -0.007 -0.001 -0.012 -0.002 0.006 -0.011 -0.002 0 0 -0.01 0.003 -0.008 -0.006 -0.001 -0.004 -0.001 -0.002 -0.007
RacCS203 -0.015 -0.019 -0.006 -0.031 -0.034 -0.018 0.003 0.005 0.004 -0.011 0.006 -0.01 -0.01 -0.027 -0.049 -0.039 0.017 0 0 0.017 -0.042 -0.009 -0.004 -0.033 -0.01
RpYN2021 -0.003 -0.001 -0.004 -0.001 -0.005 -0.005 0 0.006 -0.001 -0.003 -0.009 0 -0.008 -0.088 0 0.006 0 0.002 -0.015 -0.006 0 -0.003 0.005 0.001 0.004
RbPrC31 0 -0.003 -0.002 0.002 0.003 -0.001 0.003 -0.004 0.005 -0.002 0 0 0 0 -0.025 0 -0.003 -0.004 -0.004 -0.005 0.005 -0.001 -0.006 -0.006 -0.004
RbjCC9 -0.012 -0.014 -0.027 0.002 -0.002 -0.01 -0.009 0.005 -0.006 -0.005 -0.002 0 -0.003 0.007 -0.011 0 0.01 -0.004 -0.02 -0.014 -0.002 0.008 -0.008 -0.008 -0.041
Rp22DB159 -0.005 -0.008 -0.011 0 0.003 0 -0.002 -0.001 0 -0.002 0.004 0.002 0.003 0 0 0 -0.021 0.002 -0.022 -0.006 0.004 0.006 0.003 -0.001 -0.002
MjMP789 -0.016 -0.014 -0.016 -0.028 -0.031 -0.036 -0.012 -0.029 0.003 0.005 -0.047 -0.01 -0.022 -0.02 -0.035 -0.065 -0.024 0.006 -0.008 -0.007 -0.04 -0.012 -0.005 -0.039 -0.003
1-1833 1833-3057 3057-3804 3804-5612 5612-7437 7437-9620 9620-11542 11542-12809 12809-15191 15191-16860 16860-18088 18088-19026 19026-20559 20559-20920 20920-21079 21079-21350 21350-21631 21631-22634 22634-23252 23252-24481 24481-24978 24978-25623 25623-27422 27422-28403 28403-30226
Nucleotide position on multiple sequence alignement
-0.10 -0.05 0.00 0.05 0.10

(D)

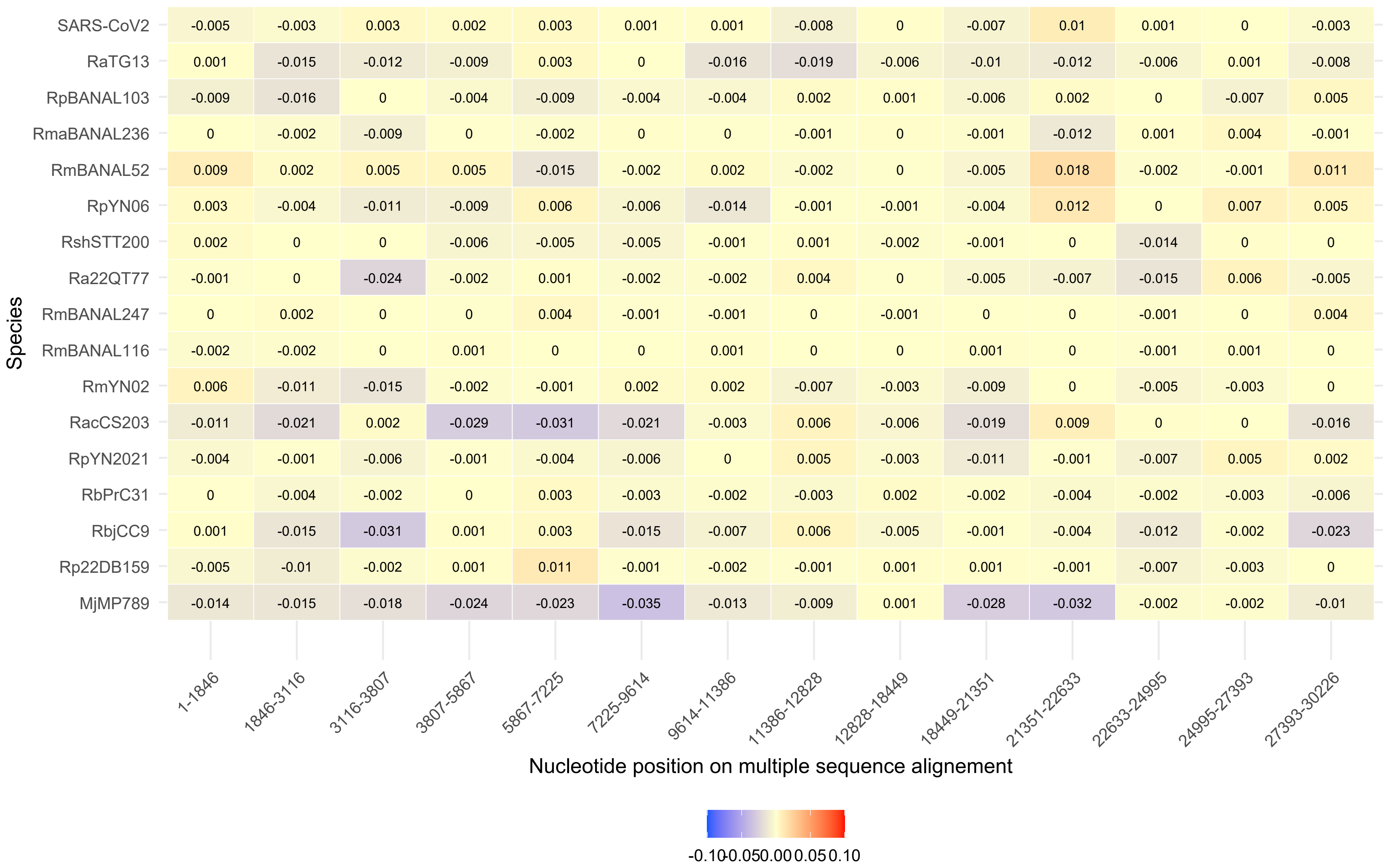
Species
Nucleotide position on multiple sequence alignement
1-1846
1846-3116
3116-3807
3807-5867
5867-7225
7225-9614
9614-11386
11386-12828
12828-18449
18449-21351
21351-22633
22633-24995
24995-27393
27393-30226
SARS-CoV2 -0.005 -0.003 0.003 0.002 0.003 0.001 0.001 -0.008 0 -0.007 0.01 0.001 0 -0.003
RaTG13 0.001 -0.015 -0.012 -0.009 0.003 0 -0.016 -0.019 -0.006 -0.01 -0.012 -0.006 0.001 -0.008
RpBANAL103 -0.009 -0.016 0 -0.004 -0.009 -0.004 -0.004 0.002 0.001 -0.006 0.002 0 -0.007 0.005
RmaBANAL236 0 -0.002 -0.009 0 -0.002 0 0 -0.001 0 -0.001 -0.012 0.001 0.004 -0.001
RmBANAL52 0.009 0.002 0.005 0.005 -0.015 -0.002 0.002 -0.002 0 -0.005 0.018 -0.002 -0.001 0.011
RpYN06 0.003 -0.004 -0.011 -0.009 0.006 -0.006 -0.014 -0.001 -0.001 -0.004 0.012 0 0.007 0.005
RshSTT200 0.002 0 0 -0.006 -0.005 -0.005 -0.001 0.001 -0.002 -0.001 0 -0.014 0 0
Ra22QT77 -0.001 0 -0.024 -0.002 0.001 -0.002 -0.002 0.004 0 -0.005 -0.007 -0.015 0.006 -0.005
RmBANAL247 0 0.002 0 0 0.004 -0.001 -0.001 0 -0.001 0 0 -0.001 0 0.004
RmBANAL116 -0.002 -0.002 0 0.001 0 0 0.001 0 0 0.001 0 -0.001 0.001 0
RmYN02 0.006 -0.011 -0.015 -0.002 -0.001 0.002 0.002 -0.007 -0.003 -0.009 0 -0.005 -0.003 0
RacCS203 -0.011 -0.021 0.002 -0.029 -0.031 -0.021 -0.003 0.006 -0.006 -0.019 0.009 0 0 -0.016
RpYN2021 -0.004 -0.001 -0.006 -0.001 -0.004 -0.006 0 0.005 -0.003 -0.011 -0.001 -0.007 0.005 0.002
RbPrC31 0 -0.004 -0.002 0 0.003 -0.003 -0.002 -0.003 0.002 -0.002 -0.004 -0.002 -0.003 -0.006
RbjCC9 0.001 -0.015 -0.031 0.001 0.003 -0.015 -0.007 0.006 -0.005 -0.001 -0.004 -0.012 -0.002 -0.023
Rp22DB159 -0.005 -0.01 -0.002 0.001 0.011 -0.001 -0.002 -0.001 0.001 0.001 -0.001 -0.007 -0.003 0
MjMP789 -0.014 -0.015 -0.018 -0.024 -0.023 -0.035 -0.013 -0.009 0.001 -0.028 -0.032 -0.002 -0.002 -0.01
-0.10-0.050.000.050.10

(E)

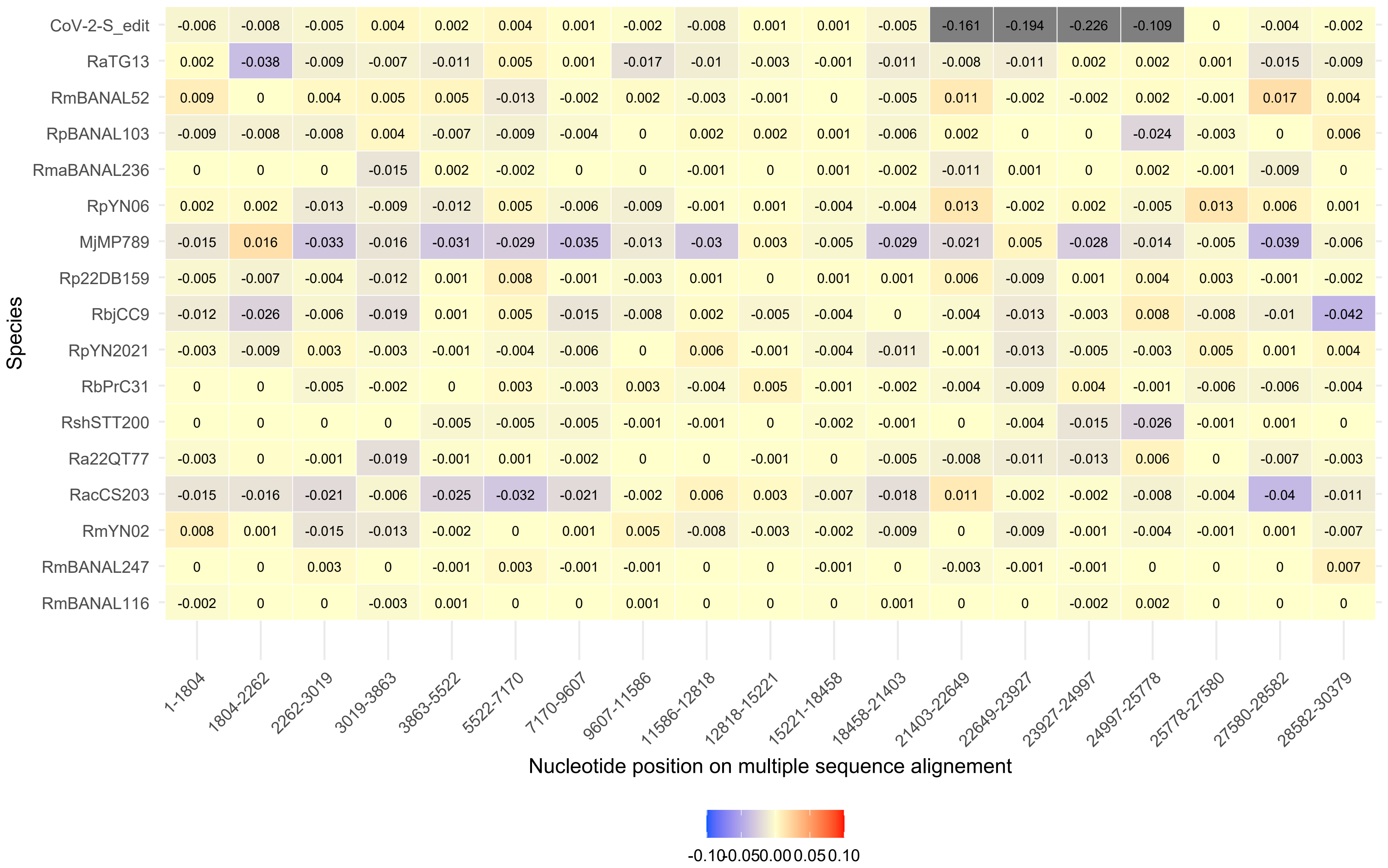
CoV-2-S_edit -0.006 -0.008 -0.005 0.004 0.002 0.004 0.001 -0.002 -0.008 0.001 0.001 -0.005 -0.161 -0.194 -0.226 -0.109 0 -0.004 -0.002
RaTG13 0.002 -0.038 -0.009 -0.007 -0.011 0.005 0.001 -0.017 -0.01 -0.003 -0.001 -0.011 -0.008 -0.011 0.002 0.002 0.001 -0.015 -0.009
RmBANAL52 0.009 0 0.004 0.005 0.005 -0.013 -0.002 0.002 -0.003 -0.001 0 -0.005 0.011 -0.002 -0.002 0.002 -0.001 0.017 0.004
RpBANAL103 -0.009 -0.008 -0.008 0.004 -0.007 -0.009 -0.004 0 0.002 0.002 0.001 -0.006 0.002 0 0 -0.024 -0.003 0 0.006
RmaBANAL236 0 0 0 -0.015 0.002 -0.002 0 0 -0.001 0 0.001 -0.002 -0.011 0.001 0 0.002 -0.001 -0.009 0
RpYN06 0.002 0.002 -0.013 -0.009 -0.012 0.005 -0.006 -0.009 -0.001 0.001 -0.004 -0.004 0.013 -0.002 0.002 -0.005 0.013 0.006 0.001
MjMP789 -0.015 0.016 -0.033 -0.016 -0.031 -0.029 -0.035 -0.013 -0.03 0.003 -0.005 -0.029 -0.021 0.005 -0.028 -0.014 -0.005 -0.039 -0.006
Rp22DB159 -0.005 -0.007 -0.004 -0.012 0.001 0.008 -0.001 -0.003 0.001 0 0.001 0.001 0.006 -0.009 0.001 0.004 0.003 -0.001 -0.002
RbjCC9 -0.012 -0.026 -0.006 -0.019 0.001 0.005 -0.015 -0.008 0.002 -0.005 -0.004 0 -0.004 -0.013 -0.003 0.008 -0.008 -0.01 -0.042
RpYN2021 -0.003 -0.009 0.003 -0.003 -0.001 -0.004 -0.006 0 0.006 -0.001 -0.004 -0.011 -0.001 -0.013 -0.005 -0.003 0.005 0.001 0.004
RbPrC31 0 0 -0.005 -0.002 0 0.003 -0.003 0.003 -0.004 0.005 -0.001 -0.002 -0.004 -0.009 0.004 -0.001 -0.006 -0.006 -0.004
RshSTT200 0 0 0 0 -0.005 -0.005 -0.005 -0.001 -0.001 0 -0.002 -0.001 0 -0.004 -0.015 -0.026 -0.001 0.001 0
Ra22QT77 -0.003 0 -0.001 -0.019 -0.001 0.001 -0.002 0 0 -0.001 0 -0.005 -0.008 -0.011 -0.013 0.006 0 -0.007 -0.003
RacCS203 -0.015 -0.016 -0.021 -0.006 -0.025 -0.032 -0.021 -0.002 0.006 0.003 -0.007 -0.018 0.011 -0.002 -0.002 -0.008 -0.004 -0.04 -0.011
RmYN02 0.008 0.001 -0.015 -0.013 -0.002 0 0.001 0.005 -0.008 -0.003 -0.002 -0.009 0 -0.009 -0.001 -0.004 -0.001 0.001 -0.007
RmBANAL247 0 0 0.003 0 -0.001 0.003 -0.001 -0.001 0 0 -0.001 0 -0.003 -0.001 -0.001 0 0 0 0.007
RmBANAL116 -0.002 0 0 -0.003 0.001 0 0 0.001 0 0 0 0.001 0 0 -0.002 0.002 0 0 0
Species
1-1804
1804-2262
2262-3019
3019-3863
3863-5522
5522-7170
7170-9607
9607-11586
11586-12818
12818-15221
15221-18458
18458-21403
21403-22649
22649-23927
23927-24997
24997-25778
25778-27580
27580-28582
28582-30379
Nucleotide position on multiple sequence alignement
-0.10 -0.05 0.00 0.05 0.10

(F)

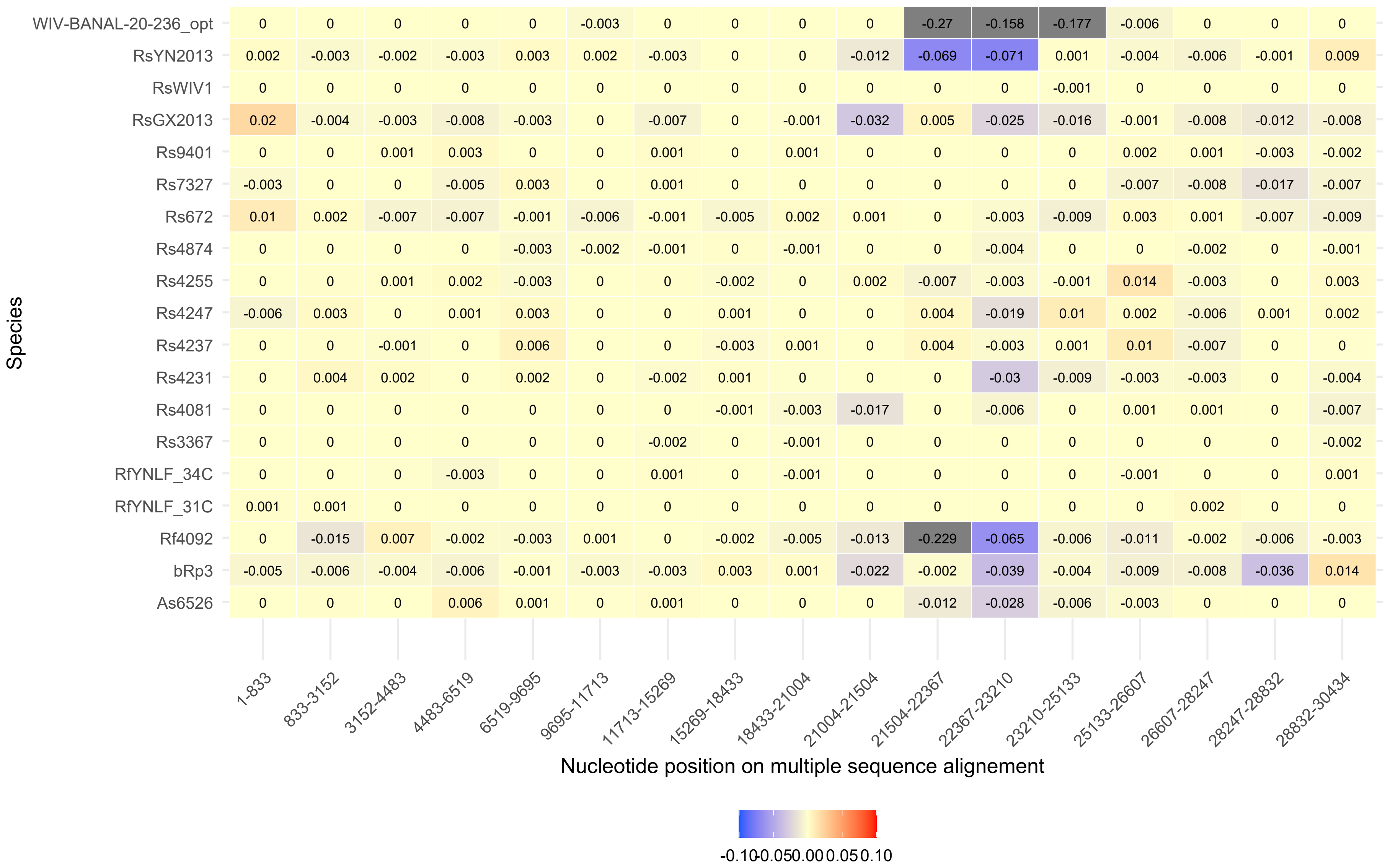
WIV-BANAL-20-236_opt 0 0 0 0 0 -0.003 0 0 0 0 -0.27 -0.158 -0.177 -0.006 0 0 0
RsYN2013 0.002 -0.003 -0.002 -0.003 0.003 0.002 -0.003 0 0 -0.012 -0.069 -0.071 0.001 -0.004 -0.006 -0.001 0.009
RsWIV1 0 0 0 0 0 0 0 0 0 0 0 0 -0.001 0 0 0 0
RsGX2013 0.02 -0.004 -0.003 -0.008 -0.003 0 -0.007 0 -0.001 -0.032 0.005 -0.025 -0.016 -0.001 -0.008 -0.012 -0.008
Rs9401 0 0 0.001 0.003 0 0 0.001 0 0.001 0 0 0 0 0.002 0.001 -0.003 -0.002
Rs7327 -0.003 0 0 -0.005 0.003 0 0.001 0 0 0 0 0 0 -0.007 -0.008 -0.017 -0.007
Rs672 0.01 0.002 -0.007 -0.007 -0.001 -0.006 -0.001 -0.005 0.002 0.001 0 -0.003 -0.009 0.003 0.001 -0.007 -0.009
Rs4874 0 0 0 0 -0.003 -0.002 -0.001 0 -0.001 0 0 -0.004 0 0 -0.002 0 -0.001
Rs4255 0 0 0.001 0.002 -0.003 0 0 -0.002 0 0.002 -0.007 -0.003 -0.001 0.014 -0.003 0 0.003
Rs4247 -0.006 0.003 0 0.001 0.003 0 0 0.001 0 0 0.004 -0.019 0.01 0.002 -0.006 0.001 0.002
Rs4237 0 0 -0.001 0 0.006 0 0 -0.003 0.001 0 0.004 -0.003 0.001 0.01 -0.007 0 0
Rs4231 0 0.004 0.002 0 0.002 0 -0.002 0.001 0 0 0 -0.03 -0.009 -0.003 -0.003 0 -0.004
Rs4081 0 0 0 0 0 0 0 -0.001 -0.003 -0.017 0 -0.006 0 0.001 0.001 0 -0.007
Rs3367 0 0 0 0 0 0 -0.002 0 -0.001 0 0 0 0 0 0 0 -0.002
RfYNLF_34C 0 0 0 -0.003 0 0 0.001 0 -0.001 0 0 0 0 -0.001 0 0 0.001
RfYNLF_31C 0.001 0.001 0 0 0 0 0 0 0 0 0 0 0 0 0.002 0 0
Rf4092 0 -0.015 0.007 -0.002 -0.003 0.001 0 -0.002 -0.005 -0.013 -0.229 -0.065 -0.006 -0.011 -0.002 -0.006 -0.003
bRp3 -0.005 -0.006 -0.004 -0.006 -0.001 -0.003 -0.003 0.003 0.001 -0.022 -0.002 -0.039 -0.004 -0.009 -0.008 -0.036 0.014
As6526 0 0 0 0.006 0.001 0 0.001 0 0 0 -0.012 -0.028 -0.006 -0.003 0 0 0
Species
1-833
833-3152
3152-4483
4483-6519
6519-9695
9695-11713
11713-15269
15269-18433
18433-21004
21004-21504
21504-22367
22367-23210
23210-25133
25133-26607
26607-28247
28247-28832
28832-30434
Nucleotide position on multiple sequence alignement
-0.10 -0.05 0.00 0.05 0.10

(F)

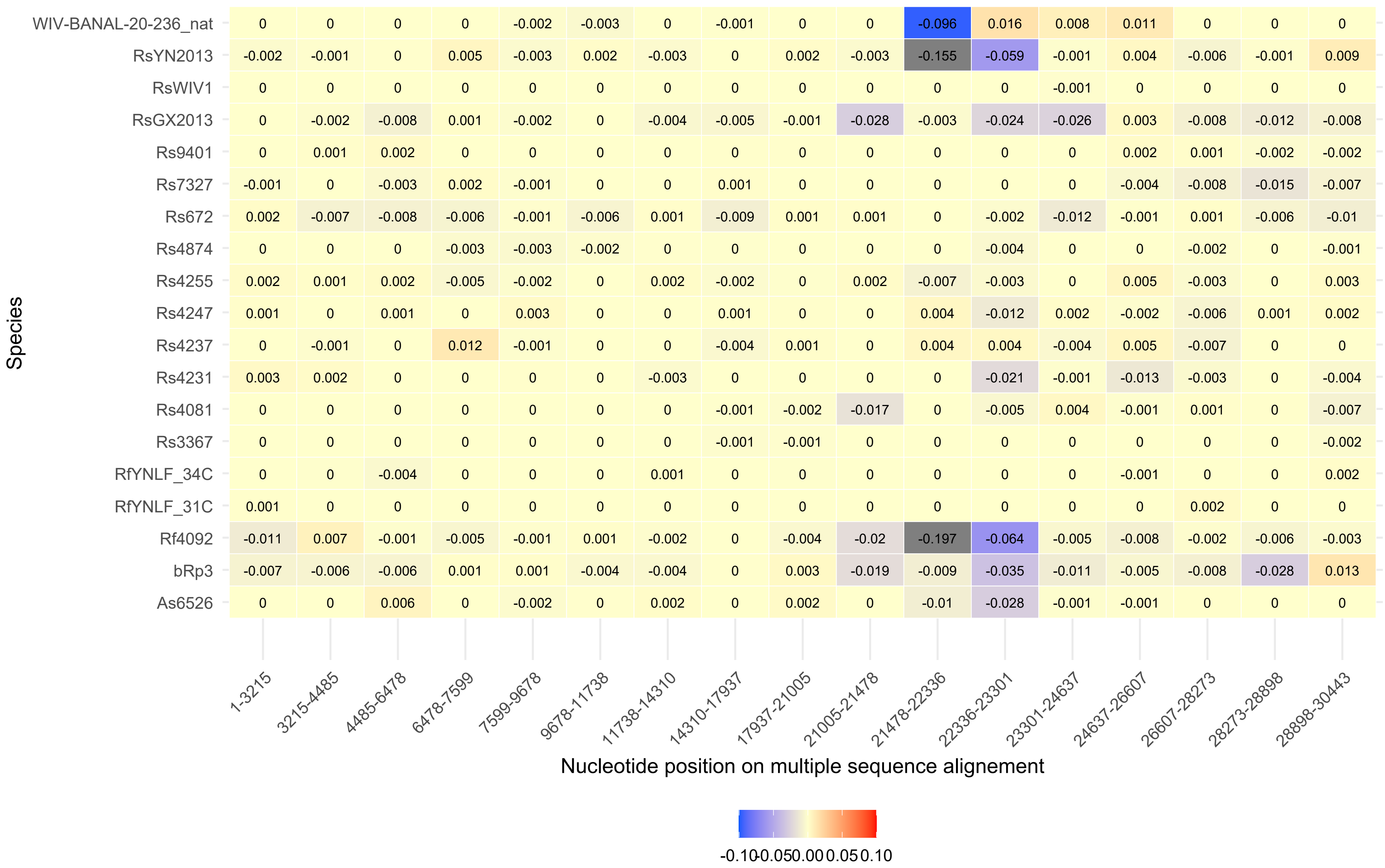
Species
Nucleotide position on multiple sequence alignement
1-3215 | 3215-4485 | 4485-6478 | 6478-7599 | 7599-9678 | 9678-11738 | 11738-14310 | 14310-17937 | 17937-21005 | 21005-21478 | 21478-22336 | 22336-23301 | 23301-24637 | 24637-26607 | 26607-28273 | 28273-28898 | 28898-30443
WIV-BANAL-20-236_nat: 0 0 0 0 -0.002 -0.003 0 -0.001 0 0 -0.096 0.016 0.008 0.011 0 0 0
RsYN2013: -0.002 -0.001 0 0.005 -0.003 0.002 -0.003 0 0.002 -0.003 -0.155 -0.059 -0.001 0.004 -0.006 -0.001 0.009
RsWIV1: 0 0 0 0 0 0 0 0 0 0 0 0 -0.001 0 0 0 0
RsGX2013: 0 -0.002 -0.008 0.001 -0.002 0 -0.004 -0.005 -0.001 -0.028 -0.003 -0.024 -0.026 0.003 -0.008 -0.012 -0.008
Rs9401: 0 0.001 0.002 0 0 0 0 0 0 0 0 0 0 0.002 0.001 -0.002 -0.002
Rs7327: -0.001 0 -0.003 0.002 -0.001 0 0 0.001 0 0 0 0 0 -0.004 -0.008 -0.015 -0.007
Rs672: 0.002 -0.007 -0.008 -0.006 -0.001 -0.006 0.001 -0.009 0.001 0.001 0 -0.002 -0.012 -0.001 0.001 -0.006 -0.01
Rs4874: 0 0 0 -0.003 -0.003 -0.002 0 0 0 0 0 -0.004 0 0 -0.002 0 -0.001
Rs4255: 0.002 0.001 0.002 -0.005 -0.002 0 0.002 -0.002 0 0.002 -0.007 -0.003 0 0.005 -0.003 0 0.003
Rs4247: 0.001 0 0.001 0 0.003 0 0 0.001 0 0 0.004 -0.012 0.002 -0.002 -0.006 0.001 0.002
Rs4237: 0 -0.001 0 0.012 -0.001 0 0 -0.004 0.001 0 0.004 0.004 -0.004 0.005 -0.007 0 0
Rs4231: 0.003 0.002 0 0 0 0 -0.003 0 0 0 0 -0.021 -0.001 -0.013 -0.003 0 -0.004
Rs4081: 0 0 0 0 0 0 0 -0.001 -0.002 -0.017 0 -0.005 0.004 -0.001 0.001 0 -0.007
Rs3367: 0 0 0 0 0 0 0 -0.001 -0.001 0 0 0 0 0 0 0 -0.002
RfYNLF_34C: 0 0 -0.004 0 0 0 0.001 0 0 0 0 0 0 -0.001 0 0 0.002
RfYNLF_31C: 0.001 0 0 0 0 0 0 0 0 0 0 0 0 0 0.002 0 0
Rf4092: -0.011 0.007 -0.001 -0.005 -0.001 0.001 -0.002 0 -0.004 -0.02 -0.197 -0.064 -0.005 -0.008 -0.002 -0.006 -0.003
bRp3: -0.007 -0.006 -0.006 0.001 0.001 -0.004 -0.004 0 0.003 -0.019 -0.009 -0.035 -0.011 -0.005 -0.008 -0.028 0.013
As6526: 0 0 0.006 0 -0.002 0 0.002 0 0.002 0 -0.01 -0.028 -0.001 -0.001 0 0 0
-0.10 -0.05 0.00 0.05 0.10

Figure S19. Graphical representation of difference in Codon Adaptation Index ($\Delta$CAI) across the segments and for all external branches of the local tree. (A) SARS-CoV-1, (B) MERS and (C) SARS-CoV-2.

(A)

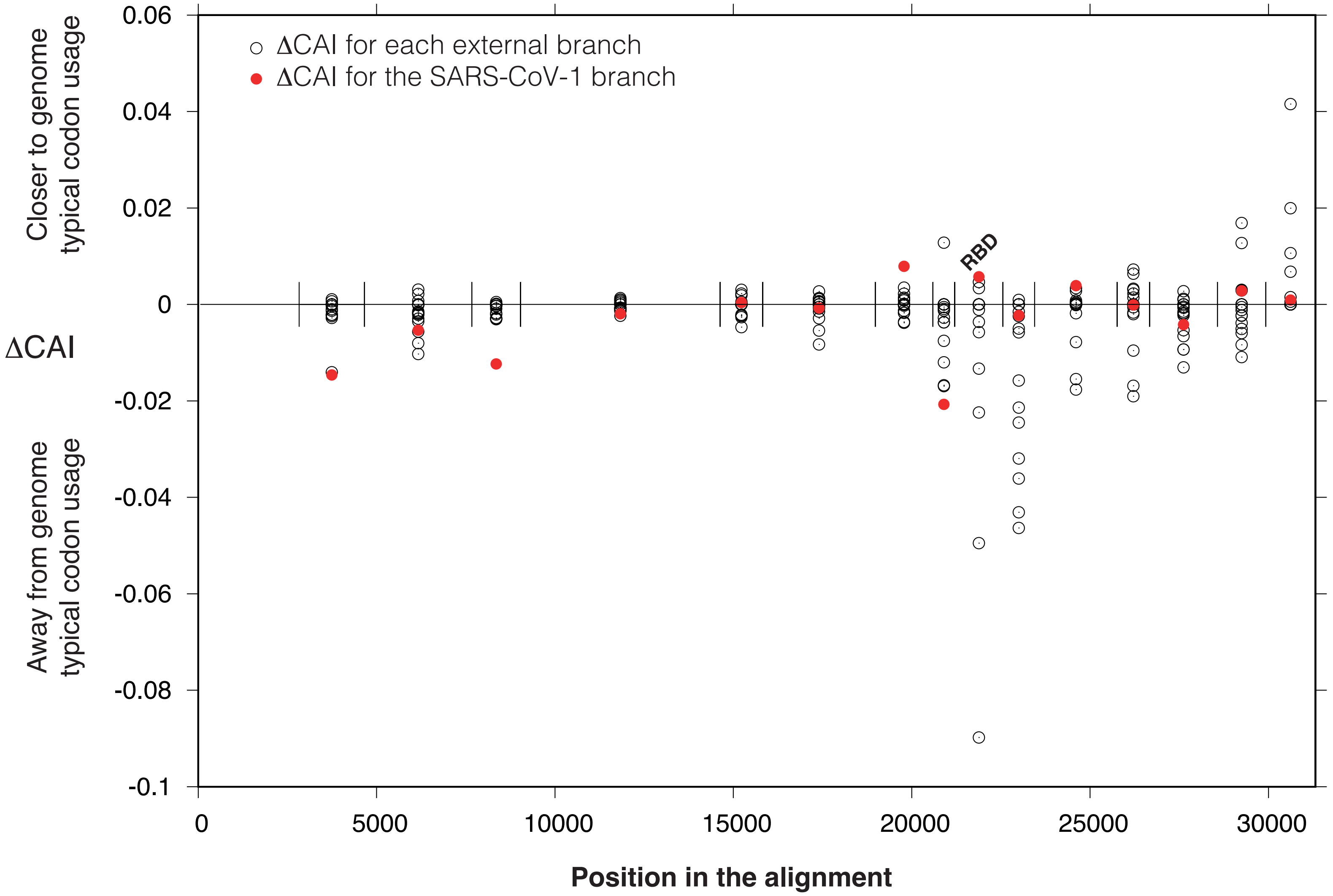

ΔCAI for each external branch
ΔCAI for the SARS-CoV-1 branch
Closer to genome typical codon usage
ΔCAI
Away from genome typical codon usage
RBD
0.06
0.04
0.02
0
-0.02
-0.04
-0.06
-0.08
-0.1
0
5000
10000
15000
20000
25000
30000
Position in the alignment

(B)

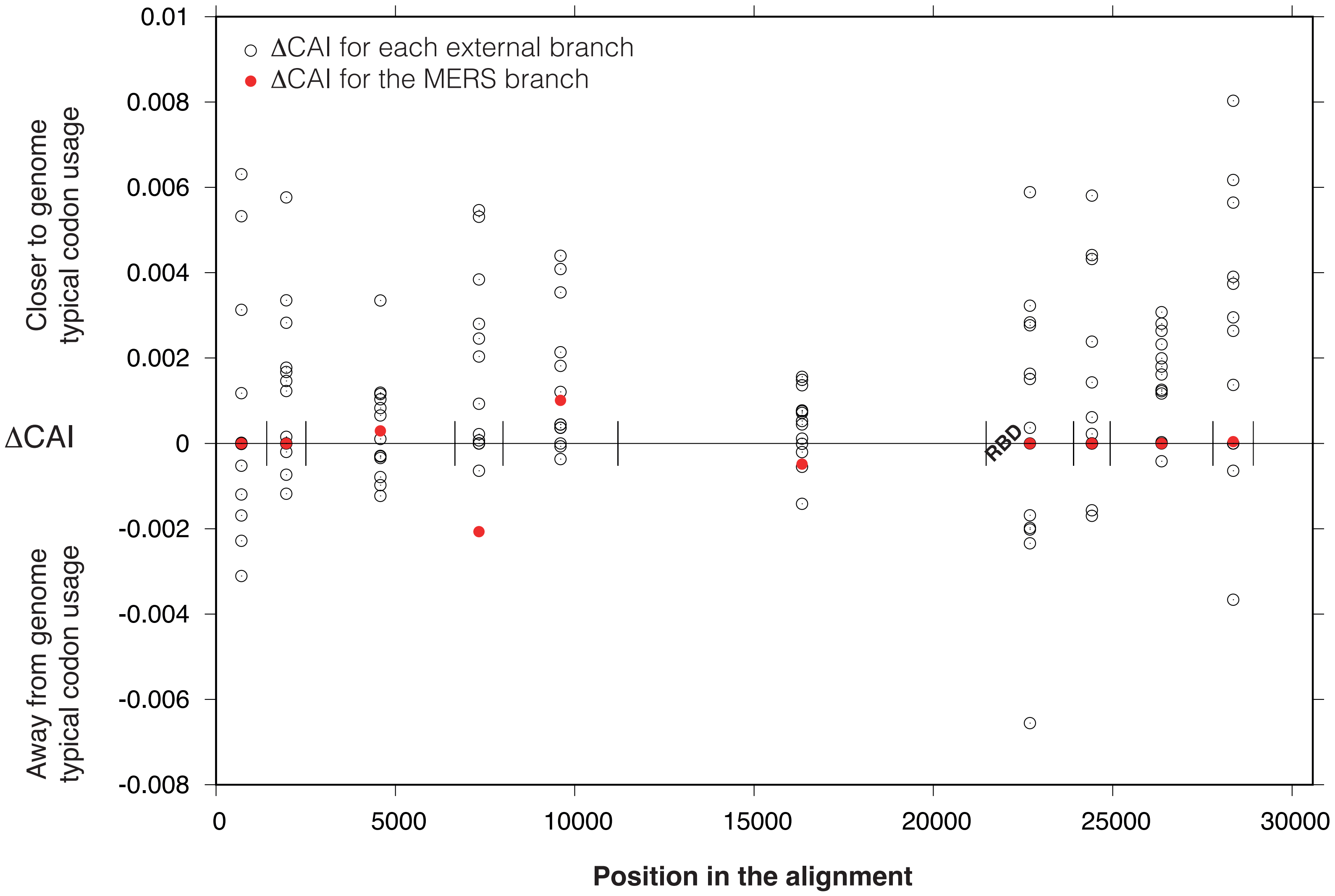

0.01
0.008
0.006
0.004
0.002
0
-0.002
-0.004
-0.006
-0.008
ΔCAI for each external branch
ΔCAI for the MERS branch
Closer to genome typical codon usage
ΔCAI
Away from genome typical codon usage
RBD
0
5000
10000
15000
20000
25000
30000
Position in the alignment

(C)

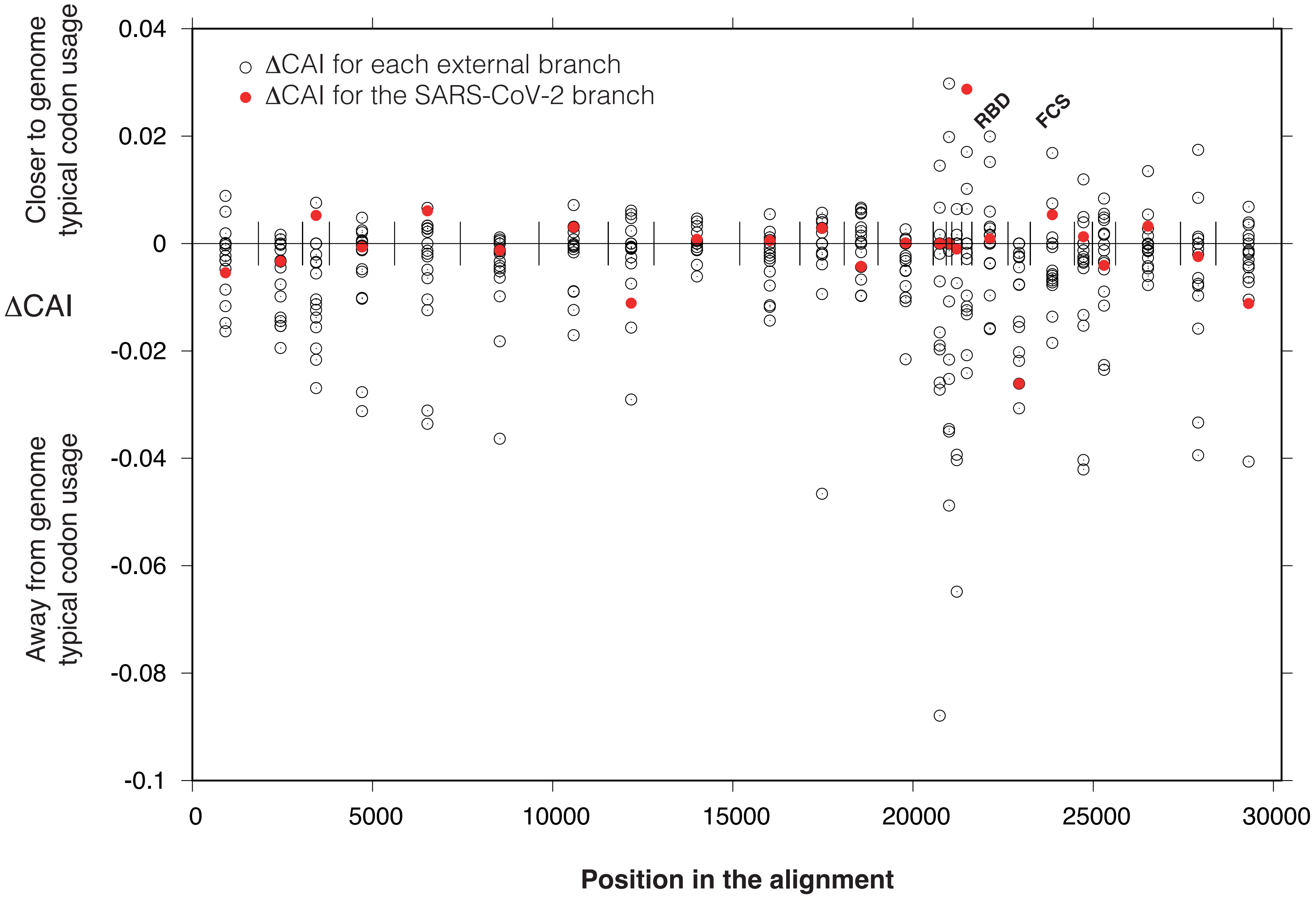

ΔCAI for each external branch
ΔCAI for the SARS-CoV-2 branch
RBD
FCS
Closer to genome typical codon usage
ΔCAI
Away from genome typical codon usage
0.04
0.02
0
-0.02
-0.04
-0.06
-0.08
-0.1
0
5000
10000
15000
20000
25000
30000
Position in the alignment

Figure S20. Estimation of the number of insertions across all the segments of the three datasets. Grey: 0; light orange: 1; dark brown: 2 or more. (A) SARS-CoV-1, (B) MERS and (C) SARS-CoV-2.

(A)

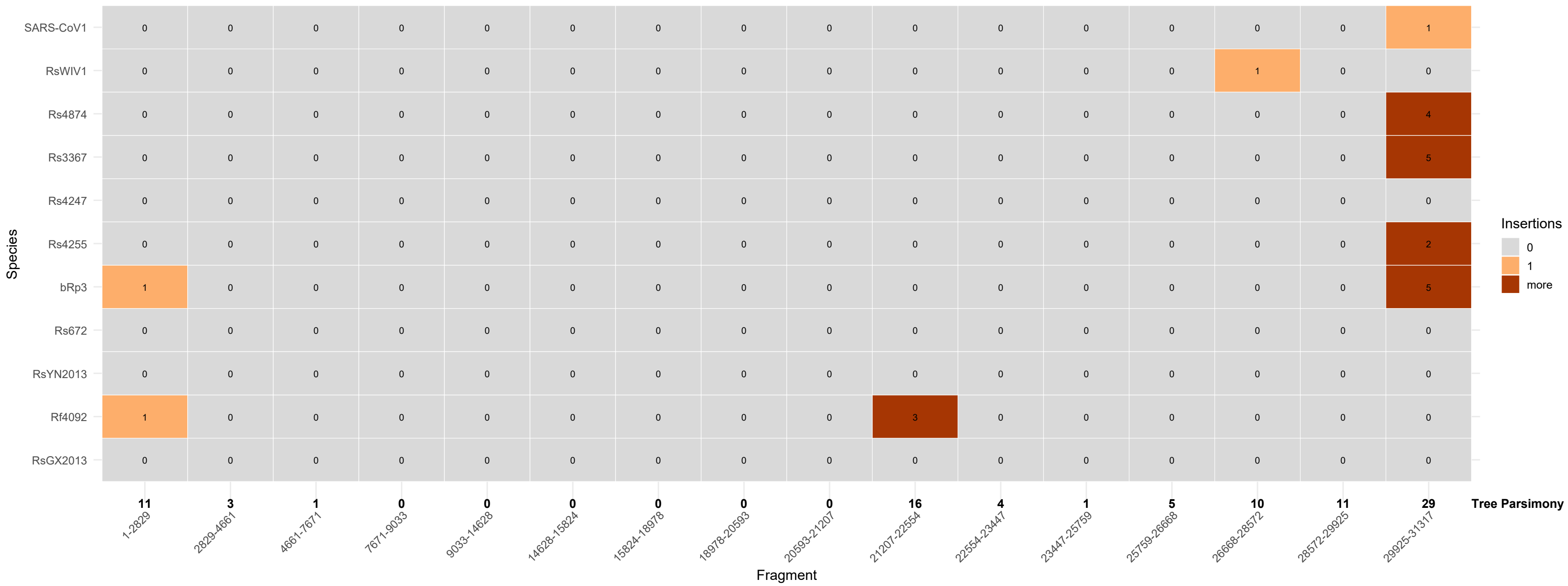
Species
SARS-CoV1
RsWIV1
Rs4874
Rs3367
Rs4247
Rs4255
bRp3
Rs672
RsYN2013
Rf4092
RsGX2013
Insertions
0
1
more
11
3
1
0
0
0
0
0
0
16
4
1
5
10
11
29
Tree Parsimony
1-2829
2829-4661
4661-7671
7671-9033
9033-14628
14628-15824
15824-18978
18978-20593
20593-21207
21207-22554
22554-23447
23447-25759
25759-26668
26668-28572
28572-29925
29925-31317
Fragment

(B)

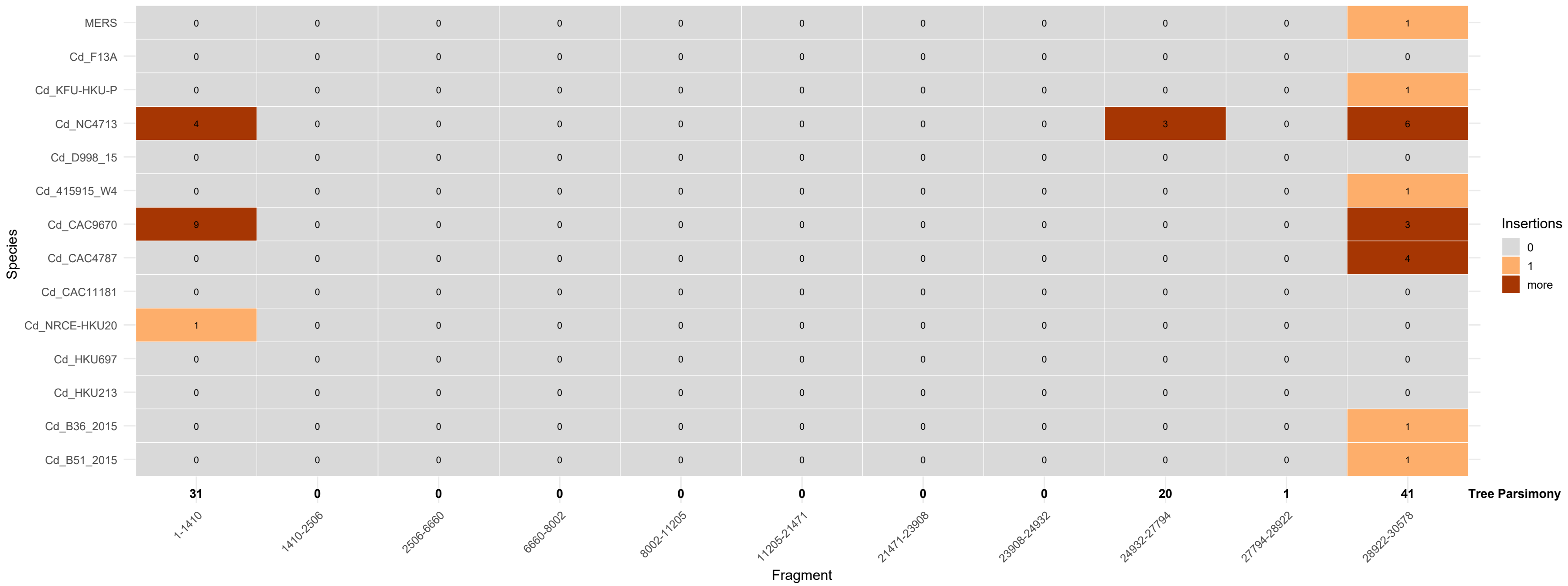

Species
MERS 0 0 0 0 0 0 0 0 0 0 1
Cd_F13A 0 0 0 0 0 0 0 0 0 0 0
Cd_KFU-HKU-P 0 0 0 0 0 0 0 0 0 0 1
Cd_NC4713 4 0 0 0 0 0 0 0 3 0 6
Cd_D998_15 0 0 0 0 0 0 0 0 0 0 0
Cd_415915_W4 0 0 0 0 0 0 0 0 0 0 1
Cd_CAC9670 9 0 0 0 0 0 0 0 0 0 3
Cd_CAC4787 0 0 0 0 0 0 0 0 0 0 4
Cd_CAC11181 0 0 0 0 0 0 0 0 0 0 0
Cd_NRCE-HKU20 1 0 0 0 0 0 0 0 0 0 0
Cd_HKU697 0 0 0 0 0 0 0 0 0 0 0
Cd_HKU213 0 0 0 0 0 0 0 0 0 0 0
Cd_B36_2015 0 0 0 0 0 0 0 0 0 0 1
Cd_B51_2015 0 0 0 0 0 0 0 0 0 0 1
31 0 0 0 0 0 0 0 20 1 41 Tree Parsimony
1-1410
1410-2506
2506-6660
6660-8002
8002-11205
11205-21471
21471-23908
23908-24932
24932-27794
27794-28922
28922-30578
Fragment
Insertions
0
1
more

(C)

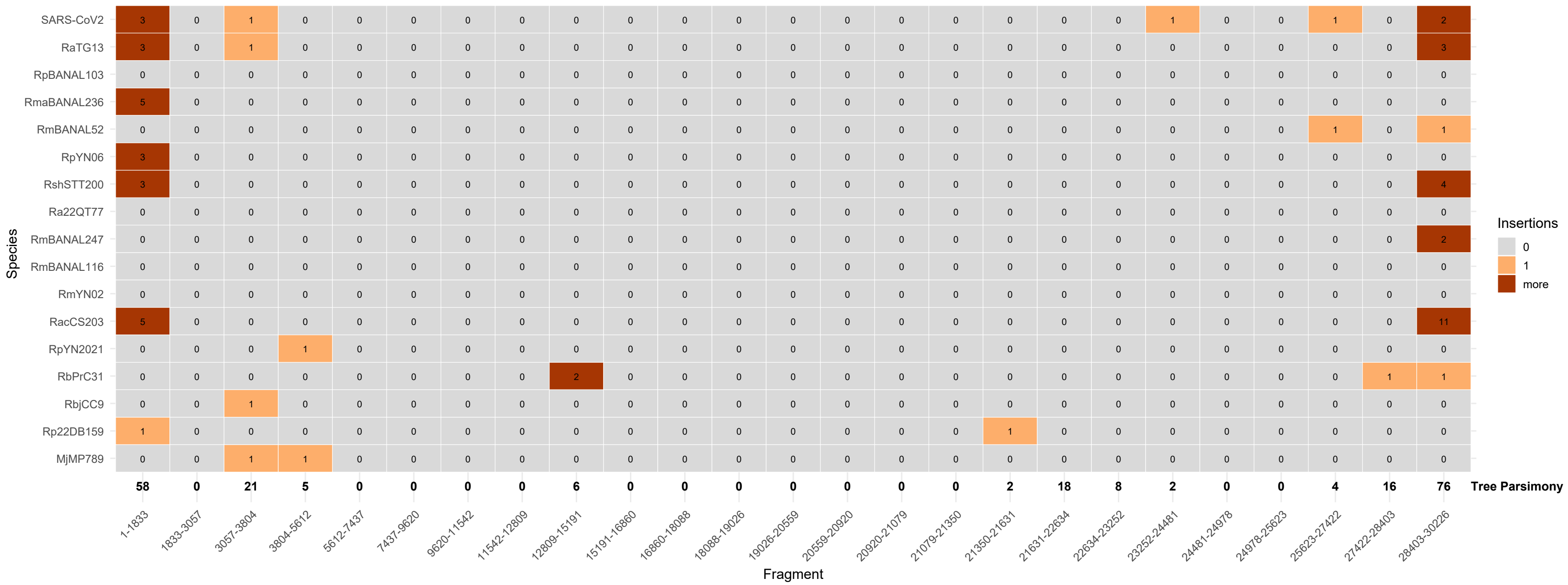

Insertions
0
1
more
Species
SARS-CoV2 3 0 1 0 0 0 0 0 0 0 0 0 0 0 0 0 0 0 0 1 0 0 1 0 2
RaTG13 3 0 1 0 0 0 0 0 0 0 0 0 0 0 0 0 0 0 0 0 0 0 0 0 3
RpBANAL103 0 0 0 0 0 0 0 0 0 0 0 0 0 0 0 0 0 0 0 0 0 0 0 0 0
RmaBANAL236 5 0 0 0 0 0 0 0 0 0 0 0 0 0 0 0 0 0 0 0 0 0 0 0 0
RmBANAL52 0 0 0 0 0 0 0 0 0 0 0 0 0 0 0 0 0 0 0 0 0 0 1 0 1
RpYN06 3 0 0 0 0 0 0 0 0 0 0 0 0 0 0 0 0 0 0 0 0 0 0 0 0
RshSTT200 3 0 0 0 0 0 0 0 0 0 0 0 0 0 0 0 0 0 0 0 0 0 0 0 4
Ra22QT77 0 0 0 0 0 0 0 0 0 0 0 0 0 0 0 0 0 0 0 0 0 0 0 0 0
RmBANAL247 0 0 0 0 0 0 0 0 0 0 0 0 0 0 0 0 0 0 0 0 0 0 0 0 2
RmBANAL116 0 0 0 0 0 0 0 0 0 0 0 0 0 0 0 0 0 0 0 0 0 0 0 0 0
RmYN02 0 0 0 0 0 0 0 0 0 0 0 0 0 0 0 0 0 0 0 0 0 0 0 0 0
RacCS203 5 0 0 0 0 0 0 0 0 0 0 0 0 0 0 0 0 0 0 0 0 0 0 0 11
RpYN2021 0 0 0 1 0 0 0 0 0 0 0 0 0 0 0 0 0 0 0 0 0 0 0 0 0
RbPrC31 0 0 0 0 0 0 0 0 2 0 0 0 0 0 0 0 0 0 0 0 0 0 0 1 1
RbjCC9 0 0 1 0 0 0 0 0 0 0 0 0 0 0 0 0 0 0 0 0 0 0 0 0 0
Rp22DB159 1 0 0 0 0 0 0 0 0 0 0 0 0 0 0 0 1 0 0 0 0 0 0 0 0
MjMP789 0 0 1 1 0 0 0 0 0 0 0 0 0 0 0 0 0 0 0 0 0 0 0 0 0
58 0 21 5 0 0 0 0 6 0 0 0 0 0 0 0 2 18 8 2 0 0 4 16 76 Tree Parsimony
1-1833 1833-3057 3057-3804 3804-5612 5612-7437 7437-9620 9620-11542 11542-12809 12809-15191 15191-16860 16860-18088 18088-19026 19026-20559 20559-20920 20920-21079 21079-21350 21350-21631 21631-22634 22634-23252 23252-24481 24481-24978 24978-25623 25623-27422 27422-28403 28403-30226
Fragment

Figure S21. Estimation of the number of deletions across all the segments of the three datasets. Grey: 0; light blue: 1; dark blue: 2 or more. (A) SARS-CoV-1, (B) MERS and (C) SARS-CoV-2.

(A)

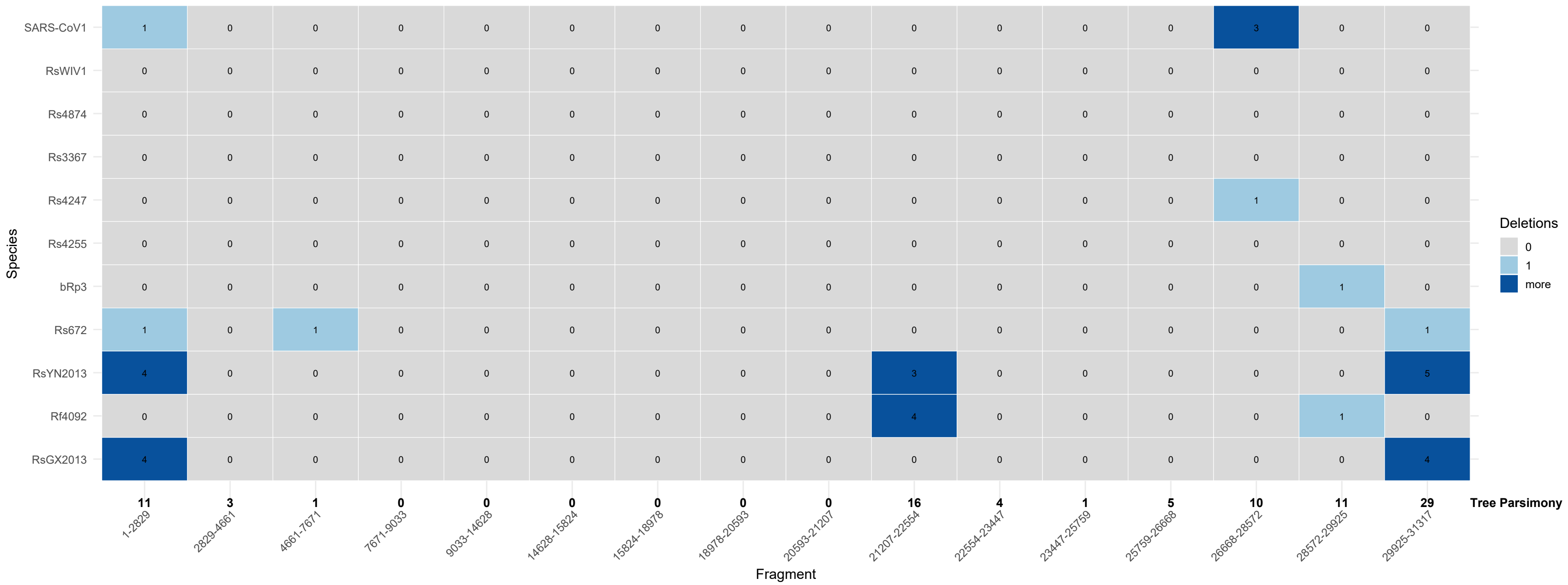
Species
SARS-CoV1
RsWIV1
Rs4874
Rs3367
Rs4247
Rs4255
bRp3
Rs672
RsYN2013
Rf4092
RsGX2013
Deletions
0
1
more
11 3 1 0 0 0 0 0 0 16 4 1 5 10 11 29
Tree Parsimony
1-2829
2829-4661
4661-7671
7671-9033
9033-14628
14628-15824
15824-18978
18978-20593
20593-21207
21207-22554
22554-23447
23447-25759
25759-26668
26668-28572
28572-29925
29925-31317
Fragment

(B)

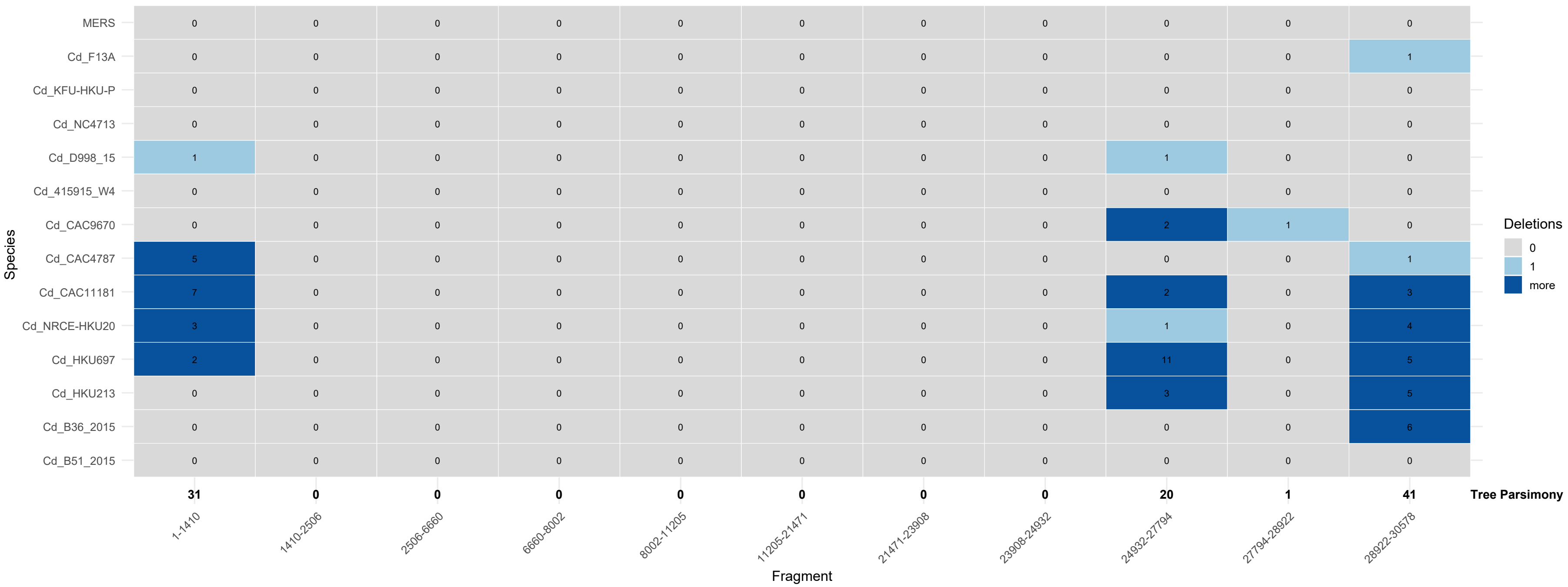
Species
MERS 0 0 0 0 0 0 0 0 0 0 0
Cd_F13A 0 0 0 0 0 0 0 0 0 0 1
Cd_KFU-HKU-P 0 0 0 0 0 0 0 0 0 0 0
Cd_NC4713 0 0 0 0 0 0 0 0 0 0 0
Cd_D998_15 1 0 0 0 0 0 0 0 1 0 0
Cd_415915_W4 0 0 0 0 0 0 0 0 0 0 0
Cd_CAC9670 0 0 0 0 0 0 0 0 2 1 0
Cd_CAC4787 5 0 0 0 0 0 0 0 0 0 1
Cd_CAC11181 7 0 0 0 0 0 0 0 2 0 3
Cd_NRCE-HKU20 3 0 0 0 0 0 0 0 1 0 4
Cd_HKU697 2 0 0 0 0 0 0 0 11 0 5
Cd_HKU213 0 0 0 0 0 0 0 0 3 0 5
Cd_B36_2015 0 0 0 0 0 0 0 0 0 0 6
Cd_B51_2015 0 0 0 0 0 0 0 0 0 0 0
31 0 0 0 0 0 0 0 20 1 41
Tree Parsimony
1-1410
1410-2506
2506-6660
6660-8002
8002-11205
11205-21471
21471-23908
23908-24932
24932-27794
27794-28922
28922-30578
Fragment
Deletions
0
1
more

(C)

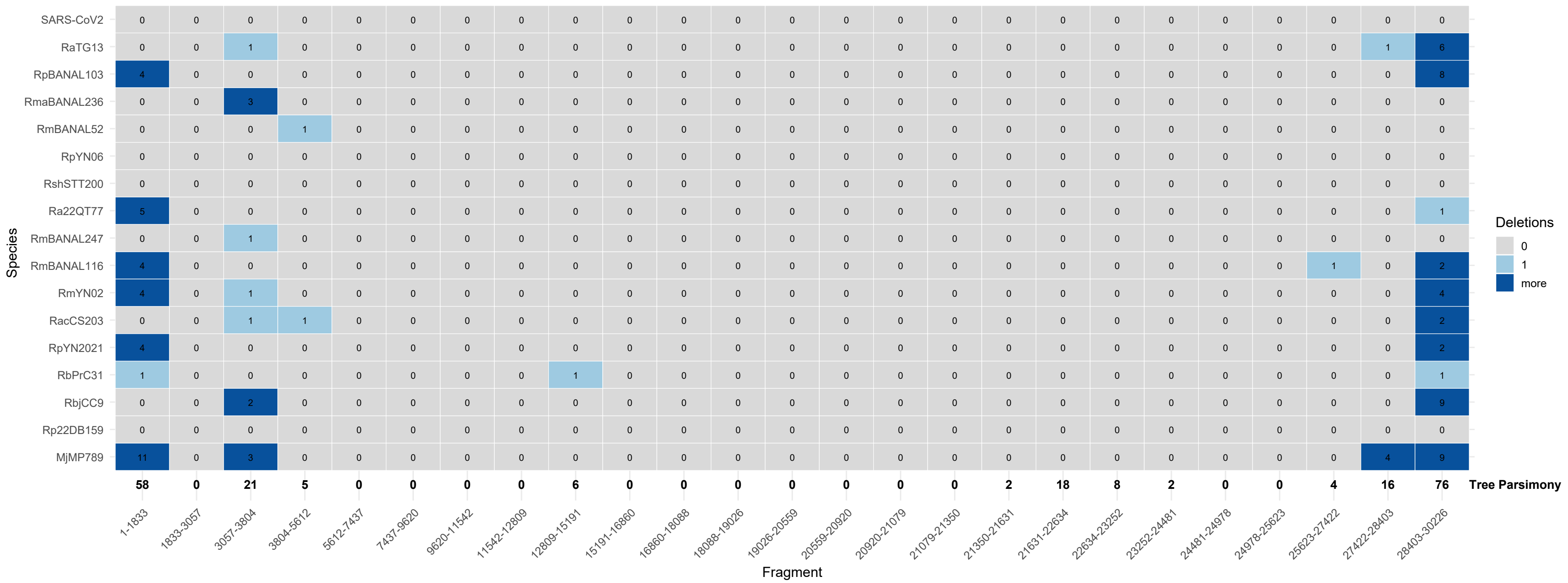
Species
SARS-CoV2 0 0 0 0 0 0 0 0 0 0 0 0 0 0 0 0 0 0 0 0 0 0 0 0 0
RaTG13 0 0 1 0 0 0 0 0 0 0 0 0 0 0 0 0 0 0 0 0 0 0 0 1 6
RpBANAL103 4 0 0 0 0 0 0 0 0 0 0 0 0 0 0 0 0 0 0 0 0 0 0 0 8
RmaBANAL236 0 0 3 0 0 0 0 0 0 0 0 0 0 0 0 0 0 0 0 0 0 0 0 0 0
RmBANAL52 0 0 0 1 0 0 0 0 0 0 0 0 0 0 0 0 0 0 0 0 0 0 0 0 0
RpYN06 0 0 0 0 0 0 0 0 0 0 0 0 0 0 0 0 0 0 0 0 0 0 0 0 0
RshSTT200 0 0 0 0 0 0 0 0 0 0 0 0 0 0 0 0 0 0 0 0 0 0 0 0 0
Ra22QT77 5 0 0 0 0 0 0 0 0 0 0 0 0 0 0 0 0 0 0 0 0 0 0 0 1
RmBANAL247 0 0 1 0 0 0 0 0 0 0 0 0 0 0 0 0 0 0 0 0 0 0 0 0 0
RmBANAL116 4 0 0 0 0 0 0 0 0 0 0 0 0 0 0 0 0 0 0 0 0 0 1 0 2
RmYN02 4 0 1 0 0 0 0 0 0 0 0 0 0 0 0 0 0 0 0 0 0 0 0 0 4
RacCS203 0 0 1 1 0 0 0 0 0 0 0 0 0 0 0 0 0 0 0 0 0 0 0 0 2
RpYN2021 4 0 0 0 0 0 0 0 0 0 0 0 0 0 0 0 0 0 0 0 0 0 0 0 2
RbPrC31 1 0 0 0 0 0 0 0 1 0 0 0 0 0 0 0 0 0 0 0 0 0 0 0 1
RbjCC9 0 0 2 0 0 0 0 0 0 0 0 0 0 0 0 0 0 0 0 0 0 0 0 0 9
Rp22DB159 0 0 0 0 0 0 0 0 0 0 0 0 0 0 0 0 0 0 0 0 0 0 0 0 0
MjMP789 11 0 3 0 0 0 0 0 0 0 0 0 0 0 0 0 0 0 0 0 0 0 0 4 9
58 0 21 5 0 0 0 0 6 0 0 0 0 0 0 0 2 18 8 2 0 0 4 16 76 Tree Parsimony
1-1833 1833-3057 3057-3804 3804-5612 5612-7437 7437-9620 9620-11542 11542-12809 12809-15191 15191-16860 16860-18088 18088-19026 19026-20559 20559-20920 20920-21079 21079-21350 21350-21631 21631-22634 22634-23252 23252-24481 24481-24978 24978-25623 25623-27422 27422-28403 28403-30226
Fragment
Deletions
0
1
more

Figure S22. Graphical representation of insertion and deletion numbers across the segments. For each segment, insertions in the external branches are displayed as positive values, whereas the deletions are indicated in the grey area under the $x$-axis. The total number of events (insertions + deletions) inferred in the local tree of each segments is represented by black triangles connected by a straight line. Values equal to 0 are not represented, except for the total number of events. (A) SARS-CoV-1, (B) MERS and (C) SARS-CoV-2. Key actors of the viral infectivity are labeled: Receptor Binding Domain (RBD) of the three viruses and Furin Cleavage Site (FCS) of the SARS-CoV-2.

(A)

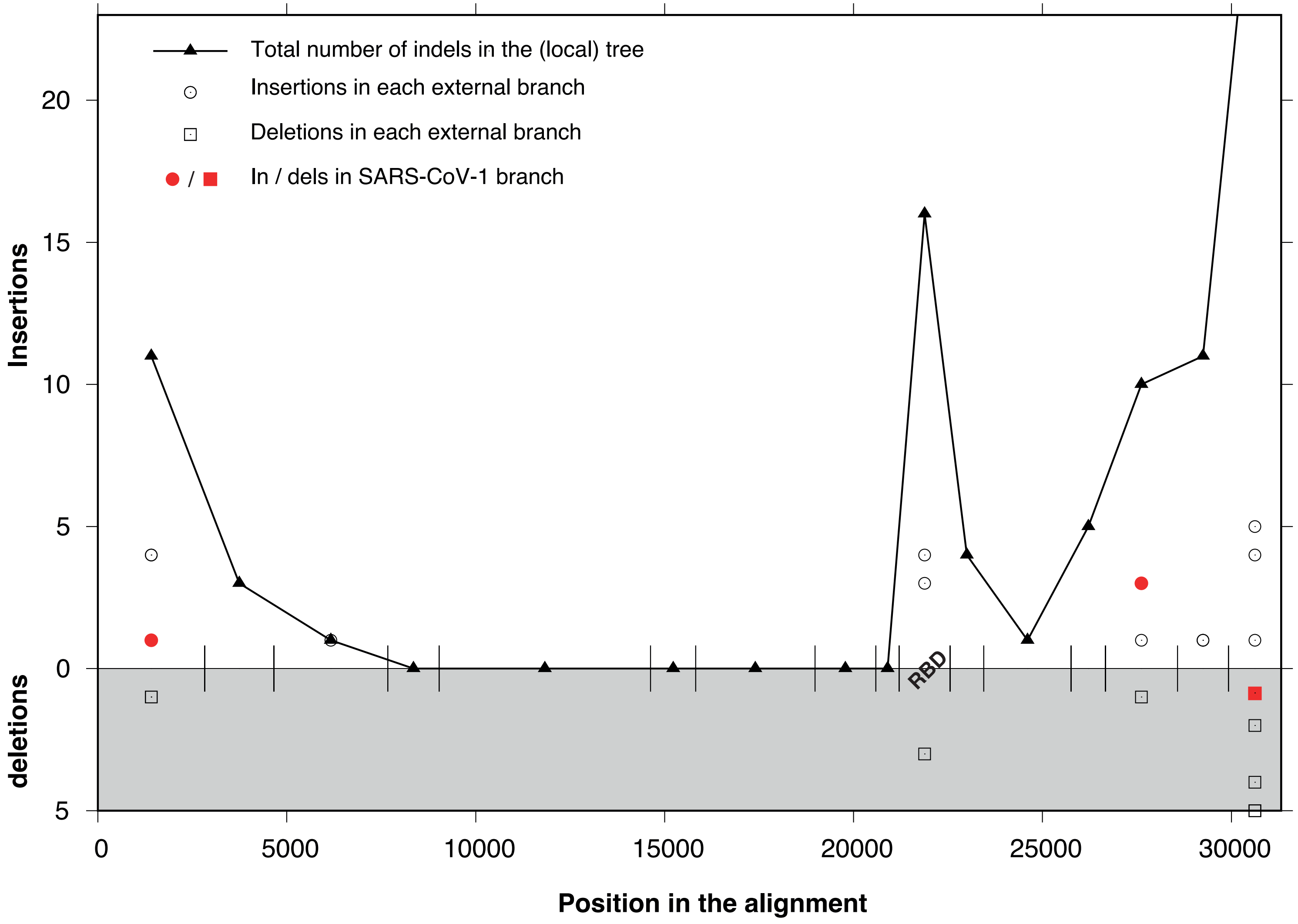
Total number of indels in the (local) tree
Insertions in each external branch
Deletions in each external branch
In / dels in SARS-CoV-1 branch
Insertions
deletions
20
15
10
5
0
5
RBD
0
5000
10000
15000
20000
25000
30000
Position in the alignment

(B)

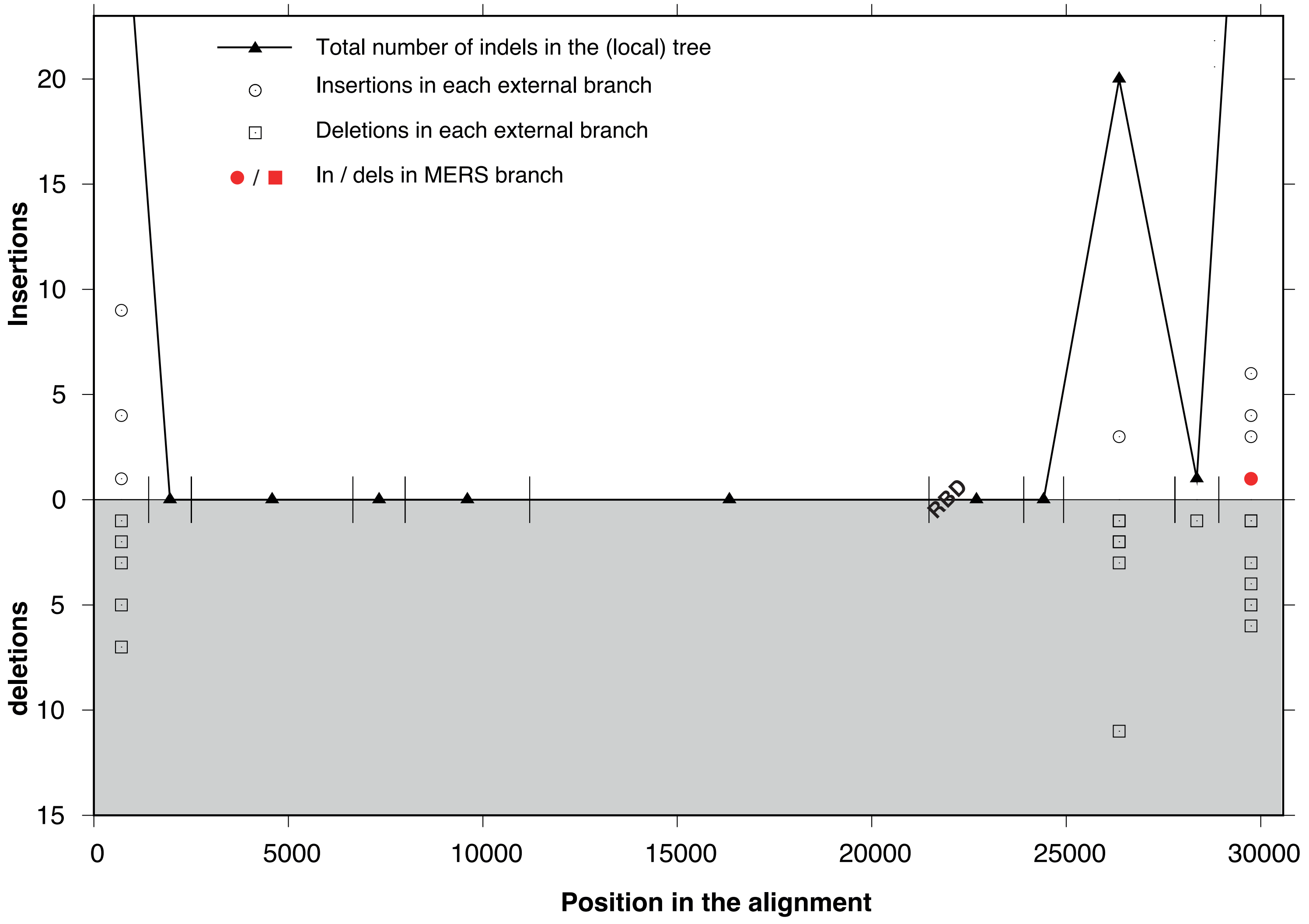

Total number of indels in the (local) tree
Insertions in each external branch
Deletions in each external branch
In / dels in MERS branch
Insertions
deletions
Position in the alignment
RBD
20
15
10
5
0
5
10
15
0
5000
10000
15000
20000
25000
30000

(C)

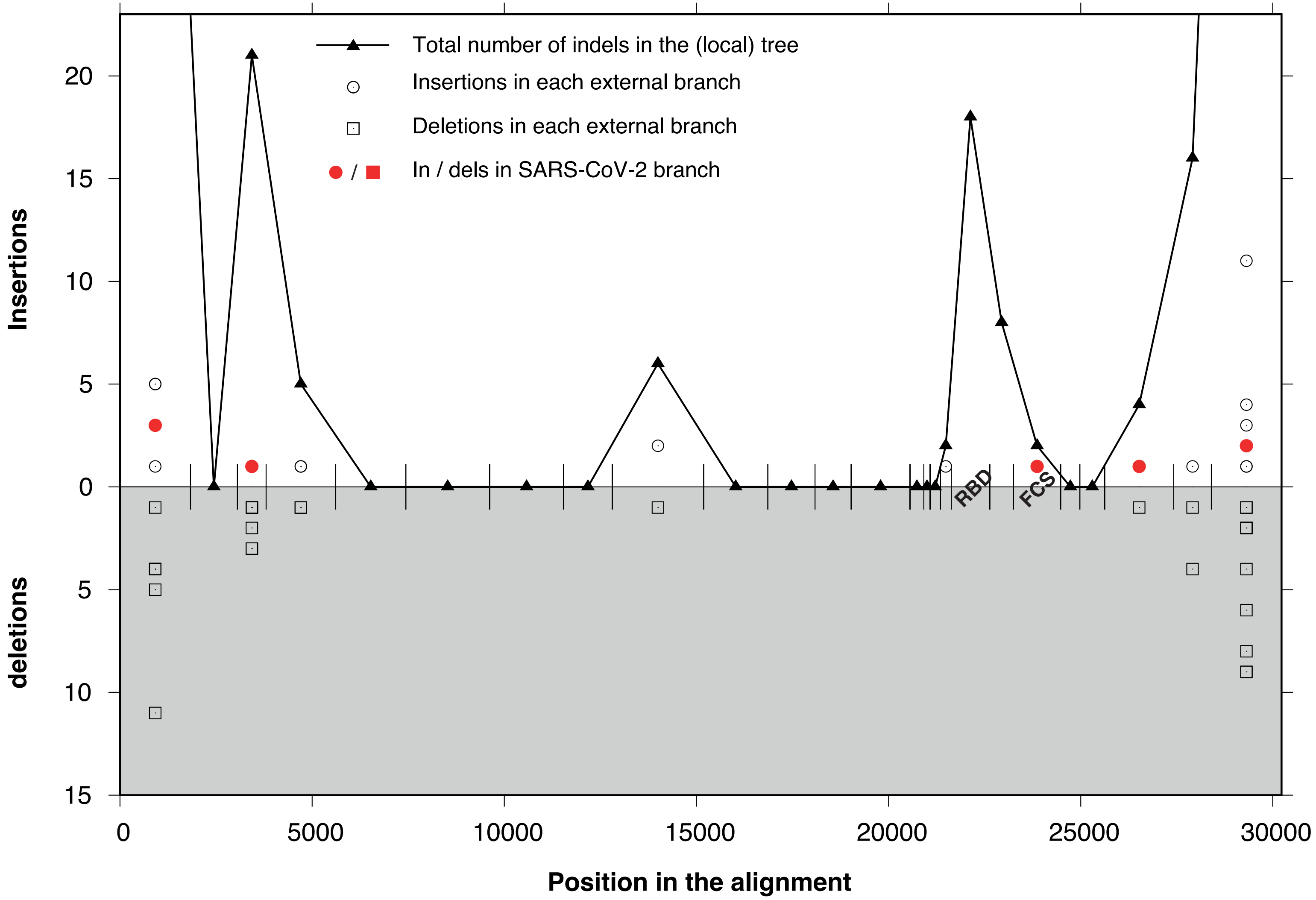

Total number of indels in the (local) tree
Insertions in each external branch
Deletions in each external branch
In / dels in SARS-CoV-2 branch
Insertions
deletions
20
15
10
5
0
5
10
15
0
5000
10000
15000
20000
25000
30000
RBD
FCS
Position in the alignment
